## Supplementary Information Mesmerize Documentation for "Mesmerize: a dynamically adaptable user-friendly analysis platform for 2D & 3D calcium imaging data"

***Release 0.6.1***

**Kushal Kolar**

**May 23, 2021**

### OVERVIEW


1 New: Video Tutorials! 3

2 Indices and tables 235

Python Module Index 237

Index 239

biorxiv: https://doi.org/10.1101/840488
GitHub: https://github.com/kushalkolar/MESmerize
Questions/Discussion: Gitter room
Contact:

CHAPTER

ONE

### NEW: VIDEO TUTORIALS!

**The Main Overview Tutorial playlist provides a quick overview that takes you from raw imaging data, to downstream analysis and interactive visualizations:**

**Additional tutorials on other aspects of Mesmerize will be placed in this playlist:** https://www.youtube.com/playlist?list=PLgofWiw2s4RF_RkGRUfflcj5k5KUTG3o_

#### 1.1 Overview

Mesmerize is a platform for the annotation and analysis of neuronal calcium imaging data. It encompasses the entire process of calcium imaging analysis from raw data to semi-final publication figures that are interactive, and aids in the creation of FAIR-functionally linked datasets. It is applicable for a broad range of experiments and is intended to be used by users with and without a programming background.

##### 1.1.1 Welcome Window

Share your analysis pipelines and figures along with your publication

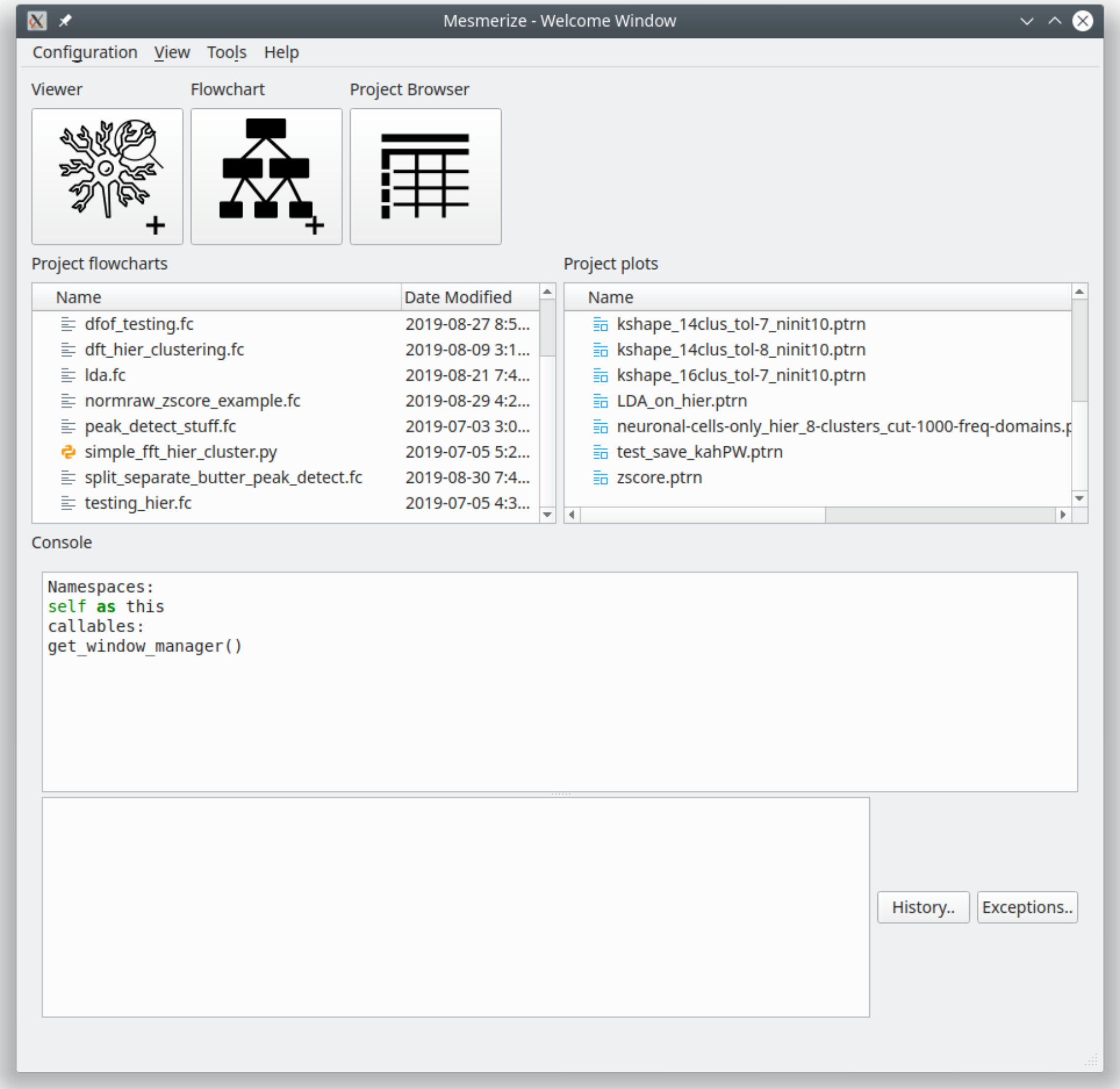


#### 1.1.2 The Viewer

Explore image sequences, and use various modules for pre-processing and signal-extraction. Annotate regions of interest with any relevant information. Map stimuli/behavior periods.

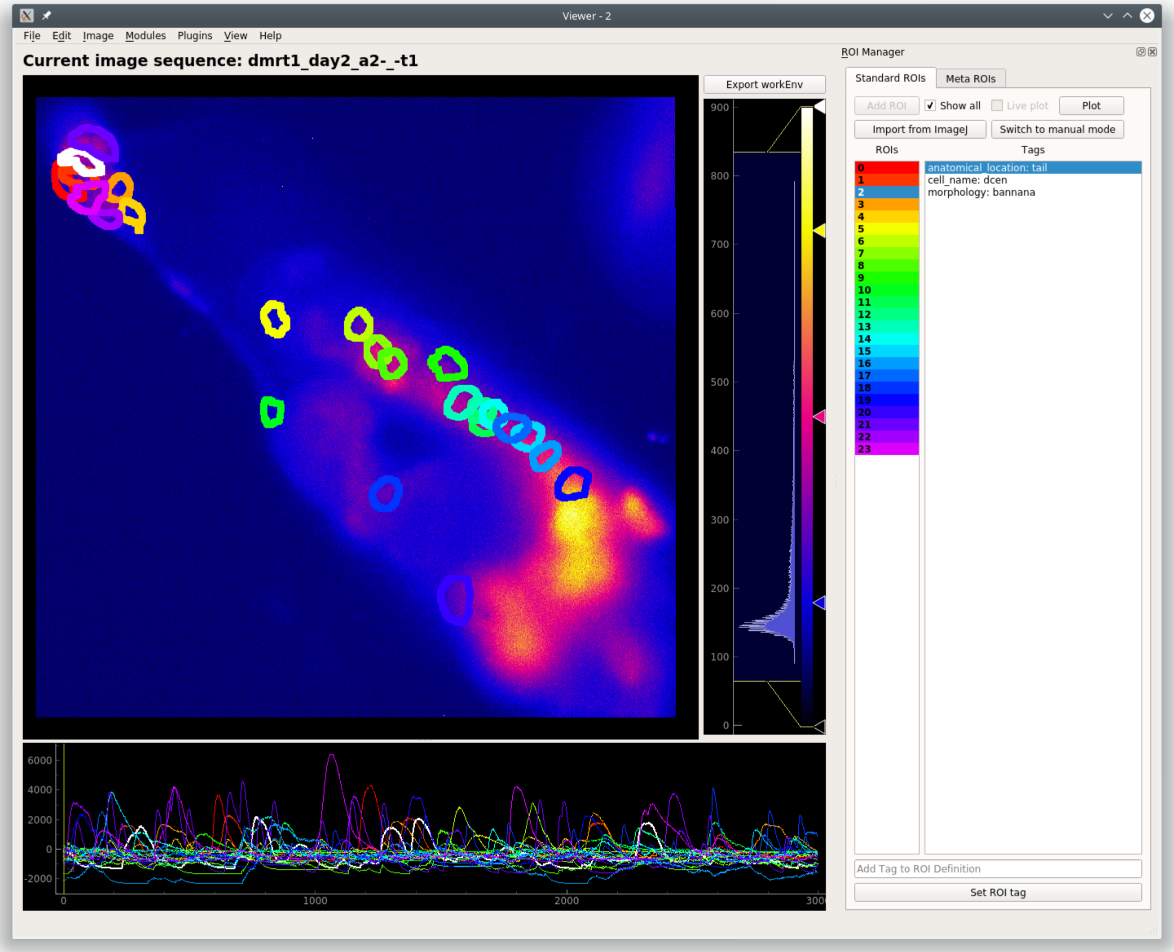


#### 1.1.3 CaImAn modules

Mesmerize contains front-end GUI modules for the CaImAn library. This makes it very easy for users without a programming background to use the library.

**CaImAn Elastic Motion Correction**

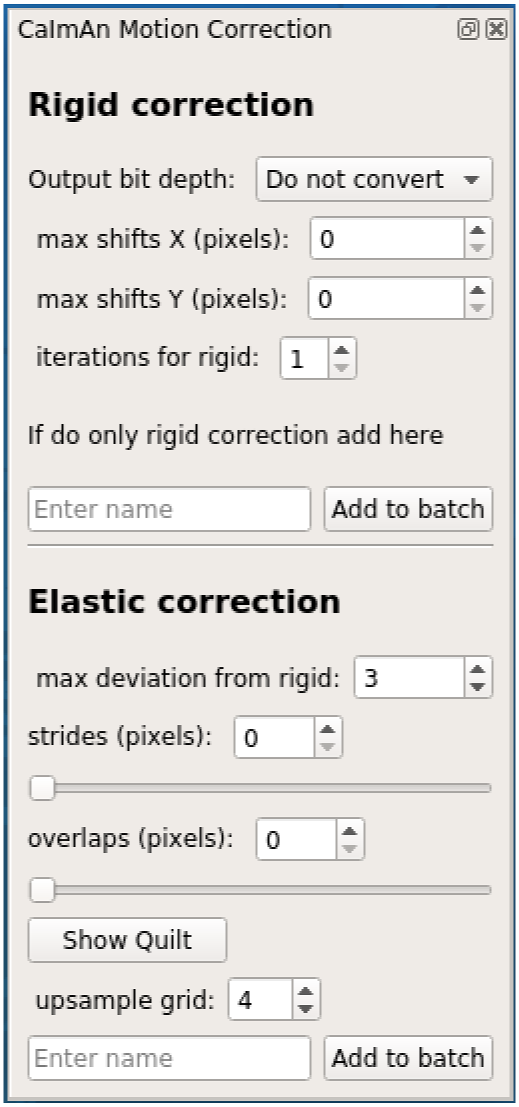
CaImAn Motion Correction
Rigid correction
Output bit depth:
Do not convert
max shifts X (pixels):
0
max shifts Y (pixels):
0
iterations for rigid:
1
If do only rigid correction add here
Enter name
Add to batch
Elastic correction
max deviation from rigid:
3
strides (pixels):
0
overlaps (pixels):
0
Show Quilt
upsample grid:
4
Enter name
Add to batch

**CNMFE**

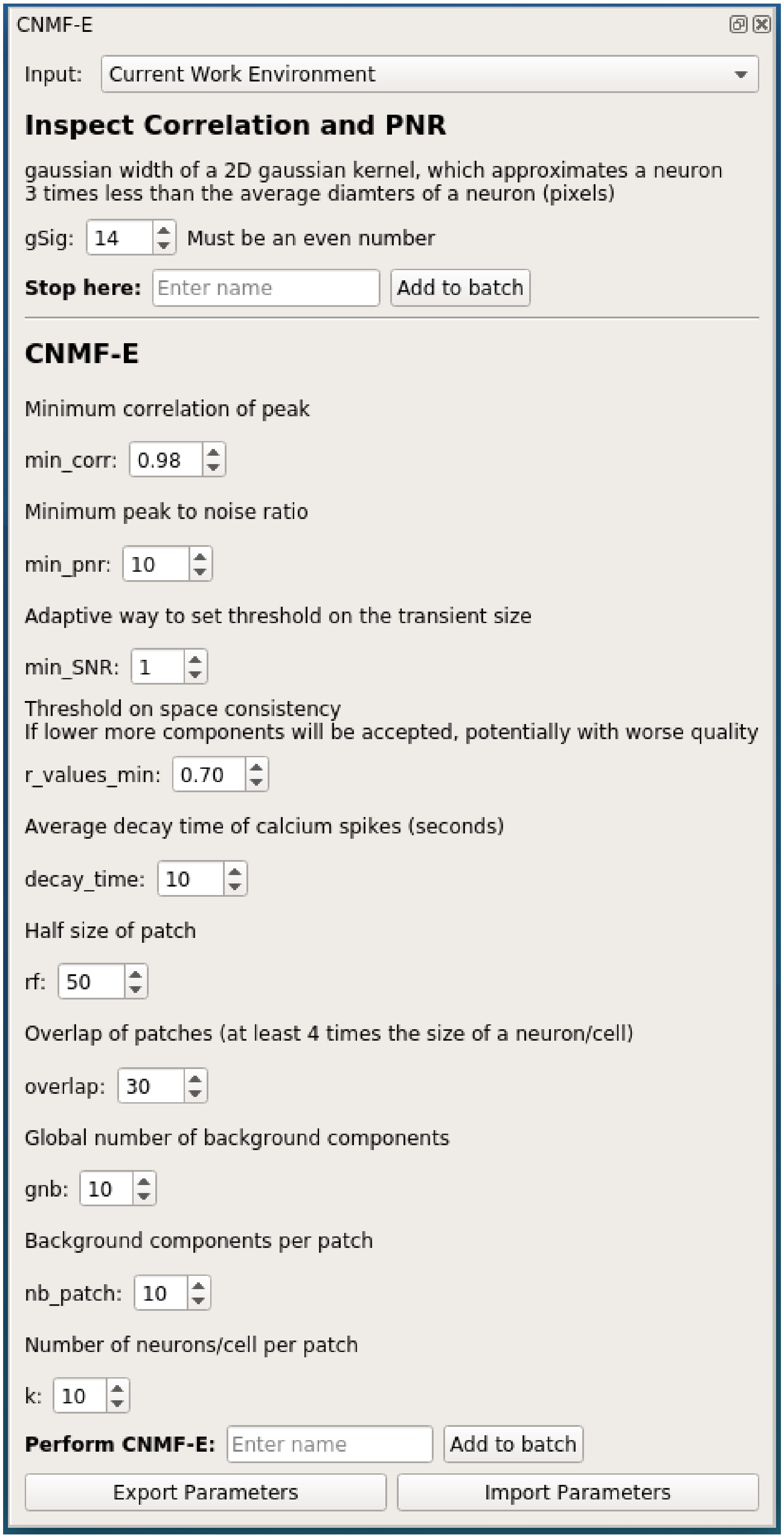
CNMF-E
Input: Current Work Environment
Inspect Correlation and PNR
gaussian width of a 2D gaussian kernel, which approximates a neuron
3 times less than the average diamters of a neuron (pixels)
gSig: 14
Must be an even number
Stop here:
Enter name
Add to batch
CNMF-E
Minimum correlation of peak
min_corr: 0.98
Minimum peak to noise ratio
min_pnr: 10
Adaptive way to set threshold on the transient size
min_SNR: 1
Threshold on space consistency
If lower more components will be accepted, potentially with worse quality
r_values_min: 0.70
Average decay time of calcium spikes (seconds)
decay_time: 10
Half size of patch
rf: 50
Overlap of patches (at least 4 times the size of a neuron/cell)
overlap: 30
Global number of background components
gnb: 10
Background components per patch
nb_patch: 10
Number of neurons/cell per patch
k: 10
Perform CNMF-E:
Enter name
Add to batch
Export Parameters
Import Parameters

##### 1.1.4 Batch Manager

Computationally intense procedures performed can be organized with the Mesmerize Batch Manager.

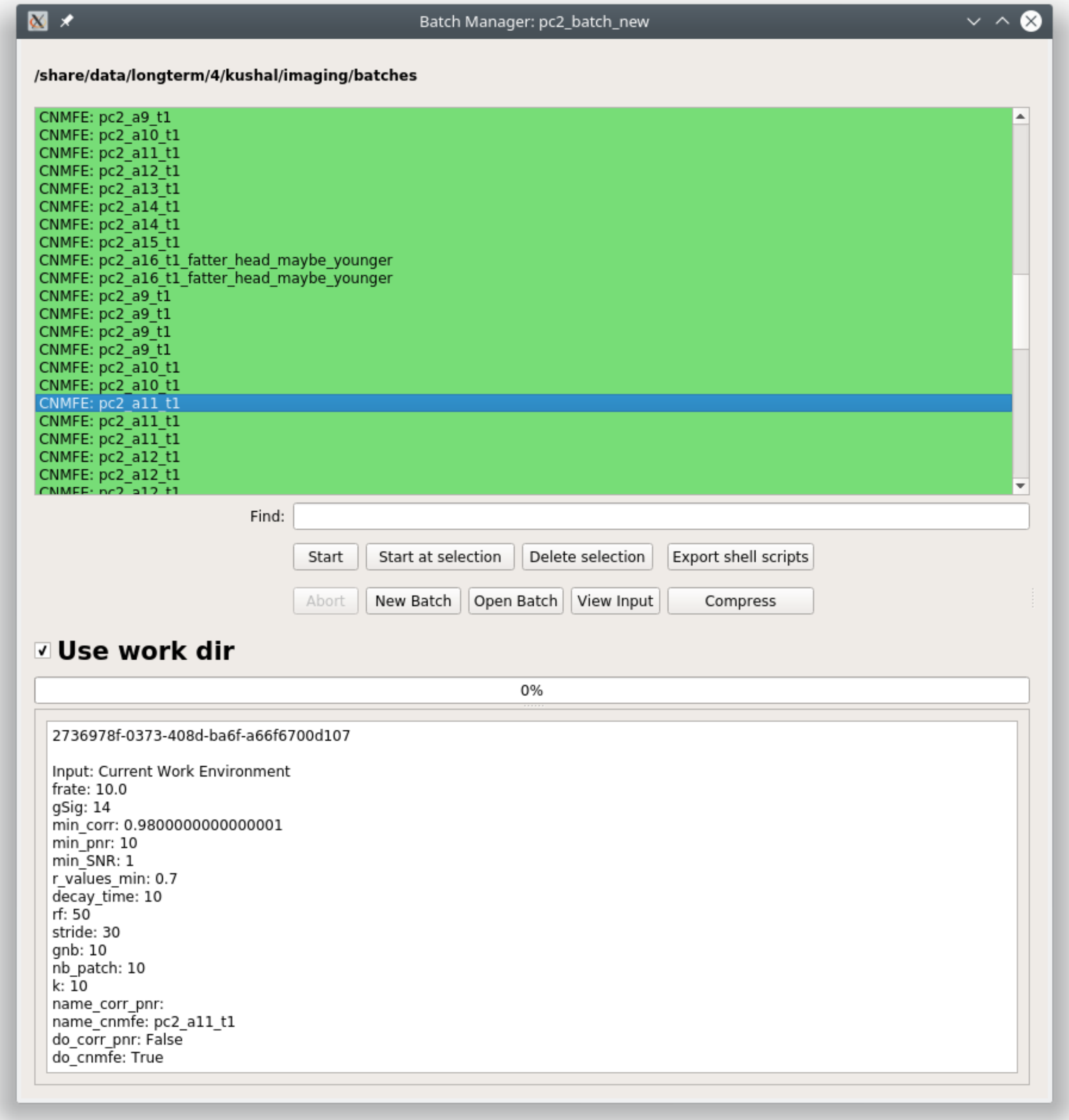

##### 1.1.5 Project Organization

Explore project data and create experimental groups.

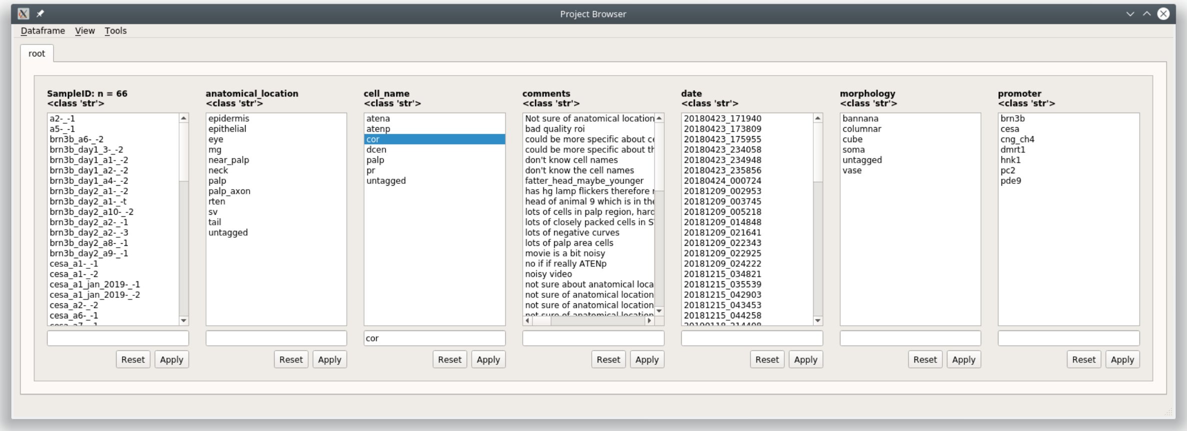


##### 1.1.6 Data analysis - pyqtgraph programmable flowcharts.

Build your own analysis piplines using flowcharts.

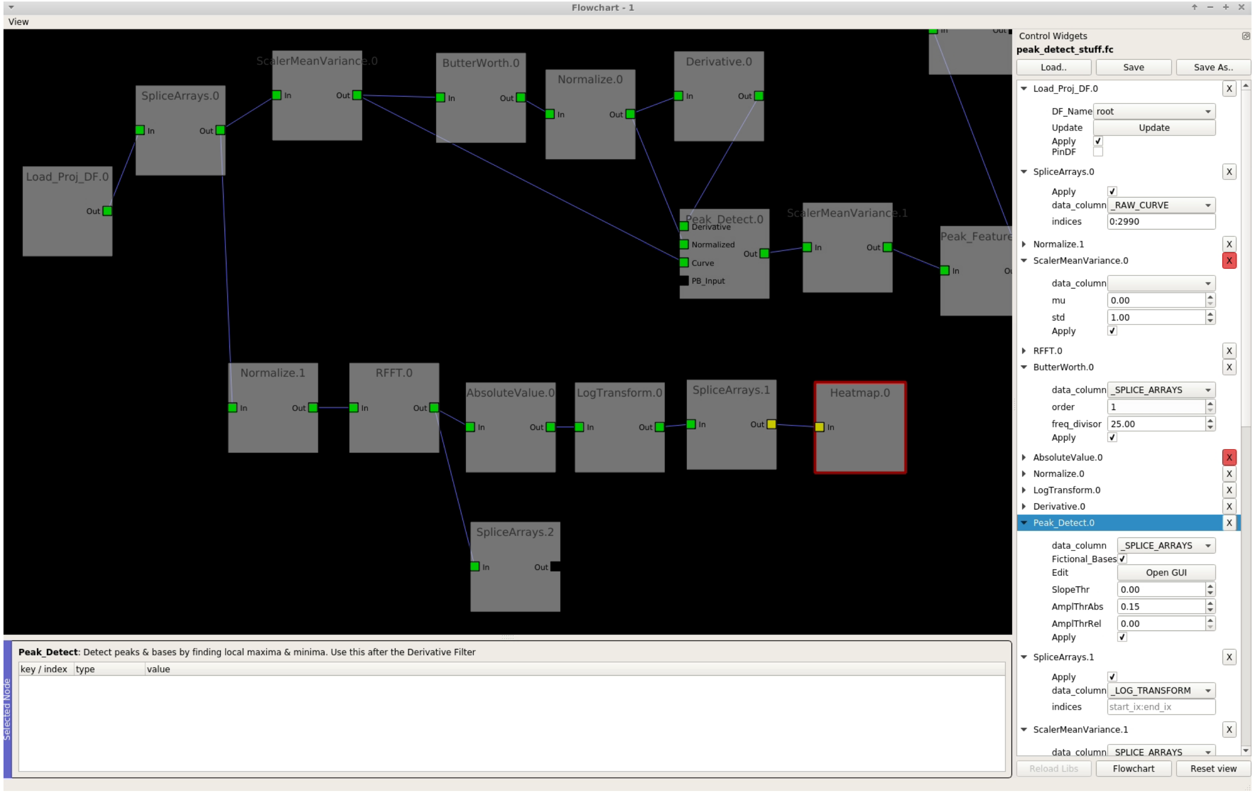

##### 1.1.7 Interactive Plots

Create shareable interactive plots where the analysis history of every datapoint is traceable. Explore information associated with a datapoint, such as the spatial localization of its parent ROI and raw data.

**Interactive Heatmaps**

**Interactive Cross-correlation analysis**

Other types of plots: Beeswarm, Violins, KShape, Proportions, Scatter

#### 1.2 Installation

Mesmerize can be installed on Linux, Mac OSX and Windows. On Windows, Mesmerize can be installed in an anaconda environment. For Mac OSX and Linux you may use either virtual environments or conda environments, but we have had much better experience with virtual environments.

##### 1.2.1 All platforms

We provide a ready to use VM with Mesmerize and all features pre-installed. You can run this VM on Windows, Mac OSX, or Linux. This is the easiest way to get started with Mesmerize if you don't want to setup anaconda or virtual environments by yourself. Just install VirtualBox and import the `mesmerize-v060-2-vm.ova` file.

- VirtualBox: https://www.virtualbox.org/wiki/Downloads
- Download the VM file `mesmerize-v060-2-vm.ova` from zenodo: https://zenodo.org/record/4738514

When you start the VM, just double click the mesmerize launcher on the desktop.

- You can setup *Shared Folders* in the settings for the VM to share data between the VM and your host computer.
- You can mount network drives etc. from within the VM.
- Do not delete the `venvs` directory, this will remove the virtual environment for Mesmerize.
- An example batch with a few examples from the caiman sample data is provided at `/home/user/example_batch`.

The details for the user account on the VM are:

username: `user` | password: `password` |

You can use the same password for `sudo`.

By default the VM is set to use 7 threads and 12GB of RAM. You may modify this according to the resources available on your host computer. You generally want to leave 2-4 threads free on your host computer.

If you get the following error when importing the VM you probably don't have enough space on your computer, I recommend importing the VM on a computer that has a few hundred gigabytes of free space:

```
E_INVALIDARG (0x80070057)
```

Video instructions:

To update Mesmerize in the VM:

```
# activate the environment
source ~/venvs/mesmerize/bin/activate
# get the latest version of mesmerize
pip install --upgrade mesmerize
```

**Note:** Virtualization features of your CPU must be enabled in your BIOS. VirtualBox will throw errors if it is not.

##### 1.2.2 Linux

###### pip (PyPI)

**You will need python==3.6 for tensorflow v1**

1. Install python 3.6:

```
# Debian & Ubuntu based
sudo apt-get install python3.6

# Fedora/CentOS
sudo dnf install python36
```

**Note:** If you're using Ubuntu 20.04 you'll need to add a PPA to get python3.6

```
sudo add-apt-repository ppa:deadsnakes/ppa
sudo apt update
sudo apt install python3.6 python3.6-dbg python3.6-dev python3.6-doc python3.6-gdbm␣
↪python3.6-gdbm-dbg python3.6-tk python3.6-tk-dbg python3.6-venv
```

1. Install build tools and other dependencies:

```
# Debian & Ubuntu based distros
sudo apt-get install build-essential python3.6-dev python3.6-venv qt5-default tcl␣
↪graphviz git llvm

# Fedora/CentOS
sudo dnf install @development-tools
sudo dnf install python3-devel tcl graphviz
sudo dnf install llvm
```

For other distributions install the equivalent meta package to get build tools.

If you're on Fedora/CentOS you'll also need `redhat-rpm-config`, install using:

```
sudo dnf install redhat-rpm-config
```

1. Create a new virtual environment:

```
python3.6 -m venv <new_venv_path>
```

2. Activate this environment:

```
source <new_venv_path/bin/activate>
```

3. Make sure you have a recent version of pip and setuptools:

```
pip install --upgrade pip setuptools
```

4. Install numpy & cython:

```
pip install numpy cython
```

5. Install `tensorflow` v1.15 (v2 is not supported for nuset) if you want to use Caiman or Nuset:

```
# CPU bound
pip install tensorflow~=1.15
# GPU
pip install tensorflow-gpu~=1.15
```

6. Install tslearn & bottleneck (optional):

```
pip install tslearn~=0.4.1 bottleneck==1.2.1
```

7. Install mesmerize:

```
pip install mesmerize
```

8. Now you should be be able to launch mesmerize from the terminal:

```
mesmerize
```

You will always need to activate the environment for Mesmerize before launching it.

1. If you want Caiman features you'll need to install caiman into the same environment as mesmerize:

```
git clone https://github.com/flatironinstitute/CaImAn
cd CaImAn/
git checkout v1.8.8
source <new_venv_path/bin/activate>
pip install -e .
```

2. You might need to setup Caiman using `caimanmanager.py`. Please see their docs for details: https://caiman.readthedocs.io/en/master/Installation.html#installation-on-macos-and-linux

3. In order to use some features that launch subprocesses, such as the batch manager, you will need to check your *System Configuration settings in Mesmerize* to make sure that it activates the environment that mesmerize is installed in. By default the pre-run commands contain `# source /<path_to_env>/activate'`, you will need to uncomment the line (remove the #) and set the path to your environment.

**Note:** Caiman=>1.8.9 requires tensorflow v2, which is currently not supported by nuset. If you want to use the latest version of caiman, you will need to install tensorflow v2 and use python3.8

##### 1.2.3 Mac OSX

This requires Anaconda and will install Mesmerize in an Anaconda environment. If you want to install into a python virtual environment use the instructions for the Linux installation from step #3 onward. Tested on macOS Catalina 10.15.1

Download Anaconda for Python 3: https://www.anaconda.com/distribution/

First make sure you have xcode:

```
xcode-select --install
```

This might take a while.

**Create an environment & install Mesmerize**

1. Create a new environment using python 3.6:

   ```
   conda create --name mesmerize python=3.6
   ```

2. Enter the environment:

   ```
   source activate mesmerize
   ```

3. Install `caiman` for Caiman features:

   ```
   conda install -c conda-forge caiman
   ```

4. Install Mesmerize. On Mac installing tslearn before mesmerize creates problems on anaconda for some reason:

   ```
   pip install mesmerize
   ```

5. Install cython, and downgrade pandas:

   ```
   conda install Cython pandas~=0.25.3
   ```

6. Install tslearn~=0.4.1:

   ```
   conda install -c conda-forge tslearn=0.4.1
   ```

7. Install bottleneck (optional):

   ```
   pip install bottleneck==1.2.1
   ```

8. To launch Mesmerize call it from the terminal:

   ```
   mesmerize
   ```

You will always need to activate the environment for Mesmerize before launching it.

**You might get a matplotlib error similar to below**:

```
Bad val 'qt5' on line #1
"backend: qt5

in file "/Users/kushal/.matplotlib/matplotlibrc"
Key backend: Unrecognized backend string 'qt5': valid strings are ['GTK3Agg', 'GTK3Cairo
↪', 'MacOSX', 'nbAgg', 'Qt4Agg', 'Qt4Cairo', 'Qt5Agg', 'Qt5Cairo', 'TkAgg', 'TkCairo',
↪'WebAgg', 'WX', 'WXAgg', 'WXCairo', 'agg', 'cairo', 'pdf', 'pgf', 'ps', 'svg',
↪'template']
```

(continues on next page)

(continued from previous page)

**To fix this, execute the following which appends the default matplotlib backend-option. Note that this will probably affect matplotlib in all your environments**:

```
echo "backend: qt5" >> ~/.matplotlib/matplotlibrc
```

You might need to setup Caiman using `caimanmanager.py`. Please see their docs for details: https://caiman.readthedocs.io/en/master/Installation.html#installation-on-macos-and-linux

In order to use some features that launch subprocesses, such as the batch manager, you will need to check your *System Configuration settings in Mesmerize* to make sure that it activates the environment that mesmerize is installed in.

##### 1.2.4 Windows

Tested on Windows 10, not sure if it'll work on earlier Windows versions.

Download & install Anaconda for Python 3: https://www.anaconda.com/distribution/

**Make sure you select the option to add anaconda to the PATH environment variable during installation.**

You will also need git: https://gitforwindows.org/

> **Warning:** It is **highly** recommended that you use Mesmerize in a new dedicated environment, even if you already have major dependencies (like caiman) installed in another environment.

**All commands are to be run in the powershell**

1. You will need anaconda to be accessible through powershell. You may need to run powershell as administrator for this step to to work. Close & open a new non-admin powershell after running this:

   ```
   conda init powershell
   ```

You will need a relatively recent version of Anaconda in order to run conda commands through the powershell.

1. Create a new anaconda environment:

   ```
   conda create -n mesmerize python=3.6
   ```

2. Activate the environment:

   ```
   conda activate mesmerize
   ```

3. Install caiman:

   ```
   conda install -c conda-forge caiman
   ```

4. Downgrade pandas, install Cython:

   ```
   conda install Cython pandas~=0.25.3
   ```

5. Install tslearn (optional):

   ```
   conda install -c conda-forge tslearn=0.4.1
   ```

6. Install bottleneck (optional):

```
pip install bottleneck==1.2.1
```

7. Install graphviz:

```
conda install graphviz
```

8. Install pywin32:

```
pip install pywin32
```

9. Install Mesmerize:

```
pip install mesmerize
```

10. Allow powershell to execute scripts. Run powershell as administrator to execute these commands. This is required for the batch manager and k-Shape GUI which launch external processes. This may affect the security of your system by allowing scripts to be executable. I'm not an expert on Windows so if someone knows a better way to do this let me know! As far as I know, I'm not sure why you would try to execute untrusted scripts so this shouldn't be a concern?:

```
Set-ExecutionPolicy RemoteSigned
Set-ExecutionPolicy Bypass -scope Process -Force
```

11. Launch Mesmerize:

```
mesmerize
```

You might need to setup Caiman using `caimanmanager.py`. Please see their docs for details: https://caiman.readthedocs.io/en/master/Installation.html#installation-on-macos-and-linux

**Note:** In order to use some features, such as the batch manager, you will need to check your *System Configuration settings in Mesmerize* to make sure that it activates the conda environment that mesmerize is installed in. By default the pre-run commands contain `# conda activate mesmerize` but you will need to uncomment the line (remove the #) or change it if you are using an environment with a different name.

##### 1.2.5 From GitHub (Development)

First, make sure you have compilers & python3.6 (see the details above for various Linux distros or Mac OSX)

1. Create a virtual environment:

```
# Choose a path to house the virtual environment
python3.6 -m venv /path/to/venv
```

2. Activate the virtual environment:

```
source /path/to/venv/bin/activate
```

3. Upgrade pip & setuptools & install some build dependencies:

```
pip install --upgrade pip setuptools
pip install Cython numpy
```

4. Install `tensorflow` or `tensorflow-gpu`, you must use version `~=1.15`:

```
pip install tensorflow~=1.15
```

5. Install tslearn (required) & bottleneck (optional):

```
pip install tslearn~=0.4.1 bottleneck==1.2.1
```

6. If you want Caiman features you'll need to install caiman into the same environment as mesmerize:

```
git clone https://github.com/flatironinstitute/CaImAn
cd CaImAn/
source <new_venv_path/bin/activate>
pip install -e .
```

7. You might need to setup Caiman using `caimanmanager.py`. Please see their docs for details: https://caiman.readthedocs.io/en/master/Installation.html#installation-on-macos-and-linux

8. Fork the main repo on github and clone it, or install from our repo:

```
git clone https://github.com/kushalkolar/MESmerize.git
# or your own form
# git clone https://github.com/<your_github_username>/MESmerize.git
cd MESmerize
```

9. Switch to new branch:

```
git checkout -b my-new-feature
```

10. Install in editable mode:

```
pip install -e .
```

11. Make your changes to the code & push to your fork:

```
git push origin my-new-feature
```

12. Create a pull request if you want to incorporate it into the main Mesmerize repo.

#### 1.3 Getting Help

If you have questions, encounter an issue, or need help:

- **Post an issue on GitHub:** https://github.com/kushalkolar/MESmerize/issues

Please provide all the details that the issue template asks for.

- **Contact us on gitter (for smaller questions or for discussion).** https://gitter.im/mesmerize_discussion/community

Please use the GitHub issue tracker for actual issues, error messages/tracebacks etc. Do not post large error messages/tracebacks in the gitter room, it gets very messy it's harder for us to help you.

#### 1.4 FAQs

##### 1.4.1 ROIs

1. **Can I delete an ROI?**
    - See *ROI Manager guide*
2. **I don't want to delete ROIs but I want to mark them for exclusion in further analysis, how can I do this?**
    - You can do this by creating an ROI type category. See <link here> *Add New ROI Type Later* which uses this as an example. You can also create this ROI Type category when you create a New Project, not necessarily when you already have a project as the example uses.
3. **Can I tag more than one piece of information to each ROI?**
    - Yes, add as many ROI Type categories as you want in the Project Configuration.

    **See also:**

    *Project Configuration*
4. **I already have a Mesmerize project with many Samples in it. Can I add a new ROI Type category?**
    - Yes, just add it to your *Project Configuration*
5. **Can some samples in my project have ROIs that originate from CNMF(E) and others that are manually drawn?**
    - Yes, but be aware that you may need to separate the CNMF(E) and manual data in downstream analysis if using flowchart nodes that work with data from specific sources.

##### 1.4.2 CNMFE

1. **I have ROIs that clearly encompass multiple cells instead of just one**
    - Increase min_coor
    - Might help to reduce gSig as well
2. **I have too many bad ROIs around random regions that are clearly noise**
    - Increase *min_pnr*
3. **Min_PNR image is completely blue and void of any signals**
    - Increase *gSig*
4. **Vmin slider is stuck in Inspect Correlation & PNR GUI.**
    - Close and reopen it. This is a matplotlib issue, not something I can fix.

##### 1.4.3 Caiman Motion Correction

1. **I have video tearing**
    - Try increasing *upsample grid*
    - It's possible that the movement is too severe to be motion corrected. When the movement is so severe that the information do not exist, it is impossible to motion correct it.
2. **My animal is growing**
    - This is growth, not motion. Unfortunately cannot be corrected for. If you have an idea for a technique I can try it out.
3. **The output actually has more motion, it has created false motion.**
    - **Try these things:**
        - Reduce *Strides* & *Overlaps* by ~25%
        - Reduce *max shifts X & Y* by ~25%
        - Reduce *max deviation from rigid* by ~25%

##### 1.4.4 Project Organization

1. **Can I modify a sample?**
    - Yes. Double click the Sample ID in the Project Browser to open it in a viewer. You can then make any modifications you want and then go to File -> Add to Project and select the "Save Changes (overwrite)" option at the bottom. If you have not changed the image sequence itself you can uncheck "Overwrite image data".
2. **Can I change the SampleID?**
    - No this is fundamentally impossible.
    - A work-around is to open that Sample in the viewer (double click it in the project browser), make any modifications if necessary, then go to File -> Add to Project, enter the the information for this sample and a new Animal ID (and Trial ID if wanted), and then select the option "Add to Project Dataframe" at the bottom and click Proceed. This will now add a new Sample to the project with this Sample ID. You can then delete the previous Sample.
3. **Can I add a new Custom Column, ROI Column, or Stimulus Column to my project when I already have samples in my pr**
    - Yes, just modify your *Project Configuration*. In the Welcome Window go to Configure -> Project Configuration. Add anything that you want, and then click "Save and Apply". **It's best to immediately restart Mesmerize whenever you change your project configuration.**
    - If you are adding a new Custom Column you can enter a "Dataframe replace value". This will allow you to set a value for all existing Samples in your project for this new column.
    - If you do not set a Dataframe replace value it will label all existing as "untagged"

#### 1.5 Citation guide

Mesmerize provides interfaces to many great tools that were created by other developers. Please cite the papers for the following Viewer Modules and analysis methods that you use in addition to citing Mesmerize. I would also suggest citing numpy, pandas, scipy, sklearn, and matplotlib.

Mesmerize relies heavily on pyqtgraph widgets. Citing pyqtgraph.

##### 1.5.1 Viewer

| Module | Cite |
|---|---|
| *CNMF* | Giovannucci A., Friedrich J., Gunn P., Kalfon J., Brown, B., Koay S.A., Taxidis J., Najafi F., Gauthier J.L., Zhou P., Baljit, K.S., Tank D.W., Chklovskii D.B., Pnevmatikakis E.A. (2019). CaImAn: An open source tool for scalable Calcium Imaging data Analysis. eLife 8, e38173. https://elifesciences.org/articles/38173<br><br>Pnevmatikakis, E.A., Soudry, D., Gao, Y., Machado, T., Merel, J., … & Paninski, L. (2016). Simultaneous denoising, deconvolution, and demixing of calcium imaging data. Neuron 89(2):285-299. http://dx.doi.org/10.1016/j.neuron.2015.11.037<br><br>Pnevmatikakis, E.A., Gao, Y., Soudry, D., Pfau, D., Lacefield, C., … & Paninski, L. (2014). A structured matrix factorization framework for large scale calcium imaging data analysis. arXiv preprint arXiv:1409.2903. http://arxiv.org/abs/1409.2903 |
| *CNMFE* | In addition to the above CNMF papers:<br>Zhou, P., Resendez, S. L., Rodriguez-Romaguera, J., Jimenez, J. C., Neufeld, S. Q., Giovannucci, A., … Paninski, L. (2018). Efficient and accurate extraction of in vivo calcium signals from microendoscopic video data. ELife, 7. doi: https://doi.org/10.7554/eLife.28728.001 |
| *Caiman Motion Correction* | Giovannucci A., Friedrich J., Gunn P., Kalfon J., Brown, B., Koay S.A., Taxidis J., Najafi F., Gauthier J.L., Zhou P., Baljit, K.S., Tank D.W., Chklovskii D.B., Pnevmatikakis E.A. (2019). CaImAn: An open source tool for scalable Calcium Imaging data Analysis. eLife 8, e38173. https://elifesciences.org/articles/38173<br><br>Pnevmatikakis, E.A., and Giovannucci A. (2017). NoRMCorre: An online algorithm for piecewise rigid motion correction of calcium imaging data. Journal of Neuroscience Methods, 291:83-92. https://doi.org/10.1016/j.jneumeth.2017.07.031 |
| Nuset Segmentation | Yang L, Ghosh RP, Franklin JM, Chen S, You C, Narayan RR, et al. (2020) NuSeT: A deep learning tool for reliably separating and analyzing crowded cells. PLoS Comput Biol 16(9): e1008193. https://doi.org/10.1371/journal.pcbi.1008193 |

##### 1.5.2 Nodes/Analysis

| Node/Method | Cite |
|---|---|
| *k-Shape clustering* | Paparrizos, J., & Gravano, L. (2016). k-Shape. ACM SIGMOD Record, 45(1), 69–76. doi: http://dx.doi.org/10.1145/2723372.2737793<br><br>Romain Tavenard, Johann Faouzi, Gilles Vandewiele and Felix Divo, Guillaume Androz, Chester Holtz, Marie Payne, Roman Yurchak, Marc Ruwurm, Kushal Kolar, & Eli Woods. (2017). Tslearn, A Machine Learning Toolkit for Time Series Data. Journal of Machine Learning Research, (118):16, 2020. http://jmlr.org/papers/v21/20-091.html |
| *Cross-correlation* | Romain Tavenard, Johann Faouzi, Gilles Vandewiele and Felix Divo, Guillaume Androz, Chester Holtz, Marie Payne, Roman Yurchak, Marc Ruwurm, Kushal Kolar, & Eli Woods. (2017). Tslearn, A Machine Learning Toolkit for Time Series Data. Journal of Machine Learning Research, (118):16, 2020. http://jmlr.org/papers/v21/20-091.html |
| *TVDiff Node* | Rick Chartrand, "Numerical Differentiation of Noisy, Nonsmooth Data," ISRN Applied Mathematics, vol. 2011, Article ID 164564, 11 pages, 2011. https://doi.org/10.5402/2011/164564. |

##### 1.5.3 Scientific Libraries

| Library | |
|---|---|
| numpy | Van Der Walt, S., Colbert, S. C. & Varoquaux, G. The NumPy array: A structure for efficient numerical computation. Comput. Sci. Eng. (2011) doi:10.1109/MCSE.2011.37 |
| pandas | McKinney, W. Data Structures for Statistical Computing in Python. Proc. 9th Python Sci. Conf. (2010) |
| scipy | Virtanen, P., Gommers, R., Oliphant, T.E. et al. SciPy 1.0: fundamental algorithms for scientific computing in Python. Nat Methods (2020). https://doi.org/10.1038/s41592-019-0686-2 |
| sklearn | Pedregosa, F. et al. Scikit-learn: Machine learning in Python. J. Mach. Learn. Res. (2011) |
| matplotlib | Hunter, J. D. Matplotlib: A 2D graphics environment. Comput. Sci. Eng. (2007) |
| pyqtgraph | http://www.pyqtgraph.org/ |

#### 1.6 Create a New Project

##### 1.6.1 Video Tutorial

This tutorial shows how to create a New Project, open images in the Viewer, use the Stimulus Mapping module and perform Caiman motion correction

##### 1.6.2 Biological Questions

**Before you create a new Mesmerize Project you must thoroughly think about the biological questions that you are interested in. Here are some thoughts to help you:**

- The effects of different types of temporary stimulation? Such as poking or odors?
- Are you interested in neuronal activity during specific behavioral periods?
- Differences in calcium dynamics between different anatomical regions?
- Chemogenetic experiments using transgenes to express DDREADs.
- Non-temporary effects of drugs (for example, if the animal is bathed in drug for longer recordings).
  - For example, if you are inducing seizures with PTZ, where you are interested in the differences between a control recording of 5 minutes and subsequent 5 minute recordings where the animal is bathed in PTZ (or whatever duration you determine is biologically meaningful). You could also be interested in a recovery stage to see what happens to the calcium dynamics when you "perfuse-back" the liquid medium (such as seawater, steinberg's solution etc.) without the drug.
- Differences in calcium dynamics between different stages during development
- Differences in calcium dynamics between different cell types using GCaMP driven by specific promoters.

---

##### 1.6.3 New Project

To create a new project click **New Project** in the *Welcome Window*. You will then be prompted to choose a location and a name for the project. This will create a directory with the chosen name in the location you previously selected.

##### 1.6.4 Project Configuration

After setting a project name you must configure it. This is where your biological questions of interest are important. You can change your project configuration later, but it is most time efficient if you enter all your categories of interest now.

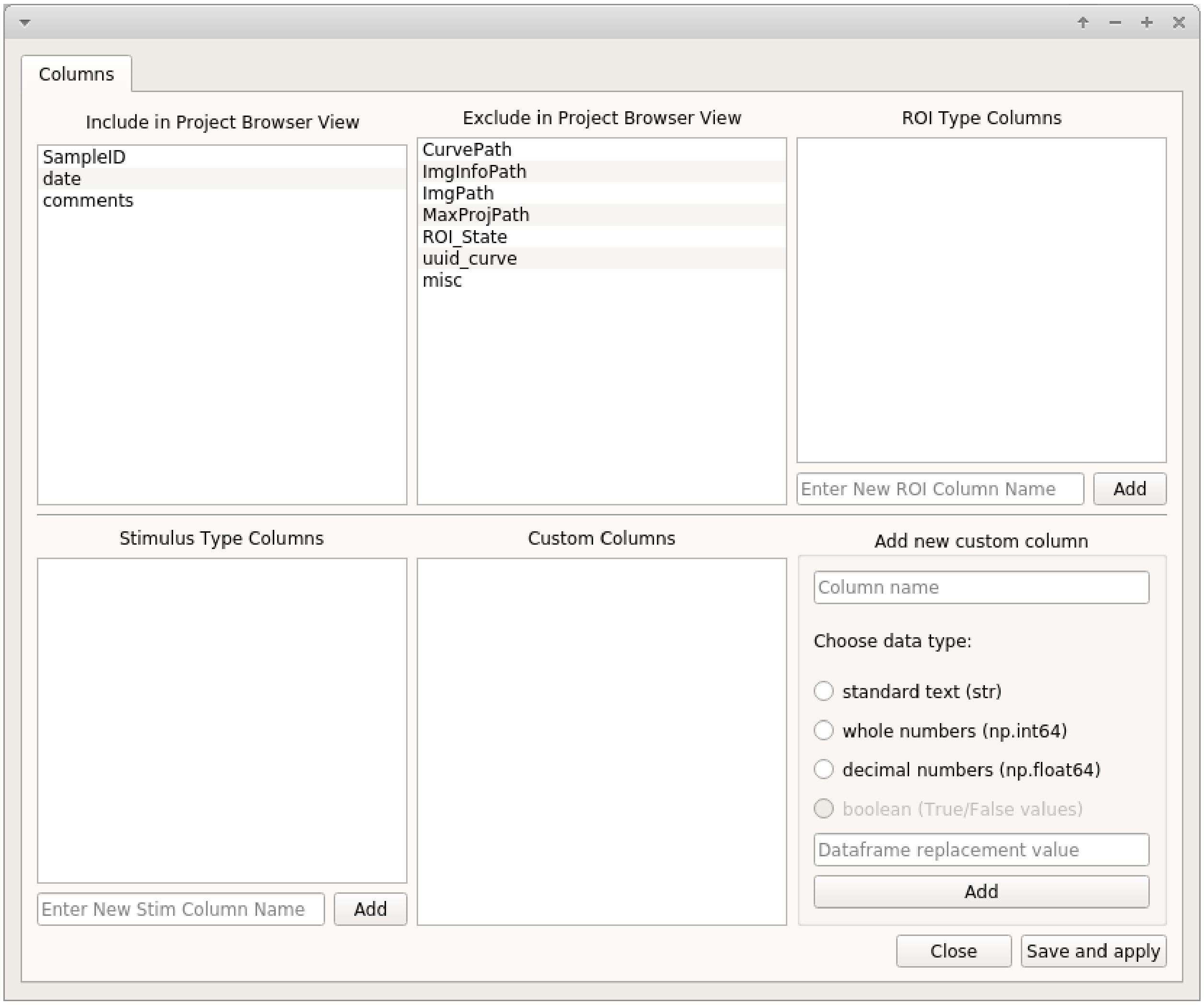


**Warning:** Restart Mesmerize whenever you change the project configuration.

**Note:** If you have Samples in your project and you change the project configuration at a later date to add new columns, all existing rows in your project DataFrame are labelled as "untagged" for the new columns.

**See also:**

*Add To Project Guide* **to understand how the project configuration relates to the addition of data samples to your project**

##### Categorical Data Columns

**Mesmerize allows you to create three main different types of categorical data columns (for the project DataFrame), and an unlimited number of each type. These categorical data columns allow you to group your data during analysis, and therefore perform comparisons between experimental groups. In essense, these categorical data columns form scaffold with which you can create your experimental groups during analysis.**

---

**Note:** You can change the project configuration at any point in the future by adding new columns or changing the visible/hidden columns.

---

**Note:** It is generally advisable to keep the names of your categorical data columns short with lowercase letters. When sharing your project you can provide a mapping for all your keys. This helps maintain consistency throughout your project and makes the data more readable.

---

###### ROI Type Columns

Create ROI-bound *categories* with which you want to group your data. Enter the desired name for the category and click **Add**. Here are some examples:

- If you are interested in calcium dynamics between different anatomical regions, you create a column named `anatomical_region`.
- You want to define defined notochord cell identities on a anterior-posterior axis, defined as "cell_1", "cell_2", ... "cell_n". You can create an ROI Type Column named `notochord_cell_id`.

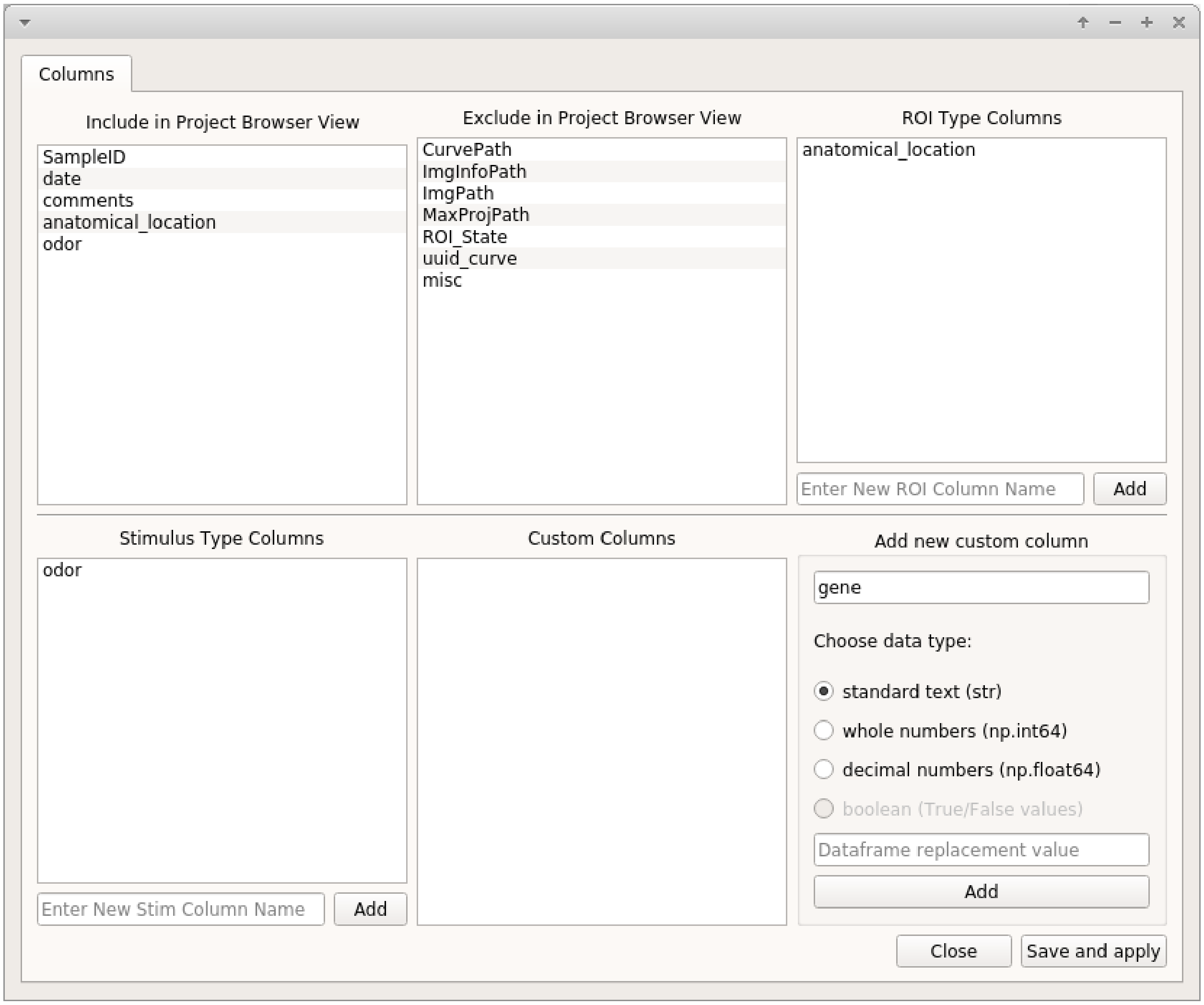


**See also:**

*ROI Manager* to understand how labels can be tagged onto ROIs using these categories that you have defined in the ROI Type Columns.

##### Stimulus Type Columns

If you're interested in mapping temporal information to your traces, such as stimuli or behavioral periods, add a "Stimulus Type column" for each type. This is only for temporary stimulation or behavioral periods that do not span the entire length of the video.

**See also:**

**<link here> Stimulus Mapping guide, to understand how stimuli can be labelled.**

##### Custom Columns

Here you can create categories to tag any other piece of useful information to each Sample. i.e. to the entire video recording. For example:

- You are studying seizures, you perform a 5 minute recording in the medium, and then subsequent 5 minute recordings in PTZ. You can create a category called "drug_state". When you add samples to your project you can tag drug states named "control", "ptz_1", "ptz_2", "ptz_recovery_1" etc.
- This is also what you would use for chemogenetics experiments if you are recording for example without CNO for 5 minutes, and then with CNO for another 5 minutes.

Three different data types can be tagged to a category, **standard text**, **whole numbers**, and **decimal numbers**.

> **Warning:** Data types cannot be changed later. If you are familiar with pandas you can manually change it, and the corresponding value in the project config file.

If you want to tag numerical information, such as the animal's development stage, it can be useful to set the data type to **whole numbers**. This allows you to sort your data numerically. For example you may want to compare dynamics of all curves between stage 48 and 72.

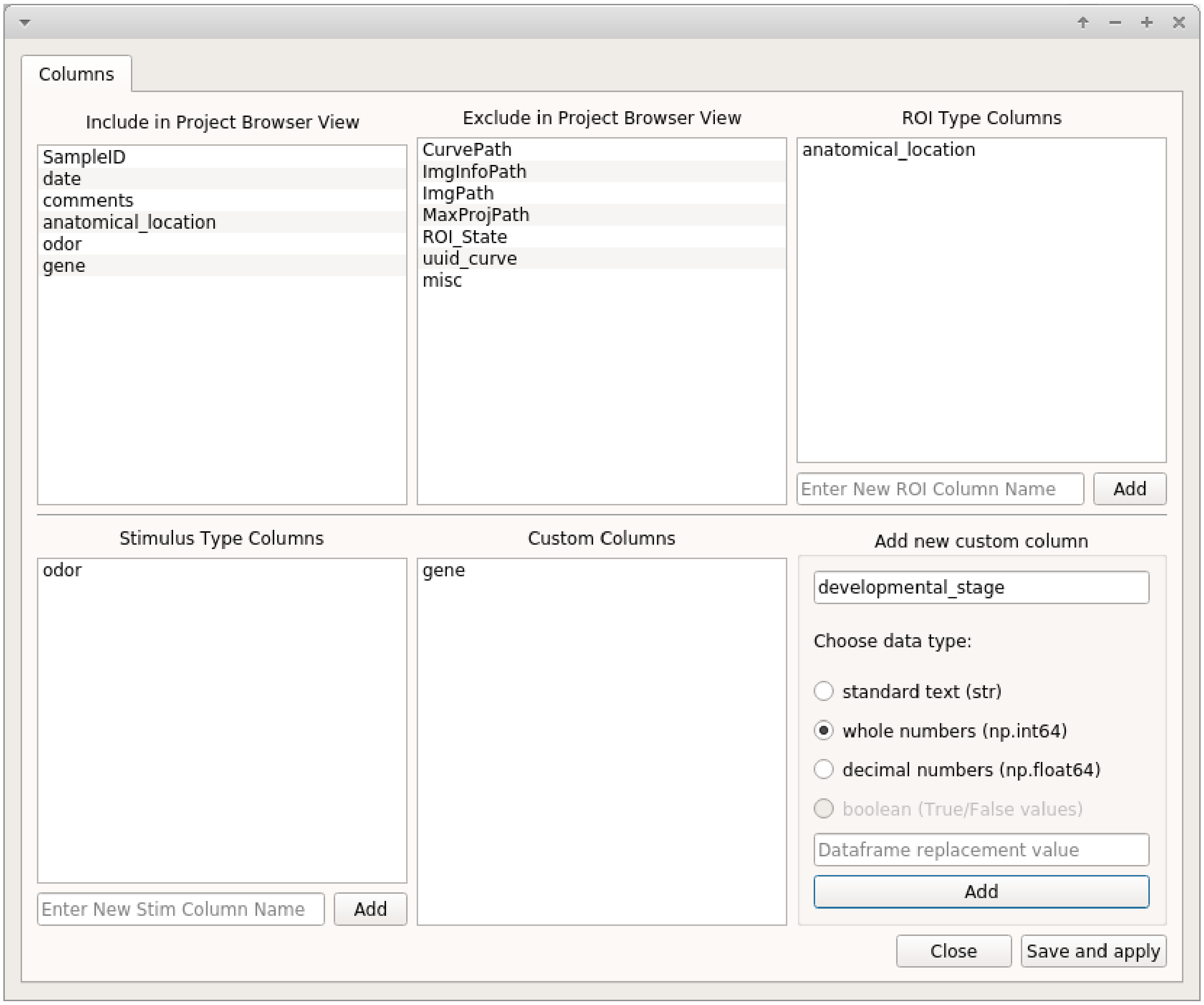

If you are interested in dynamics between different cell types for which you are using specific GCaMP promoters, you can create a custom column called `promoter` or `cell_type` and select **standard text** as the data type.

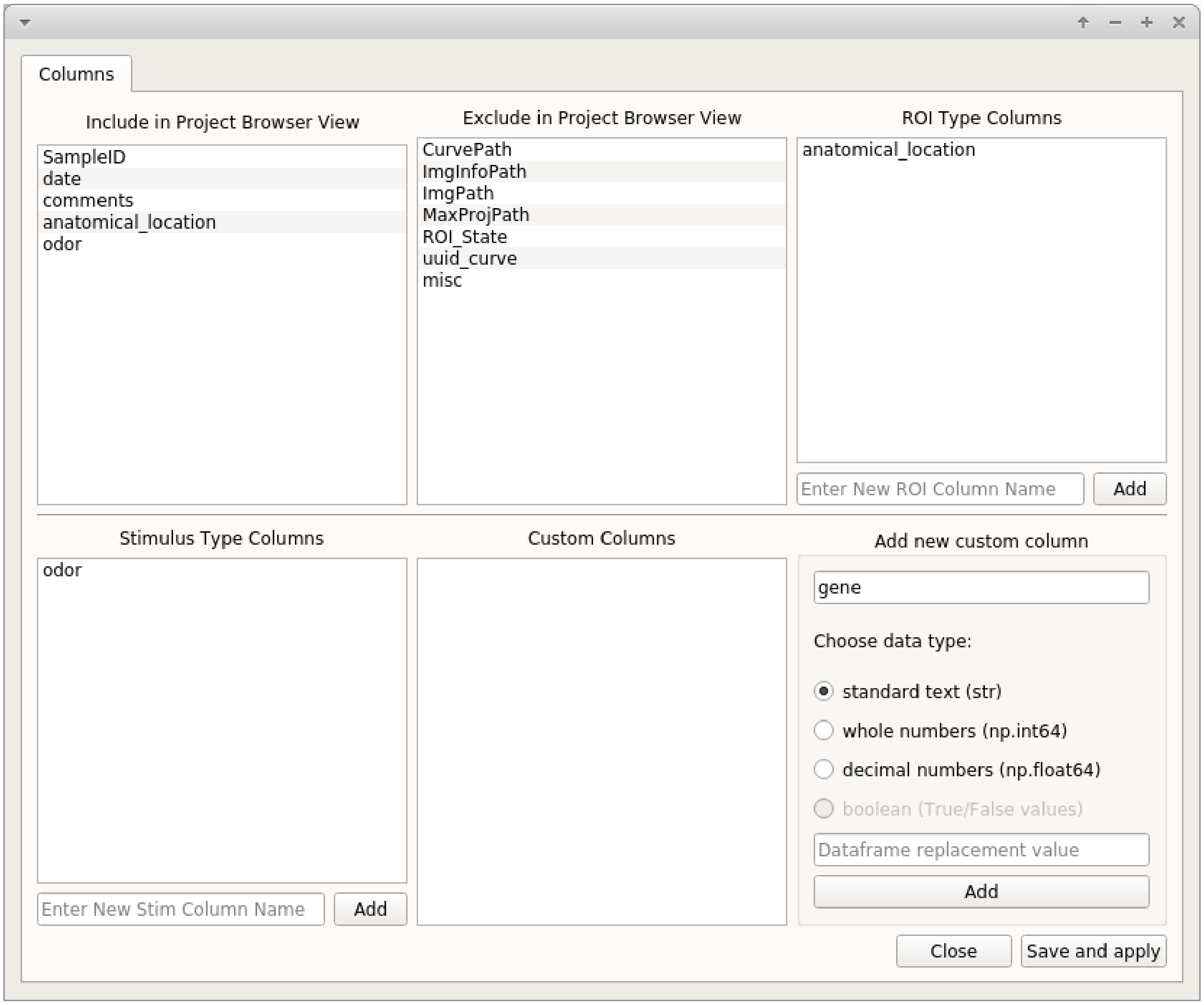


When you add samples to your project from the viewer, you will be prompted to enter information that is directly based on the Custom Columns that you create here.

**See also:**

*Add to Project guide*

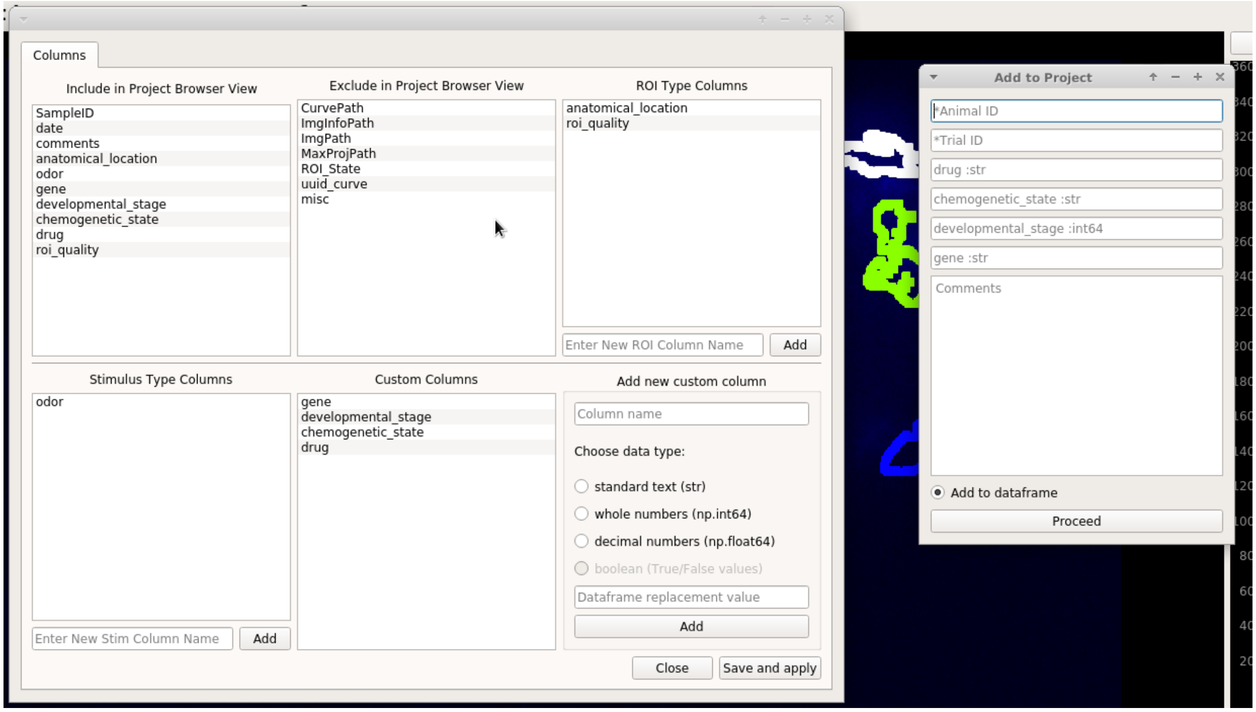


##### Visible / Hidden in Project Browser

You can drag and drop items (column names) between these two lists to set which ones are visible in the Project Browser. This is just to avoid clutter.

**See also:**

*Project Browser*

#### 1.7 Project Browser

**Browse, edit and sort the project DataFrame**

You can open the Project Browser from the *Welcome Window* after you have opened a project.

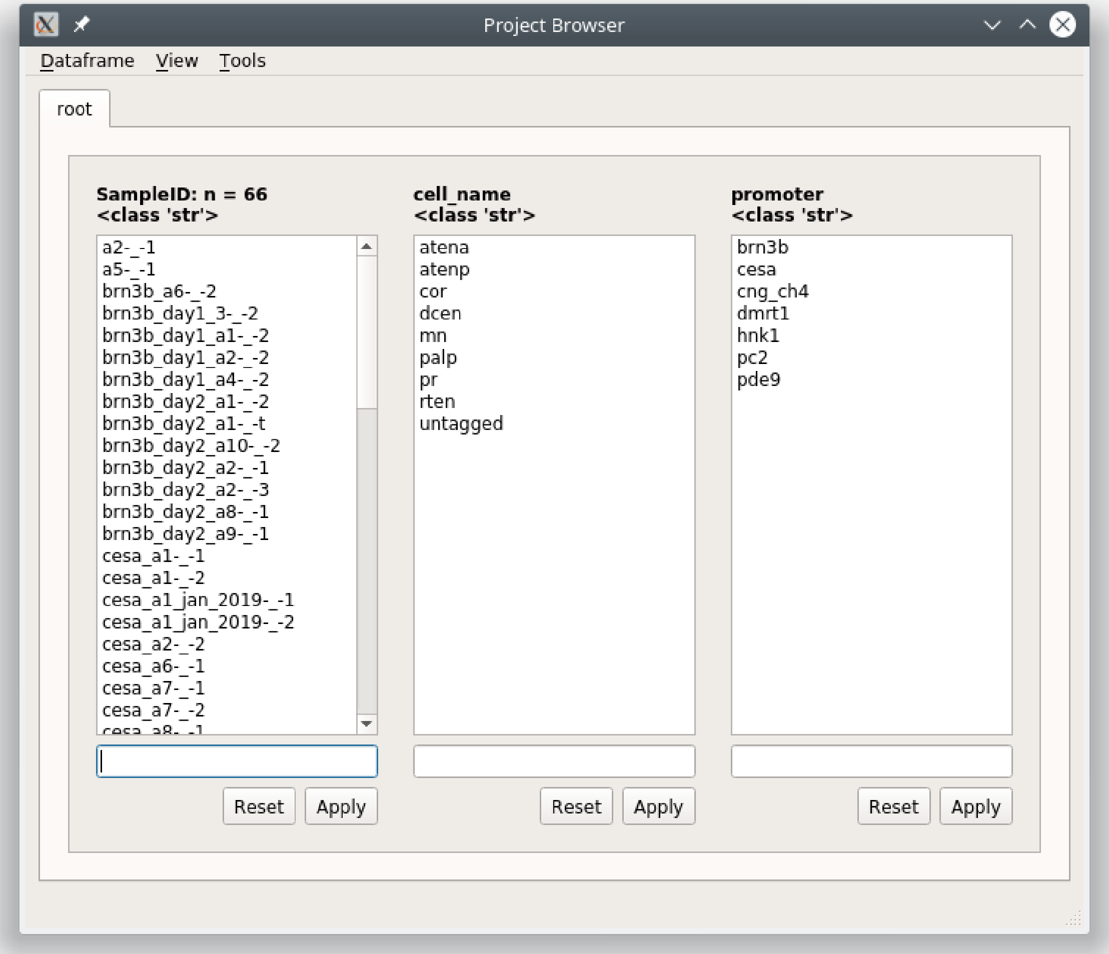


The columns that are visible in the Project Browser Window correspond to the *Project Configuration*. For each column you will see a list which is a set of unique elements from that column in the project DataFrame.

Functions

##### 1.7.1 Open Sample

Double-click on a Sample in the *SampleID* column to open it in the *Viewer*.

In the viewer you can make changes and then save it by going to File -> Add to Project. You will see a "Save changes (overwrite)" option which will overwrite the data for this project Sample with the current data in the viewer work environment. If you have not changed the image sequence data you can uncheck the "Overwrite image data" checkbox, useful if your image sequences are large.

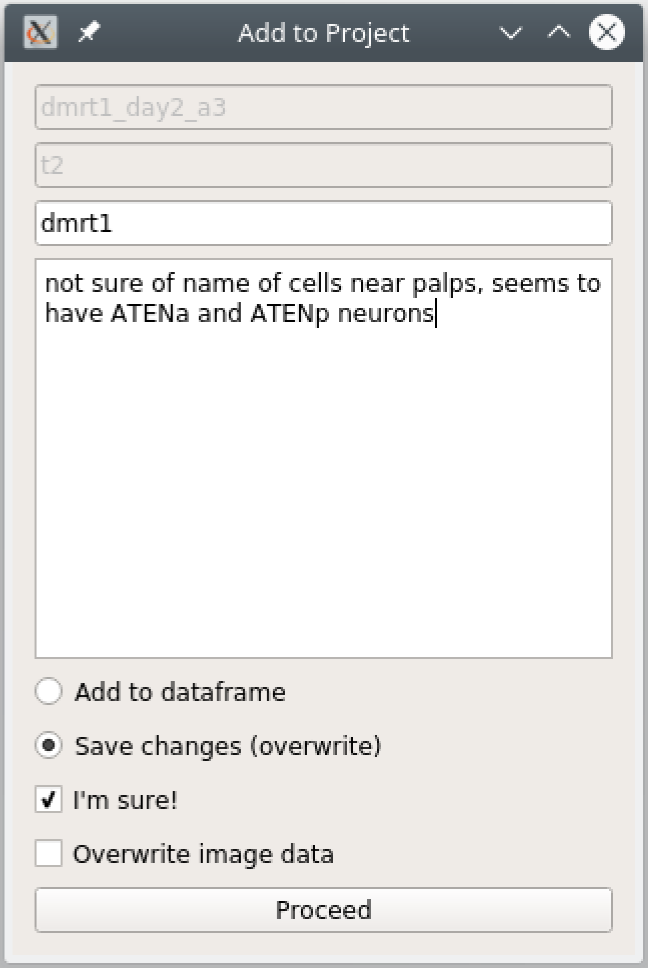


**Note:** You can make any changes that you want to the Sample. This may include things such as changing or adding new tags to ROIs, changing stimulus maps, tagging a new custom column etc.

**Warning:** You can never change the AnimalID or TrialID (i.e. SampleID) since these are partially used as unique identifiers. A workaround is described in the *FAQ for Project Organization*.

##### 1.7.2 Filter

You can sort your Project DataFrame into different groups (such as experimental groups) using text and numerical filters. Type a filter into the text entries that are below the list of elements for a column. You can also click on one or many elements in a column to set those elements as the filters.

If you filter out of the root tab, it will always create a new tab with a name of your choice. If you filter out of any other tab it will apply the filter in-place unless you right click on the "Apply" button and choose "Apply in new tab"

###### Text filters

###### Partial match

To filter out a group based on partial text matches just enter the text into the filter text entry below the column(s) of interest and click "Apply"

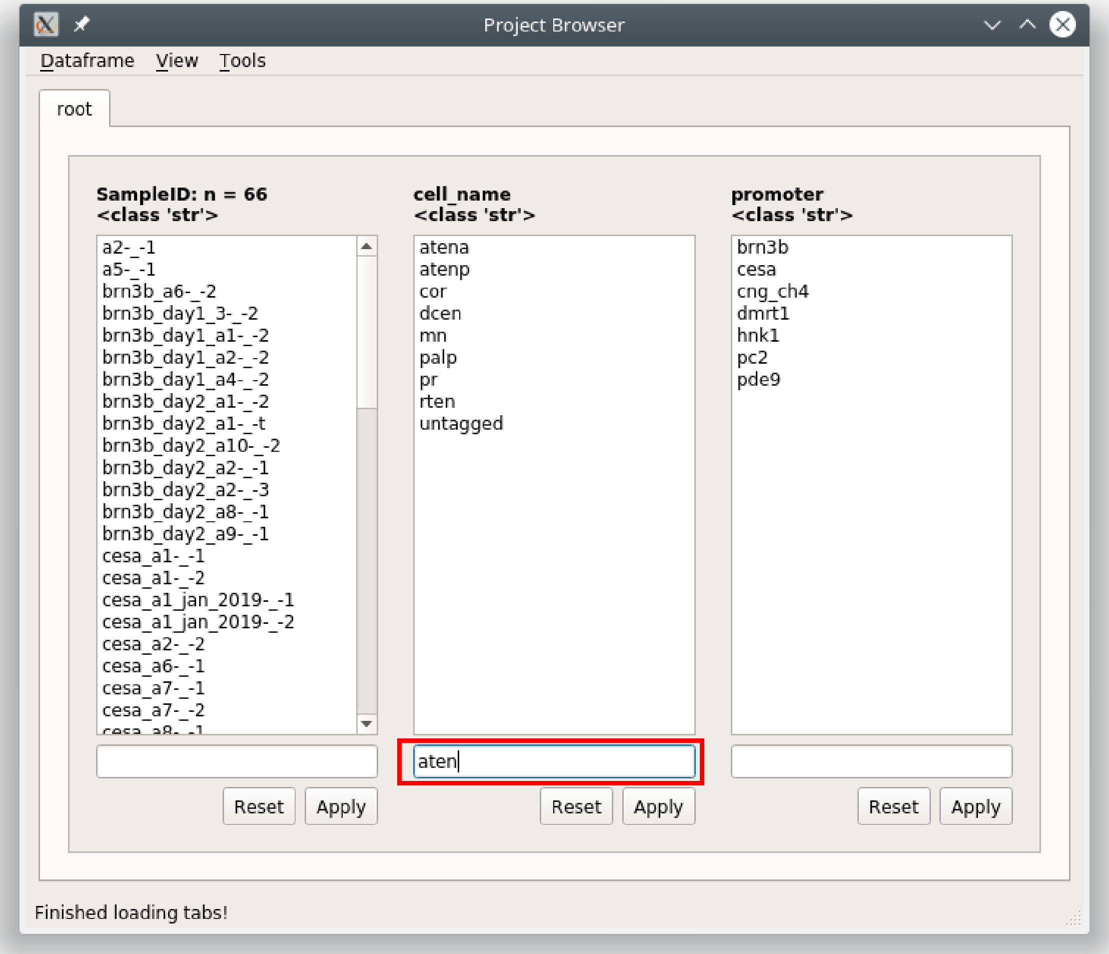


Since this is filtering out of the root tab, you will be prompted to give a name for a new tab that will be created based on the filter you have entered.

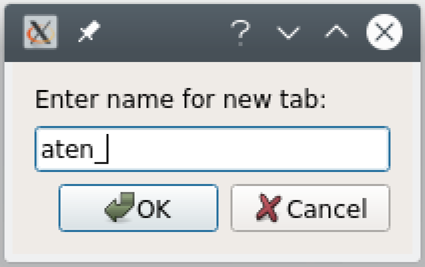


The result is a DataFrame containing all rows where the cell_name contains 'aten'

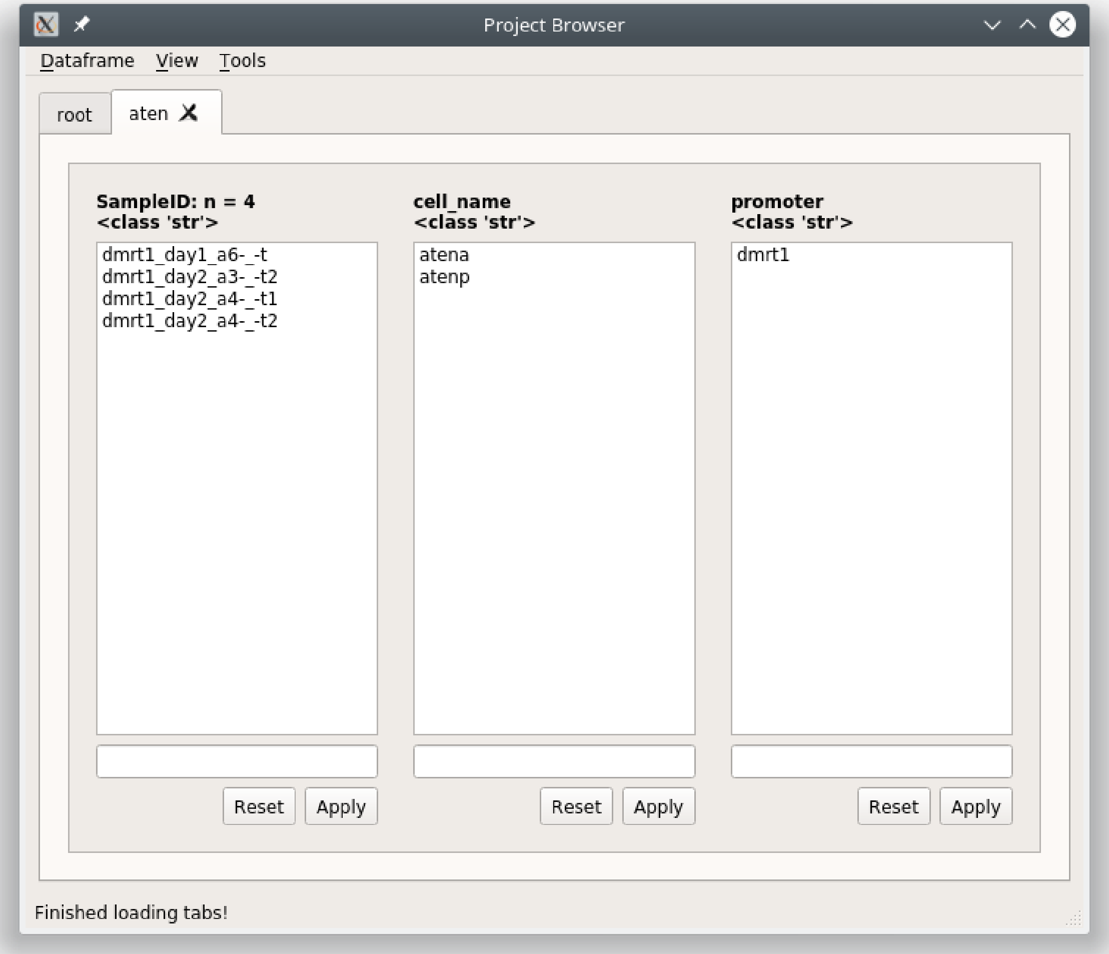


If you go to View -> Current dataframe you can see the whole dataframe.

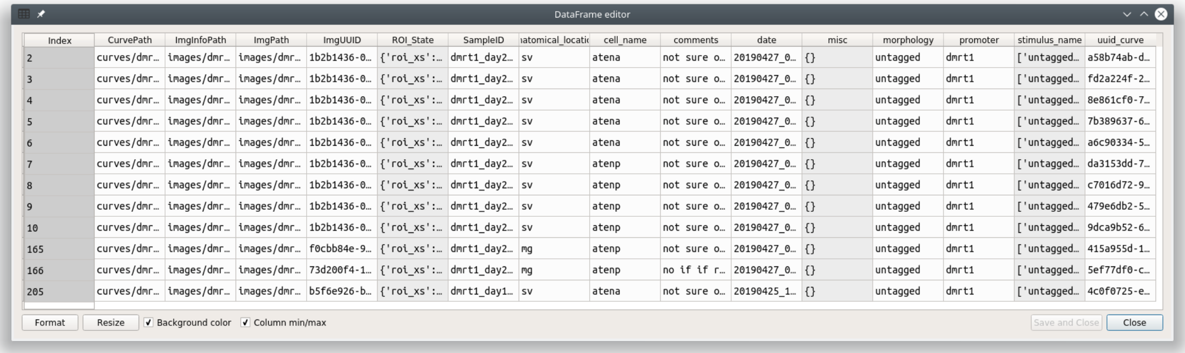


To see how the filter translates to pandas commands go to View -> Current tab filter history

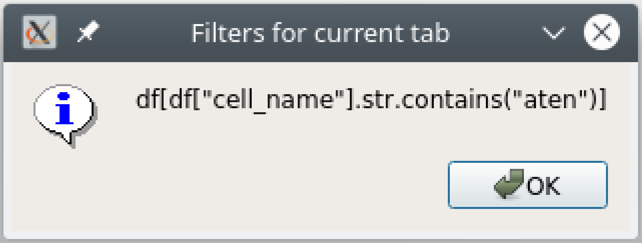


#### Multiple filters

You can combine filters together by using the | seperator. The | acts as an “or” operator.

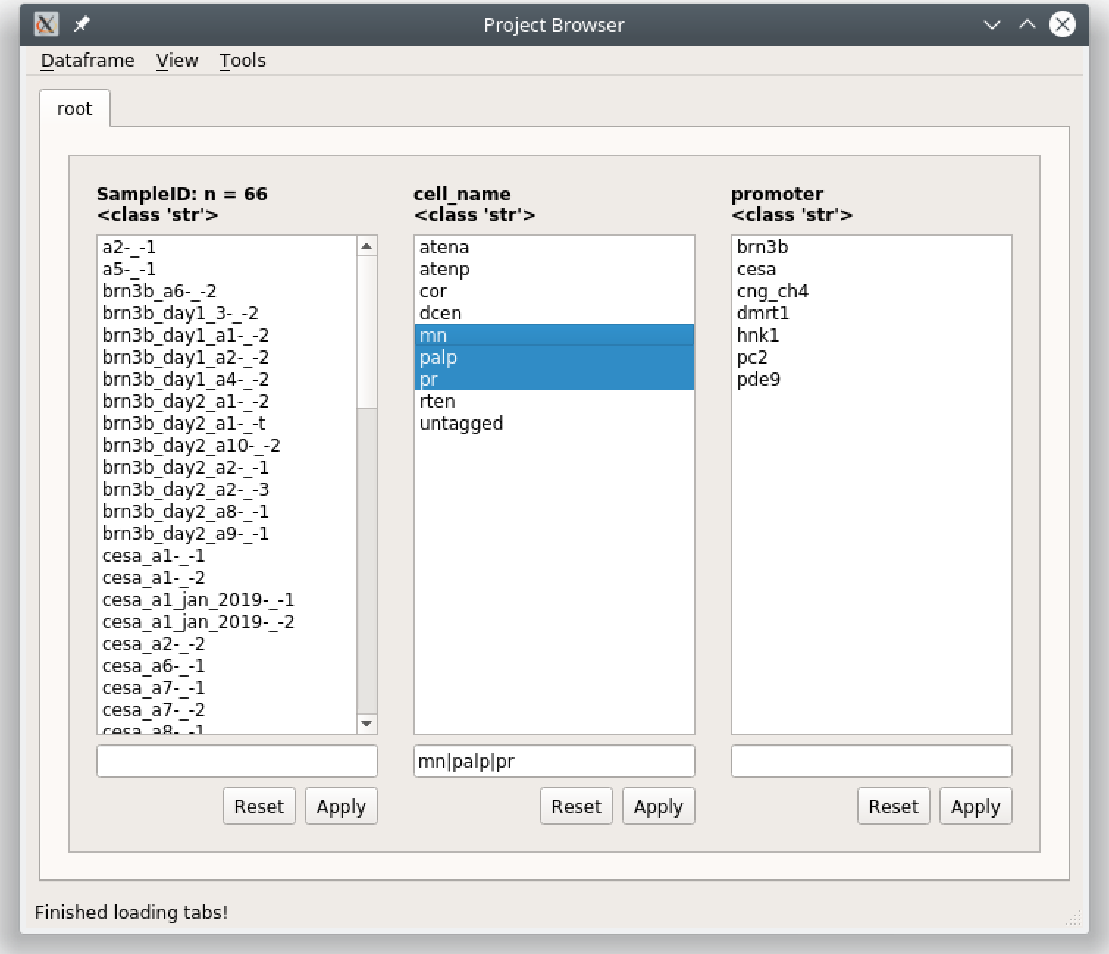


The result is all rows where mn, palp, or pr are in the cell_name column.

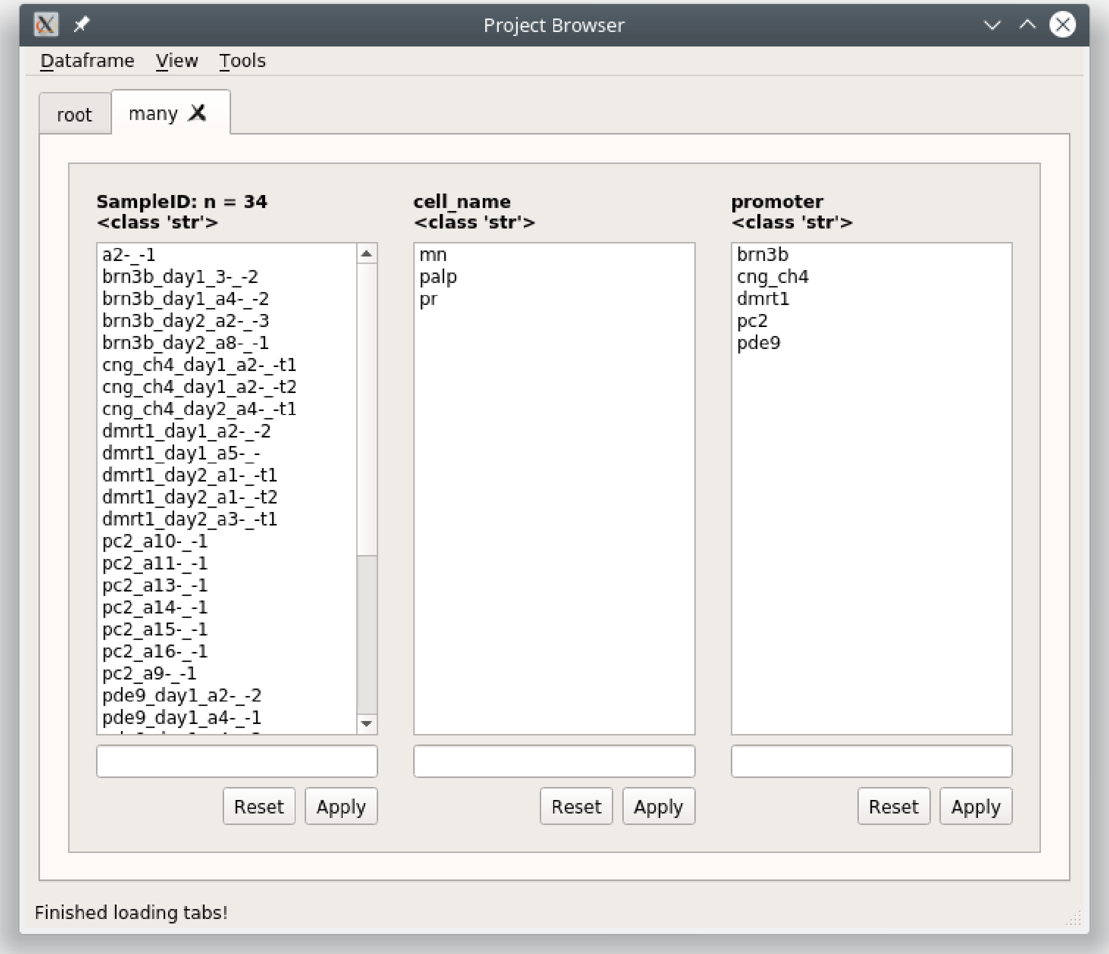


**Note:** This can be combined with *Modifiers*

##### Filter multiple columns

You can filter multiple columns simultaneously if you are not in the root tab. You can create a new tab that is essentially the same as the root by just keeping the filter entries blank and clicking "Apply".

Filter out all rows where the cell_name column contains 'pr' and promoter column contains 'pc2' or 'cng_ch4'.

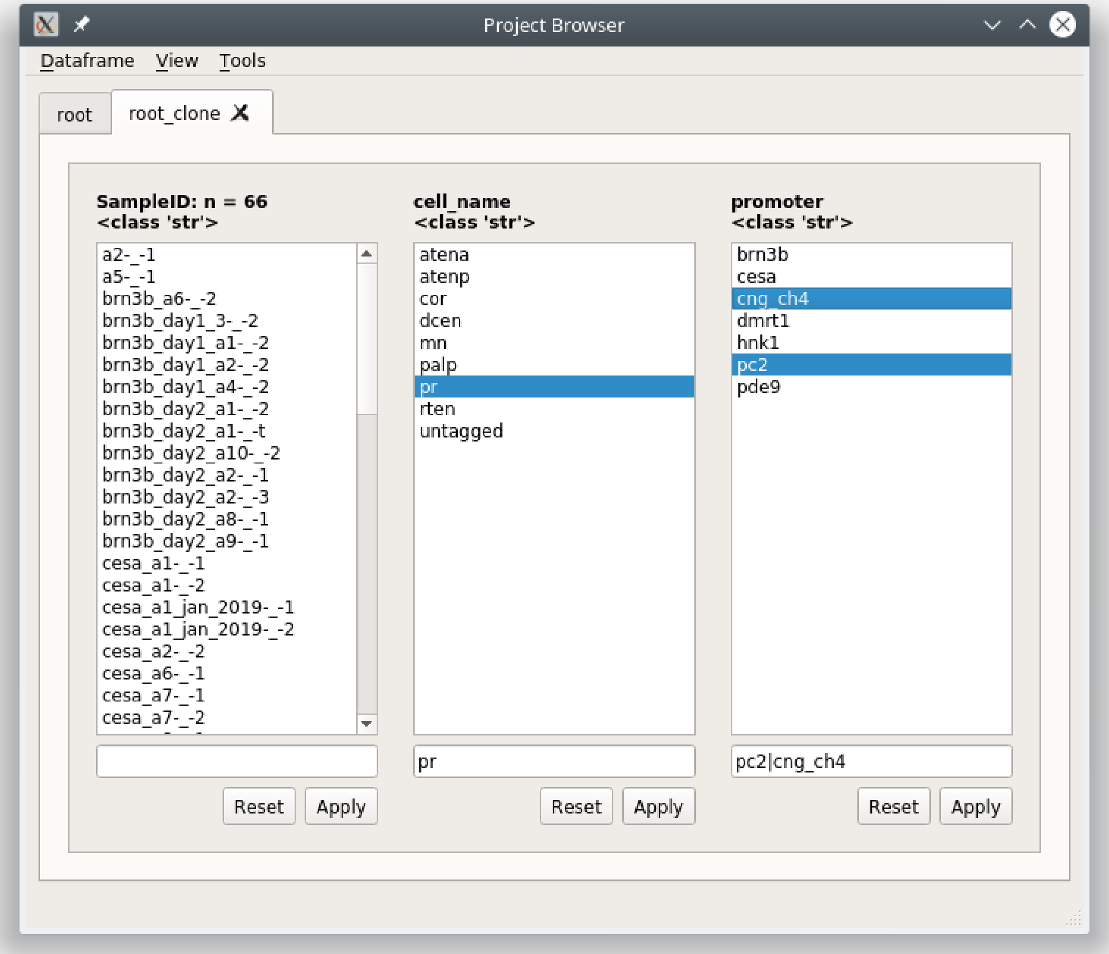


Right click on the “Apply” button and choose “Apply all” or “Apply all in new tab”

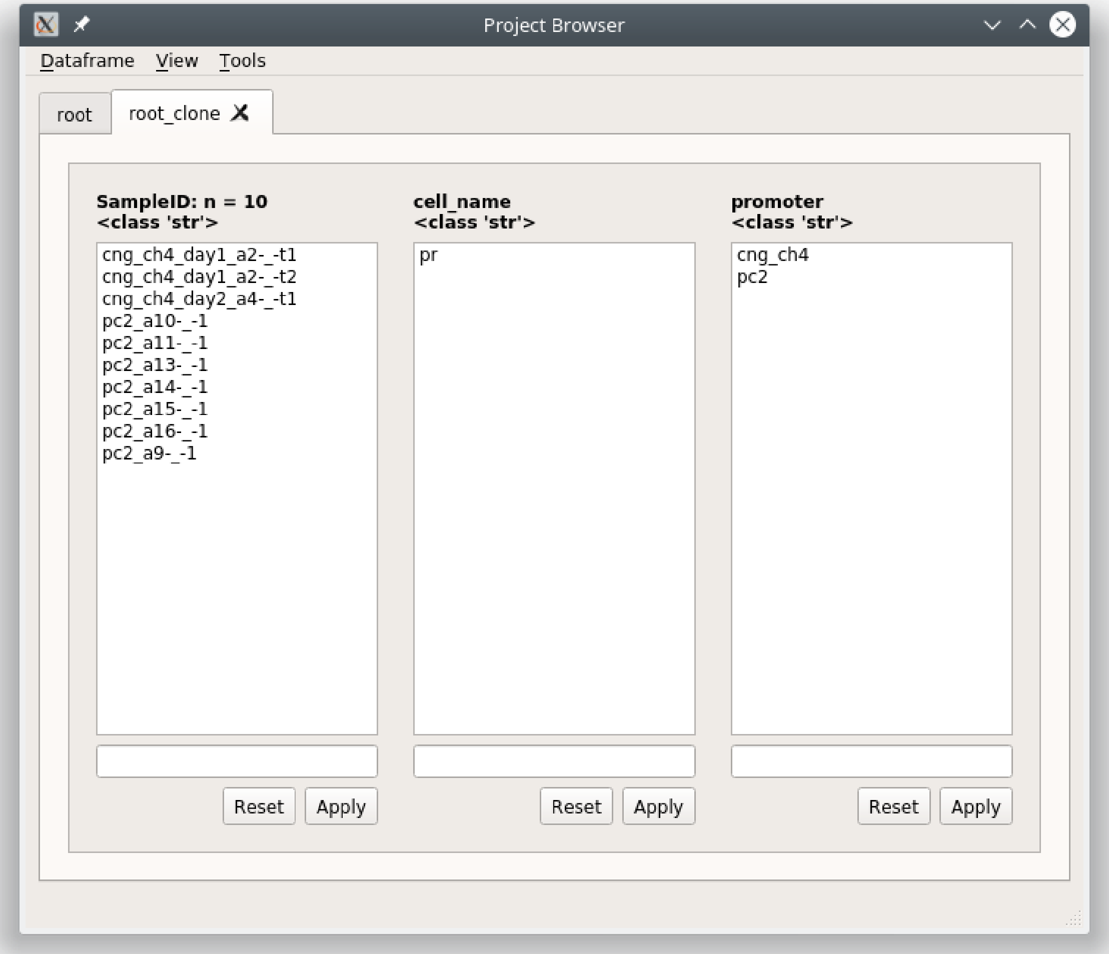


If you view the pandas filter history (View -> Current tab filter history) you can see that the filters for each column are simply applied sequentially.

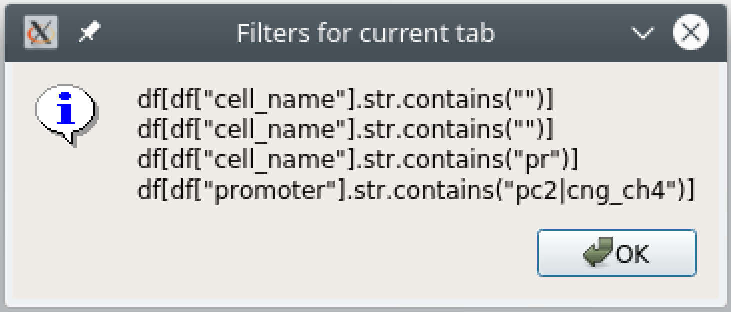


The dataframe

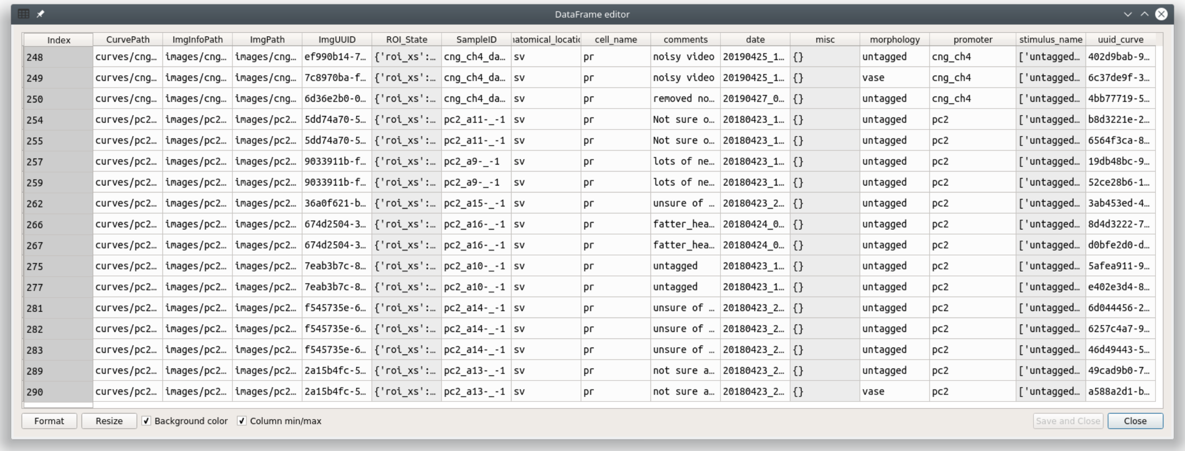


| Index | CurvePath | ImgInfoPath | ImgPath | ImgUUID | ROI_State | SampleID | ıatomical_locatic | cell_name | comments | date | misc | morphology | promoter | stimulus_name | uuid_curve |
|---|---|---|---|---|---|---|---|---|---|---|---|---|---|---|---|
| 248 | curves/cng… | images/cng… | images/cng… | ef990b14-7… | {'roi_xs':… | cng_ch4_da… | sv | pr | noisy video | 20190425_1… | {} | untagged | cng_ch4 | ['untagged… | 402d9bab-9… |
| 249 | curves/cng… | images/cng… | images/cng… | 7c8970ba-f… | {'roi_xs':… | cng_ch4_da… | sv | pr | noisy video | 20190425_1… | {} | vase | cng_ch4 | ['untagged… | 6c37de9f-3… |
| 250 | curves/cng… | images/cng… | images/cng… | 6d36e2b0-0… | {'roi_xs':… | cng_ch4_da… | sv | pr | removed no… | 20190427_0… | {} | untagged | cng_ch4 | ['untagged… | 4bb77719-5… |
| 254 | curves/pc2… | images/pc2… | images/pc2… | 5dd74a70-5… | {'roi_xs':… | pc2_a11-_-1 | sv | pr | Not sure o… | 20180423_1… | {} | untagged | pc2 | ['untagged… | b8d3221e-2… |
| 255 | curves/pc2… | images/pc2… | images/pc2… | 5dd74a70-5… | {'roi_xs':… | pc2_a11-_-1 | sv | pr | Not sure o… | 20180423_1… | {} | untagged | pc2 | ['untagged… | 6564f3ca-8… |
| 257 | curves/pc2… | images/pc2… | images/pc2… | 9033911b-f… | {'roi_xs':… | pc2_a9-_-1 | sv | pr | lots of ne… | 20180423_1… | {} | untagged | pc2 | ['untagged… | 19db48bc-9… |
| 259 | curves/pc2… | images/pc2… | images/pc2… | 9033911b-f… | {'roi_xs':… | pc2_a9-_-1 | sv | pr | lots of ne… | 20180423_1… | {} | untagged | pc2 | ['untagged… | 52ce28b6-1… |
| 262 | curves/pc2… | images/pc2… | images/pc2… | 36a0f621-b… | {'roi_xs':… | pc2_a15-_-1 | sv | pr | unsure of … | 20180423_2… | {} | untagged | pc2 | ['untagged… | 3ab453ed-4… |
| 266 | curves/pc2… | images/pc2… | images/pc2… | 674d2504-3… | {'roi_xs':… | pc2_a16-_-1 | sv | pr | fatter_hea… | 20180424_0… | {} | untagged | pc2 | ['untagged… | 8d4d3222-7… |
| 267 | curves/pc2… | images/pc2… | images/pc2… | 674d2504-3… | {'roi_xs':… | pc2_a16-_-1 | sv | pr | fatter_hea… | 20180424_0… | {} | untagged | pc2 | ['untagged… | d0bfe2d0-d… |
| 275 | curves/pc2… | images/pc2… | images/pc2… | 7eab3b7c-8… | {'roi_xs':… | pc2_a10-_-1 | sv | pr | untagged | 20180423_1… | {} | untagged | pc2 | ['untagged… | 5afea911-9… |
| 277 | curves/pc2… | images/pc2… | images/pc2… | 7eab3b7c-8… | {'roi_xs':… | pc2_a10-_-1 | sv | pr | untagged | 20180423_1… | {} | untagged | pc2 | ['untagged… | e402e3d4-8… |
| 281 | curves/pc2… | images/pc2… | images/pc2… | f545735e-6… | {'roi_xs':… | pc2_a14-_-1 | sv | pr | unsure of … | 20180423_2… | {} | untagged | pc2 | ['untagged… | 6d044456-2… |
| 282 | curves/pc2… | images/pc2… | images/pc2… | f545735e-6… | {'roi_xs':… | pc2_a14-_-1 | sv | pr | unsure of … | 20180423_2… | {} | untagged | pc2 | ['untagged… | 6257c4a7-9… |
| 283 | curves/pc2… | images/pc2… | images/pc2… | f545735e-6… | {'roi_xs':… | pc2_a14-_-1 | sv | pr | unsure of … | 20180423_2… | {} | untagged | pc2 | ['untagged… | 46d49443-5… |
| 289 | curves/pc2… | images/pc2… | images/pc2… | 2a15b4fc-5… | {'roi_xs':… | pc2_a13-_-1 | sv | pr | not sure a… | 20180423_2… | {} | untagged | pc2 | ['untagged… | 49cad9b0-7… |
| 290 | curves/pc2… | images/pc2… | images/pc2… | 2a15b4fc-5… | {'roi_xs':… | pc2_a13-_-1 | sv | pr | not sure a… | 20180423_2… | {} | vase | pc2 | ['untagged… | a588a2d1-b… |



#### Modifiers

You can perform other types of matches, such as exact matches, negations, and exact negations. Enter the filter and then right click on the text entry to see available modifiers and choose the desired modifier.

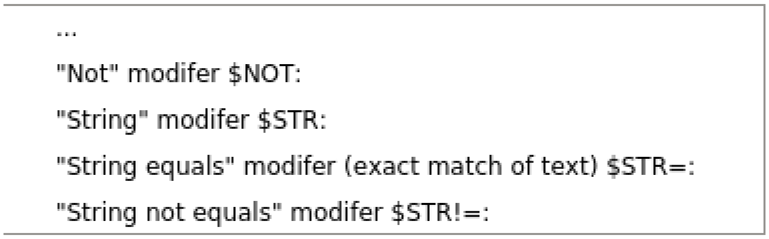


| Modifier | Description |
|---|---|
| $NOT: | Results in the negation of partial matches |
| $STR: | Treats the filter as a str, same as Partial Match (see above sub-section) |
| $STR=: | Exact text match |
| $STR!=: | Negation of exact text match |

##### Numerical filters

By default the filters in all entires are treated as text. If your column contains numerical data you have additional options for modifiers. The first four modifiers are the *same as explained above*. The rest are self explanatory.

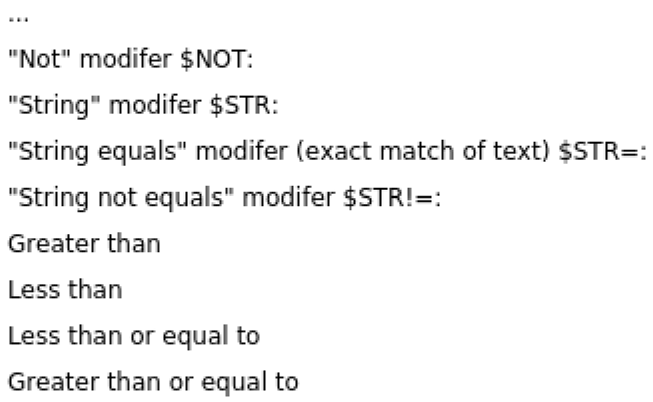


#### 1.7.3 Editor

You can view and edit the Project DataFrame directly in a GUI using the DataFrame editor.

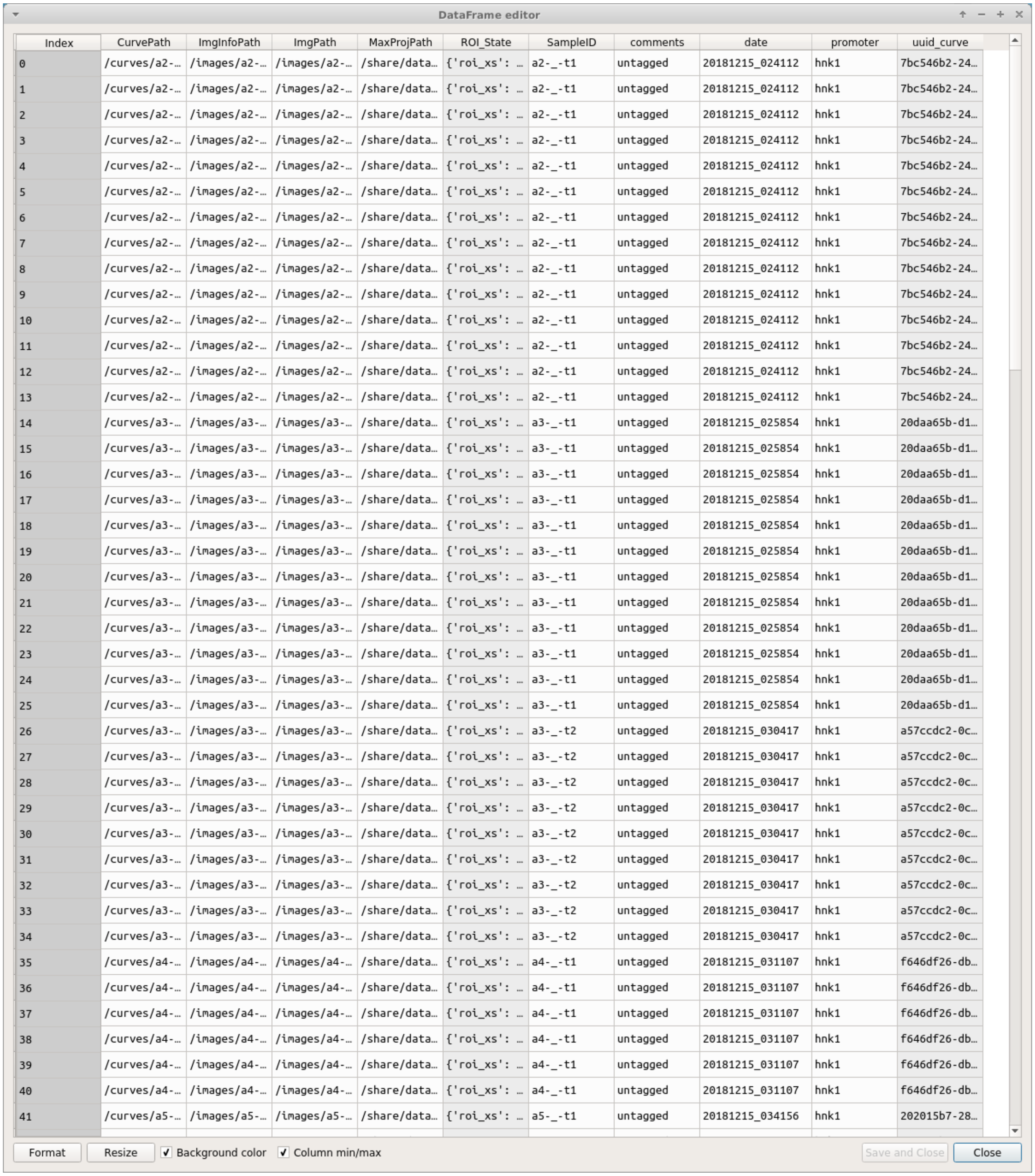
DataFrame editor

| Index | CurvePath | ImgInfoPath | ImgPath | MaxProjPath | ROI_State | SampleID | comments | date | promoter | uuid_curve |
|---|---|---|---|---|---|---|---|---|---|---|
| 0 | /curves/a2-… | /images/a2-… | /images/a2-… | /share/data… | {'roi_xs': … | a2-_-t1 | untagged | 20181215_024112 | hnk1 | 7bc546b2-24… |
| 1 | /curves/a2-… | /images/a2-… | /images/a2-… | /share/data… | {'roi_xs': … | a2-_-t1 | untagged | 20181215_024112 | hnk1 | 7bc546b2-24… |
| 2 | /curves/a2-… | /images/a2-… | /images/a2-… | /share/data… | {'roi_xs': … | a2-_-t1 | untagged | 20181215_024112 | hnk1 | 7bc546b2-24… |
| 3 | /curves/a2-… | /images/a2-… | /images/a2-… | /share/data… | {'roi_xs': … | a2-_-t1 | untagged | 20181215_024112 | hnk1 | 7bc546b2-24… |
| 4 | /curves/a2-… | /images/a2-… | /images/a2-… | /share/data… | {'roi_xs': … | a2-_-t1 | untagged | 20181215_024112 | hnk1 | 7bc546b2-24… |
| 5 | /curves/a2-… | /images/a2-… | /images/a2-… | /share/data… | {'roi_xs': … | a2-_-t1 | untagged | 20181215_024112 | hnk1 | 7bc546b2-24… |
| 6 | /curves/a2-… | /images/a2-… | /images/a2-… | /share/data… | {'roi_xs': … | a2-_-t1 | untagged | 20181215_024112 | hnk1 | 7bc546b2-24… |
| 7 | /curves/a2-… | /images/a2-… | /images/a2-… | /share/data… | {'roi_xs': … | a2-_-t1 | untagged | 20181215_024112 | hnk1 | 7bc546b2-24… |
| 8 | /curves/a2-… | /images/a2-… | /images/a2-… | /share/data… | {'roi_xs': … | a2-_-t1 | untagged | 20181215_024112 | hnk1 | 7bc546b2-24… |
| 9 | /curves/a2-… | /images/a2-… | /images/a2-… | /share/data… | {'roi_xs': … | a2-_-t1 | untagged | 20181215_024112 | hnk1 | 7bc546b2-24… |
| 10 | /curves/a2-… | /images/a2-… | /images/a2-… | /share/data… | {'roi_xs': … | a2-_-t1 | untagged | 20181215_024112 | hnk1 | 7bc546b2-24… |
| 11 | /curves/a2-… | /images/a2-… | /images/a2-… | /share/data… | {'roi_xs': … | a2-_-t1 | untagged | 20181215_024112 | hnk1 | 7bc546b2-24… |
| 12 | /curves/a2-… | /images/a2-… | /images/a2-… | /share/data… | {'roi_xs': … | a2-_-t1 | untagged | 20181215_024112 | hnk1 | 7bc546b2-24… |
| 13 | /curves/a2-… | /images/a2-… | /images/a2-… | /share/data… | {'roi_xs': … | a2-_-t1 | untagged | 20181215_024112 | hnk1 | 7bc546b2-24… |
| 14 | /curves/a3-… | /images/a3-… | /images/a3-… | /share/data… | {'roi_xs': … | a3-_-t1 | untagged | 20181215_025854 | hnk1 | 20daa65b-d1… |
| 15 | /curves/a3-… | /images/a3-… | /images/a3-… | /share/data… | {'roi_xs': … | a3-_-t1 | untagged | 20181215_025854 | hnk1 | 20daa65b-d1… |
| 16 | /curves/a3-… | /images/a3-… | /images/a3-… | /share/data… | {'roi_xs': … | a3-_-t1 | untagged | 20181215_025854 | hnk1 | 20daa65b-d1… |
| 17 | /curves/a3-… | /images/a3-… | /images/a3-… | /share/data… | {'roi_xs': … | a3-_-t1 | untagged | 20181215_025854 | hnk1 | 20daa65b-d1… |
| 18 | /curves/a3-… | /images/a3-… | /images/a3-… | /share/data… | {'roi_xs': … | a3-_-t1 | untagged | 20181215_025854 | hnk1 | 20daa65b-d1… |
| 19 | /curves/a3-… | /images/a3-… | /images/a3-… | /share/data… | {'roi_xs': … | a3-_-t1 | untagged | 20181215_025854 | hnk1 | 20daa65b-d1… |
| 20 | /curves/a3-… | /images/a3-… | /images/a3-… | /share/data… | {'roi_xs': … | a3-_-t1 | untagged | 20181215_025854 | hnk1 | 20daa65b-d1… |
| 21 | /curves/a3-… | /images/a3-… | /images/a3-… | /share/data… | {'roi_xs': … | a3-_-t1 | untagged | 20181215_025854 | hnk1 | 20daa65b-d1… |
| 22 | /curves/a3-… | /images/a3-… | /images/a3-… | /share/data… | {'roi_xs': … | a3-_-t1 | untagged | 20181215_025854 | hnk1 | 20daa65b-d1… |
| 23 | /curves/a3-… | /images/a3-… | /images/a3-… | /share/data… | {'roi_xs': … | a3-_-t1 | untagged | 20181215_025854 | hnk1 | 20daa65b-d1… |
| 24 | /curves/a3-… | /images/a3-… | /images/a3-… | /share/data… | {'roi_xs': … | a3-_-t1 | untagged | 20181215_025854 | hnk1 | 20daa65b-d1… |
| 25 | /curves/a3-… | /images/a3-… | /images/a3-… | /share/data… | {'roi_xs': … | a3-_-t1 | untagged | 20181215_025854 | hnk1 | 20daa65b-d1… |
| 26 | /curves/a3-… | /images/a3-… | /images/a3-… | /share/data… | {'roi_xs': … | a3-_-t2 | untagged | 20181215_030417 | hnk1 | a57ccdc2-0c… |
| 27 | /curves/a3-… | /images/a3-… | /images/a3-… | /share/data… | {'roi_xs': … | a3-_-t2 | untagged | 20181215_030417 | hnk1 | a57ccdc2-0c… |
| 28 | /curves/a3-… | /images/a3-… | /images/a3-… | /share/data… | {'roi_xs': … | a3-_-t2 | untagged | 20181215_030417 | hnk1 | a57ccdc2-0c… |
| 29 | /curves/a3-… | /images/a3-… | /images/a3-… | /share/data… | {'roi_xs': … | a3-_-t2 | untagged | 20181215_030417 | hnk1 | a57ccdc2-0c… |
| 30 | /curves/a3-… | /images/a3-… | /images/a3-… | /share/data… | {'roi_xs': … | a3-_-t2 | untagged | 20181215_030417 | hnk1 | a57ccdc2-0c… |
| 31 | /curves/a3-… | /images/a3-… | /images/a3-… | /share/data… | {'roi_xs': … | a3-_-t2 | untagged | 20181215_030417 | hnk1 | a57ccdc2-0c… |
| 32 | /curves/a3-… | /images/a3-… | /images/a3-… | /share/data… | {'roi_xs': … | a3-_-t2 | untagged | 20181215_030417 | hnk1 | a57ccdc2-0c… |
| 33 | /curves/a3-… | /images/a3-… | /images/a3-… | /share/data… | {'roi_xs': … | a3-_-t2 | untagged | 20181215_030417 | hnk1 | a57ccdc2-0c… |
| 34 | /curves/a3-… | /images/a3-… | /images/a3-… | /share/data… | {'roi_xs': … | a3-_-t2 | untagged | 20181215_030417 | hnk1 | a57ccdc2-0c… |
| 35 | /curves/a4-… | /images/a4-… | /images/a4-… | /share/data… | {'roi_xs': … | a4-_-t1 | untagged | 20181215_031107 | hnk1 | f646df26-db… |
| 36 | /curves/a4-… | /images/a4-… | /images/a4-… | /share/data… | {'roi_xs': … | a4-_-t1 | untagged | 20181215_031107 | hnk1 | f646df26-db… |
| 37 | /curves/a4-… | /images/a4-… | /images/a4-… | /share/data… | {'roi_xs': … | a4-_-t1 | untagged | 20181215_031107 | hnk1 | f646df26-db… |
| 38 | /curves/a4-… | /images/a4-… | /images/a4-… | /share/data… | {'roi_xs': … | a4-_-t1 | untagged | 20181215_031107 | hnk1 | f646df26-db… |
| 39 | /curves/a4-… | /images/a4-… | /images/a4-… | /share/data… | {'roi_xs': … | a4-_-t1 | untagged | 20181215_031107 | hnk1 | f646df26-db… |
| 40 | /curves/a4-… | /images/a4-… | /images/a4-… | /share/data… | {'roi_xs': … | a4-_-t1 | untagged | 20181215_031107 | hnk1 | f646df26-db… |
| 41 | /curves/a5-… | /images/a5-… | /images/a5-… | /share/data… | {'roi_xs': … | a5-_-t1 | untagged | 20181215_034156 | hnk1 | 202015b7-28… |



> **Warning:** Make sure you know what you are doing when you directly modify the Project DataFrame. Changes cannot be undone but you can restore a backup from the project's *dataframe directory*. For example, do not modify data under the following columns: CurvePath, ImgInfoPath, ImgPath, ROI_State, any uuid column.

**See also:**

Uses the Spyder object editor

##### 1.7.4 Console

If you are familiar with pandas you can interact with the project DataFrame directly. If you are unfamiliar with pandas it's very easy to learn.

**See also:**

Pandas documentation

**Useful Callables**

| Callable | Purpose |
|---|---|
| get_dataframe() | returns dataframe of the current project browser tab |
| get_root_dataframe() | always returns dataframe of the root tab (entire project DataFrame) |
| set_root_dataframe() | pass a pandas.DataFrame instance to set it as the project DataFrame |

###### Usage

General usage to modify the project DataFrame would be something like this:

```
# Get a copy the project DataFrame to modify
df = get_root_dataframe().copy()

# Do stuff to df
...

# Set the project DataFrame with the modified one
set_root_dataframe(df)
```

###### Example

Let's say you have been inconsistent in naming "ATENA" ROI Tags in the "cell_name" column. You can rename all occurances of 'atena' to 'ATENA'

```
# Get a copy of the project DataFrame
>>> df = get_root_dataframe().copy()

# View all occurances of 'atena'
>>> df.cell_name[df.cell_name == 'atena']
2      atena
3      atena
4      atena
5      atena
6      atena
205    atena
Name: cell_name, dtype: object

# Rename all occurances of 'atena' to 'ATENA'
>>> df.cell_name[df.cell_name == 'atena'] = 'ATENA'

# Check that there are more occurances of 'atena'
>>> df.cell_name[df.cell_name == 'atena']
```

(continues on next page)

(continued from previous page)

```
Series([], Name: cell_name, dtype: object)

# Check that we have renamed the 'atena' occurances to 'ATENA'
# Indices 2-6 and 205 were named 'atena'
>>> df.cell_name
0      untagged
1      untagged
2         ATENA
3         ATENA
4         ATENA
5         ATENA
6         ATENA
7         atenp
...
Name: cell_name, Length: 311, dtype: object

# Check index 205
>>> df.cell_name.iloc[205]
'ATENA'

# Finally set the changed DataFrame as the root (project) DataFrame
>>> set_root_dataframe(df)
```

#### 1.8 Viewer overview

Based on the pyqtgraph ImageView widget.

**The Viewer allows you to do the following things:**

- Examine your calcium movies
- Use modules to perform things like motion correction, CNMF(E), ROI labeling, and stimulus mapping. See their respective guides for details.
- You can also make modifications to an existing Sample in your project by opening it in the Viewer. See Modify Sample and Overwrite guide.

##### 1.8.1 Video Tutorial

This tutorial shows how to create a New Project, open images in the Viewer, use the Stimulus Mapping module and perform Caiman motion correction

##### 1.8.2 Layout

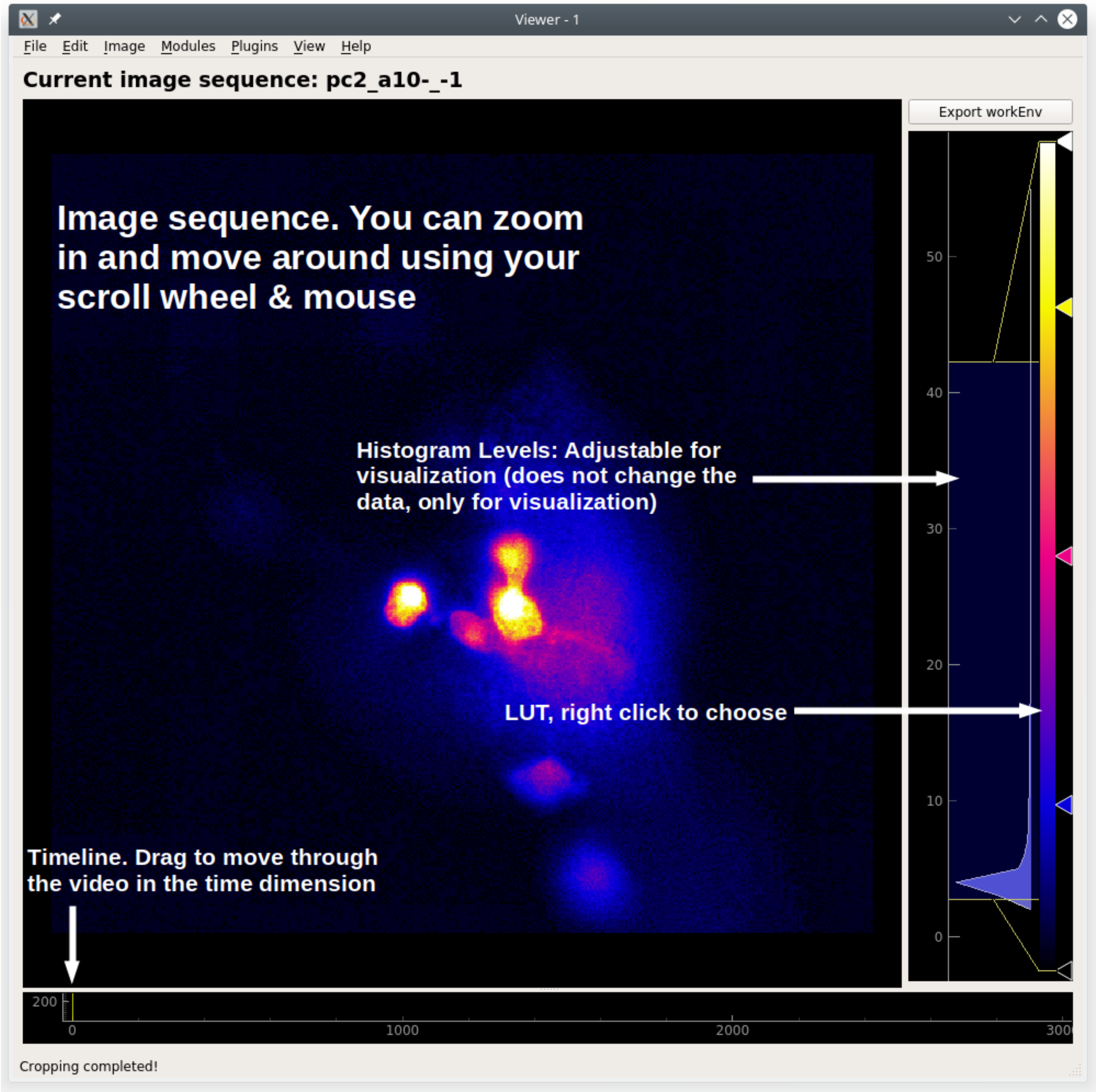


To access Viewer modules choose the module you want to run from the Modules menu at the top. All modules, except the Batch Manager, are small floating windows which you can dock into the Viewer by dragging them to an edge of the viewer.

##### 3D data

When viewing 3D data a slider on the left allows you to move through the z axis.

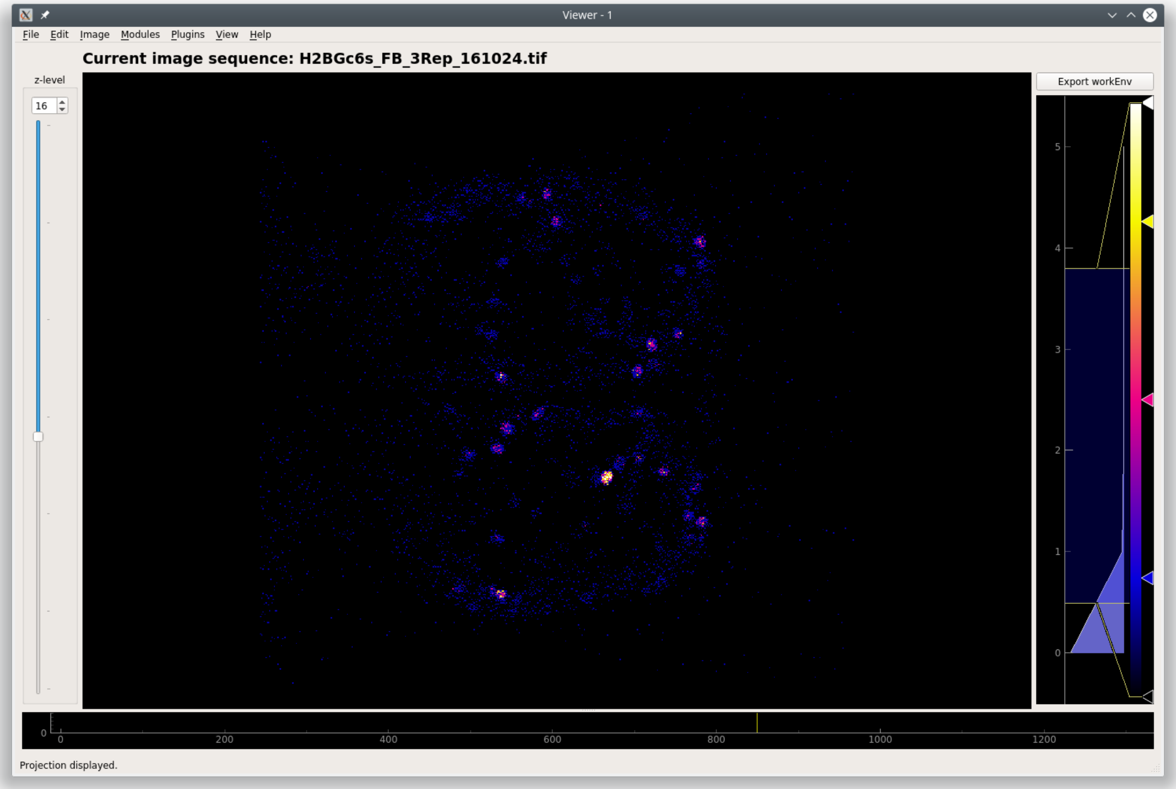


The image stack shown above is from Martin Haesemeyer's dataset from the following paper:

> Haesemeyer M, Robson DN, Li JM, Schier AF, Engert F. A Brain-wide Circuit Model of Heat-Evoked Swimming Behavior in Larval Zebrafish. Neuron. 2018;98(4):817-831.e6. doi:10.1016/j.neuron.2018.04.013

#### 1.8.3 Work Environment

Everything in the viewer is stored in a Work Environment object. The main data attributes of the viewer work environment are outlined below.

**See also:**

*ViewerWorkEnv API*

| Attribute | Description |
|---|---|
| imgdata | *ImgData object* containing the Image Sequence and meta data from the imaging source |
| roi_manager | The back-end *ROI Manager* that is currently in use |
| sample_id | SampleID, if opened from a project Sample |
| stim_maps | Stimulus maps, if any are defined |
| history_trace | History log, currently used for logging *caiman motion correction*, *CNMF* and *CNMFE* history. |
| UUID | If opened from a project Sample, it refers to the ImgUUID |

You can view everything in the current work environment by going to View -> Work Environment Editor. You cannot edit through this GUI at this time.

##### 1.8.4 Menubar

###### File

###### Add to Project

Add the current *work environment* as a Sample to the project.

###### Open Work Environment

Deprecated

###### Save Work Environment

Deprecated

###### Clear Work Environment

Clear the current *work environment*. Useful for freeing up RAM.

###### Edit

Deprecated

###### Image

###### Reset Scale

Reset the scale of the image ViewBox

###### Resize

Resize the image sequence using interpolation.

##### Crop

Crop the image sequence.

**Usage**

1. When you click this option a square crop region will appear in the top left corner of the image sequence.
2. You can change its shape using the handle in the bottom right corner.
3. To crop to the selection, in the menubar go to Image -> Crop. To cancel cropping right click in the crop region and click "Remove ROI".

##### Measure

Measure the distance (in pixels) between two points in the image sequence.

**Usage**

1. After clicking this option in the menubar, click on a point in the image sequence. You will not see anything yet.
2. Click on a second point in the image sequence, a line will appear connecting the first and second point that you clicked.
3. You can use the handles at the endpoints of the line to change the line.
4. The displacement in the x, y, and along the line will be displayed in the status bar (bottom left corner of the Viewer Window) when you hover over a measuring line.
5. You can create as many measuring lines as you want.
6. To remove a measuring line, right click on a handle and click "Remove ROI"

##### Change dtype

Not implemented yet. You can change the dtype through the console.

##### Projections

View Mean, Max, and Standard Deviation projections of the current image sequence in the work environment. If the data are 3D, the projection is of the current plane.

##### Modules

Default Viewer Modules. These are explained in more details in the Viewer Modules chapters.

##### Plugins

Custom viewer modules.

##### View

##### Work Envionment Editor

Explore the data in your work environment using a GUI.

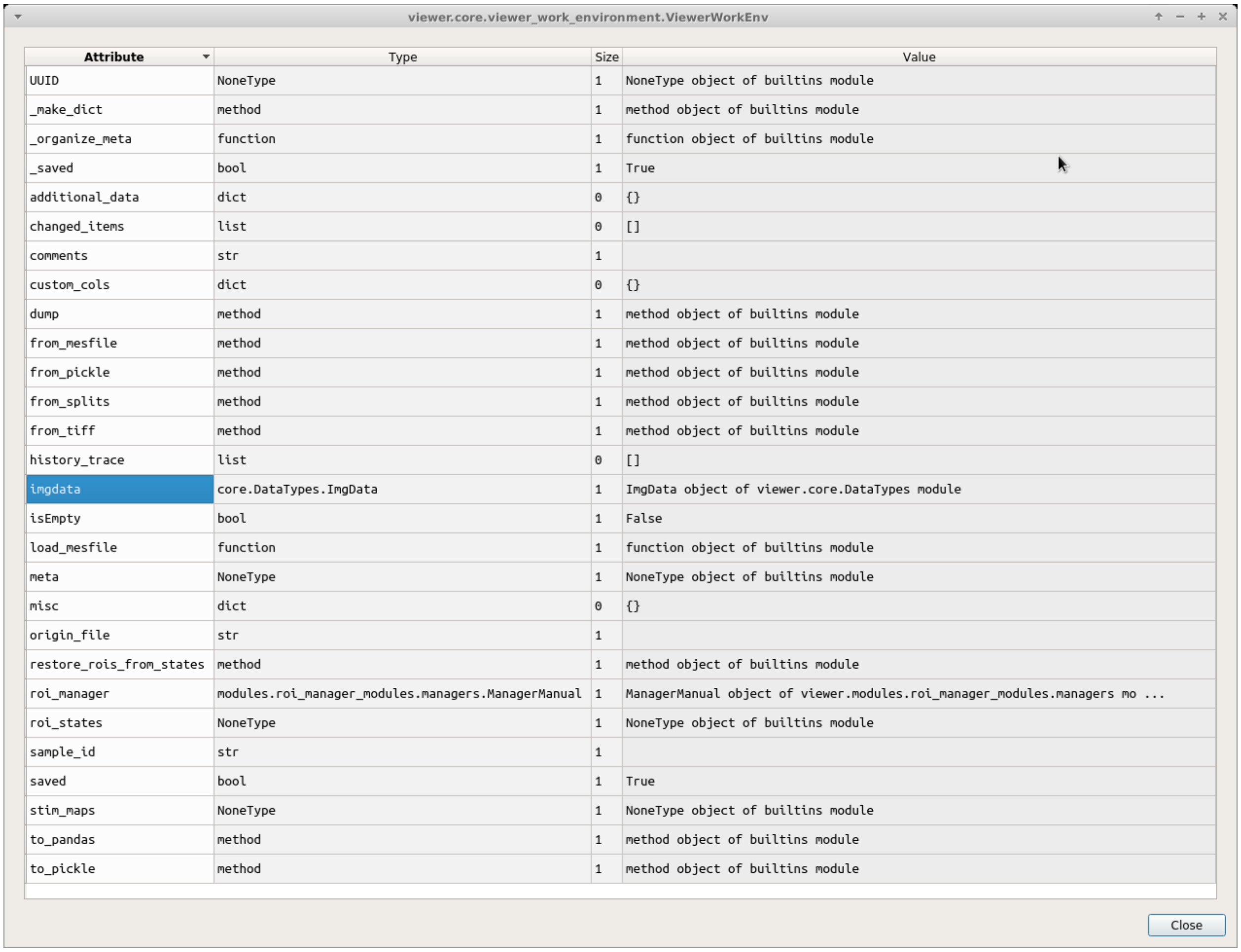


| Attribute | Type | Size | Value |
|---|---|---|---|
| UUID | NoneType | 1 | NoneType object of builtins module |
| _make_dict | method | 1 | method object of builtins module |
| _organize_meta | function | 1 | function object of builtins module |
| _saved | bool | 1 | True |
| additional_data | dict | 0 | {} |
| changed_items | list | 0 | [] |
| comments | str | 1 | |
| custom_cols | dict | 0 | {} |
| dump | method | 1 | method object of builtins module |
| from_mesfile | method | 1 | method object of builtins module |
| from_pickle | method | 1 | method object of builtins module |
| from_splits | method | 1 | method object of builtins module |
| from_tiff | method | 1 | method object of builtins module |
| history_trace | list | 0 | [] |
| imgdata | core.DataTypes.ImgData | 1 | ImgData object of viewer.core.DataTypes module |
| isEmpty | bool | 1 | False |
| load_mesfile | function | 1 | function object of builtins module |
| meta | NoneType | 1 | NoneType object of builtins module |
| misc | dict | 0 | {} |
| origin_file | str | 1 | |
| restore_rois_from_states | method | 1 | method object of builtins module |
| roi_manager | modules.roi_manager_modules.managers.ManagerManual | 1 | ManagerManual object of viewer.modules.roi_manager_modules.managers mo ... |
| roi_states | NoneType | 1 | NoneType object of builtins module |
| sample_id | str | 1 | |
| saved | bool | 1 | True |
| stim_maps | NoneType | 1 | NoneType object of builtins module |
| to_pandas | method | 1 | method object of builtins module |
| to_pickle | method | 1 | method object of builtins module |

**Note:** This is read only, you cannot edit via this GUI.

For example if you want to see your meta data, double click on "imgdata" and then you can see that "imgdata" has two things, the image sequence (i.e. your video) and the meta data.

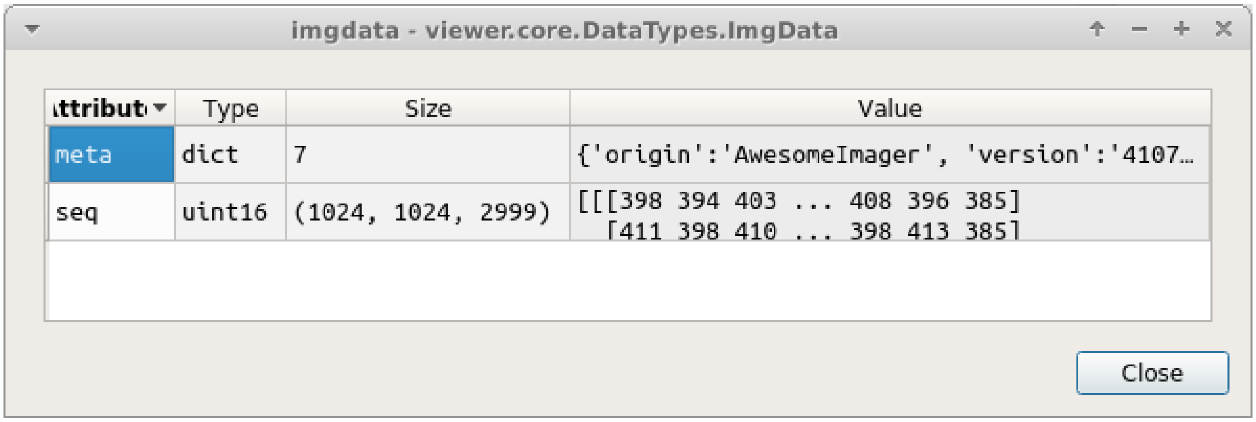


If you double click on "meta" above you can see your meta data.

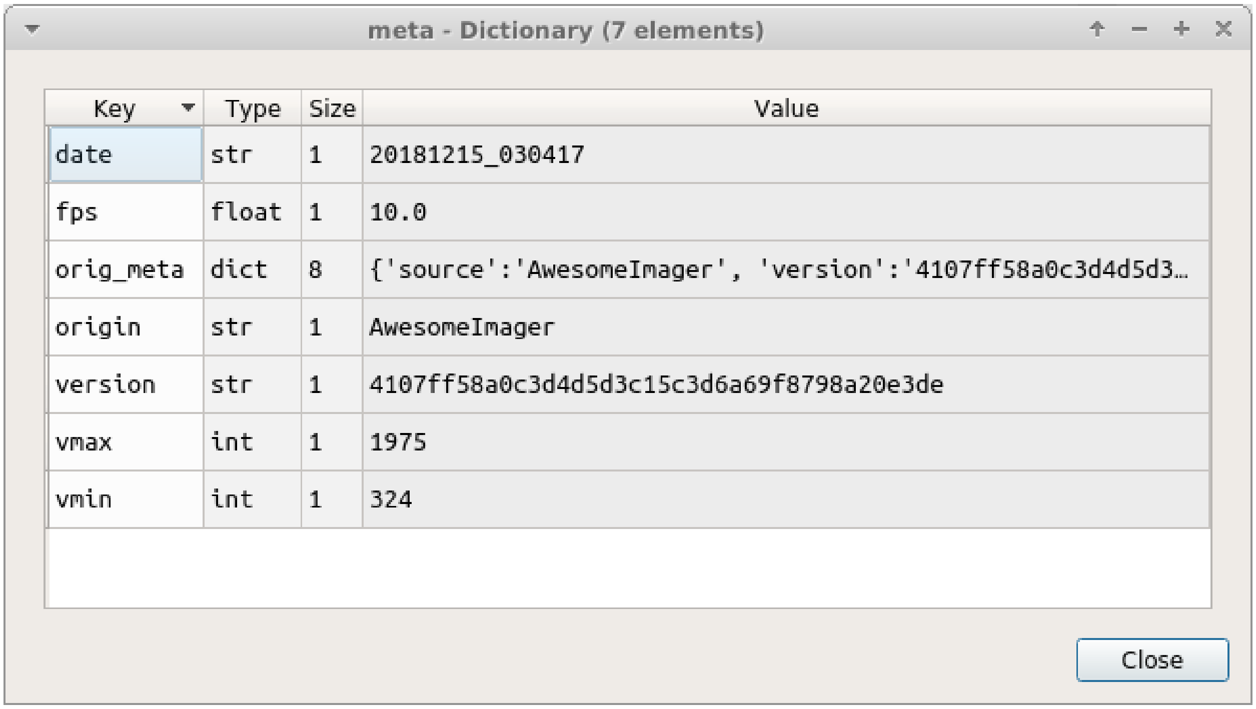


You can view your meta data more quickly using the console.

Open the console by going to View -> Console. You can then call `get_meta()` to print the meta data dict.

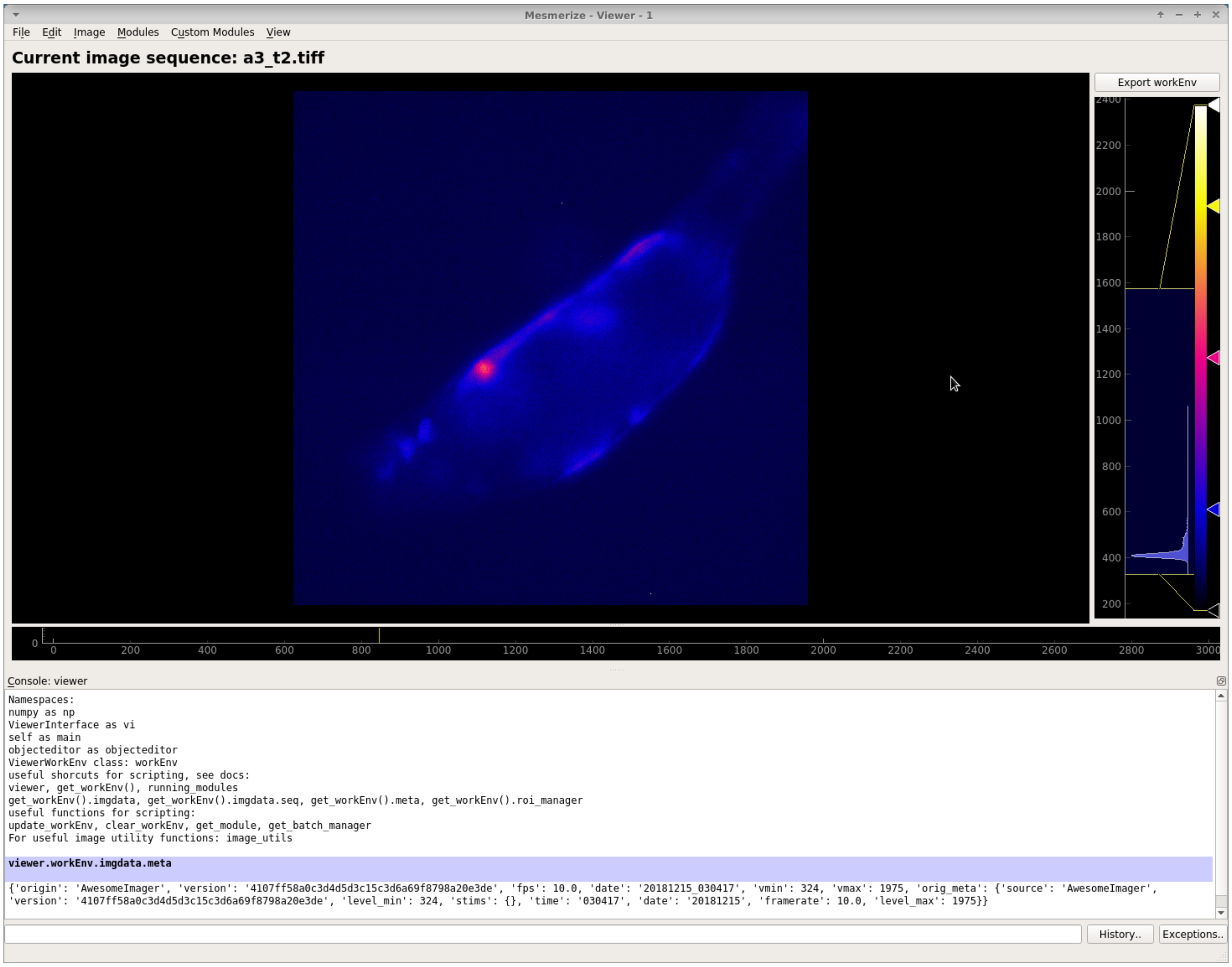


#### Console

View/hide the viewer console

#### Help

##### Open docs

Open these docs

##### 1.8.5 Console

You can interact directly with the *work environment* using the console.

**See also:**

*Viewer Core API*, *Overview on consoles*

###### Namespace

| Reference | Description |
|---|---|
| vi | Instance of *ViewerUtils*. Use this to interact with the viewer. |
| all_modules | List all available modules (includes default and any available plugins/custom modules) |
| Viewer-WorkEnv | Use for creating new instances of *ViewerWorkEnv* |
| ImgData | Use for creating new instances of *ImgData* |
| get_workEnv() | Get the current viewer *work environment* (instance of *ViewerWorkEnv*) |
| get_image() | Get the current image sequence (returns current *ViewerWorkEnv.imgdata.seq*). If the data are 3D it returns the current plane only. |
| get_meta() | Get the current meta data |
| get_module(<name>) | Pass the name of a module as a string. Returns that module if it is available. |
| get_batch_manager() | Get the batch manager. |
| up-date_workEnv() | Update the viewer GUI with the viewer work environment (vi.viewer.workEnv) |
| clear_workEnv() | Clear the current work envionment, cleanup the GUI and free the RAM |

###### Video Tutorial

###### Examples

These examples can be run in the *Viewer Console*.

###### Working with meta data

```
# view meta data
>>> get_meta()

{'origin': 'AwesomeImager', 'version': '4107ff58a0c3d4d5d3c15c3d6a69f8798a20e3de', 'fps
↪': 10.0, 'date': '20190426_152034', 'vmin': 323, 'vmax': 1529, 'orig_meta': {'source':
↪'AwesomeImager', 'version': '4107ff58a0c3d4d5d3c15c3d6a69f8798a20e3de', 'level_min':␣
↪323, 'stims': {}, 'time': '152034', 'date': '20190426', 'framerate': 10.0, 'level_max
↪': 1529}}

# manually set meta data entries
>>> get_meta()['fps'] = 30.0
```

##### Open image

Use the *Viewer Core API* to open any arbitrary image

This example loads an image stored using numpy.save(), but this is applicable to images stored in any format that can eventually be represented as a numpy array in python. For example, you could also load image files stored in HDF5 format and load the numpy array that represents your image sequence.

```
import numpy as np

# clear the viewer work environment
clear_workEnv()

a = np.load('/path_to_image.npy')

# check what the axes order is
a.shape

# (1000, 512, 512) # for example
# looks like this is in [t, x, y]
# this can be transposed so we get [x, y, t]
# ImgData takes either [x, y, t] or [x, y, t, z] axes order

# Define a meta data dict
meta = \
    {
        "origin":      "Tutorial example",
        "fps":         10.0,
        "date":        "20200629_171823",
        "scanner_pos": [0, 1, 2, 3, 4, 5, 6]
    }

# Create ImgData instance
imgdata = ImgData(a.T, meta)  # use a.T to get [x, y, t]

# Create a work environment instance
work_env = ViewerWorkEnv(imgdata)

# Set the current Viewer Work Environment from this new instance
vi.viewer.workEnv = work_env

# Update the viewer with the new work environment
# this MUST be run whenever you replace the viewer work environment (the previous line)
update_workEnv()
```

##### Image data

Image sequences are simply numpy arrays. For example extract the image sequence between frame 1000 and 2000.

**See also:**

Numpy array indexing

```python
# Get the current image sequence
seq = get_image()

# Trim the image sequence
trim = seq[:, :, 1000:2000]

# Set the viewer work environment image sequence to the trim one
vi.viewer.workEnv.imgdata.seq = trim

# Update the GUI with the new work environment
update_workEnv()
```

##### View analysis log

View the analysis log, such as batch manager processing history.

```
>>> get_workEnv().history_trace

[{'caiman_motion_correction': {'max_shifts_x': 32, 'max_shifts_y': 32, 'iters_rigid': 1, 'name_rigid': 'Does not matter', 'max_dev': 20, 'strides': 196, 'overlaps': 98, 'upsample': 4, 'name_elas': 'a1_t2', 'output_bit_depth': 'Do not convert', 'bord_px': 5}}, {'cnmfe': {'Input': 'Current Work Environment', 'frate': 10.0, 'gSig': 10, 'bord_px': 5, 'min_corr': 0.9600000000000001, 'min_pnr': 10, 'min_SNR': 1, 'r_values_min': 0.7, 'decay_time': 2, 'rf': 80, 'stride': 40, 'gnb': 8, 'nb_patch': 8, 'k': 8, 'name_corr_pnr': 'a8_t1', 'name_cnmfe': 'a1_t2', 'do_corr_pnr': False, 'do_cnmfe': True}}, {'cnmfe': {'Input': 'Current Work Environment', 'frate': 10.0, 'gSig': 10, 'bord_px': 5, 'min_corr': 0.9600000000000001, 'min_pnr': 14, 'min_SNR': 1, 'r_values_min': 0.7, 'decay_time': 4, 'rf': 80, 'stride': 40, 'gnb': 8, 'nb_patch': 8, 'k': 8, 'name_corr_pnr': '', 'name_cnmfe': 'a1_t2', 'do_corr_pnr': False, 'do_cnmfe': True}}]
```

##### Running scripts

You can use the *Script Editor* to run scripts in the Viewer console for automating tasks such as batch creation. It basically allows you to use the *viewer console* more conveniently with a text editor. The execution environment of the viewer console and script editor are identical.

Some example are provided for caiman modules and *stimulus mapping*.

#### 1.9 Add a Sample to the Project

When you are happy with the ROIs in the viewer for the current CNMF(E) derived or manually created ROIs, you can add this as a *Sample* to your project.

Each sample in your project contains the following:

- The imaging data from which ROIs were extracted (the video)
- All the ROIs with their spatial location, temporal dynamics, and any tags that you have entered in the ROI Manager.
- Stimulus mappings, if your project is configured for this.
- Meta data (that were associated with the imaging video), the date, video framerate.
- Any further information that you have chosen to add based on your Project Configuration

**Note:** If your ROIs were obtained through CNMF/CNMFE the following attributes from the final cnm object are stored: cnm.A, cnm.b, cnm.C, cnm.f, cnm.YrA

##### 1.9.1 How to

To add the current *viewer work environment* (see above) as a sample to your project, go to File -> Add To Project. You will be presented with a window similar to this:

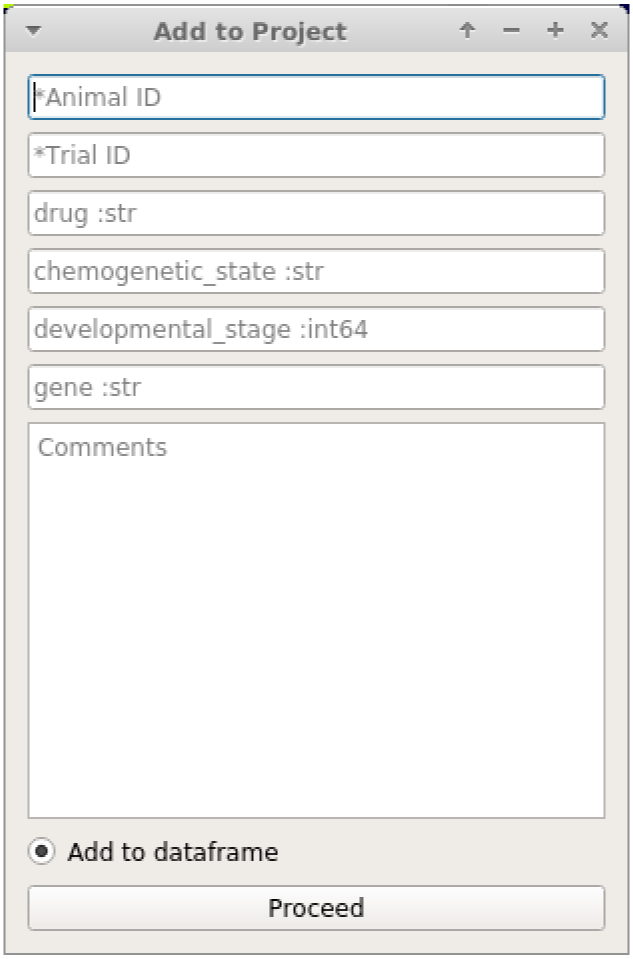


The entries that you are prompted with directly correspond to the custom columns in your Project Configuration.

**See also:**

*Project Configuration*

Every Sample in a project has a unique **SampleID** which is the combination of **AnimalID** + **TrialID**.

> **Warning: You can never change the AnimalID or TrialID (i.e. SampleID) since these are partially used as unique identifiers.** A workaround is described in the *FAQ for Project Organization*.

> **Warning:** AnimalID and TrialID are separated by the `-_-` character combination when stored as a SampleID. Therefore do not use that character combination within your AnimalID or TrialID.

##### 1.9.2 Video Tutorial

#### 1.10 Tiff file module

To open a tiff file go to Modules -> Load Images -> Tiff files.

**Note:** You can also use this module through the console and scripts. See *Tiff module API*.

To open tiff files first click the "Select file" button and choose your file. You can also drag and drop a tiff file (drag and drop doesn't work properly on Windows).

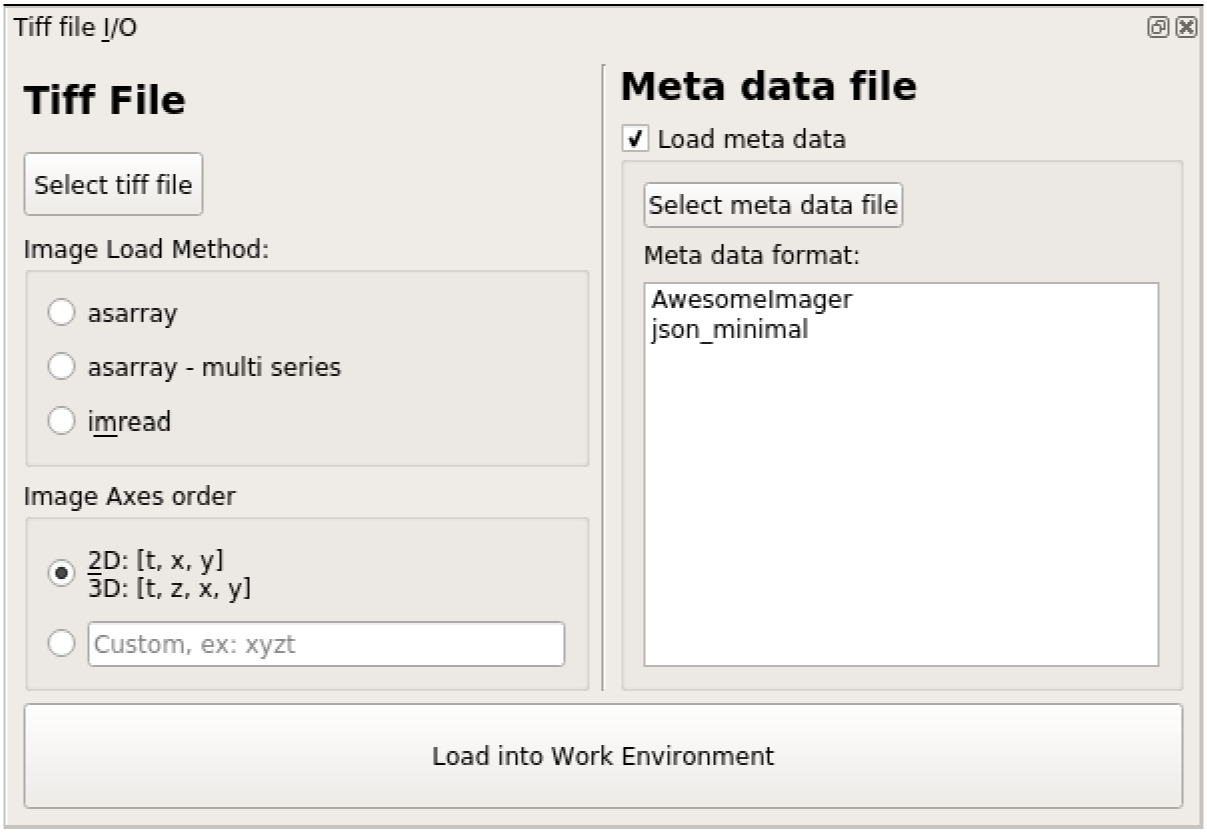


Next, you must select an appropriate Image Load Method (see next section). You can also import meta data associated with your recording.

**Certain meta data, such as the sampling rate of the data, are necessary for some downstream analysis procedures.** There are a few ways to import your meta data into the Viewer Work Environment:

- Simple JSON files, see *json_minimal* under the table in the *Meta data* section
- Define your own *Custom functions* to open meta in other file formats
- Manually create a meta data dictionary using the *Console*

##### 1.10.1 Load Method

The options for "Load Method" correspond to the tifffile library method that is used for loading the images.

If you are not sure which method you should use, try all of them and see which one loads your data appropriately. If none of them work, create an issue on GitHub.

- **asarray:** Should work for most tiff files, fast method
- **asarray - multi series:** Also fast. Use this if it is a multi-page tiff. For example if the tiff file was created by a program that appends each frame to a file as they are being acquired by the camera.
- **imread:** Usually slower, should work for most tiff files.

##### 1.10.2 Axes order

Choose the default axes order or manually enter the axes order if your tiff file uses a different order.

##### 1.10.3 Meta data

Check the "Load meta data" checkbox if you want to load meta data. Alternatively, you can uncheck this box and create a meta data dictionary manually using the console (see the *Console* section)

You can select a meta data format from the list. This list of formats correspond to the functions available in the module: `mesmerize.viewer.core.organize_meta`. When you select a meta data format, it will automatically try to find a file with the extension specified by the selected format if it has the same name as the selected tiff file.

If you have questions on meta data formats feel free to drop a message in the Gitter room

**Default list of formats that are recognized:**

| Name | extension | Description |
|---|---|---|
| json_minimal | .json | Recognizes a json file that contains at least the minimal set of necessary keys: `origin`, `fps` and `date`.<br><br>All other keys in the JSON file are placed in a sub-dictionary with the key `orig_meta`<br><br>See *Minimal dict* below for more info. |
| AwesomeImager | .json | Used for 1p imaging in the Chatzigeorgiou group at the Sars Center |

##### Custom functions

You may define your own function to organize your meta data. It MUST return a dict which has at least the following keys: `origin`, `fps` and `date`.

- `origin` is a `str` describing the software or microscope the recording comes from. This is for your own record.
- `fps` is the sampling rate of the recording as a `float` or `int`
- `date` is the date & time represented by a `str` in the following format: `"YYYYMMDD_HHMMSS"`

In addition to these 3 keys, you may include any additional keys as you wish.

If you think your meta data organizing function will be useful for others I'll be happy to review a pull request and it can be included by default in Mesmerize. We're happy help you create a meta data function, just contact us on Gitter or create an issue on GitHub.

##### Minimal dict

Example of a minimal meta data dict.

```
{
    "origin": "microscope or software origin",  # must be a str
    "fps":     10.0,                            # must be a int or float
    "date":    "20201123_172345"                # must be a str formatted as "YYYYMMDD_
↪HHMMSS"
}
```

##### Function outline

Basic outline of a function that you can add to `mesmerize.viewer.core.organize_meta` for organizing your meta data:

1. The function can only take the `path` to the meta data file as the argument.
2. The expected file extension for the meta data must be specified. The files of a single format are allowed to have multiple different file extension but you must only specify the most common one.
3. The function would generally open the meta data file specified by the `path`, using any python libraries or other code of your choice, and finally return a dictionary that contains the minimal complement of keys, i.e. `origin`, `fps` and `date` with values of the appropriate types (see previous section).

```
def my_meta_organizer(path: str) -> dict:
    """.ext""" # define the file ext in the docstring

    raw_meta = function_to_load_my_file(path)

    # do stuff to organize the raw_meta

    meta = ... # stuff to organize raw meta
    return meta
    # return the organized meta data dict
    # that mesmerize can use
```

##### 1.10.4 Console/Script usage

You can also load tiff files through the *Viewer Console* or *Script Editor*.

This example can be run line-by-line through the *Viewer Console*, or from the *Script Editor*.

```
image_path = # path to tiff file
meta_path = # path to json meta data file

clear_workEnv() # Prevents a confirmation dialog from appearing

# Get the tiff module
tio = get_module('tiff_io', hide=True)

# Load the tiff file
tio.load(image_path, method='imread', axes_order='txy', meta_path=meta_path, meta_format=
↪'json_minimal)
```

**Alternatively, you may manually create a meta data dictionary after loading a tiff file:**

```
image_path = # path to tiff file

clear_workEnv() # Prevents a confirmation dialog from appearing

# Get the tiff module
tio = get_module('tiff_io', hide=True)

# Load the tiff file
tio.load(image_path, method='imread', axes_order='txy')

meta_dict = \
    {
        "origin":   "my_microscope_software",   # must a str
        "fps":      17.25,                      # must be a int or float
        "date"      "20201123_172345"           # must be a str formatted as "YYYYMMDD_
↪HHMMSS"/
    }

get_workEnv().imgdata.meta = meta_dict
```

**See also:**

*Tiff module API*, *Viewer Core API*, *Overview on consoles*

#### 1.11 Batch Manager

**Batch process computationally intensive tasks.**

**See also:**

*Batch Manager API*

##### 1.11.1 Video Tutorial

This tutorial shows how to create a New Project, open images in the Viewer, use the Stimulus Mapping module and perform Caiman motion correction

This is currently used for *Caiman Motion Correction*, *CNMF*, *CNMF 3D* and *CNMFE*.

The Batch Manager can be accessed in the viewer through Modules -> Batch Manager. If you don't have a batch open you will be prompted with a dialog to open a batch or to select a location for a new batch.

> **Warning:** The full path to the batch directory must not contain spaces or special characters, only a-z, A-Z, 0-9 and underscores.

The Batch Manager processes the batch items in external processes, allowing you to add batch items when that batch is being processed.

##### 1.11.2 Layout

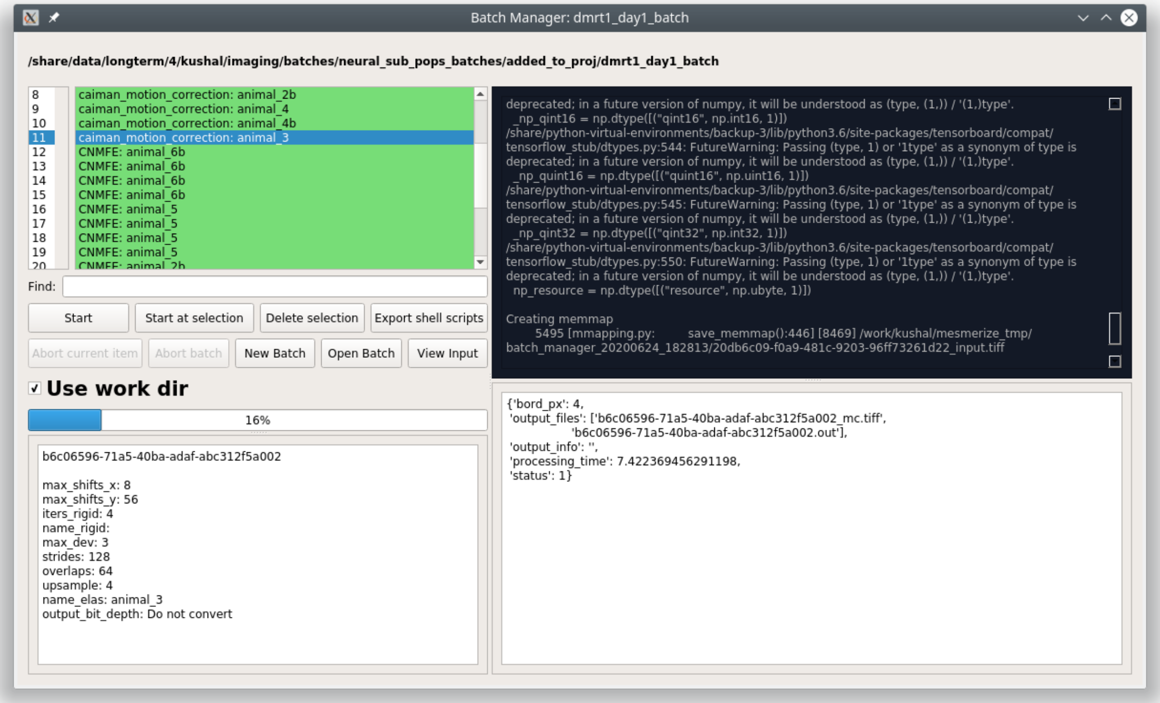


Window title: Name of batch directory

**Top:** Parent directory of batch directory

**Top left:** list of batch items and some controls.

| Colors | Description |
|---|---|
| Green | Finished without exceptions |
| Red | Did not finish, click on the item to see the exceptions in the bottom right information area |
| Yellow | Currently being processed |
| Orange | Item aborted by user |
| Blue | Output data for this item are being moved from the work dir to the batch dir. |

| Button | Description |
|---|---|
| Start | Process the batch from the first item. |
| Start at selection | Process the batch starting from the item that is currently selected in the list. |
| Delete selection | Delete the item that is currently being selected along with the associated data in the batch dir. |
| Export shell scripts | Export bash scripts so that the batch items can be run on a computing cluster |
| Abort current item | Abort the current batch item and move on to the next item |
| Abort batch | Abort the current item and stop processing the batch |
| New batch | Create a new batch |
| Open batch | Open a batch |
| View Input | Open the input work environment, in the viewer, for the currently selected item |

**Use work dir:** Check this box to use the work dir that has been set in the *System Configuration*. This feature is only available on Linux & Mac OSX.

**Top right:** Standard out from the external processes that are processing the batch items.

**Bottom left:** Parameters for the selected batch item. The first line is the UUID of the batch item.

**Bottom right:** Output information area for the currently selected item.

##### 1.11.3 Scheduling

You can schedule a batch to run at a later time using the following bash script. Doesn't work for a snap installation yet.

`mesmerize-scheduler`

**Usage:**

```
Usage: mesmerize-scheduler -b <batch> -i <start item> -t <start time>

    -b      full batch path in quotes, no spaces
    -i      uuid of the batch item to start from, no quotes
    -t      time at which to start the batch, no quotes

    examples of how to specify time:
            23:00  7:30Feb30
            use 24hr time and no spaces

Full usage example:
    mesmerize-scheduler -b "/share/data/temp/kushal/pc2_batch" -i a80d1923-e490-4eb3-
↪ba4f-7e651d4cf938 -t 2:00
```

#### 1.12 Stimulus Mapping

*API Reference*

##### 1.12.1 Video Tutorial

This tutorial shows how to create a New Project, open images in the Viewer, use the Stimulus Mapping module and perform Caiman motion correction

**Map temporal information such as stimulus or behavioral periods.**

Stimulus Mapping Module

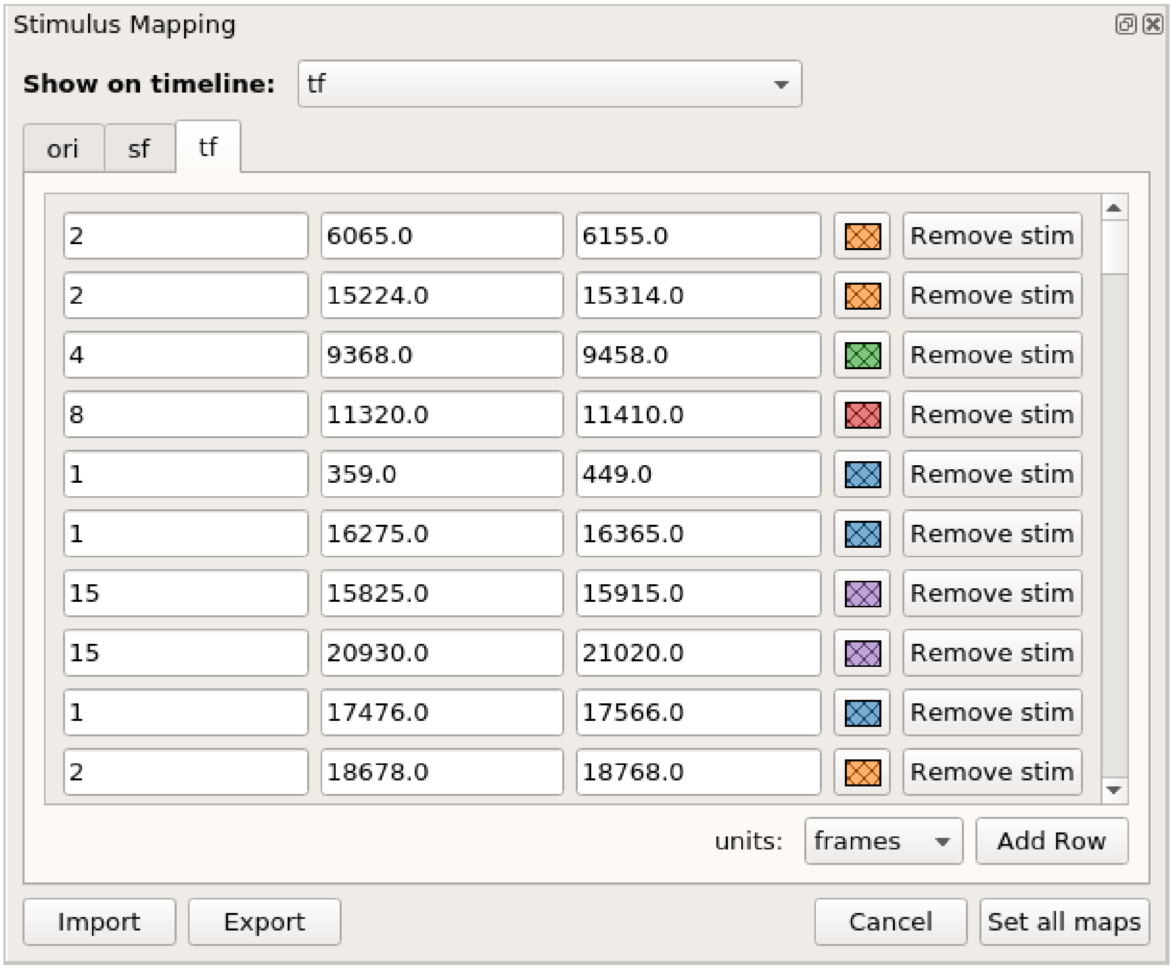


Stimulus periods illustrated on the viewer timeline

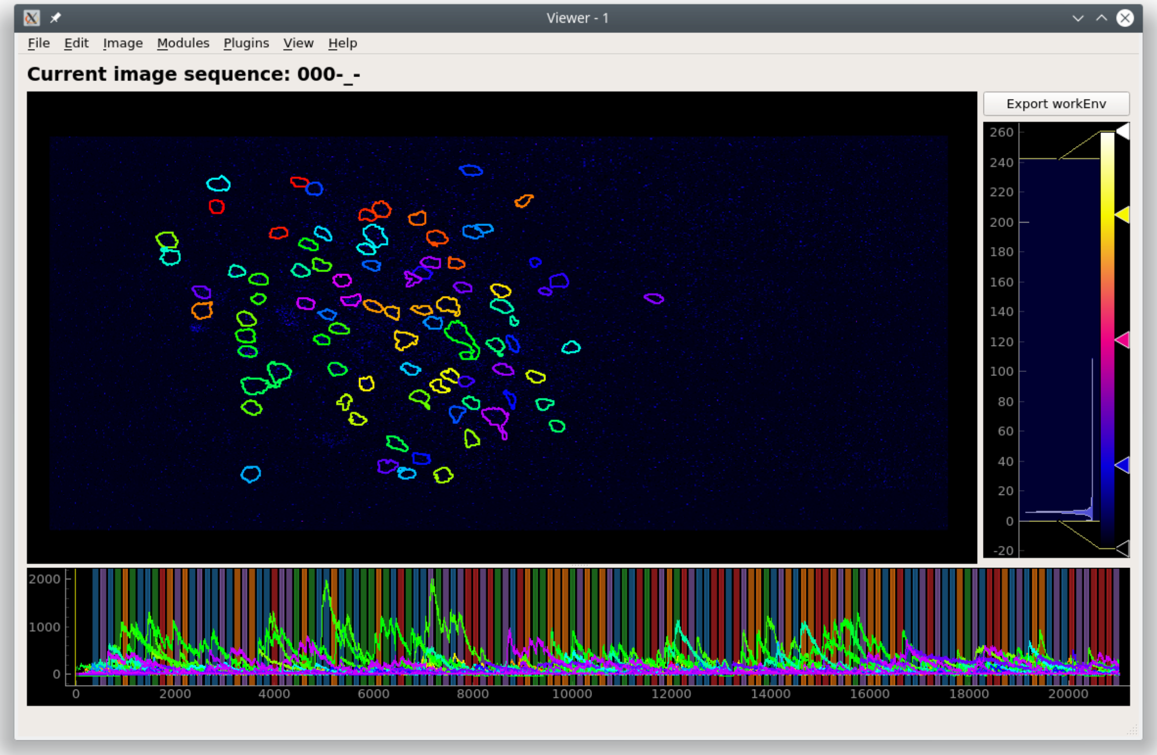


The tabs that are available in the stimulus mapping module corresponds to the stimulus types in your *Project Configuration*.

You can add stimulus periods either manually or through a script.

#### 1.12.2 Manual Annotation

1. To add a stimulus manually click the "Add Row" button. This will add an empty row to the current tab page.
2. Enter a name for the stimulus, start time, end time, and pick a color for illustrating the stimulus periods on the Viewer timeline.
3. To remove a stimulus click the "Remove stim" button. Stimulus periods do not have to be added in chronological order.
4. Click "Set all maps" to set the mappings for all stimulus types. You can then choose to illustrate a stimulus on the viewer timeline by selecting it from "Show on timeline"

Import and Export are not implemented yet.

**Warning:** At the moment, only "frames" are properly supported for the time units.

**Note:** It is generally advisable to keep your stimulus names short with lowercase letters. When sharing your project you can provide a mapping for all your keys. This helps maintain consistency throughout your project and makes the data more readable.

##### 1.12.3 Script

**See also:**

*API Reference*

You can also use the *Stimulus Mapping module's API* to set the stimulus mappings from a pandas DataFrame.

This example creates a pandas DataFrame from a csv file to set the stimulus mappings. It uses the csv file from the pvc-7 dataset availble on CRCNS: http://dx.doi.org/10.6080/K0C8276G

You can also download the csv here: `stimulus_pvc7.csv`

This example is meant to be run through the *Viewer Script Editor*

```
import pandas as pd
from mesmerize.plotting.utils import get_colormap

# Load dataframe from CSV
df = pd.read_csv('path_to_csv_file')

# Sort according to time
df.sort_values(by='start').reset_index(drop=True, inplace=True)

# Trim off the stimulus periods that are not in the current image sequence
trim = get_image().shape[2]
df = df[df['start'] <= trim]

# get one dataframe for each of the stimulus types
ori_df = df.drop(columns=['sf', 'tf', 'contrast'])  # contains ori stims
sf_df = df.drop(columns=['ori', 'tf', 'contrast'])  # contains sf stims
tf_df = df.drop(columns=['sf', 'ori', 'contrast'])  # contains tf stims

# Rename the stimulus column of interest to "name"
ori_df.rename(columns={'ori': 'name'}, inplace=True)
sf_df.rename(columns={'sf': 'name'}, inplace=True)
tf_df.rename(columns={'tf': 'name'}, inplace=True)


# Get the stimulus mapping module
smm = get_module('stimulus_mapping')

# set the stimulus map in Mesmerize for each of the 3 stimulus types
for stim_type, _df in zip(['ori', 'sf', 'tf'], [ori_df, sf_df, tf_df]):
    # data in the name column must be `str` type for stimulus mapping module
    _df['name'] = _df['name'].apply(str)

    # Get the names of the stimulus periods
    stimuli = _df['name'].unique()
    stimuli.sort()

    # Create colormap with the stimulus names
    stimuli_cmap = get_colormap(stimuli, 'tab10', output='pyqt', alpha=0.6)

    # Create a column with colors that correspond to the stimulus names
    # This is for illustrating the stimulus periods in the viewer plot
```

(continues on next page)

(continued from previous page)

```
    _df['color'] = _df['name'].map(stimuli_cmap)

    # Set the data in the Stimulus Mapping module
    smm.maps[stim_type].set_data(_df)
```

#### 1.13 ROI Manager

*API Reference*

**Manage and annotate ROIs**

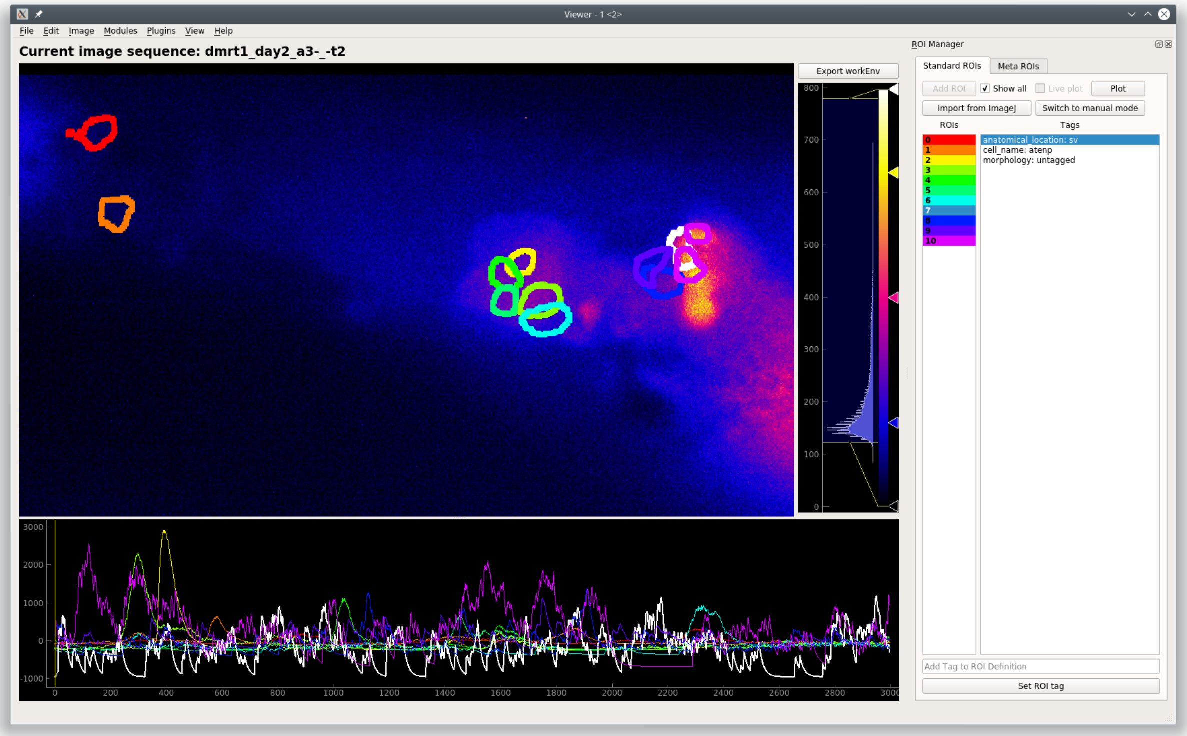


The ROI Manager has a manual mode, to draw ROIs manually, and a CNMF(E) mode where ROIs can be imported from CNMF(E) outputs.

**See also:**

*CNMF*, *CNMF 3D*, and *CNMFE*.

**Note:** You cannot combine manual and CNMF(E) ROIs in the same sample.

The ImageJ ROI import uses the read-roi package by Hadrien Mary https://pypi.org/project/read-roi/

##### 1.13.1 Video Tutorial

##### 1.13.2 Layout

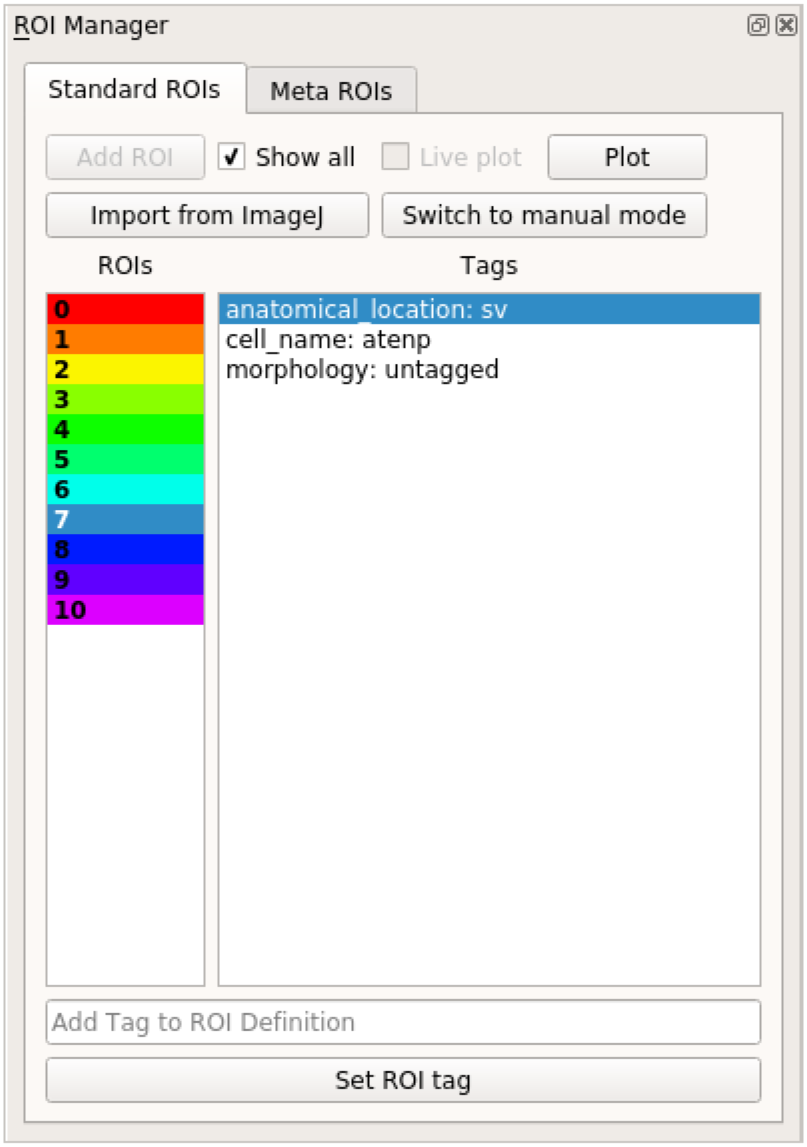


**Controls**

| UI | Description |
|---|---|
| Add ROI button | Add Polygon ROI (Manual mode)<br><br>Right click this button to add an elliptical ROI |
| Show all | Show all ROIs in the viewer |
| Live plot | Live update of the curve plot with changes (Manual mode) |
| Plot | Plot the curves (Manual mode) |
| Import from ImageJ | Import ROIs from an ImageJ ROIs zip file (Manual mode) |
| Switch to manual … | Switch to Manual mode. Clears CNMF(E) ROIs. |
| ROIs list | Color-coded list of ROIs.<br><br>Left click to highlight the ROI in the viewer<br><br>Right click to show the context menu allowing you to delete the selected ROI |
| Tags list | List of tags for the selected ROI<br><br>Correspond to the *ROI Type Columns of the Project Configuration* |
| Add Tag to ROI Def… | Set the tag for the current selection in the Tags list |
| Set ROI Tag | Click to set the tag, or just press return in the text entry above |

**Note:** It is generally advisable to keep your ROI tags short with lowercase letters. When sharing your project you can provide a mapping for all your keys. This helps maintain consistency throughout your project and makes the data more readable.

**Note:** When using 3D data, the ROIs are colored randomly along the list (not linearly as shown in the image). If you want to set the colors linearly call this in the Viewer Console: `get_workEnv().roi_manager.roi_list.reindex_colormap(random_shuffle=False)`

**Warning:** Importing several *thousands* of ROIs can take 15-30 minutes. You will be able to track the progress of the import in the Viewer Window's status bar.

**Keyboard shortcuts**.

These only work when the ROI manager is docked within the Viewer and while you are typing in the *Add Tag to ROI Definition* text entry.

| Key | Description |
|---|---|
| Page Up | Select previous ROI |
| Page Down | Select next ROI |
| Right Arrow | Play the video at high speed |
| Left Arrow | Play the video backwards at high speed |
| Home | Go to the beginning of the video |
| End | Go to the end of the video |

##### 1.13.3 Manual ROI Mode

When you click the "Add ROI" button to add a Manual Polygon ROI, a new rectangular ROI will be add in the top left corner of the image. You can add new vertices to this polygon by clicking on any of its edges. You can drag the vertices to change the shape of the polygon, and you can drag the entire ROI as well by clicking and dragging within the ROI region. Similarly you can reshape elliptical ROIs.

Hovering over the ROI selects it in the ROI list.

##### 1.13.4 Console

These examples can be run through the viewer console or *Script editor* to interact with the ROIs.

**See also:**

*Back-end ROI Manager APIs*, *ROIList API*, *ROI Type APIs*

Get the back-end ROI Manager, see *ROI Manager APIs*

```
>>> get_workEnv().roi_manager

<mesmerize.viewer.modules.roi_manager_modules.managers.ManagerCNMFROI object at␣
↪0x7f01b8780668>``
```

Get the ROI List, see *ROIList API*

```
>>> get_workEnv().roi_manager.roi_list

[<mesmerize.viewer.modules.roi_manager_modules.roi_types.CNMFROI object at␣
↪0x7f01bc78b278>, <mesmerize.viewer.modules.roi_manager_modules.roi_types.CNMFROI␣
↪object at 0x7f01bc817630>, <mesmerize.viewer.modules.roi_manager_modules.roi_types.
↪CNMFROI object at 0x7f01bc817668>, <mesmerize.viewer.modules.roi_manager_modules.roi_
↪types.CNMFROI object at 0x7f01bc7c5438>, <mesmerize.viewer.modules.roi_manager_modules.
↪roi_types.CNMFROI object at 0x7f01bc7c5208>]
```

Work with an ROI object, see *ROI Type APIs*

```
# Get the curve data of an ROI
>>> get_workEnv().roi_manager.roi_list[3].curve_data

(array([   0,     1,     2, ..., 2995, 2996, 2997]), array([ -207.00168389,  -161.78229208,
↪  -157.62522988, ..., -1017.73174502,
   -1030.27047731, -1042.26989668]))

# Get the tags of an ROI
```

(continues on next page)

(continued from previous page)

```
>>> get_workEnv().roi_manager.roi_list[2].get_all_tags()

{'anatomical_location': 'tail', 'cell_name': 'dcen', 'morphology': 'untagged'}

# Get a single tag
>>> get_workEnv().roi_manager.roi_list[2].get_tag('cell_name')

'dcen'
```

#### 1.14 Caiman Motion Correction

Perform motion correction using the NoRMCorre implementation in the CaImAn library.

**I highly recommend going through the following before using this module**

- **NoRMCorre paper** Pnevmatikakis, E. A., & Giovannucci, A. (2017). NoRMCorre: An online algorithm for piecewise rigid motion correction of calcium imaging data. Journal of Neuroscience Methods, 291, 83–94.
- **The CaImAn demo notebook, the implementation in Mesmerize is basically from the demo** https://github.com/flatironinstitute/CaImAn/blob/master/demos/notebooks/demo_motion_correction.ipynb

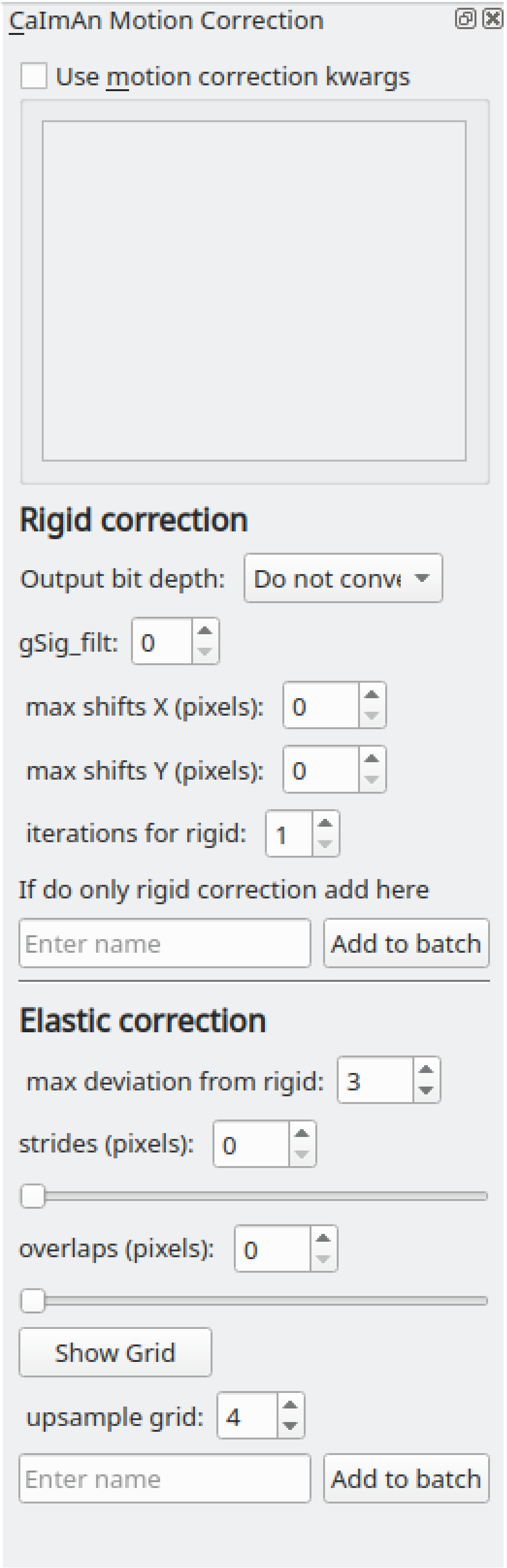


**Parameters**

Output bit depth: The motion corrected image sequences are of float32 type. You can optionally convert the output to

8 or 16 bit uint types to save disk space. **This doesn't always work from my experience, values might get clipped**.

For all other parameters please see the demo notebook mentioned above.

You can also enter parameters as keyword arguments (kwargs) in the text box if you select "Use motion correction kwargs". This is useful if you want to enter parameters that cannot be entered in the GUI for example. **Use single quotes if you want to enter string kwargs, do not use double quotes.**

##### 1.14.1 Usage

This module adds a "caiman motion correction" *item* to the batch. Set the desired parameters (see demo notebook) and then enter a name to add it as an *item* to the batch. After the batch item is processed, double-click the batch item to open the motion corrected image sequence in the viewer. You can then use this motion corrected image sequence for further analysis.

**See also:**

This modules uses the *Batch Manager*.

**Note:** The parameters used for motion correction are stored in the work environment of the viewer and this log is carried over and saved in the *Project Sample* as well. To see the parameters that were used for motion correction in the viewer, execute `get_workEnv().history_trace` in the viewer console and look for the `caiman_motion_correction` entry.

**Warning:** If you're using Windows, large *memmap* files will linger in your batch dir or work dir, you can clean them out periodically.

##### 1.14.2 Script Usage

A script can be used to add caiman motion correction batch items. This is much faster than using the GUI.

**See also:**

*Script Editor*

###### Add items

This example shows how to add all tiff files (of image sequences) from a directory as batch items with 3 different variants of parameters.

**See also:**

This example uses the *Caiman Motion Correction Module API*, *ViewerWorkEnv API*, and *Batch Manager API*

```
# Import glob so we can get all tiff files in a dir
from glob import glob
# Import os to get filenames from paths
import os

# Motion correction params.

mc_kwargs = \
```

(continues on next page)

(continued from previous page)

```
{
    "max_shifts":            (6, 6),
    "niter_rig":             2,
    "max_deviation_rigid":   3,
    "strides":               (196, 196),
    "overlaps":              (98, 98),
    "upsample_factor_grid": 4,
    "gSig_filt":             (10, 10)  # Set to `None` for 2p data
}

params = \
{
    'mc_kwargs':         mc_kwargs,  # the kwargs we set above
    'item_name':         "will set later per file",
    'output_bit_depth': "Do not convert"  # can also set to `8` or `16` if you want the␣
↪output in `8` or `16` bit
}

# Path to the dir containing images
files = glob("/full_path_to_raw_images/*.tiff")
# Sort in alphabetical order (should also work for numbers)
files.sort()

# Open each file, crop, and add to batch with 3 diff mot cor params
for i, path in enumerate(files):
    print("Working on file " + str(i + 1) + " / " + str(len(files)))

    # get json file path for the meta data
    meta_path = path[:-5] + ".json"

    # Create a new work environment with this image sequence
    work_env = ViewerWorkEnv.from_tiff(path, "asarray-multi", meta_path)

    # set it as the current work environment
    vi.viewer.workEnv = work_env
    vi.update_workEnv()

    # Get caiman motion correction module, hide=False to not show GUI
    mc_module = get_module("caiman_motion_correction", hide=True)

    # Set name for this video file
    name = os.path.basename(path)[:-5]
    params["item_name"] = name

    # First variant of params
    params["mc_kwargs"]["strides"] = (196, 196)
    params["mc_kwargs"]["overlaps"] = (98, 98)

    # Add one variant of params for this video to the batch
    mc_module.add_to_batch(params)

    # Try another variant of params
```

(continues on next page)

(continued from previous page)

```
    params["mc_kwargs"]["strides"] = (256, 256)
    params["mc_kwargs"]["overlaps"] = (128, 128)

    # Set these params and add to batch
    mc_module.add_to_batch(params)

    # Try one more variant of params
    params["mc_kwargs"]["strides"] = (296, 296)
    params["mc_kwargs"]["overlaps"] = (148, 148)

    # Set these params and add to batch
    mc_module.add_to_batch(params)

# If you want to process the batch after adding the items uncomment the following lines
#bm = get_batch_manager()
#bm.process_batch(clear_viewers=True)
```

##### Crop and add items

This example shows how to crop videos prior to adding them as batch items. This is useful if you want to crop-out large unchanging regions of your movides. It uses either simple thresholding or spectral salieny on a standard deviation projection to determine the bounding box for cropping.

**See also:**

This example uses the *Caiman Motion Correction Module API*, *ViewerWorkEnv API*, and *Batch Manager API*

```
# Import glob so we can get all tiff files in a dir
from glob import glob
# Import os to get filenames from paths
import os

# Just get a shortcut reference to the auto_crop function
auto_crop = image_utils.auto_crop

# Parameters for cropping, these should work for everything
# These worked well for various different constructs
# If you get non-specific cropping (too much black) try "method" as "spectral_saliency"␣
↪(See below)
crop_params = \
{
    "projection":       "max+std",
    "method":           "threshold",
    "denoise_params":   (32, 32),
}

# Spectral saliency is another method
# You can try and play around with the parameters
# If the cropping is insufficient, you can set "projection" to just "max" or "std"
# If you get too much junk blackness around the animal try increasing denoise_params
# or reduce padding. Default padding is 30 (when nothing is specified like above)
crop_params_salient = \
```

(continues on next page)

(continued from previous page)

```
{
    "projection":       "max+std",
    "method":           "spectral_saliency",
    "denoise_params":   (16, 16),
    "padding":          40
}

# Motion correction params.
mc_kwargs = \
{
    "max_shifts":           (6, 6),
    "niter_rig":            2,
    "max_deviation_rigid":  3,
    "strides":              (196, 196),
    "overlaps":             (98, 98),
    "upsample_factor_grid": 4,
    "gSig_filt":            (10, 10)  # Set to `None` for 2p data
}

params = \
{
    'mc_kwargs':        mc_kwargs,  # the kwargs we set above
    'item_name':        "will set later per file",
    'output_bit_depth': "Do not convert"  # can also set to `8` or `16` if you want the output in `8` or `16` bit
}

# Path to the dir containing images
files = glob("/full_path_to_raw_images/*.tiff")
# Sort in alphabetical order (should also work for numbers)
files.sort()

# Open each file, crop, and add to batch with 3 diff mot cor params
for i, path in enumerate(files):
    print("Working on file " + str(i + 1) + " / " + str(len(files)))

    # get json file path for the meta data
    meta_path = path[:-5] + ".json"

    # Create a new work environment with this image sequence
    work_env = ViewerWorkEnv.from_tiff(path, "asarray-multi", meta_path)

    # autocrope the image sequence in the work environment
    raw_seq = work_env.imgdata.seq
    # Auto crop the image sequence
    print("Cropping file: " + str(i + 1))

    cropped = auto_crop.crop(raw_seq, crop_params)
    # Set work env img seq to the cropped one and update
    work_env.imgdata.seq = cropped

    # update thew work environment
```

(continues on next page)

(continued from previous page)

```
    vi.viewer.workEnv = work_env
    vi.update_workEnv()

    # Get caiman motion correction module, hide=False to not show GUI
    mc_module = get_module("caiman_motion_correction", hide=True)

    # Set name for this video file
    name = os.path.basename(path)[:-5]
    params["item_name"] = name

    # First variant of params
    params["mc_kwargs"]["strides"] = (196, 196)
    params["mc_kwargs"]["overlaps"] = (98, 98)

    # Add one variant of params for this video to the batch
    mc_module.add_to_batch(params)

    # Try another variant of params
    params["mc_kwargs"]["strides"] = (256, 256)
    params["mc_kwargs"]["overlaps"] = (128, 128)

    # Set these params and add to batch
    mc_module.add_to_batch(params)

    # Try one more variant of params
    params["mc_kwargs"]["strides"] = (296, 296)
    params["mc_kwargs"]["overlaps"] = (148, 148)

    # Set these params and add to batch
    mc_module.add_to_batch(params)

# If you want to process the batch after adding the items uncomment the following lines
#bm = get_batch_manager()
#bm.process_batch(clear_viewers=True)
```

#### 1.15 CNMF

Perform CNMF using the implementation provided by the CaImAn library. This modules basically provides a GUI for parameter entry.

**I highly recommend going through the following before using this module**

- **CNMFE builds upon CNMF** Pnevmatikakis, E. A., Gao, Y., Soudry, D., Pfau, D., Lacefield, C., Poskanzer, K., … Paninski, L. (2014). A structured matrix factorization framework for large scale calcium imaging data analysis, 1–16.

  Pnevmatikakis, E. A., Soudry, D., Gao, Y., Machado, T. A., Merel, J., Pfau, D., … Paninski, L. (2016). Simultaneous Denoising, Deconvolution, and Demixing of Calcium Imaging Data. Neuron, 89(2), 285.

- **CaImAn demo notebook, the implementation in Mesmerize is basically from the demo. The second half of the notebook d**[illegible] https://github.com/flatironinstitute/CaImAn/blob/master/demos/notebooks/demo_pipeline.ipynb

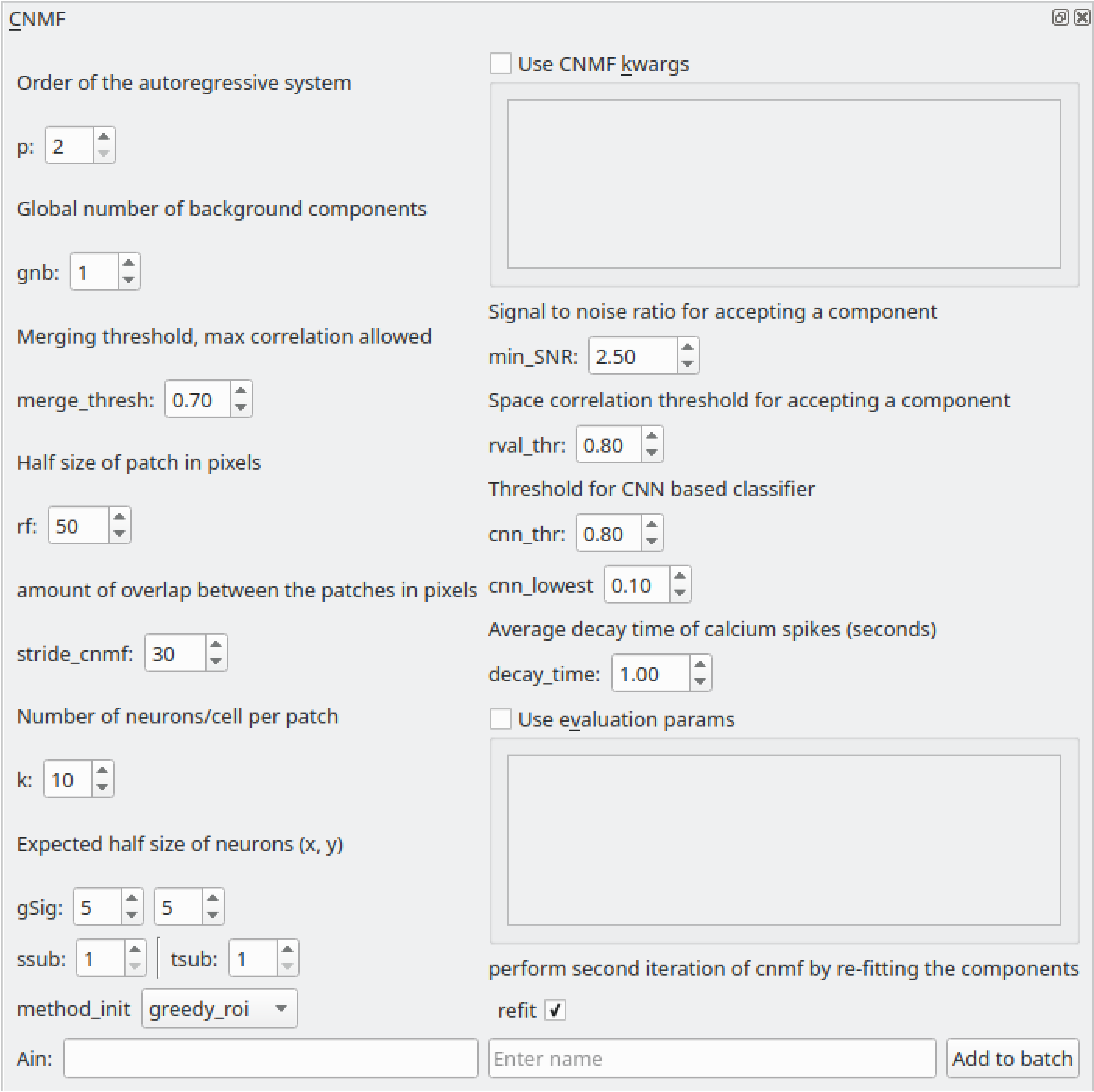


###### Parameters

Please see the CaImAn demo notebook mentioned above to understand the parameters. The Caiman docs also provide descriptions of the parameters: https://caiman.readthedocs.io/

You can also enter parameters for CNMF and component evaluation as keyword arguments (kwargs) in the the respective text boxes if you select "Use CNMF kwrags" or "Use evaluation params". This is useful if you want to enter parameters that cannot be entered in the GUI for example. **Use single quotes if you want to enter string kwargs, do not use double quotes.**

##### 1.15.1 Usage

This module adds a "CNMF" *item* to the batch. Set the desired parameters (see Caiman docs & demos) and then enter a name to add it as an *item* to the batch. After the batch item is processed, **double-click the batch item** to import the CNMF output into a Viewer. You can then annotate and curate ROIs, and add the data as a *Sample* to your project.

**See also:**

This modules uses the *Batch Manager*.

> **Warning:** It's recommended to open a new Viewer when you want to import 3D CNMF data. Full garbage collection of 3D data in the Viewer Work environment is a WIP for when you want to clear & import 3D data into the same viewer. However when you close the Viewer entirely it is garbage collected entirely.

**Note:** The parameters used for CNMF are stored in the work environment of the viewer and this log is carried over and saved in *Project Samples* as well. To see the parameters that were used for CNMF in the viewer, execute `get_workEnv().history_trace` in the viewer console and look for the 'cnmf' entry.

> **Warning:** Importing several *thousands* of ROIs into the Viewer can take 15-30 minutes. You will be able to track the progress of the import in the Viewer Window's status bar.

> **Warning:** If you're using Windows, large *memmap* files will linger in your batch dir or work dir, you can clean them out periodically.

##### 1.15.2 Script usage

A script can be used to add CNMF batch items. This is much faster than using the GUI. This example sets the work environment from the output of a batch item. See the *Caiman Motion Correction script usage examples* for how to load images if you want to add CNMF items from images that are not in a batch.

**See also:**

*Script Editor*

```python
def reset_params():
    # CNMF Params that we will use for each item
    cnmf_kwargs = \
    {
        'p': 2,
        'gnb': 1,
        'merge_thresh': 0.25,
        'rf': 70,
        'stride': 40,
        'k': 16,
        'gSig': (8, 8),
        'gSiz': (33, 33)
    }

```

(continues on next page)

(continued from previous page)

```
    # component evaluation params
    eval_kwargs = \
    {
        'min_SNR': 2.5,
        'rval_thr': 0.8,
        'min_cnn_thr': 0.8,
        'cnn_lowest': 0.1,
        'decay_time': 2.0,
    }

    # the dict that will be passed to the mesmerize caiman module
    params = \
    {
        "cnmf_kwargs":  cnmf_kwargs,
        "eval_kwargs":  eval_kwargs,
        "refit":        True,  # if you want to perform a refit
        "item_name":    "will set later per file",
    }

    return params

# Get the batch manager
bm = get_batch_manager()
cnmf_mod = get_module('cnmf', hide=True)

# Start index if we want to start processing the new items after they have been added
start_ix = bm.df.index.size + 1

# This example uses motion corrected output items from the batch manager
# You can also open image files directly from disk, see the motion correction
# script examples to see how to open images from disk.
for ix, r in bm.df.iterrows():
    # Use output of items 6 - 12
    # for example if items 6 - 12 were motion correction items
    if ix < 6:
        continue
    if ix > 12: # You need to set a break point, else the batch grows infinitely
        break

    # get the first variant of params
    params = reset_parmas()

    # Get the name of the mot cor item
    name = r['name']

    # Set the name for the new cnmf item
    params['item_name'] = name

    # Load the mot cor output
    bm.load_item_output(module='caiman_motion_correction', viewers=viewer, UUID=r['uuid
↪'])

```

(continues on next page)

(continued from previous page)

```
    # Set the sampling rate of the data
    params['eval_kwargs']['fr'] = vi.viewer.workEnv.imgdata.meta['fps']

    # Get the border_pix value from the motion correction output
    # skip this if loading files that don't have NaNs on the image borders
    history_trace = vi.viewer.workEnv.history_trace
    border_pix = next(d for ix, d in enumerate(history_trace) if 'caiman_motion_
↪correction' in d)['caiman_motion_correction']['bord_px']

    # Set the border_pix values
    params['border_pix'] = border_pix
    params['cnmf_kwargs']['border_pix'] = border_pix

    # Add to batch
    cnmf_mod.add_to_batch(params)

    # change some of the params and add this variant to batch
    params['cnmf_kwargs']['gSig'] = (10, 10)
    params['cnmf_kwargs']['gSiz'] = (41, 41)

    # Add to batch with this params variant
    cnmf_mod.add_to_batch(params)

    # another parameter variant
    params['eval_kwargs']['rval_thr'] = 0.7
    params['eval_kwargs']['min_cnn_thr'] = 0.65

    # Add to batch with this params variant
    cnmf_mod.add_to_batch(params)

# Cleanup the work environment
vi._clear_workEnv()

# Uncomment the last two lines to start the batch as well
#bm.process_batch(start_ix, clear_viewers=True)
```

#### 1.16 CNMF 3D

Perform 3D CNMF using the implementation provided by the CaImAn library. This modules basically provides a GUI for parameter entry.

**I highly recommend going through the following before using this module**

- **CNMFE builds upon CNMF** Pnevmatikakis, E. A., Gao, Y., Soudry, D., Pfau, D., Lacefield, C., Poskanzer, K., … Paninski, L. (2014). A structured matrix factorization framework for large scale calcium imaging data analysis, 1–16.

  Pnevmatikakis, E. A., Soudry, D., Gao, Y., Machado, T. A., Merel, J., Pfau, D., … Paninski, L. (2016). Simultaneous Denoising, Deconvolution, and Demixing of Calcium Imaging Data. Neuron, 89(2), 285.

- **CaImAn demo notebook, the implementation in Mesmerize is basically from the demo.** https://github.com/flatironinstitute/CaImAn/blob/master/demos/notebooks/demo_caiman_cnmf_3D.ipynb

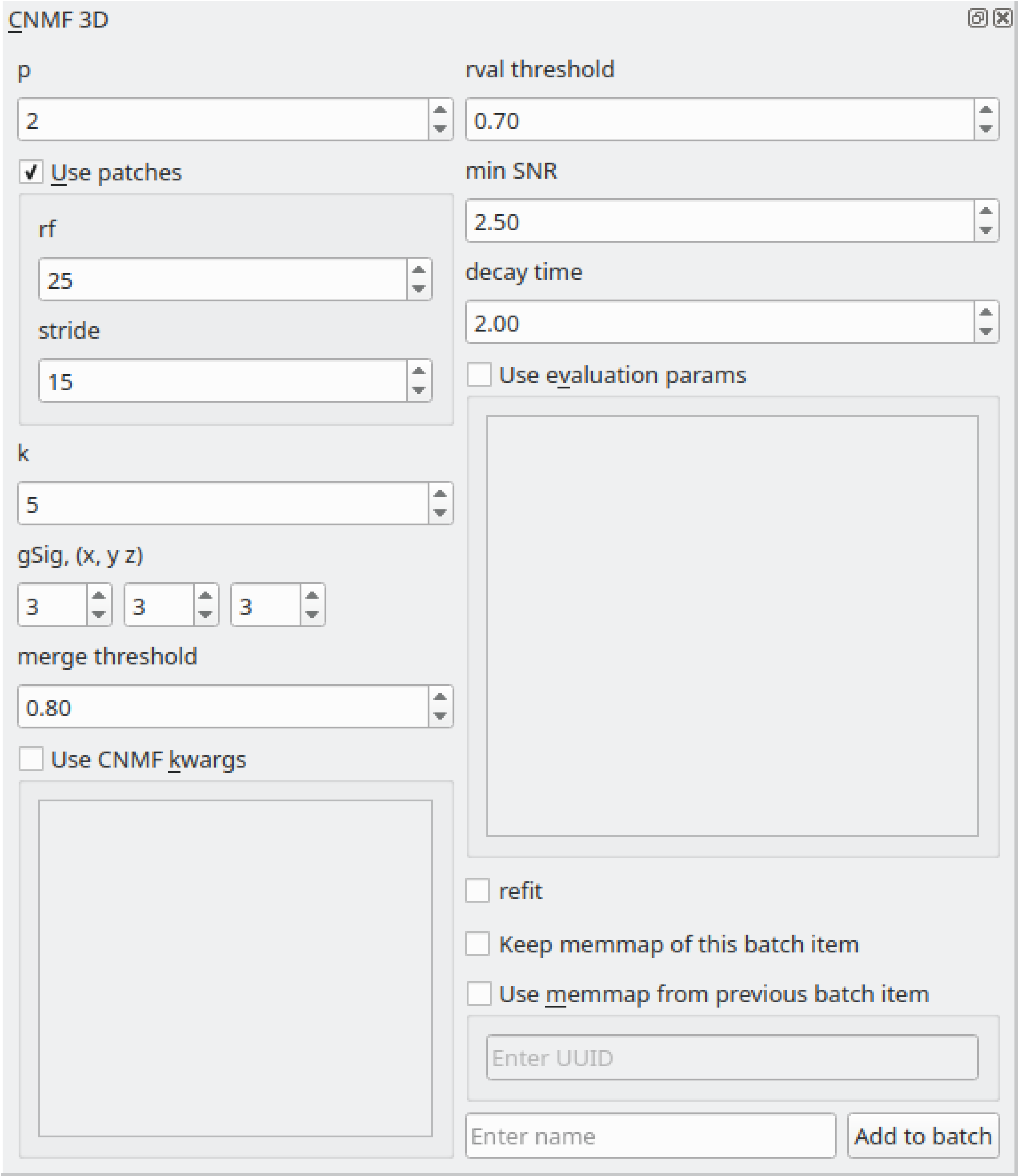


**Parameters**

Please see the CaImAn demo notebook mentioned above to understand the parameters. The Caiman docs also provide descriptions of the parameters: https://caiman.readthedocs.io/

You can also enter parameters for CNMF and component evaluation as keyword arguments (kwargs) in the the respective text boxes if you select "Use CNMF kwrags" or "Use evaluation params". This is useful if you want to enter parameters that cannot be entered in the GUI for example. **Use single quotes if you want to enter string kwargs, do not use double quotes.**

**Note:** The parameters used for 3D CNMF are stored in the work environment of the viewer and this log is carried over and saved in *Project Samples* as well. To see the parameters that were used for 3D CNMF in the viewer, execute `get_workEnv().history_trace` in the viewer console and look for the 'cnmf_3d' entry.

---

> **Warning:** Importing several *thousands* of ROIs into the Viewer can take 15-30 minutes. You will be able to track the progress of the import in the Viewer Window's status bar.

##### 1.16.1 Usage

This module adds a "CNMF_3D" *item* to the batch. Set the desired parameters (see Caiman docs & demos) and then enter a name to add it as an *item* to the batch. After the batch item is processed, **double-click the batch item** to import the CNMF output into a Viewer. You can then annotate and curate ROIs, and add the data as a *Sample* to your project.

**See also:**

This modules uses the *Batch Manager*.

> **Warning:** It's recommended to open a new Viewer when you want to import 3D CNMF data. Full garbage collection of 3D data in the Viewer Work environment is a WIP for when you want to clear & import 3D data into the same viewer. However when you close the Viewer entirely it is garbage collected entirely.

##### 1.16.2 Script Usage

A script can be used to add CNMF batch items. This is much faster than using the GUI. This example sets the work environment from the output of a batch item.

**See also:**

*Script Editor*

This example loads 3D sequences from disk & adds them to a batch with 3 parameter variants.

```python
# just so we can reset the params for each new image file
def reset_params():
    # CNMF Params that we will use for each item
    cnmf_kwargs = \
    {
        'p': 2,
        'merge_thresh': 0.8,
        'k': 50,
        'gSig': (10, 10, 1),
        'gSiz': (41, 41, 4)
    }

    # component evaluation params
    eval_kwargs = \
    {
        'min_SNR': 3.0,
        'rval_thr': 0.75,
        'decay_time': 1.0,
    }

```

(continues on next page)

(continued from previous page)

```
    # the dict that will be passed to the mesmerize caiman module
    params = \
    {
        "cnmf_kwargs":  cnmf_kwargs,
        "eval_kwargs":  eval_kwargs,
        "refit":        True,  # if you want to perform a refit
        "item_name":    "will set later per file",
        "use_patches":  False,
        "use_memmap":   False,  # re-use the memmap from a previous batch item, reduces computation time
        "memmap_uuid:   None,   # UUID (as a str) of the batch item to use the memmap from
        "keep_memmmap": False   # keep the memmap of this batch item

    }

    return params

# get the 3d cnmf module
cnmf_mod = get_module('cnmf_3d', hide=True)

# Path to the dir containing images
files = glob("/full_path_to_raw_images/*.tiff")
# Sort in alphabetical order (should also work for numbers)
files.sort()

# Open each file, crop, and add to batch with 3 diff mot cor params
for i, path in enumerate(files):
    print("Working on file " + str(i + 1) + " / " + str(len(files)))

    # get json file path for the meta data
    meta_path = path[:-5] + ".json"

    # Create a new work environment with this image sequence
    vi.viewer.workEnv = ViewerWorkEnv.from_tiff(path=path,           # tiff file path
                                                method='imread',     # use imread
                                                meta_path=meta_path, # json metadata file path
                                                axes_order=None)     # default axes order
                                                                     # see Mesmerize Tiff file module docs for more info on axes order

    # update the work environment
    vi.update_workEnv()

    # get the first variant of params
    params = reset_parmas()

    # Set name for this video file
    name = os.path.basename(path)[:-5]
    params["item_name"] = name

```

(continues on next page)

(continued from previous page)

```
    # add batch item with one variant of params
    u = cnmf_mod.add_to_batch(params)

    # add the same image but change some params
    params["cnmf_kwargs"]["gSig"] = (12, 12, 1)
    params["eval_kwargs"]["min_SNR"] = 2.5

    # use the same memmap as the previous batch item
    # since it's the same image
    params["use_memmap"] = True
    params["memmap_uuid"] = str(u)

    # add this param variant to the batch
    cnmf_mod.add_to_batch(params)

    # one more variant of params
    params["eval_kwargs"]["min_SNR"] = 2.0

    # add this param variant to the batch
    cnmf_mod.add_to_batch(params)
```

#### 1.17 CNMFE

Perform CNMFE using the implementation provided by the CaImAn library.

**I highly recommend going through the following before using this module**

- **The paper on CNMF-E** Zhou, P., Resendez, S. L., Rodriguez-Romaguera, J., Jimenez, J. C., Neufeld, S. Q., Stuber, G. D., … Paninski, L. (2016). Efficient and accurate extraction of in vivo calcium signals from microendoscopic video data. ELife, 1–37.
- **CNMFE builds upon CNMF** Pnevmatikakis, E. A., Gao, Y., Soudry, D., Pfau, D., Lacefield, C., Poskanzer, K., … Paninski, L. (2014). A structured matrix factorization framework for large scale calcium imaging data analysis, 1–16.

  Pnevmatikakis, E. A., Soudry, D., Gao, Y., Machado, T. A., Merel, J., Pfau, D., … Paninski, L. (2016). Simultaneous Denoising, Deconvolution, and Demixing of Calcium Imaging Data. Neuron, 89(2), 285.
- **CaImAn CNMF-E demo notebook, the implementation in Mesmerize is basically from the demo** https://github.com/flatironinstitute/CaImAn/blob/master/demos/notebooks/demo_pipeline_cnmfE.ipynb

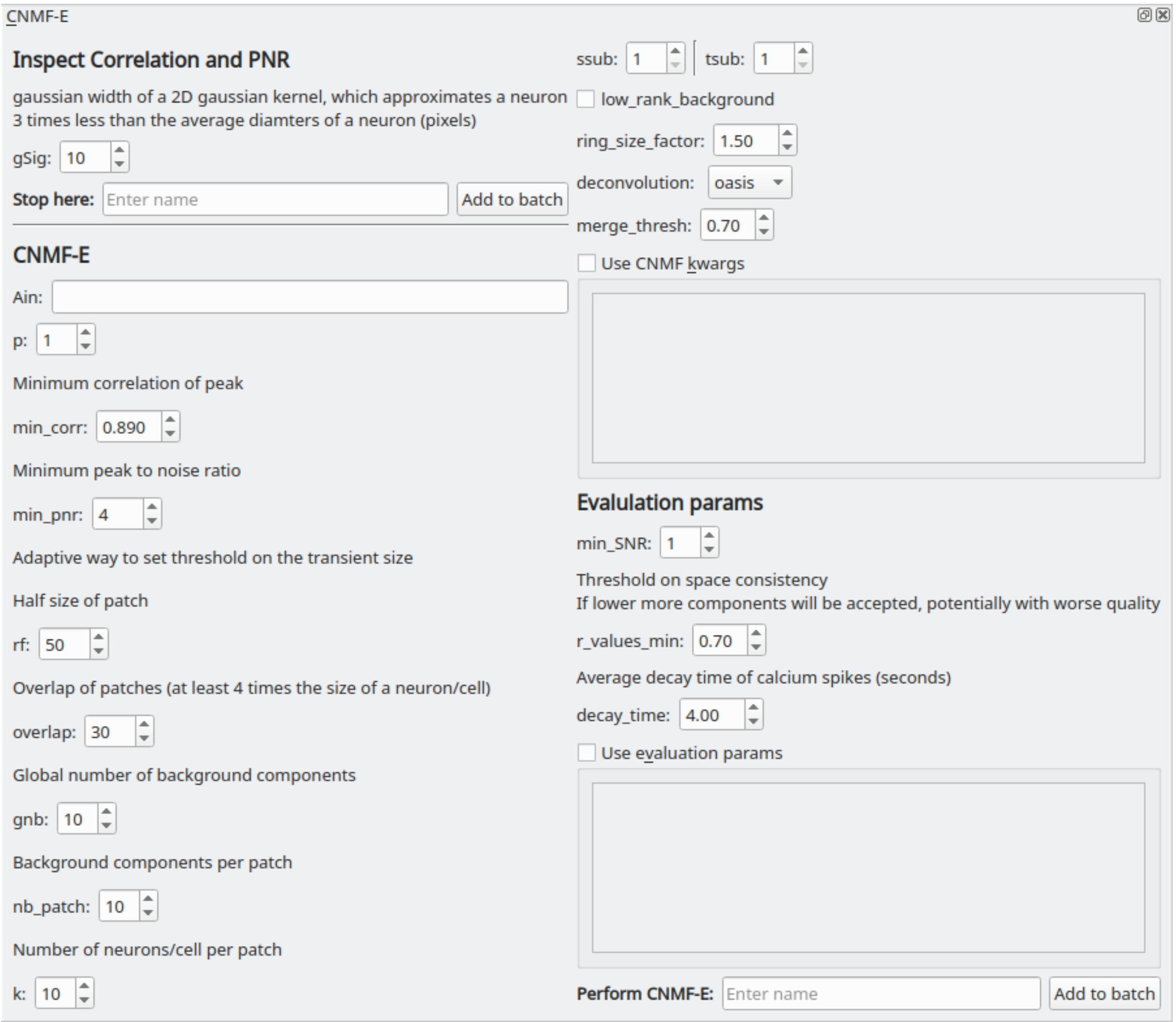


**Parameters**

**Ain**: Seed spatial components from another CNMFE item by entering its UUID here.

Please see the CaImAn demo notebook mentioned above to understand the rest of the parameters. The Caiman docs also provide descriptions of the parameters: https://caiman.readthedocs.io/

You can also enter parameters for CNMF and component evaluation as keyword arguments (kwargs) in the the respective text boxes if you select "Use CNMF kwrags" or "Use evaluation params". This is useful if you want to enter parameters that cannot be entered in the GUI for example. **Use single quotes if you want to enter string kwargs, do not use double quotes.**

##### 1.17.1 Usage

This module creates two types of batch items, one where you can inspect the Correlation & PNR images and another that performs CNMFE and extracts components. Here is an outline of typical usage:

- Enter a *gSig* parameter value and a name for "Inspect Correlation and PNR", the text entry for "Stop here". Click "Add to batch". Run the batch item.
- Double-click the batch item, you will be presented with a GUI to help optimize *min_corr* and *min_pnr*. For the correlation image use the vmin slider to optimize the seperation of cells and set the *min_corr* parameter to this value. Likewise, optimize the value for the PNR until the PNR image mostly contains regions that show real signal and no or few regions that are likely to be just noise and set this vmin value as the *min_pnr* parameter. You may need to try slightly different variations to optimize the parameters.

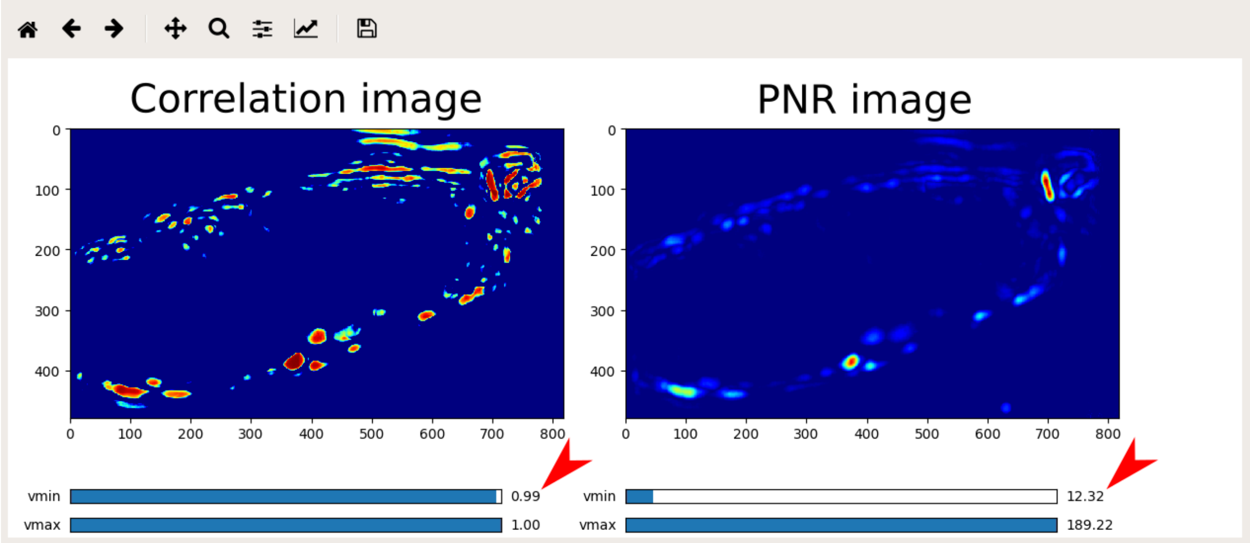


- Enter the rest of the parameters and give a name under "Perform CNMF-E", click "Add to batch" and run the item.
- Double-click the batch item and you will be presented with 3 options. The first option will display the correlation-pnr images and the second option is currently non-functional (matplotlib Qt issue). The last option will import the components extracted by CNMFE into an open Viewer. The components are managed by the ROI Manager.

**See also:**

*ROI Manager*

**See also:**

This modules uses the *Batch Manager*.

**Note:** The parameters used for CNMFE are stored in the work environment of the viewer and this log is carried over and saved in *Project Samples* as well. To see the parameters that were used for CNMFE in the viewer, execute `get_workEnv().history_trace` in the viewer console and look for the 'cnmfe' entry.

**Warning:** If you're using Windows, large *memmap* files will linger in your batch dir or work dir, you can clean them out periodically.

##### 1.17.2 Script Usage

A script can be used to add CNMFE batch items. This is much faster than using the GUI.

**See also:**

*Script Editor*.

###### Add Corr PNR items

Add Corr PNR batch items from a batch that contains motion corrected items. This example add 2 variants of parameters (just gSig) for each motion corrected item.

**See also:**

This example uses the *Caiman CNMFE module API* and *Batch Manager API*

**See also:**

*Caiman Motion Correction script usage examples* for how to load images if you want to add Corr PNR items from images that are not in a batch.

```
# Get the batch manager
bm = get_batch_manager()

# Get the CNMFE module
cnmfe_mod = get_module('cnmfe', hide=True)

# Start index to start processing the new items after they have been added
start_ix = bm.df.index.size + 1

for ix, r in bm.df.iterrows():
        if ix == start_ix:
                break

        # Load the output of the motion corrected batch item
        # The output will load into the viewer that this script
        # is running in.
        bm.load_item_output(module='caiman_motion_correction', viewers=viewer, UUID=r[
↪'uuid'])

        # Get the currently set params
        # You just need the dict with all the correct keys
        # You will just modify the "gSig" and "item_name" keys
        params = cnmfe_mod.get_params(item_type='corr_pnr', group_params=True)

        # Get the name of the mot cor item
        name = r['name']
        params['item_name'] = name

        params['border_pix'] = border_pix

        # Set the gSig and name params
        params['corr_pnr_kwargs']['gSig'] = 8

        # Add to batch
```

(continues on next page)

(continued from previous page)

```
        cnmfe_mod.add_to_batch_corr_pnr(params)

        # Another variant of params
        params['corr_pnr_kwargs']['gSig'] = 10

        # Add to batch with this variant of params
        cnmfe_mod.add_to_batch_corr_pnr(params)

# Cleanup the work environment
vi._clear_workEnv()

# Start the batch from the start_ix
bm.process_batch(start_ix, clear_viewers=True)
```

##### CNMFE

#### 1.18 Caiman HDF5 Importer

You can import HDF5 files containing CNMF results that were produced externally by Caiman. The ROIs produced by CNMF, 3D-CNMF or CNMFE will be imported into the current *work environment* and placed onto the image that is currently open.

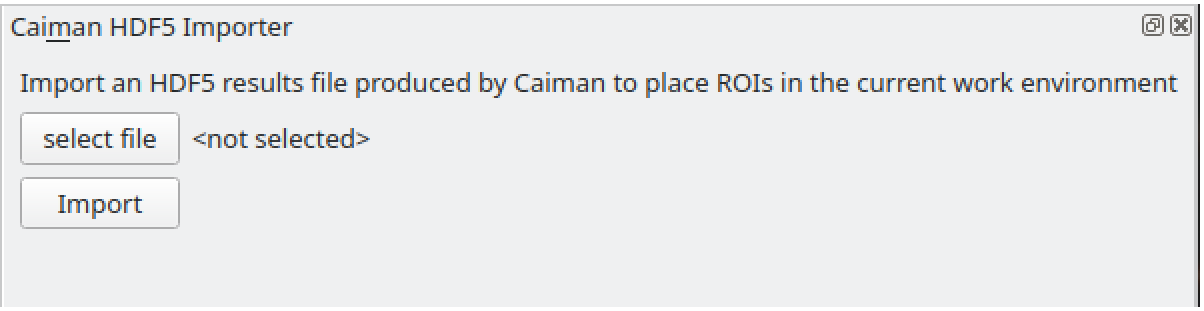


You can also use this module through the *viewer console*, or in the *Script Editor* instead of clicking buttons.

**Example**

```
# get the module, hide the GUI
caiman_importer = get_module('caiman_importer', hide=True)

# import the file
caiman_importer.import_file('/path/to/file.hdf5')
```

#### 1.19 Suite2p Importer

You can load Suite2p output files to import ROIs into the current *work environment*. This places the Suite2p-derived ROIs onto the image that is currently open.

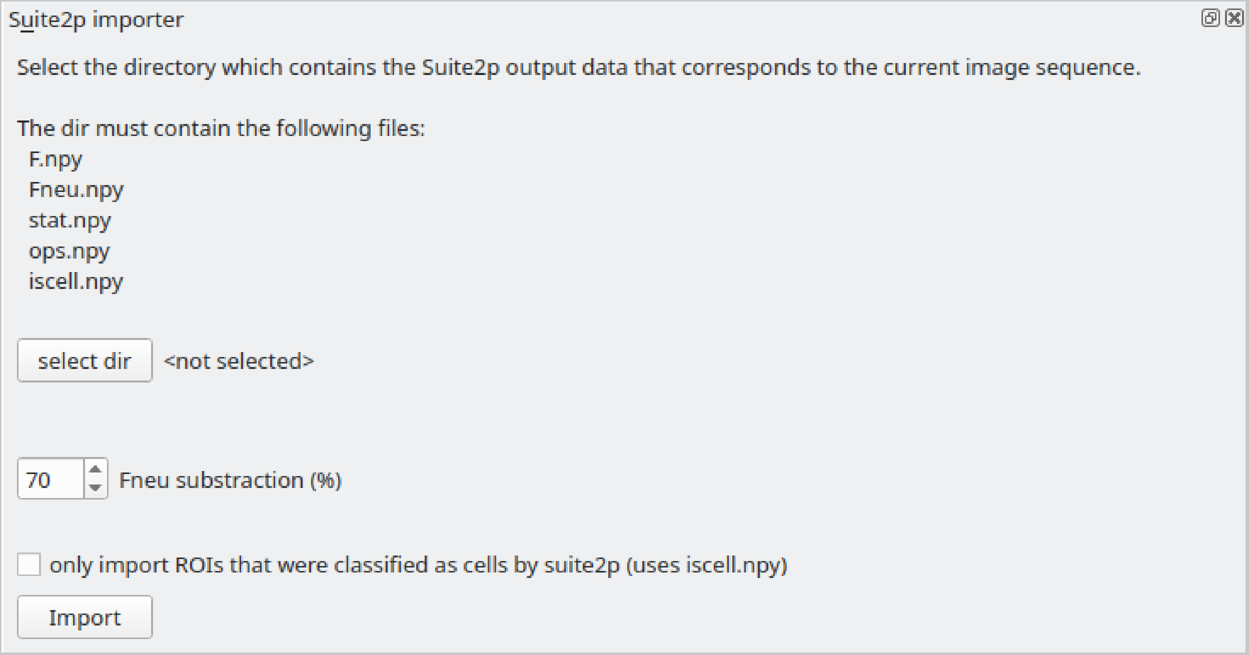


##### 1.19.1 Video Tutorial

##### 1.19.2 Script Usage

You can also use this module through the *viewer console*, or in the *Script Editor* instead of clicking buttons.

**Example**

```
# get the module, hide the GUI
s2p_importer = get_module('suite2p_importer', hide=True)

# set the path to the dir containing the suite2p output files
s2p_importer.data.set_dir('/path/to/dir')

# set the amount of neuropil contamination to subtract
s2p_importer.data.Fneu_sub = 0.7

# import the suite2p data into the current work environment
s2p_importer.import_rois()

# clear the data from the importer before importing another directory
# this doesn't do anything to the viewer work environment, just clears the importer data
s2p_importer.data.clear()
```

#### 1.20 Nuset Segmentation

Deep learning based segmentation, useful for nuclear localized indicators. ROIs segmented through this module can be imported into the Viewer Work Environment.

**Note:** If you use this tool, please cite the Nuset paper in addition to citing Mesmerize: Yang L, Ghosh RP, Franklin JM, Chen S, You C, Narayan RR, et al. (2020) NuSeT: A deep learning tool for reliably separating and analyzing crowded cells. PLoS Comput Biol 16(9): e1008193. https://doi.org/10.1371

##### 1.20.1 Parameters

###### Projection

Choose a projection which maximizes the visibility of your regions of interest

###### Pre-process

| Parameter | Description |
|---|---|
| do_preprocess | perform pre-processing |
| do_sigmoid | perform sigmoid correction |
| sig-moid_cutoff | cutoff, lower values will increase the exposure |
| sigmoid_gain | gain, high values can be thought of as increasing contrast |
| sig-moid_invert | invert the image if necessary. Regions of interesting should be bright, background should be dark |
| do_equalize | perform adaptive histogram equalization |
| equal-ize_lower | Set a lower limit, this helps remove background & increase contrast |
| equal-ize_upper | Upper limit for the histogram |
| equal-ize_kernel | kernel size, increase if the pre-processed image is grainy. Start with a value ~1/16-1/8 the size of the image |

###### NuSeT

| Parameter | Description |
|---|---|
| watershed | wastershed the image, useful if your cells are tightly packed. Uncheck if cells are large and/or sparse. |
| min_score | Decreasing this value will cause more regions to be found, i.e. cells tend to split more |
| nms_threshold | Increasing this value will cause more regions to be found, i.e. cells tend to split more |
| rescale_ratio | Use smaller values less than 1.0 if you have large bright cells, If you have smaller or dim cells use values higher than 1.0 |

**Note:** min_score & nms_threshold work in opposing ways

**Note:** Segmentation will utilize all threads available on your system (regardless of the value set in your System Configuration). However it only takes a few seconds or a few minutes if segmenting a large 3D stack.

**Note:** high **rescale_ratio** values will increase the time required for segmentation. Values around 3.0 take about ~1 minute for 512x512 sized images on ~16 core CPUs.

###### Post-process

##### 1.20.2 Export

If you export using a Convex Hull masks containing only a few pixels, which may be noise, will be removed.

**Note:** Segmentation will utilize all threads available on your system (regardless of the value set in your System Configuration). However it only takes a few seconds if exporting a 2D image, and make take ~10 minutes if exporting a large 3D stack.

#### 1.21 Script Editor

A simple text editor for writing scripts that can be run in the *viewer console*

The scripts are simply ran in the *viewer console* and all output will also be visible in the *viewer console*.

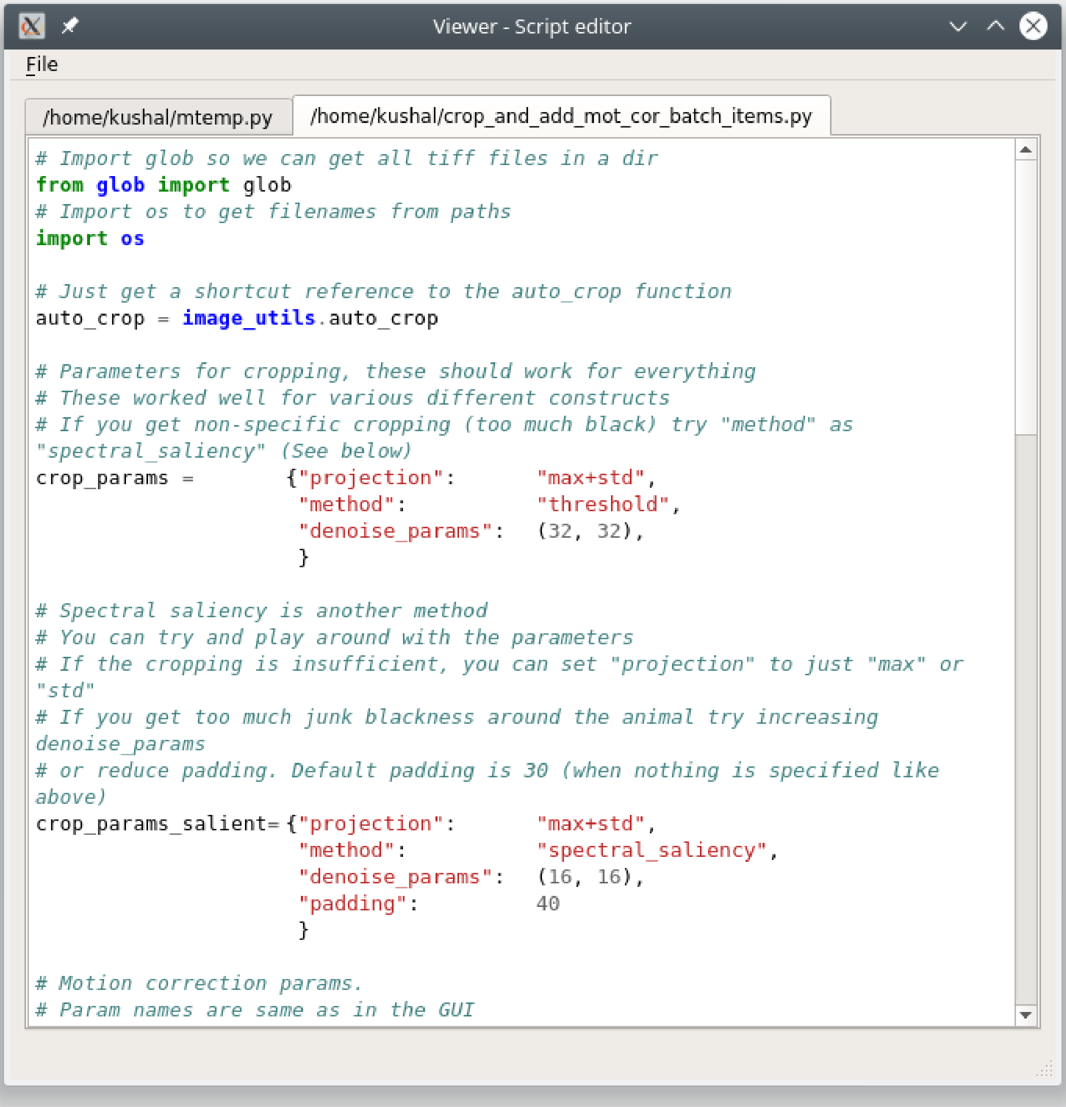


**See also:**

*Viewer Core API*

**Warning:** There is no auto-save function

#### 1.22 Femtonics Importers

You can import `.mes` and `.mesc` files containing data recorded by a Femtonics microscope. Access these modules in Viewer through Modules -> Load images -> Femtonics

##### 1.22.1 mesc files

You can explore the contents of a `.mesc` file using the module's GUI show on the left in the image below. To load a recording just double click on a selection under *Channel*. If the recording is an image sequence it will be imported into the Viewer. If the recording is a Curve a plot will open in a new window to display the curve.

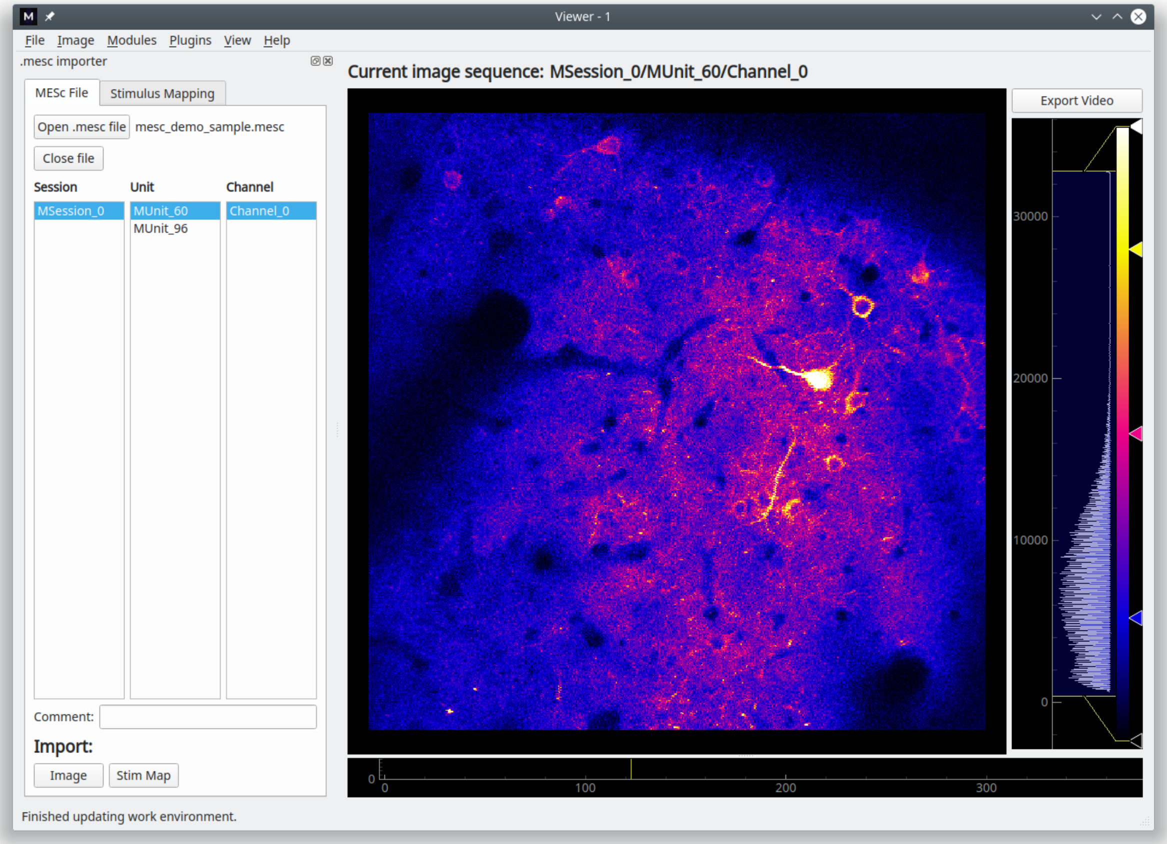


##### 1.22.2 mes files

You can import recordings from a `.mes` using this module, and you can also map metadata from the microscope to specific stimuli.

To load an image sequence into the Viewer work environment, just double click the desired recording from the list.

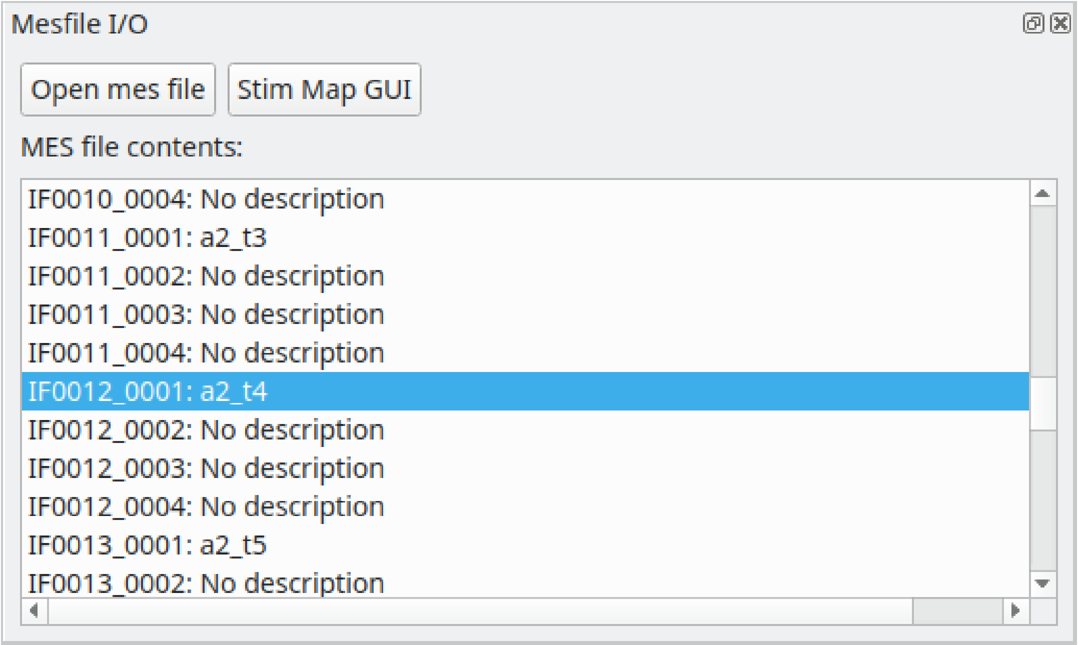


You can map voltage data from various microscope channels (such as auxiliary outputs) to specific stimuli. The stimulus types which you can choose from will correspond to the Stimulus Type columns in your *Project Configuration*. You can view & edit the imported stimulus data using the *Stimulus Mapping Module*

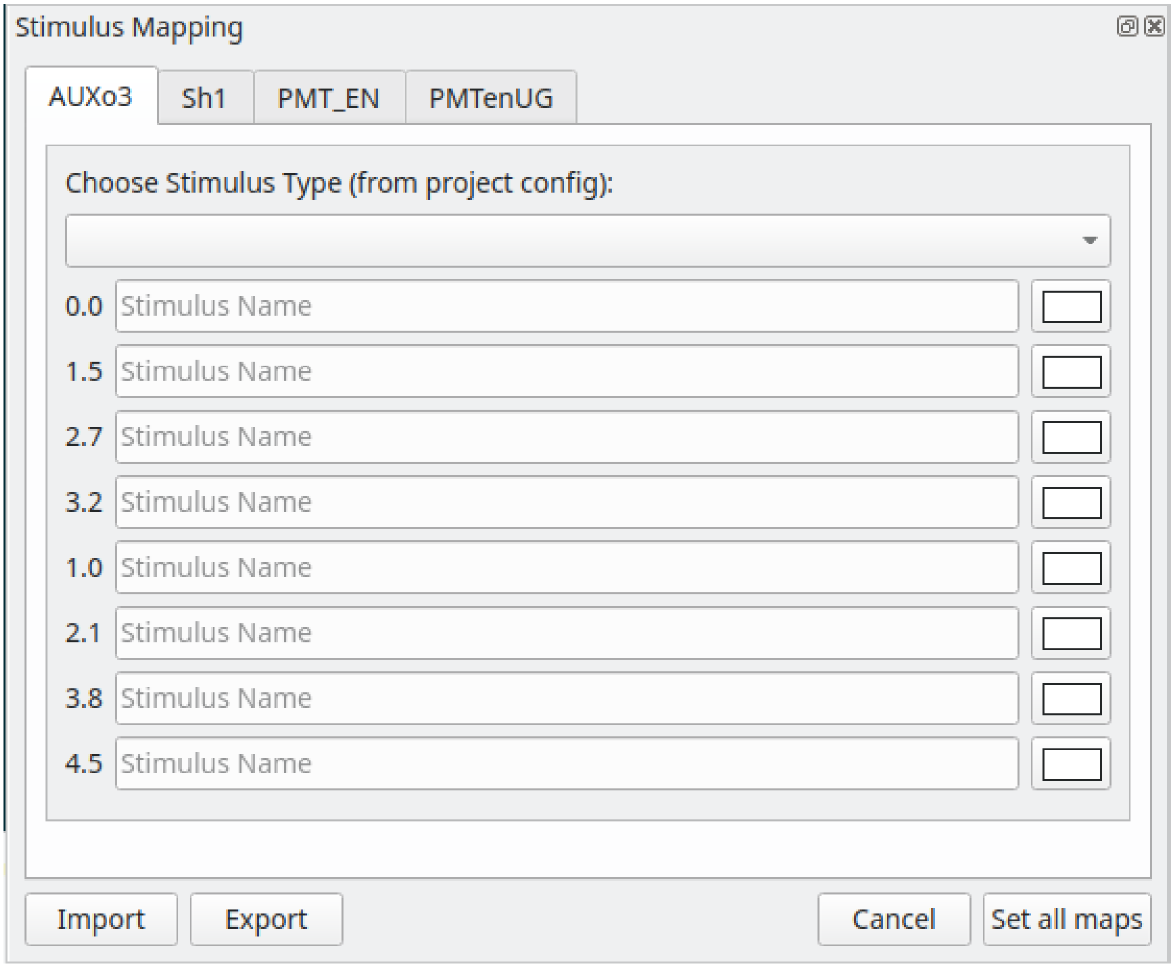


#### 1.23 Flowchart Overview

The flowchart allows you to analyze samples in your project and create plots by arranging analysis nodes. Each node takes an input, performs an operation, and produces an output. For example the *Derivative* node takes use-specified numerical arrays, computes the derivative of these arrays, and then outputs the result.

The Flowchart is based on the pyqtgraph flowchart widgets

**Flowchart Window**

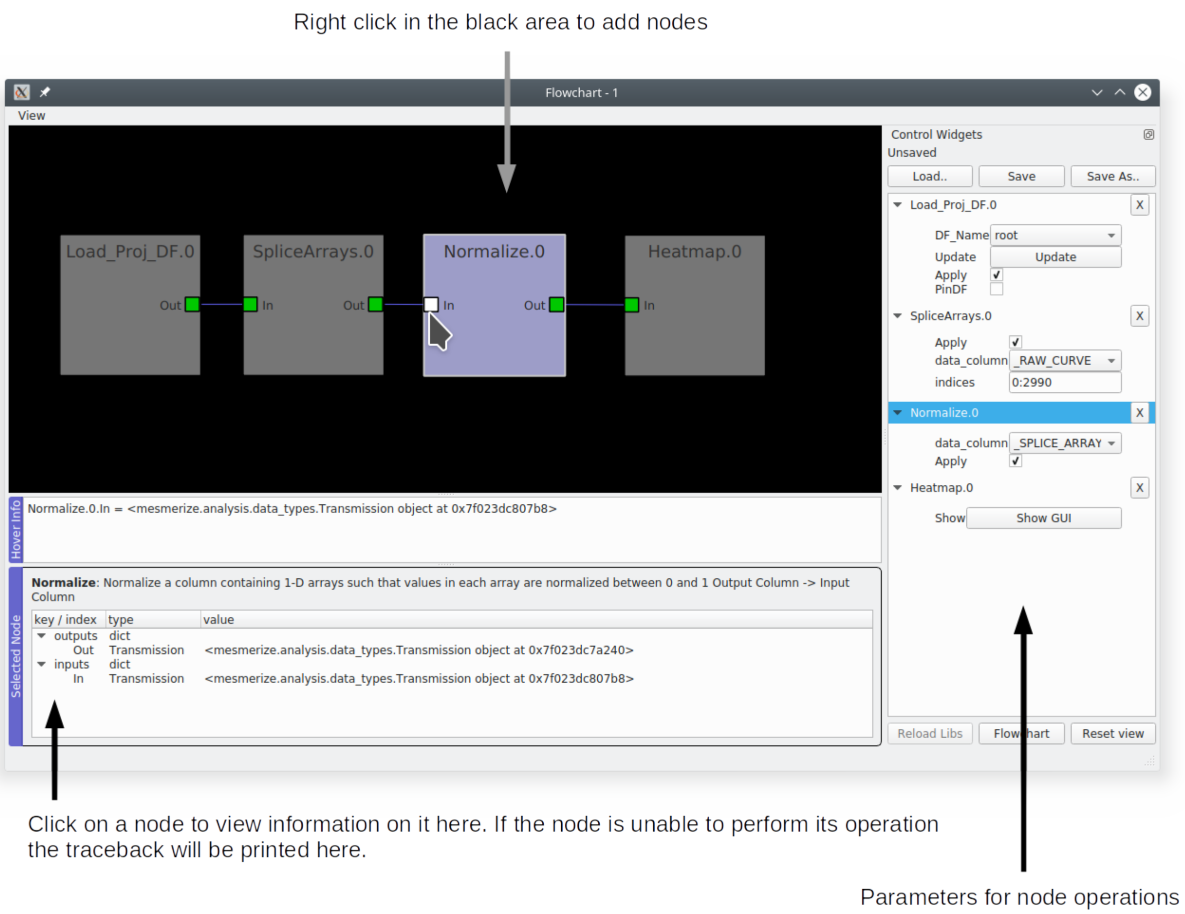


**Add node**: Right click -> Add node -> Choose from selection

Click on a node to highlight the Control Widget

**Remove node**: Right click -> Remove node

**Connecting nodes**: Click on a node terminal and drag to another terminal

**Save the flowchart layout**: Click "Save as…" to save the layout to a new file. You must specify the file extension as ".fc". If you save this file within the *"flowcharts" directory* of your project it will show up in the *Welcome Window* when you open your project.

**Note:** This does not save the data, use the *Save* node to save data.

**Warning:** Due to a weird Qt or pyqtgraph bug certain parameter values (such as those in drop-down menus) can't be saved. Similarly, parameters values are lost when you save to an existing .fc file. If you're interested take a look at `pyqtgraphCore.WidgetGroup`. Anyways you shouldn't be using the flowchart layout to save this information, that's what the History Trace in Transmission objects is for.

**Load an .fc file**: Click the "Load" button.

**Reset View button**: Reset the view, for example if you zoom out or pan too far.

##### 1.23.1 Video Tutorial

Part 5 - 9 of the Main Tutorial series also provide various examples for how the flowchart can be used: https://www.youtube.com/playlist?list=PLgofWiw2s4REPxH8bx8wZo_6ca435OKqg

##### 1.23.2 Transmission

*API Reference*

Almost every node uses a Transmission object for input and output. A Transmission is basically a DataFrame and a History Trace (analysis log) of the data within the DataFrame.

**Transmission DataFrame**

The Transmission DataFrame is created from your *Project DataFrame* (or sub-DataFrame) by the *Load_Proj_DF node*. This initial DataFrame will contain the same columns as your Project DataFrame, and a new column named **_RAW_CURVE**. Each element (row) in the **_RAW_CURVE** column is a 1-D numerical array representing a single raw curve extracted from an ROI.

A new column named **_BLOCK_** is also added which contains the UUID for logging the analysis history of this newly created block of DataFrame rows, known as a *data block*. This allows you to merge Transmissions (see *Merge node*) and maintain their independent analysis logs prior to the merge.

**Naming conventions for DataFrame columns according to the data types**

- *numerical data*: single leading underscore ( _ ). All caps if produced by a flowchart node.
- *categorial data*: no leading underscore. All caps if produced by flowhchart node.
- *special cases*: Peak detection data are placed in a column named **peaks_bases** where each element is a DataFrame.
- *uuid data*: has uuid or UUID in the name

**Note:** _BLOCK_ is an exception, it contains UUIDs not numerical data.

**History Trace**

The History Trace of a Transmission is a log containing the discrete analysis steps, known as operations, along with their parameters and any other useful information. When a flowchart node performs an operation it stores the output(s) data in the Transmission DataFrame and appends the operation parameters to this log. A seperate log is kept for each data block present in the Transmission DataFrame.

##### 1.23.3 Console

You have direct access to the data within the nodes through the console in the flowchart. To show the console go to View -> Console.

**See also:**

If you are unfamiliar with the console see the overview on *Consoles*

Call `get_nodes()` to view a dict of all nodes in the flowchart. You can access the output Transmission in most nodes through the attribute *t*. You can access the transmission dataframe through `t.df`.

**See also:**

See the *Transmission API* for more information. Sources for the nodes at mesmerize/pyqtgraphCore/flowchart/library.

**Example, directly accessing DataFrame elements through the flowchart console**

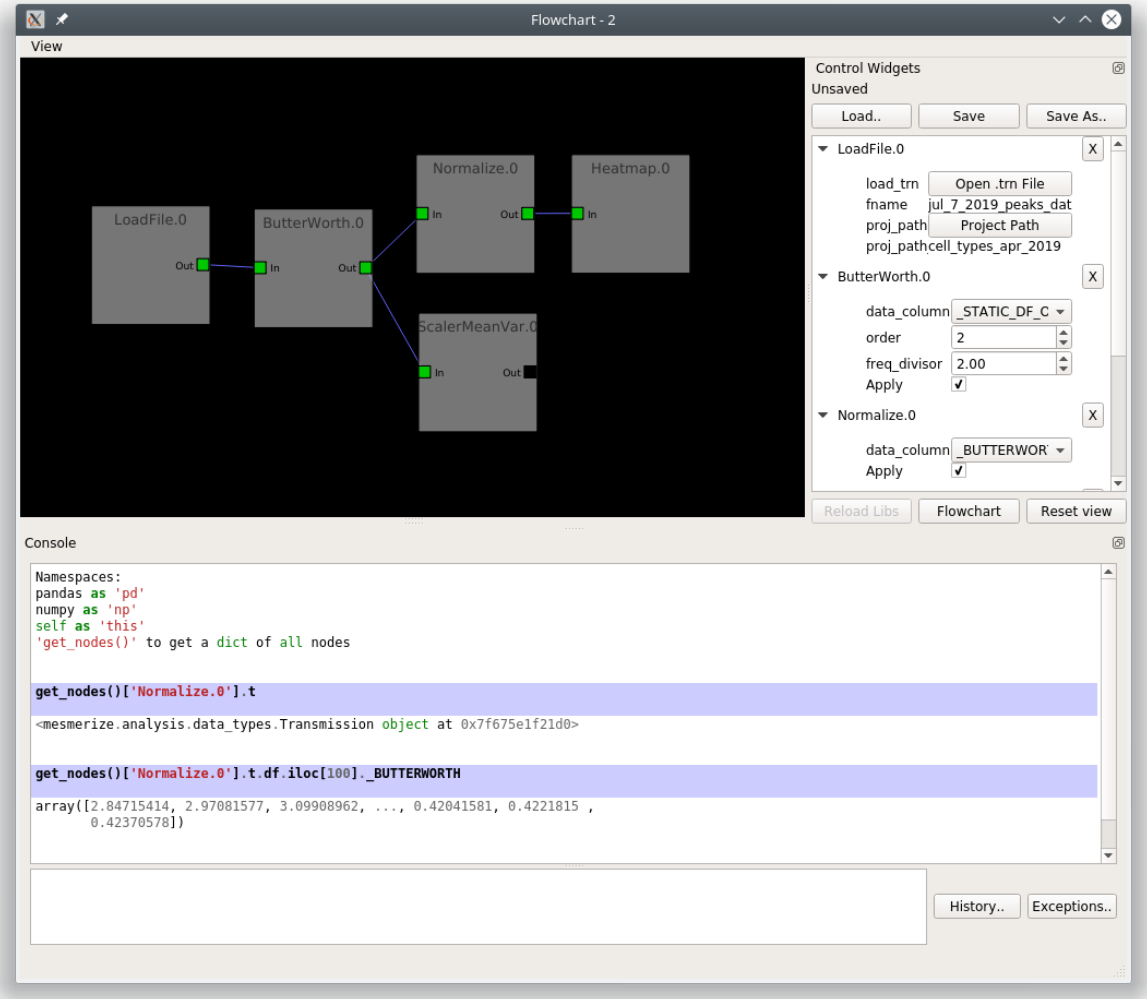


#### 1.23.4 Transmission Files

You can save a Transmission files using the *Save node* and work with the data directly in scripts, jupyter notebooks etc. You can also save them through the flowchart console (and plot consoles) through *Transmission.to_hdf5*.

##### Working with Transmission files

Load a saved Transmission instance using *Transmission.from_hdf5*

```
>>> from mesmerize import Transmission
>>> from uuid import UUID

# load transmission file
>>> t = Transmission.from_hdf5('/share/data/temp/kushal/data.trn')
<mesmerize.analysis.data_types.Transmission at 0x7f4d42f386a0>
```

(continues on next page)

(continued from previous page)

```

# The DataFrame is always the 'df' attribute
>>> t.df.head()

                                         CurvePath  ... FCLUSTER_LABELS
0  curves/a2-_-1-_-843c2d43-75f3-421a-9fef-483d1e...  ...               8
1  curves/brn3b_a6-_-2-_-21557a64-6868-4ff4-8db1-...  ...               4
2  curves/brn3b_a6-_-2-_-21557a64-6868-4ff4-8db1-...  ...               5
3  curves/brn3b_day1_3-_-2-_-ff3e95df-0e15-495c-9...  ...               8
4  curves/brn3b_day1_3-_-2-_-ff3e95df-0e15-495c-9...  ...               6

[5 rows x 27 columns]

# the `df` is just a pandas dataframe
# View a list of samples in the current file
>>> t.df.SampleID.unique()

array(['a2-_-1', 'a5-_-1', 'brn3b_a6-_-2', 'brn3b_day1_3-_-2',
    'brn3b_day1_a1-_-2', 'brn3b_day1_a2-_-2', 'brn3b_day1_a4-_-2',
    'brn3b_day2_a1-_-2', 'brn3b_day2_a1-_-t', 'brn3b_day2_a10-_-2',
    'brn3b_day2_a2-_-1', 'brn3b_day2_a2-_-3', 'brn3b_day2_a8-_-1',
    'cesa_a1-_-1', 'cesa_a1-_-2', 'cesa_a1_jan_2019-_-1',
    'cesa_a1_jan_2019-_-2', 'cesa_a2-_-2', 'cesa_a6-_-1',
    'cesa_a7-_-1', 'cesa_a7-_-2', 'cesa_a8-_-1', 'cesa_a9-_-1',
    'cng_ch4_day1_a2-_-t1', 'cng_ch4_day1_a2-_-t2',
    'cng_ch4_day2_a4-_-t1', 'dmrt1_day1_a2-_-2', 'dmrt1_day1_a4-_-t2',
    'dmrt1_day1_a5-_-', 'dmrt1_day1_a6-_-t', 'dmrt1_day1_a6-_-t2',
    'dmrt1_day2_a1-_-t1', 'dmrt1_day2_a1-_-t2', 'dmrt1_day2_a2-_-t1',
    'dmrt1_day2_a3-_-t1', 'dmrt1_day2_a3-_-t2', 'dmrt1_day2_a4-_-t1',
    'dmrt1_day2_a4-_-t2', 'hnk1_a5-_-2', 'hnk1_a6-_-1', 'hnk1_a7-_-1',
    'hnk1_a7-_-2', 'hnk1_a8-_-1', 'pc2_a10-_-1', 'pc2_a11-_-1',
    'pc2_a13-_-1', 'pc2_a14-_-1', 'pc2_a15-_-1', 'pc2_a16-_-1',
    'pc2_a9-_-1', 'pde9_day1_a2-_-2', 'pde9_day1_a3-_-1',
    'pde9_day1_a4-_-1', 'pde9_day1_a4-_-2', 'pde9_day2_a2-_-t2',
    'pde9_day2_a2-_-t4', 'pde9_day2_a4-_-t1', 'pde9_day2_a4-_-t2',
    'pde9_day2_a4-_-t3', 'pde9_day2_a5-_-t1', 'pde9_day2_a5-_-t2',
    'pde9_day2_a6-_-t1', 'pde9_day2_a7-_-t1', 'pde9_day2_a7-_-t2'],
    dtype=object)

# Show data associated with a single sample
>>> t.df[t.df['SampleID'] == 'brn3b_day1_a1-_-2']

                                         CurvePath  ... FCLUSTER_LABELS
6   curves/brn3b_day1_a1-_-2-_-d3c5f225-7039-4abd-...  ...               6
7   curves/brn3b_day1_a1-_-2-_-d3c5f225-7039-4abd-...  ...               6
8   curves/brn3b_day1_a1-_-2-_-d3c5f225-7039-4abd-...  ...               5
9   curves/brn3b_day1_a1-_-2-_-d3c5f225-7039-4abd-...  ...               7
10  curves/brn3b_day1_a1-_-2-_-d3c5f225-7039-4abd-...  ...               5

# View the data associated with one ROI
# the `uuid_curve` is a unique identifier for each curve/ROI
>> t.df[t.df['SampleID'] == 'brn3b_day1_a1-_-2'].iloc[0]
```

(continues on next page)

(continued from previous page)

```
CurvePath                curves/brn3b_day1_a1-_-2-_-d3c5f225-7039-4abd-...
ImgInfoPath              images/brn3b_day1_a1-_-2-_-d3c5f225-7039-4abd-...
ImgPath                  images/brn3b_day1_a1-_-2-_-d3c5f225-7039-4abd-...
ImgUUID                               d3c5f225-7039-4abd-a7a1-5e9ef2150013
ROI_State                {'roi_xs': [554, 553, 553, 552, 552, 551, 551,...
SampleID                                               brn3b_day1_a1-_-2
anatomical_location                                                 palp
cell_name                                                           palp
comments                                                        untagged
date                                                     20190425_110103
dorso_ventral_axis                                              untagged
misc                                                                  {}
morphology                                                      untagged
promoter                                                           brn3b
rostro_caudal_axis                                              untagged
stimulus_name                                                 [untagged]
uuid_curve                            f44fbd3d-6eaa-4e19-a677-496908565fde
_RAW_CURVE               [81.41972198848178, 75.61356993008134, 70.0493...
meta                     {'origin': 'AwesomeImager', 'version': '4107ff...
stim_maps                                                       [[None]]
_BLOCK_                               3e069e2d-d012-47ee-830c-93d85197e2f4
_SPLICE_ARRAYS           [2.646593459501195, 1.8252819116136887, 1.7422...
_NORMALIZE               [0.0681729940259753, 0.06533186950232853, 0.06...
_RFFT                    [443.19357880089615, -66.8777897472859, 55.244...
_ABSOLUTE_VALUE          [443.19357880089615, 66.8777897472859, 55.2443...
_LOG_TRANSFORM           [2.646593459501195, 1.8252819116136887, 1.7422...
FCLUSTER_LABELS                                                        6
Name: 6, dtype: object

# Show the ROI object data
>>> t.df[t.df['SampleID'] == 'brn3b_day1_a1-_-2'].iloc[0]['ROI_State']

{'roi_xs': array([554, 553, 553, 552, 552, 551, 551, 551, 551, 550, 550, 550, 549,
    548, 547, 547, 546, 546, 545, 545, 544, 543, 543, 542, 541, 541,
    540, 540, 539, 539, 538, 537, 536, 535, 534, 533, 532, 531, 531,
    530, 529, 528, 527, 527, 526, 526, 525, 525, 525, 524, 524, 523,
    522, 522, 521, 521, 520, 521, 521, 521, 521, 521, 522, 522, 522,
    522, 522, 522, 522, 522, 521, 521, 521, 521, 521, 521, 522, 523,
    524, 524, 525, 525, 525, 526, 526, 527, 528, 528, 529, 529, 529,
    530, 530, 531, 532, 532, 533, 534, 535, 535, 536, 536, 537, 538,
    539, 540, 540, 541, 541, 542, 542, 543, 544, 545, 546, 546, 547,
    548, 548, 549, 549, 549, 549, 550, 550, 550, 550, 551, 551, 551,
    552, 552, 552, 553, 553, 553, 554, 554, 554, 553, 554, 554, 554,
    554, 554]),
'roi_ys': array([155, 156, 156, 157, 157, 158, 159, 160, 160, 161, 162, 162, 162,
        162, 163, 163, 164, 164, 165, 165, 165, 166, 166, 166, 167, 167,
        167, 166, 167, 167, 167, 167, 167, 167, 167, 167, 167, 168, 168,
        168, 168, 168, 168, 167, 167, 166, 166, 165, 164, 164, 163, 163,
        163, 162, 162, 161, 161, 160, 160, 159, 158, 157, 156, 156, 155,
        154, 153, 152, 151, 150, 150, 149, 148, 147, 146, 145, 144, 144,
        144, 144, 143, 143, 142, 141, 141, 140, 140, 140, 139, 139, 138,
```

(continues on next page)

(continued from previous page)

```
        137, 137, 136, 136, 136, 135, 135, 135, 136, 136, 137, 137, 137,
        137, 137, 138, 138, 138, 137, 137, 136, 136, 136, 136, 137, 137,
        137, 138, 138, 139, 140, 141, 141, 142, 143, 144, 144, 145, 146,
        146, 147, 148, 148, 149, 150, 150, 151, 151, 152, 152, 153, 154,
        155, 155]),
'curve_data': (array([   0,    1,    2, ..., 2996, 2997, 2998]),
array([ 81.41972199,  75.61356993,  70.04934883, ..., 195.4416283 ,
        184.8844155 , 174.76708104])),
'tags': {'anatomical_location': 'palp',
'cell_name': 'palp',
'morphology': 'untagged'},
'roi_type': 'CNMFROI',
'cnmf_idx': 2}
```

##### View History Log

Transmissions have a *history_trace* attribute which is an instance of *HistoryTrace*.

Use the *get_data_block_history* and *get_operations_list* methods to view the history log of a data block.

```
# To view the history log, first get the block UUID of the dataframe row of which you want the history log

# Block UUIDs are stored in the _BLOCK_ column
>>> bid = t.df.iloc[10]._BLOCK_
>>> bid

'248a6ece-e60e-4a09-845e-188a5199d262'

# Get the history log of this data block
# HistoryTrace.get_operations_list() returns a list of operations, without parameters
# HistoryTrace.get_data_block_history() returns the operations list with the parameters
>>> t.history_trace.get_operations_list(bid)

['spawn_transmission',
 'splice_arrays',
 'normalize',
 'rfft',
 'absolute_value',
 'log_transform',
 'splice_arrays',
 'fcluster']

# View the entire history log with all params
>>> t.history_trace.get_data_block_history(bid)

[{'spawn_transmission': {'sub_dataframe_name': 'neuronal',
'dataframe_filter_history': {'dataframe_filter_history': ['df[~df["promoter"].isin([\'cesa\', \'hnk1\'])]',
    'df[~df["promoter"].isin([\'cesa\', \'hnk1\'])]',
    'df[~df["cell_name"].isin([\'not_a_neuron\', \'non_neuronal\', \'untagged\', \'ependymal\'])]']}}},
```

(continues on next page)

(continued from previous page)

```
{'splice_arrays': {'data_column': '_RAW_CURVE',
'start_ix': 0,
'end_ix': 2990,
'units': 'time'}},
{'normalize': {'data_column': '_SPLICE_ARRAYS', 'units': 'time'}},
{'rfft': {'data_column': '_NORMALIZE',
'frequencies': [0.0,
    0.0033444816053511705,
    0.0033444816053511705,
    0.006688963210702341,
    ...

# Get the parameters for the 'fcluster' operation
>>> fp = t.history_trace.get_operation_params(bid, 'fcluster')

# remove the linkage matrix first so we can view the other params
>>> fp.pop('linkage_matrix');fp

{'threshold': 8.0,
 'criterion': 'maxclust',
 'depth': 1,
 'linkage_params': {'method': 'complete',
 'metric': 'wasserstein',
 'optimal_ordering': True}}

# Draw the analysis history as a graph
# This will open your defeault pdf viewer with the graph
>>> t.history_trace.draw_graph(bid, view=True)

# If you are using the API to perform analysis on
# transmission files, you can use the `HistoryTrace`
# to log the analysis history
# For example, add a number `3.14` to all datapoints in a curve
>>> t.df['_RAW_CURVE'] = t.df['_RAW_CURVE'].apply(lambda x: x + 3.14)

# Append the analysis log
>>> t.history_trace.add_operation(data_block_id='all', operation='addition', parameters={
↪'value': 3.14}
```

#### 1.24 Nodes

##### 1.24.1 Data

**These nodes are for performing general data related operations**

#### LoadFile

*Source*

Loads a save Transmission file. If you have a Project open it will automatically set the project path according to the open project. Otherwise you must specify the project path. You can specify a different project path to the project that is currently open (this is untested, weird things could happen). You should not merge Transmissions originating from different projects.

**Note:** You can also load a saved Transmission file by dragging & dropping it into the Flowchart area. It will create a LoadFile node with the name of the dropped.

| Terminal | Description |
|---|---|
| Out | Transmission loaded from the selected file. |

| Parameters | Description |
|---|---|
| load_trn | Button to choose a .trn file (Transmission) to load |
| proj_trns | Load transmission file located in the project's "trns" directory |
| proj_path | Button to select the Mesmerize project that corresponds to the chosen .trn file. |

**Note:** The purpose of specifying the Project Path when you load a save Transmission file is so that interactive plots and the *Datapoint Tracer* can find raw data that correspond to datapoints.

#### LoadProjDF

*Source*

Load the entire Project DataFrame (root) of the project that is currently open, or a sub-DataFrame that corresponds a tab that you have created in the *Project Browser*.

**Output Data Column** *(numerical)*: _RAW_CURVE

Each element in this output column contains a 1-D array representing the trace extracted from an ROI.

| Terminal | Description |
|---|---|
| Out | Transmission created from the Project DataFrame or sub-DataFrame. |

| Parameters | Description |
|---|---|
| DF_Name | DataFrame name. List correponds to *Project Browser* tabs. |
| Update | Re-create Transmission from corresponding *Project Browser* tab. |
| Apply | Process data through this node |

**Note:** The **DF_Name** options do not update live with the removal or creation of tabs in the *Project Browser*, you must create a new node to reflect these types of changes.

##### Save

*Source*

Save the input Transmission to a file so that the Transmission can be used re-loaded in the Flowchart for later use.

**Usage:** Connect an input Transmission to this node's **In** terminal, click the button to choose a path to save a new file to, and then click the Apply checkbox to save the input Transmission to the chosen file.

| Terminal | Description |
|---|---|
| In | Transmission to be saved to file |

| Parameters | Description |
|---|---|
| saveBtn | Button to choose a filepath to save the Transmission to. |
| Apply | Process data through this node |

**Note:** You must always save a Transmission to a new file (pandas with hdf5 exihibts weird behavior if you overwrite, this is the easiest workaround). If you try to overwrite the file you will be presented with an error saying that the file already exists.

##### Merge

*Source*

Merge multiple Transmissions into a single Transmission. The DataFrames of the individual Transmissions are concatenated using pandas.concat and History Traces are also merged. The History Trace of each indidual input Transmission is kept separately.

**Warning:** At the moment, if you create two separate data streams that originate from the same Transmission and then merge them at a later point, the analysis log (History Trace) of the individual data streams are not maintained. See the information about data blocks in the *Transmission*.

| Terminal | Description |
|---|---|
| In | Transmissions to be merged |
| Out | Merged Transmission |

##### ViewTransmission

*Source*

View the input Transmission object using the spyder Object Editor. For example you can explore the Transmission DataFrame and HistoryTrace.

##### ViewHistory

*Source*

View the HistoryTrace of the input Transmission in a nice Tree View GUI.

##### TextFilter

*Source*

Include or Exclude Transmission DataFrame rows according to a text filter in a categorical column.

**Usage Example:** If you want to select all traces that are from photoreceptor cells and you have a categorical column, named cell_types for example, containing cell type labels, choose "cell_type" as the *Column* parameter and enter "photoreceptor" as the *filter* parameter, and select *Include*. If you want to select everything that are not photoreceptors select *Exclude*.

**Note:** It is recommended to filter and group your data beforehand using the *Project Browser* since it allows much more sophisticated filtering.

| Terminal | Description |
|---|---|
| In | Input Transmission |
| Out | Transmission its DataFrame filtered accoring parameters |

| Parameters | Description |
|---|---|
| Column | Categorical column that contains the text filter to apply |
| filter | Text filter to apply |
| Include | Include all rows matching the text filter |
| Exclude | Exclude all rows matching the text filter |
| Apply | Process data through this node |

**HistoryTrace output structure:** Dict of all the parameters for this node

##### SpliceArrays

*Source*

Splice arrays derived in the specified numerical data column and place the spliced output arrays in the output column.

**Output Data Column** *(numerical)*: _SPLICE_ARRAYS

| Terminal | Description |
|---|---|
| In | Input Transmission |
| Out | Transmission with arrays from the input column spliced and placed in the output column |

| Parameters | Description |
|---|---|
| data_column | Numerical data column containing the arrays to be spliced |
| indices | The splice indices, "start_index:end_index" |
| Apply | Process data through this node |

##### DropNa

*Source*

Drop NaNs and Nones (null) from the Transmission DataFrame. Uses DataFrame.dropna and DataFrame.isna methods.

- If you choose "row" or "column" as axis, entire rows or columns will be dropped if any or all (see params) of the values are NaN/None.
- If you choose to drop NaNs/Nones according to a specific column, it will drop the entire row if that row has a NaN/None value for the chosen column.

| Terminal | Description |
|---|---|
| In | Input Transmission |
| Out | Transmission NaNs and None's removed according to the params |

| Parameters | Description |
|---|---|
| axis | Choose to rows, columns, or a rows according to a specific column. |
| how | *any:* Drop if any value in the row/column is NaN/None<br><br>*all:* Drop only if all values in the row/column are Nan/None<br><br>ignored if "axis" parameter is set to a specific column |
| Apply | Process data through this node |

##### NormRaw

`Source`

Scale the raw data such that the min and max values are set to the min and max values derived from the raw spatial regions of the image sequences they originate from. Only for CNMFE data.

The arrays in the **_RAW_CURVE** column are scaled and the output is placed in a new column named **_NORMRAW**

| Terminal | Description |
|---|---|
| In | Input Transmission |
| Out | Transmission with the result placed in the output column |

| Parameter | Description |
|---|---|
| option | Derive the raw min & max values from one of the following options:<br>*top_5:* Top 5 brightest pixels<br>*top_10:* Top 10 brighest pixels<br>*top_5p:* Top 5% of brightest pixels<br>*top_10p:* Top 10% of brightest pixels<br>*top_25p:* Top 25% of brightest pixels<br>*full_mean:* Full mean of the min and max array |
| Apply | Process data through this node |

**Note:** If the raw min value is higher than the raw max value the curve will be excluded in the output. You will be presented with a warning box with the number of curves that were excluded due to this.

#### 1.24.2 Display

**These nodes connect input Transmission(s) to various plots for visualization**

The actual Plot Widget instance that these nodes use can be accessed through the `plot_widget` attribute in the flowchart console.

For example

```
# Get a heatmap node that is named "Heatmap.0"
>>> hn = get_nodes()['Heatmap.0']

# the plot widget instance
>>> hn.plot_widget

<mesmerize.plotting.widgets.heatmap.widget.HeatmapTracerWidget object at 0x7f26e5d29678>
```

##### BeeswarmPlots

*Source*

Based on pqytgraph Beeswarm plots.

Visualize data points as a pseudoscatter and as corresponding Violin Plots. This is commonly used to visualize peak features and compare different experimental groups.

For information on the plot widget see *Beeswarm Plots*

| Terminal | Description |
|---|---|
| In | Input Transmission<br><br>The DataFrame column(s) of interest must have single numerical values, not arrays |

##### Heatmap

*Source*

Used for visualizing numerical arrays in the form of a heatmap. Also used for visualizing a hieararchical clustering tree (dendrogram) along with a heatmap with row order corresponding to the order leaves of the dendrogram.

For information on the plot widget see *Heat Plot*

| Terminal | Description |
|---|---|
| In | Input Transmission<br><br>The arrays in the DataFrame column(s) of interest **must** be of the same length |

**Note:** Arrays in the DataFrame column(s) of interest **must** be of the same length. If they are not, you must splice them using the *SpliceArrays* node.

##### CrossCorr

*Source*

Perform Cross-Correlation analysis. For information on the plot widget see *CrossCorrelation Plot*

##### Plot

*Source*

For information on the plot widget see *Simple Plot*

A simple plot.

| Terminal | Description |
|---|---|
| In | Input Transmission |

| Parameters | Description |
|---|---|
| data_column | Data column to plot, must contain numerical arrays |
| Show | Show/hide the plot window |
| Apply | Process data through this node |

##### Proportions

*Source*

Plot stacked bar chart of one categorical variable vs. another categorical variable.

For information on the plot widget see *Proportions Plot*

##### ScatterPlot

*Source*

Create scatter plot of numerical data containing [X, Y] values

For information on the plot widget see *Scatter Plot*

---

#### 1.24.3 Signal

**Routine signal processing functions**

I recommend this book by Tom O'Haver if you are unfamiliar with basic signal processing: https://terpconnect.umd.edu/~toh/spectrum/TOC.html

##### Butterworth

Source

Creates a Butterworth filter using scipy.signal.butter and applies it using scipy.signal.filtfilt.

The Wn parameter of scipy.signal.butter is calculated by dividing the sampling rate of the data by the *freq_divisor* parameter (see below).

**Output Data Column** *(numerical)*: _BUTTERWORTH

| Terminal | Description |
|---|---|
| In | Input Transmission |
| Out | Transmission with filtered signals in the output data column |

| Parameters | Description |
|---|---|
| data_column | Data column containing numerical arrays to be filtered |
| order | Order of the filter |
| freq_divisor | Divisor for dividing the sampling frequency of the data to get Wn |
| Apply | Process data through this node |

##### SavitzkyGolay

*Source*

Savitzky Golay filter. Uses scipy.signal.savgol_filter.

**Output Data Column** *(numerical)*: _SAVITZKY_GOLAY

| Terminal | Description |
|---|---|
| In | Input Transmission |
| Out | Transmission with filtered signals in the output data column |

| Parameters | Description |
|---|---|
| data_column | Data column containing numerical arrays to be filtered |
| window_length | Size of windows for fitting the polynomials. Must be an odd number. |
| polyorder | Order of polynomials to fit into the windows. Must be less than *window_length* |
| Apply | Process data through this node |

##### PowSpecDens

##### Resample

*Source*

Resample the data in numerical arrays. Uses scipy.signal.resample.

**Output Data Column** *(numerical)*: _RESAMPLE

| Terminal | Description |
|---|---|
| In | Input Transmission |
| Out | Transmission with resampled signals in the output data column |

| Parameters | Description |
|---|---|
| data_column | Data column containing numerical arrays to be resampled |
| Rs | New sampling rate in *Tu* units of time. |
| Tu | Time unit |
| Apply | Process data through this node |

**Note:** If Tu = 1, then Rs is the new sampling rate in Hertz.

##### ScalerMeanVariance

*Source*

Uses tslearn.preprocessing.TimeSeriesScalerMeanVariance

**Output Data Column** *(numerical)*: _SCALER_MEAN_VARIANCE

| Terminal | Description |
|---|---|
| In | Input Transmission |
| Out | Transmission with scaled signals in the output column |

| Parameters | Description |
|---|---|
| data_column | Data column containing numerical arrays to be scaled |
| mu | Mean of the output time series |
| std | Standard Deviation of the output time series |
| Apply | Process data through this node |

**Note:** if mu = 0 and std = 1, the output is the z-score of the signal.

##### Normalize

*Source*

Normalize the signal so that all values are between 0 and 1 based on the min and max of the signal.

**Output Data Column** *(numerical)*: _NORMALIZE

| Terminal | Description |
|---|---|
| In | Input Transmission |
| Out | Transmission with scaled signals in the output column |

| Parameters | Description |
|---|---|
| data_column | Data column containing numerical arrays to be scaled |
| Apply | Process data through this node |

##### RFFT

*Source*

Uses scipy.fftpack.rfft. “Discrete Fourier transform of a real sequence”

**Output Data Column** *(numerical)*: _RFFT

| Terminal | Description |
|---|---|
| In | Input Transmission |
| Out | Transmission with the RFT of signals in the output column |

| Parameters | Description |
|---|---|
| data_column | Data column containing numerical arrays |
| Apply | Process data through this node |

#### iRFFT

*Source*

Uses scipy.fftpack.irfft. "inverse discrete Fourier transform of real sequence x"

**Output Data Column** *(numerical)*: _IRFFT

#### PeakDetect

*Source*

Simple Peak Detection using derivatives. The "Differentiation" chapter of Tom O'Haver's book has a section on Peak Detection which I recommend reading. https://terpconnect.umd.edu/~toh/spectrum/TOC.html

**Output Data Column** *(DataFrame)*: peaks_bases

**See also:**

*Peak Editor GUI*

| Terminal | Description |
|---|---|
| Derivative | Transmission with derivatives of signals. Must have **_DERIVATIVE** column.<br><br>It's recommended to use a derivative from a normalized filtered signal. |
| Normalized | Transmission containing Normalized signals, used for thresholding<br><br>See *Normalize* node |
| Curve | Transmission containing original signals.<br><br>Usually not filtered to avoid distortions caused by filtering |
| PB_Input *(optional)* | Transmission containing peaks & bases data (peaks_bases column).<br><br>Useful for visualizing a saved Transmission that has peaks & bases data |
| Out | Transmission with the detected peaks & bases as DataFrames in the output column |

> **Warning:** The *PB_Input* terminal overrides all other terminals. Do not connect inputs to *PB_Input* and other terminals simultaneously.

| Parameter | Description |
|---|---|
| data_column | Data column of the input *Curve* Transmission for placing peaks & bases onto |
| Fictional_Bases | Add bases to beginning and end of signal if first or last peak is lonely |
| Edit | Open Peak Editor GUI, see *Peak Editor* |
| SlopeThr | Slope threshold |
| AmplThrAbs | Absolute amplitude threshold |
| AmplThrRel | Relative amplitude threshold |
| Apply | Process data through this node |

##### PeakFeatures

*Source*

Compute peak features. The DataFrame of the ouput Transmission contains one row for each peak.

| Output Data Column | Description |
|---|---|
| _pf_peak_curve | array representing the peak |
| _pf_ampl_rel_b_ix_l | peak amplitude relative to its left base |
| _pf_ampl_rel_b_ix_r | peak amplitude relative to its right base |
| _pf_ampl_rel_b_mean | peak amplitude relative to the mean of its bases |
| _pf_ampl_rel_zero | peak amplitude relative to zero |
| _pf_area_rel_zero | Simpson's Rule Integral of the curve |
| _pf_area_rel_min | Simpson's Rule Integral relative to the minimum value of the curve<br>Substracts the minimum values of the peak curve before computing the integral |
| _pf_rising_slope_avg | slope of the line drawn from the left base to the peak |
| _pf_falling_slope_avg | slope of the line drawn from the right base to the peak |
| _pf_duration_base | distance between the left and right base |
| _pf_p_ix | index of the peak maxima in the parent curve |
| _pf_uuid | peak UUID |
| _pf_b_ix_l | index of the left base in the parent curve |
| _pf_b_ix_r | index of the right base in the parent curve |

**See also:**

`mesmerize/analysis/compute_peak_features` for the code that computes the peak features.

| Terminal | Description |
|---|---|
| In | Input Transmission. Must contain *peak_bases* column that contains peak_bases DataFrames. |
| Out | Transmission with peak features in various output columns |

| Parameter | Description |
|---|---|
| data_column | Data column containing numerical arrays from which to compute peak features. |
| Apply | Process data through this node |

**Warning:** If there are issues with a particular peak a user warning will be displayed in the terminal that Mesmerize is running and the peak will be ignored. This happens when a peak is 1) not flanked by bases on both sides, 2) a peak or base is out of bounds for the parent curve from teh chosen *data_column* or 3) other index issues w.r.t. the peak. In the terminal, the number after the progress bar will show the index of the parent curve, for example here the parent curve is 319: 41%| | 319/771. The index of the offending peak within the parent curve will be printed below the progress bar along with a statement that may specific the issue with the peak.

#### 1.24.4 Math

**Nodes for performing basic Math functions**

##### Derivative

*Source*

Computes the first derivative.

**Output Data Column** *(numerical)*: _DERIVATIVE

| Terminal | Description |
|---|---|
| In | Input Transmission |
| Out | Transmission with the derivative placed in the output column |

| Parameter | Description |
|---|---|
| data_column | Data column containing numerical arrays |
| Apply | Process data through this node |

##### TVDiff

*Source*

Based on Numerical Differentiation of Noisy, Nonsmooth Data. Rick Chartrand. (2011).. Translated to Python by Simone Sturniolo.

##### XpowerY

*Source*

Raises each element of the numerical arrays in the data_column to the exponent Y

**Output Data Column** *(numerical)*: _X_POWER_Y

| Terminal | Description |
|---|---|
| In | Input Transmission |
| Out | Transmission with the result placed in the output column |

| Parameter | Description |
|---|---|
| data_column | Data column containing numerical arrays |
| Y | Exponent |
| Apply | Process data through this node |

##### AbsoluteValue

*Source*

Element-wise absolute values of the input arrays. Computes root mean squares if input arrays are complex.

**Output Data Column** *(numerical)*: _ABSOLUTE_VALUE

| Terminal | Description |
|---|---|
| In | Input Transmission |
| Out | Transmission with the result placed in the output column |

| Parameter | Description |
|---|---|
| data_column | Data column containing numerical arrays |
| Apply | Process data through this node |

##### LogTransform

*Source*

Perform Logarithmic transformation of the data.

**Output Data Column** *(numerical)*: _LOG_TRANSFORM

| Terminal | Description |
|---|---|
| In | Input Transmission |
| Out | Transmission with the result placed in the output column |

| Parameter | Description |
|---|---|
| data_column | Data column containing numerical arrays |
| transform | *log10*: Base 10 logarithm<br><br>*ln*: Natural logarithm<br><br>*modlog10*: $sign(x) * \log_{10}(\|x\| + 1)$<br><br>*modln*: $sign(x) * \ln(\|x\| + 1)$ |
| Apply | Process data through this node |

##### ArrayStats

*Source*

Perform a few basic statistical functions.

**Output Data Column** *(numerical)*: Customizable by user entry

Output data are single numbers, not arrays

| Terminal | Description |
|---|---|
| In | Input Transmission |
| Out | Transmission with the result placed in the output column |

The desired function is applied to each 1D array in the *data_column* and the output is placed in the Output Data Column.

| Parameter | Description |
|---|---|
| data_column | Data column containing numerical arrays |
| function | *amin*: Return the minimum of the input array<br>*amax*: Return the maximum of the input array<br>*nanmin*: Return the minimum of the input array, ignore NaNs<br>*nanmax*: Return the maximum of the input array, ignore NaNs<br>*ptp*: Return the range (max - min) of the values of the input array<br>*median*: Return the median of the input array<br>*mean*: Return the mean of the input array<br>*std*: Return the standard deviation of the input array<br>*var*: Return the variance of the input array<br>*nanmedian*: Return the median of the input array, ignore NaNs<br>*nanmean*: Return the mean of the input array, ignore NaNs<br>*nanstd*: Return the standard deviation of the input array, ignore NaNs<br>*nanvar*: Return the variance of the input array, ignore NaNs |
| group_by *(Optional)* | Group by a categorial variable, for example get the mean array of a group |
| group_by_sec *(Optional)* | Group by a secondary categorical variable |
| output_col | Enter a name for the output column |
| Apply | Process data through this node |

##### ArgGroupStat

*Source*

Group by a categorial variable and return the value of any other column based on a statistic. Basically creates sub-dataframes for each group and then returns based on the sub-dataframe.

Group by column "group_by" and return value from column "return_col" where data in *data_column* fits "stat"

**Output Data Column** *(Any)*: ARG_STAT

| Terminal | Description |
|---|---|
| In | Input Transmission |
| Out | Transmission with the result placed in the output column |

| Parameter | Description |
|---|---|
| data_column | Data column containing single numbers (not arrays for now) |
| group_by | Group by column (categorical variables) |
| return_col | Return value from this column (any data) |
| stat | "max" or "min" |
| Apply | Process data through this node |

##### ZScore

*Source*

Compute Z-Scores of the data. Uses scipy.stats.zscore. The input data are are divided into groups according to the *group_by* parameter. Z-Scores are computed for the data in each group with respect to the data only in that group.

**Output Data Column** *(numerical)*: _ZSCORE

| Terminal | Description |
|---|---|
| In | Input Transmission |
| Out | Transmission with the result placed in the output column |

| Parameter | Description |
|---|---|
| data_column | Input data column containing numerical arrays |
| group_by | Categorial data column to group by. |
| Apply | Process data through this node |

##### LinRegress

*Source*

Basically uses scipy.stats.linregress

Performs Linear Regression on numerical arrays and returns slope, intercept, r-value, p-value and standard error

| Terminal | Description |
|---|---|
| In | Input Transmission |
| Out | Transmission with the result placed in the output column |

| Parameter | Description |
|---|---|
| data_column | Data column containing 1D numerical arrays. The values are used as the y values and indices as the x values for the regression |

**Output Columnns:** Single numbers, *_SLOPE*, *_INTERCEPT*, *_R-VALUE*, *_P-VALUE*, *_STDERR* as decribed in scipy.stats.linregress

##### 1.24.5 Biology

**Nodes for some biologically useful things which I couldn't categorize elsewhere**

###### ExtractStim

*Source*

Extract the portions of a trace corresponding to stimuli that have been temporally mapped onto it. It outputs one row per stimulus period.

**Note:** Stimulus extraction is currently quite slow, will be optimized after some planned changes in the Transmission object.

| Output Data Column | Description |
|---|---|
| ST_TYPE | Stimulus type, corresponds to your *Project Config* |
| ST_NAME | Name of the stimulus |
| _ST_CURVE | The extracted array based on the parameters |
| _ST_START_IX | Start index of the stimulus period in the parent curve |
| _ST_END_IX | End index of the stimulus period in the parent curve |
| ST_uuid | UUID assigned for the extracted stimulus period |

| Parameter | Description |
|---|---|
| data_column | Data column containing the signals to be extracted based on the stimulus maps |
| Stim_Type | Type of stimulus to extract |
| start_offset | Offset the start index of the stimulus mapping by a value (in frames) |
| end_offset | Offset the end index of the stimulus mapping by a value (in frames) |
| zero_pos | Zero index of the extracted signal<br><br>*start_offset*: extraction begins at the *start_offset* value, stops at the *end_offset*<br><br>*stim_end*: extraction begins at the end of the stimulus, stops at the *end_offset*.<br><br>*stim_center*: extraction begins at the midpoint of the stimulus period plus the *start_offset*, stops at *end_offset* |

###### DetrendDFoF

Source

Uses the detrend_df_f function from the CaImAn library. This node does not use any of the numerical data in a Transmission DataFrame to compute the detrended $\Delta F/F_0$. It directly uses the CNMF output data for the Samples that are present in the Transmission DataFrame.

**Output Data Column** *(numerical)*: _DETREND_DF_O_F

###### StaticDFoFo

*Source*

Perform $\frac{F-F_0}{F_0}$ without a rolling window. $F$ is an input array and $F_0$ is the minimum value of the input array.

**Output Data Column** *(numerical)*: _STATIC_DF_O_F

| Terminal | Description |
|---|---|
| In | Input Transmission |
| Out | Transmission with the result placed in the output column |

| Parameter | Description |
|---|---|
| data_column | Data column containing numerical arrays |
| Apply | Process data through this node |

###### StimTuning

*Source*

Stimulus Tuning analysis. For more information see *Stimulus Tuning Plot*

##### 1.24.6 Clustering

###### KShape

*Source*

Perform KShape clustering. For more information see *KShape plot*.

###### KMeans

*Source*

Basically sklearn.cluster.KMeans.

##### 1.24.7 Hierarchical

These nodes allow you to perform Hierarchical Clustering using scipy.cluster.hierarchy.

If you are unfamiliar with Hierarchical Clustering I recommend going through this chapter from Michael Greenacre: http://www.econ.upf.edu/~michael/stanford/maeb7.pdf

**Note: Some of these nodes do not use Transmission objects for some inputs/outputs.**

###### Linkage

*Source*

Compute a linkage matrix which can be used to form flat clusters using the *FCluster* node.

Based on scipy.cluster.hierarchy.linkage

| Terminal | Description |
|---|---|
| In | Input Transmission |
| Out | dict containing the Linkage matrix and parameters, **not a Transmission object** |

| Parameters | Description |
|---|---|
| data_column | Numerical data column used for computing linkage matrix |
| method | linkage method |
| metric | metric for computing distance matrix |
| optimal_order | minimize distance between successive leaves, more intuitive visualization<br><br>Click here for more info |
| Apply | Process data through this node |

###### FCluster

*Source*

"Form flat clusters from the hierarchical clustering defined by the given linkage matrix."

Based on scipy.cluster.hierarchy.fcluster

**Output Data Column** *(categorial)*: FCLUSTER_LABELS

| Terminal | Description |
|---|---|
| Linkage | Linkage matrix, output from *Linkage* node. |
| Data | Input Transmission, usually the same input Transmission used for the *Linkage* node. |
| IncM *(optional)* | Inconsistency matrix, output from *Inconsistent* |
| Monocrit *(optional)* | Output from *MaxIncStat* or *MaxInconsistent* |
| Out | Transmission with clustering data that can be visualized using the *Heatmap* |

**Parameters:** Exactly as desribed in scipy.cluster.hierarchy.fcluster

**HistoryTrace output structure:** Dict of all the parameters for this node, as well as the parameters used for creating the linkage matrix and the linkage matrix itself from the *Linkage node*.

###### Inconsistent

###### MaxIncStat

###### MaxInconsistent

---

##### 1.24.8 Transform

Nodes for transforming data

###### LDA

*Source*

Perform Linear Discriminant Analysis. Uses sklearn.discriminant_analysis.LinearDiscriminantAnalysis

| Terminal | Description |
|---|---|
| train_data | Input Transmission containing the training data |
| predict | Input Transmission containing data on which to predict |
| T | Transmission with Transformed data and decision function. Output columns outlined below:<br>**_LDA_TRANSFORM:** The transformed data, can be visualized with a *Scatter Plot* for instance<br>**_LDA_DFUNC:** Decision function (confidence scores). Can be visualized with a *Heatmap* |
| coef | Transmission with LDA Coefficients. Output columns outlined below:<br>**classes:** The categorical labels that were trained against<br>**_COEF:** LDA Coefficients (weight vectors) for the classes. Can be visualized with a *Heatmap* |
| means | Transmission with LDA Means. Output columns outlined below:<br>**classes:** The categorical labels that were trained against<br>**_MEANS:** LDA means for the classes. Can be visualized with a *Heatmap* |
| predicted | Transmission containing predicted class labels for the data.<br>The class labels are placed in a column named **LDA_PREDICTED_LABELS**<br>The names of the class labels correspond to the labels from the training labels<br>*optional* |

| Parameter | Description |
|---|---|
| train_data | Single or multiple data columns that contain the input features. |
| labels | Data column containing categorical labels to train to |
| solver | *svd:* Singular Value Decomposition<br>*lsqr:* Least Squares solution<br>*eigen*: Eigen decomposition |
| shrinkage | Can be used with *lsqr* or *eigen* solvers. |
| shrinkage_val | shrinkage value if *shrinkage* is set to "value" |
| n_components | Number of components to output |
| tol | Tolereance threshold exponent. The used value is 10^<tol> |
| score | Displays mean score of the classification (read only) |
| predict_on | Single or multiple data columns that contain the data that are used for predicting on<br>Usually the same name as the data column(s) used for the training data.<br>*optional* |

**HistoryTrace output structure:** Dict of all the parameters for this node

#### 1.25 Examples

##### 1.25.1 Datasets

You can view examples of flowcharts in the demo dataset or one of the other datasets associated with the paper:

Demo dataset: https://doi.org/10.6084/m9.figshare.11370183

C. intestinalis dataset: https://doi.org/10.6084/m9.figshare.10289162

C. elegans dataset: https://doi.org/10.6084/m9.figshare.10287113

PVC-7 as a Mesmerize dataset: https://doi.org/10.6084/m9.figshare.10293041

##### 1.25.2 Video Tutorials

##### 1.25.3 Screenshots

Flowchart screenshots from the C. intestinalis dataset.

###### Z-score

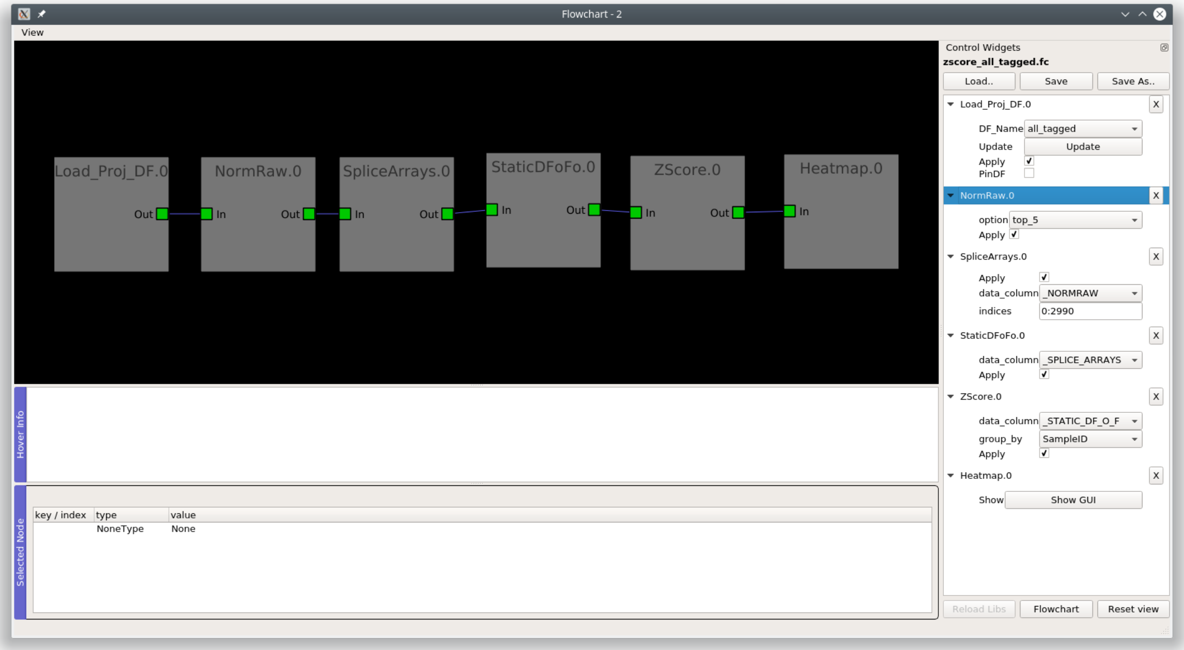

#### Peak detection

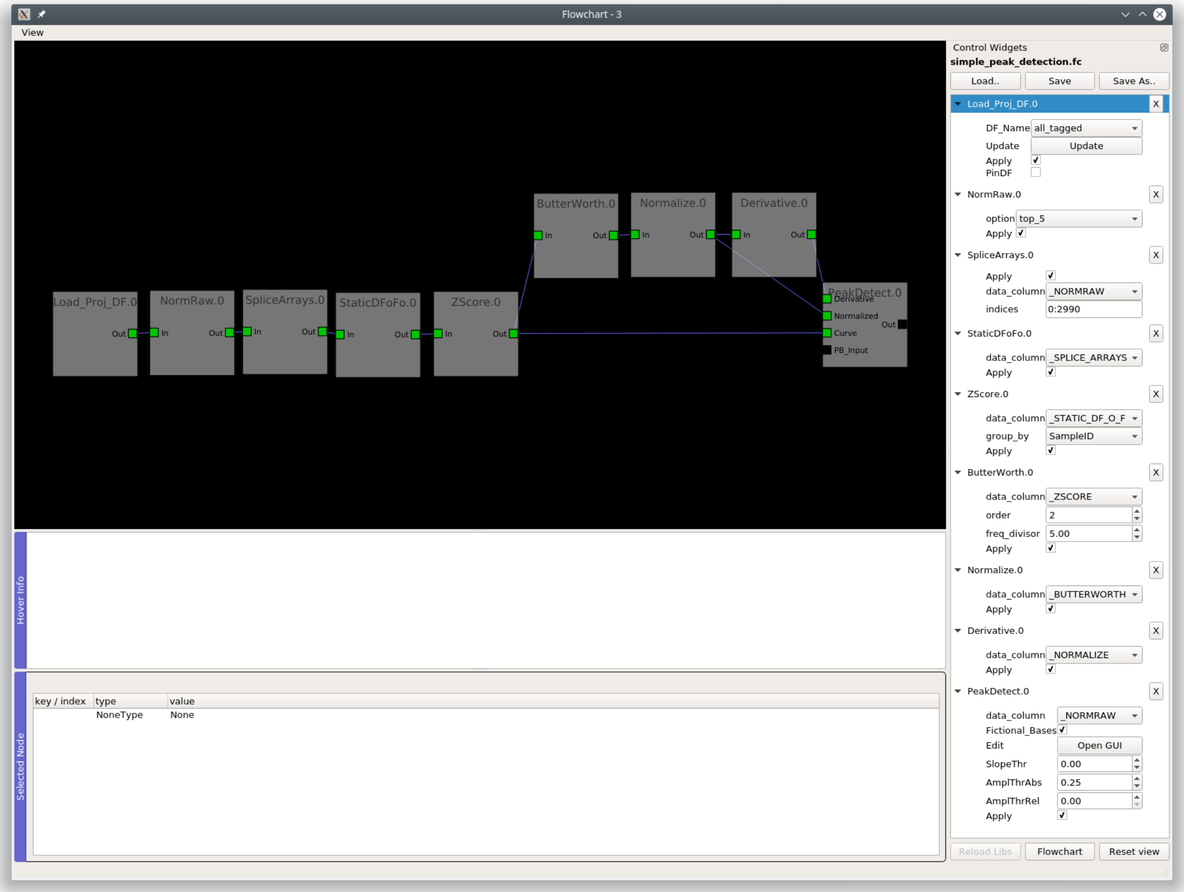

##### Hierarchical clustering

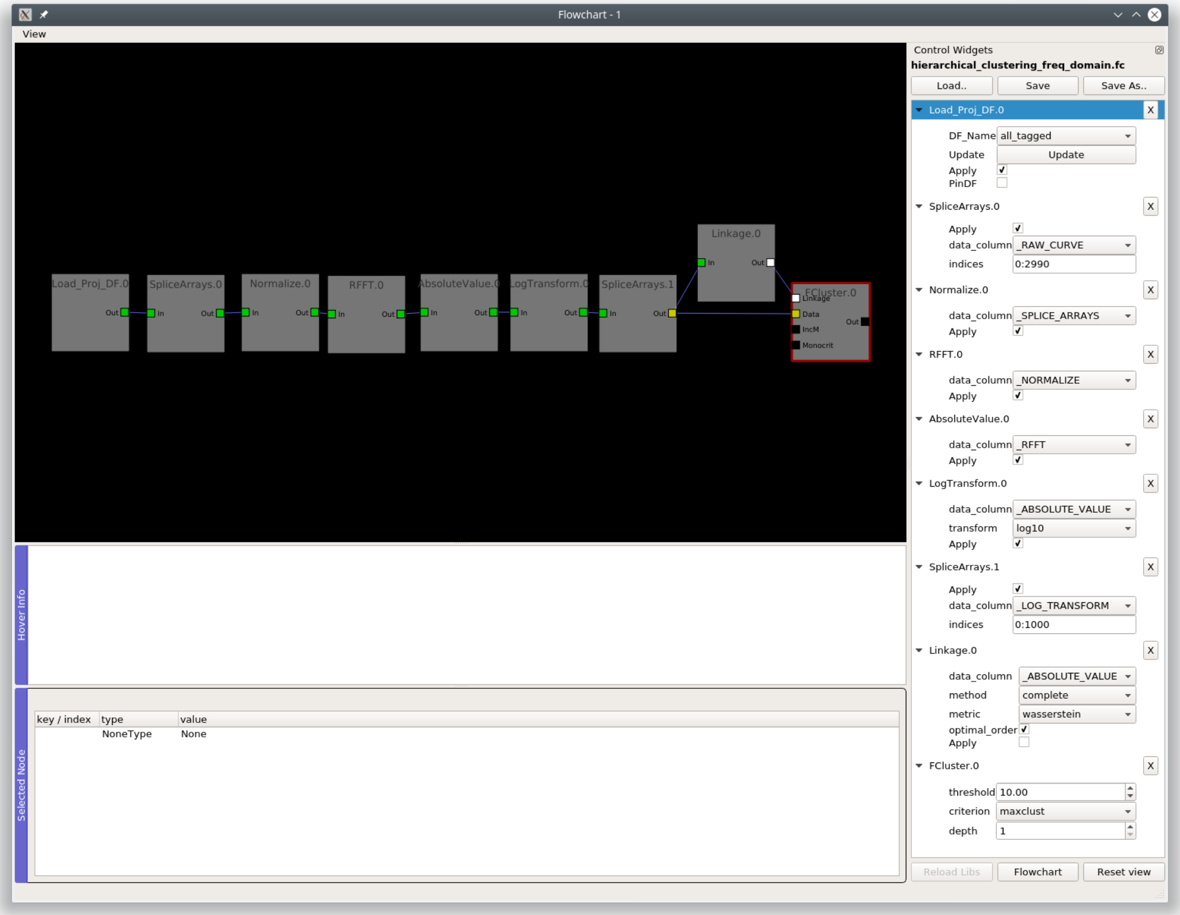


#### 1.26 Beeswarm

Used for visualization of data points using a pseudo-scatter and violin plots.

**Layout**

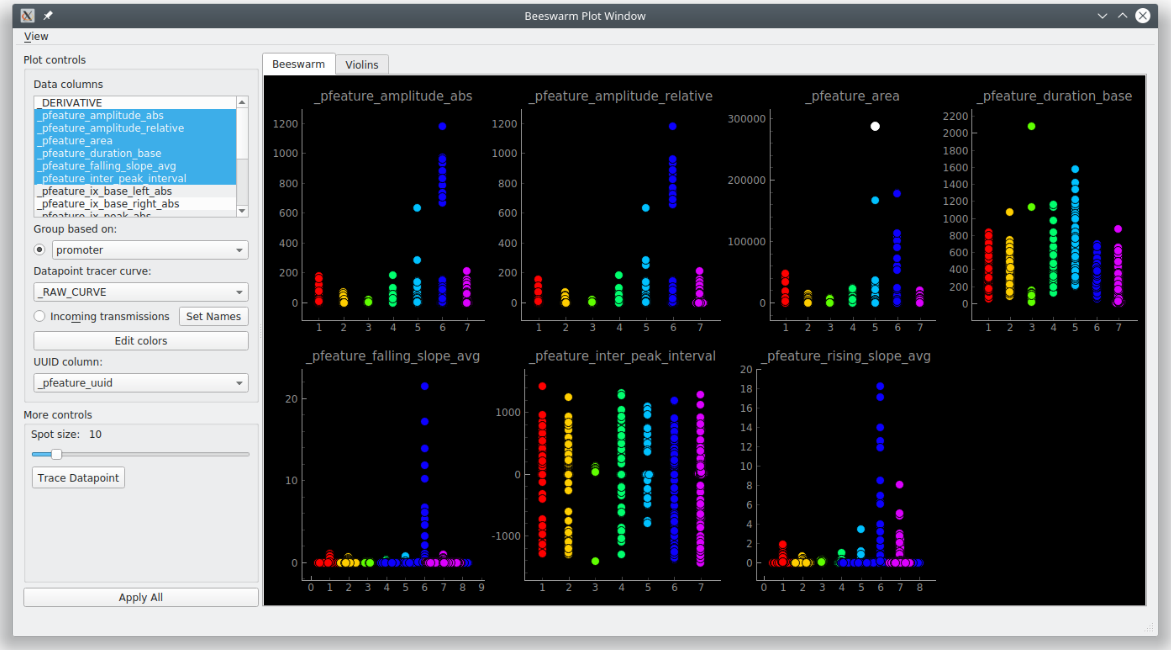


You can click on individual datapoints and view the associated data using the *Datapoint Tracer*. To show the Datapoint Tracer, in the menubar go to View -> Live datapoint tracer

##### 1.26.1 Parameters

| Parameter | Description |
|---|---|
| Data columns | Multi-select data columns to plot<br><br>They must have single numerical values, not arrays |
| Group based on | Categorical data column used for grouping the data |
| Datapoint tracer curve | Data column, containing numerical arrays, that is shown in the *Datapoint Tracer* |
| UUID column | Column containing the UUIDs that correspond to the data in the selected data column(s) |
| Apply all | Apply the plot parameters and draw the plot |

#### 1.27 Consoles

Currently the *Heatmap*, *Scatter*, *Proportions plot* and *KShape* follow a uniform structure allowing internal access to the data and plot axes. Refer to their *Base API*. For example, through their consoles you can access the *Transmission* containing data for the current plot, manually save the plot, etc.

#### 1.28 Cross Correlation

Explore Cross-correlation functions of all curves from a sample. Normalized cross-correlations are computed using `tslearn.cycc.normalized_cc`

This is an interactive widget. You can click on the individual cells in the heatmap to view the individual curves, the cross-correlation function of the two curves, and the spatial localization of the ROI that they originate from.

##### 1.28.1 Layout

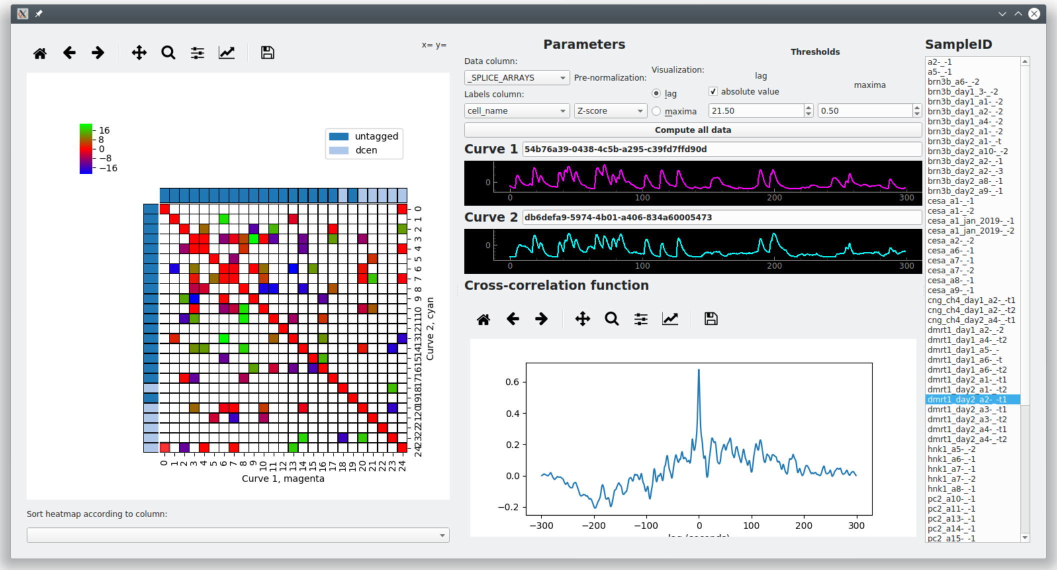


**Left**: Lag or Maxima Matrix (see below) with thresholds applied and visualized as a heatmap. When you click on the individual cells it will open/update the *Datapoint Tracer* according to the two curves the cell corresponds to.

**Top Center**: Parameters.

**Center**: When you click on a cell in the heatmap you will see Curve 1 (x-axis of heatmap), and Curve 2 (y-axis of heatmap) and their cross-correlation function. **The units are in seconds for all of these**

**Right**: List of Samples. Click on a Sample to select it as the current sample.

##### 1.28.2 Lag Matrix

**Computed as follows:**

1. A 2D array is created where each element is a cross-correlation function (represented by a 1D numerical array).
2. The x-distance (time) between zero and the global maxima of the cross-correlation function (called $lag$) is computed for each of these elements.
3. The 2D array of cross-correlation functions is reduced to a 2D array of these $lag$ values.

The result is a matrix where each element is the x-distance between zero and the global maxima of the cross-correlation of the two curves the element represents.

##### 1.28.3 Maxima Matrix

Similar to computation of the Lag Matrix above, but instead of using the $lag$ between zero and the global maxima it uses the y-value of the global maxima.

##### 1.28.4 Parameters

**Data column:** The data column, containing *numerical* arrays, that are used as the input curves for computing cross-correlations.

**Labels column:** The labels column, containing *categorical* labels, that are used for the row and column labels in the heatmaps.

**Pre-normalization:** Option to perform 0 - 1 Normalization (Same method as the *Normalize*) or Z-Score of the input curves prior to computing their cross-correlation functions.

**Compute all data:** Appy the parameters and compute cross-correlation data for all Samples in the DataFrame of the input transmission.

###### Thresholds

Apply thresholds for $lag$ and the maxima value. The colormap limits of the heatmap are set according to this threshold and all data under are set to white on the heatmap (you can still click and explore them).

Thresholds are applied live onto the heatmap.

#### 1.29 Datapoint Tracer

*API Reference*

The Datapoint Tracer is attached to many plots, allowing you to interactively explore the data associated to the data-points. You can explore the analysis history, the spatial localization of the ROI it originates from, associated numerical or categorical data, and view an additional numerical column (such as the raw trace).

The Datapoint Tracer is embedded in some plots, and in others you can open it by going to View -> Live Datapoint Tracer.

##### 1.29.1 Video Tutorial

This tutorial shows how the Heatmap plot can be used along with the Datapoint Tracer during the latter half of this tutorial.

Part 5, 6 & 8 of the main tutorial series also show how the Datapoint Tracer can be used along with other types of plots: https://www.youtube.com/playlist?list=PLgofWiw2s4REPxH8bx8wZo_6ca435OKqg

##### 1.29.2 Layout

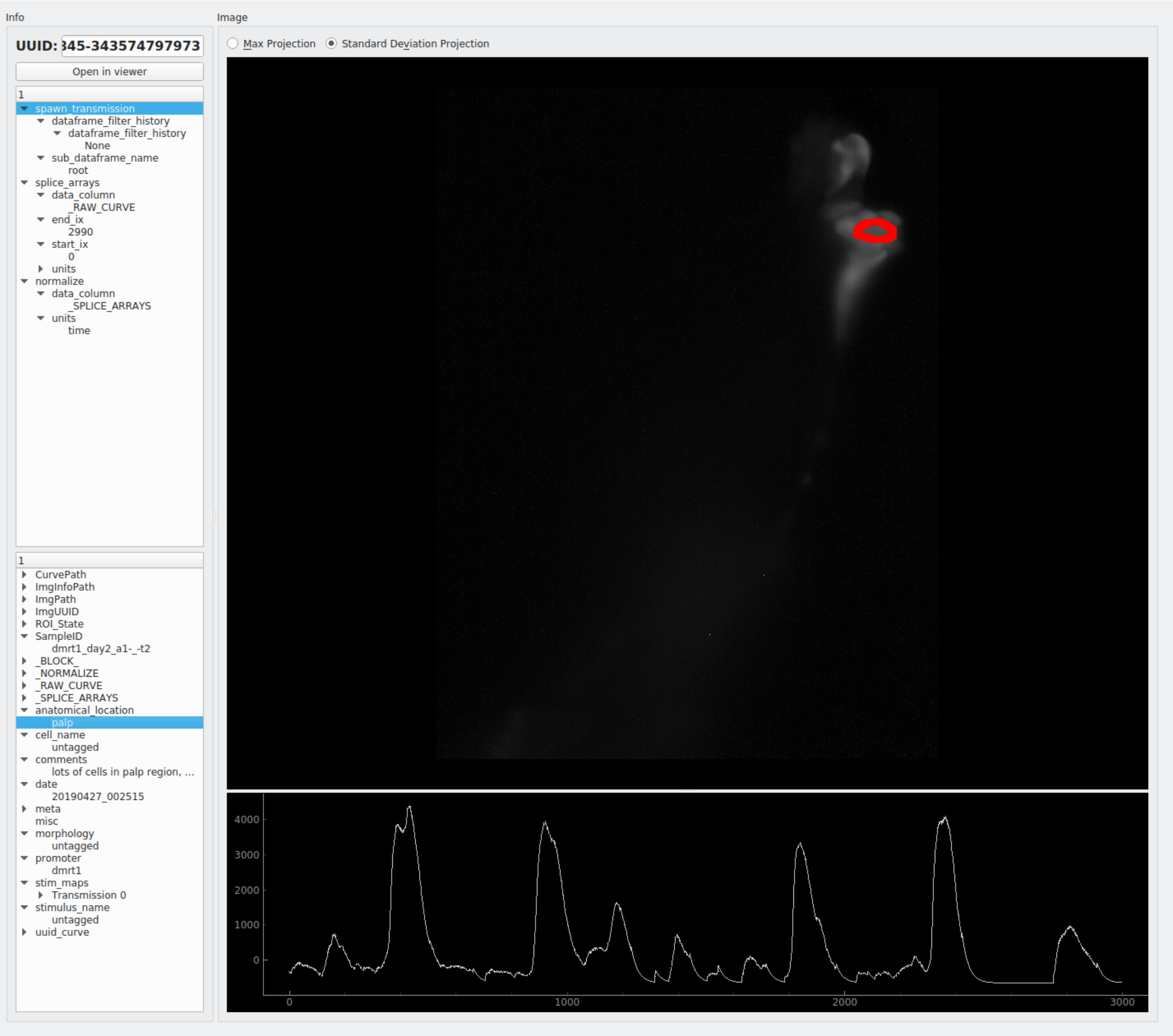


**Top right**: Max Projection or Standard Deviation Project of the image sequence.

**Bottom right**: Numerical data, based on the *"DPT Curve column"* that the user has specified in the plot controls. If exploring peak feature based data the temporal span of the peak will be highlighted.

**Top left**: Analysis log, a ordered list of operations and their parameters.

**Bottom left**: All other data associated with this datapoint (the data present in the other columns of the row this datapoint is present in, see *Transmission*)

**Open in viewer button**: Open the parent Sample of the current datapoint in the viewer.

### 1.30 Heatmap

*API Reference*

**Note:** This plot can be saved in an interactive form, see *Saving plots*

Visualize numerical arrays in the form of a heatmap. Also used for visualization of Hierarchical clusterting dendrograms. *Datapoint Tracer* is embedded.

#### 1.30.1 Video Tutorial

This tutorial shows how the Heatmap plot can be used along with the Datapoint Tracer during the latter half of this tutorial.

#### 1.30.2 Layout

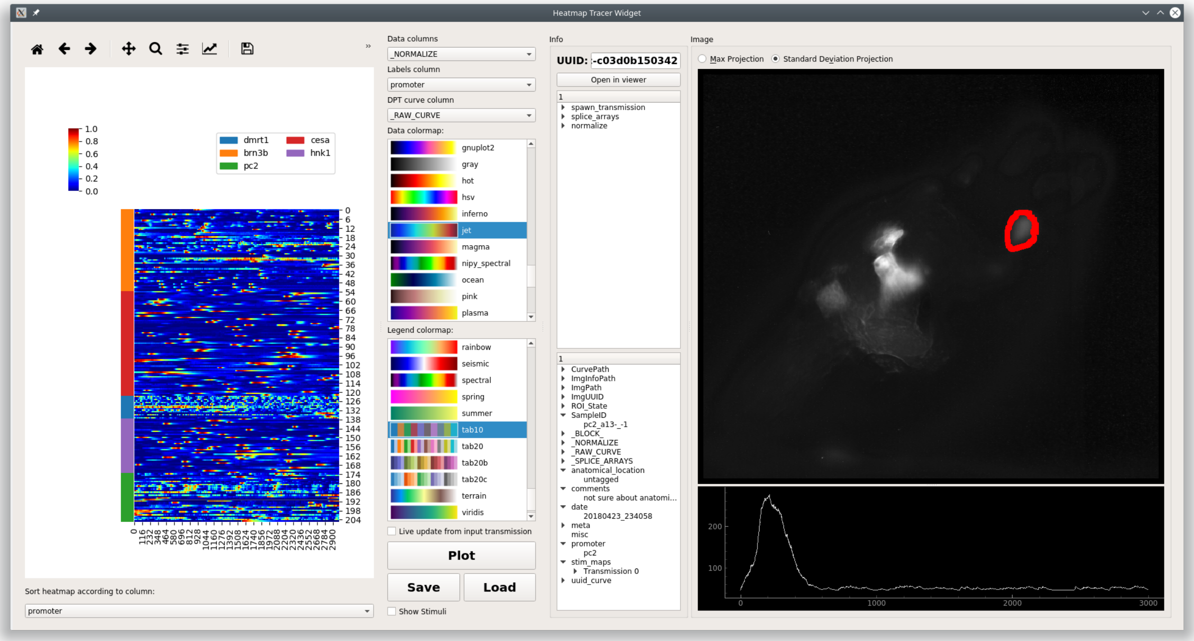


**Left:** The heatmap. Clicking the heatmap highlights the selected row and upates the *Datapoint Tracer*. Right click on the heatmap to clear the selection highlight on the heatmap. You can zoom and pan the heatmap using the tools above the plot area. You can zoom/pan in the legend and heatmap. The up and down keys on your keyboard can be used to move the current row selection.

**Bottom left:** Set the row order of the heatmap according to a categorical column.

**Middle:** Plot controls.

**Very bottom:** Status label - displays any issues that were raised while setting the plot data. Click on the status label to see more information.

##### 1.30.3 Parameters

**Data column:** Data column, numerical arrays, that contain the data for the heatmap. Each row of this data column (a 1D array) is represented as a row on the heatmap.

**Labels column:** Column containing categorical labels that are used to create the row legend for the heatmap.

**DPT curve column:** Data column, containing numerical arrays, that is shown in the *Datapoint Tracer*.

**Data colormap:** Colormap used for representing the data in the heatmap. Default is 'jet'.

**Legend colormap:** Colormap used for the row legend.

**Live update from input transmission:** If checked this plots receives live updates from the *flowchart*.

**Plot:** Updates data input from the flowchart.

**Save:** *Save the plot data and state in an interactive form*

**Load:** Load a plot that has been saved as a ".ptrn" file.

**Layout to visualize Hierarchical Clustering**

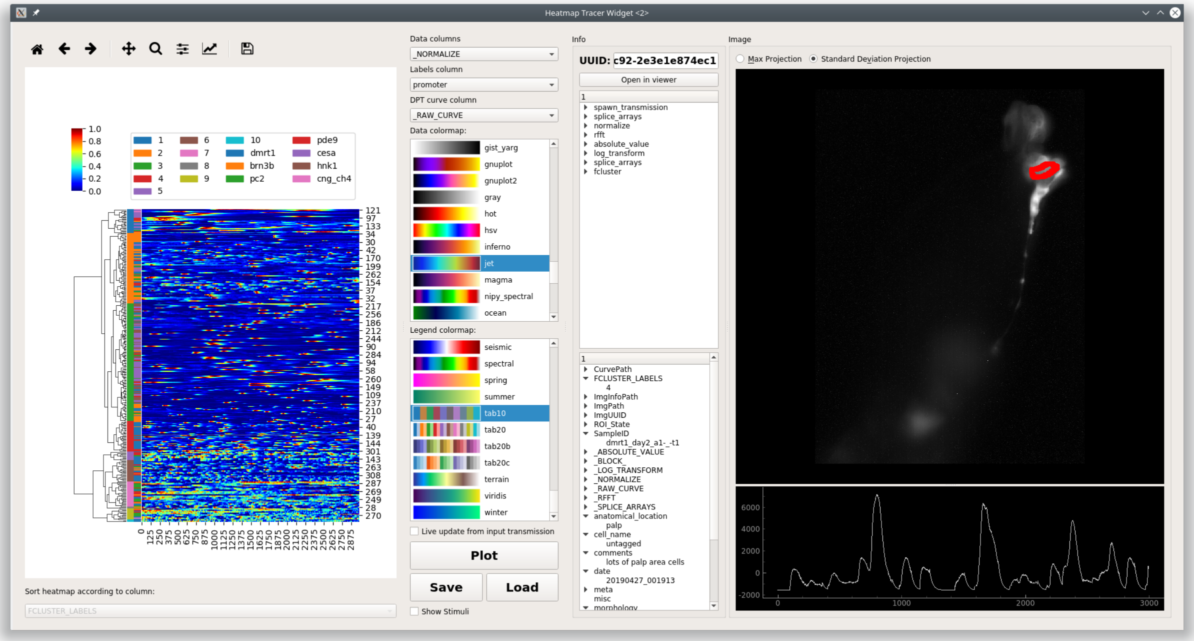


This plot widget can also be used to visualize a dendrogram on top of a heatmap of data.

The differences are:

1. There are two legend bars
   - Left: Cluster label
   - Right: Corresponds to Labels column parameter.
2. You can also zoom/pan the dendrogram in addition to the legends and heatmap.
3. Sorting the heatmap rows is disabled because this wouldn't make sense

##### 1.30.4 Console

You can directly access the heatmap widget through the console. This is useful for plot customization and exporting with specific parameters.

Toggle the console's visibility by clicking on the "Show/Hide Console" button at the bottom of the controls.

**See also:**

*API Reference*

###### Namespace

| reference | Description |
|---|---|
| this | The higher-level *HeatmapTracerWidget* instance, i.e. the entire widget |
| this.transmission | Current input *Transmission* |
| get_plot_area() | Returns the lower-level *Heatmap* variant instance, basically the actual plot area |
| get_plot_area().plot | Returns the seaborn ClusterGrid instance containing the axes |
| get_plot_area().fig | Returns the matplotlib Figure instance |

**Attributes of** `get_plot_area().plot`

For example, the heatmap axes object can be retrieved through `get_plot_area().plot.ax_heatmap`. See the usage examples.

| | |
|---|---|
| ax_heatmap | Heatmap axes |
| ax_row_dendrogram | Row dendrogram axes |
| ax_col_dendrogram | Used for the legend |
| cax | Colorbar axes |

###### Examples

###### Export

**See also:**

matplotlib API for: Figure.savefig, Figure.set_size_inches, Figure.get_size_inches

```
# Desired size (width, height)
size = (2.0, 2.5)

# Get the figure
fig = get_plot_area().fig

# original size to reset the figure after we save it
orig_size = fig.get_size_inches()

#Set the desired size
fig.set_size_inches(size)

# Save the figure as a png file with 1200 dpi
fig.savefig('/share/data/temp/kushal/amazing_heatmap.png', dpi=1200, bbox_inches='tight',
↪ pad_inches=0)
```

(continues on next page)

(continued from previous page)

```

# Reset the figure size and draw()
fig.set_size_inches(orig_size)
get_plot_area().draw()
```

**Note:** The entire plot area might go gray after the figure is reset to the original size. I think this is a Qt-matplotlib issue. Just resize the window a bit and the plot will be visible again!

**Warning:** From my experience I have not been able to open clustermap SVG files saved with very high DPI (600+). Even with 32 cores & 128GB of RAM both inkscape and illustrator just hang ¯_(ツ)_/¯. Try png or other formats.

##### x tick labels

**See also:**

matplotlib.axes.Axes.set_xticklabels, matplotlib.axes.Axes.set_xticks.

If the data are in the time domain:

```
from mesmerize.analysis import get_sampling_rate
import numpy as np

# Get the sampling rate of the data
sampling_rate = get_sampling_rate(this.transmission)

# Number of frames currently displayed in the heatmap
num_frames = get_plot_area().data.shape[1]

# Set an appropriate interval
interval = 30 # This is in seconds, not frames

# Get the recording time in seconds
recording_time = int(num_frames / sampling_rate)

# Set the new ticks
get_plot_area().plot.ax_heatmap.set_xticks(np.arange(0, num_frames, interval * sampling_
↪rate))

# Set the tick labels
# You can change the fontsize here
get_plot_area().plot.ax_heatmap.set_xticklabels(np.arange(0, recording_time, interval),␣
↪fontdict={'fontsize': 4})

# Set a title for the x axis. You can change the fontsize here
get_plot_area().plot.ax_heatmap.set_xlabel('Time (seconds)', fontdict={'fontsize': 6})

# Draw the plot with these changes
get_plot_area().draw()
```

**Note:** You may need to resize the dock widget that the plot is present in to display the newly drawn plot, this is a Qt-matplotlib issue.

If the data are in the frequency domain:

```
from mesmerize.analysis import get_frequency_linspace
import numpy as np

# Get frequency linspace and Nyquist frequency
freqs, nf = get_frequency_linspace(this.transmission)

# Get the number of frequencies currently shown in the heatmap
num_freqs = get_plot_area().data.shape[1]

# The max frequency currently display in the heatmap
max_freq = freqs[num_freqs - 1]

# Set an appropriate interval
interval = 0.25 # This is in Hertz

# Set the tick labels
# Set the new ticks
get_plot_area().plot.ax_heatmap.set_xticks(np.arange(0, num_freqs, (num_freqs * interval) / max_freq))

# You can change the fontsize here
get_plot_area().plot.ax_heatmap.set_xticklabels(np.arange(0, max_freq, interval), fontdict={'fontsize': 4})

# Set a title for the x axis. You can change the fontsize here
get_plot_area().plot.ax_heatmap.set_xlabel('Frequency (Hertz)', fontdict={'fontsize': 6})

# Draw the plot with these changes
get_plot_area().draw()
```

**Note:** You may need to resize the dock widget that the plot is present in to display the newly drawn plot, this is a Qt-matplotlib issue.

###### Colorbar label

```
get_plot_area().plot.cax.set_title('norm. z-score', x=-0.25, y=0.65, fontdict={'fontsize
↪': 6}, rotation=90)
get_plot_area().draw()
```

###### Axes visibility

Hide/show legend

```
get_plot_area().plot.ax_col_dendrogram.set_visible(False)
get_plot_area().draw()
```

Hide/show y axis (similar for x axis)

```
get_plot_area().plot.ax_heatmap.get_yaxis().set_visible(False)
get_plot_area().draw()
```

Hide/show colorbar

```
get_plot_area().plot.cax.set_visible(False)
get_plot_area().draw()
```

#### 1.31 KShape

Perform KShape clustering.

I recommend reading the paper on it: Paparrizos, John, and Luis Gravano. "k-Shape: Efficient and Accurate Clustering of Time Series." In Proceedings of the 2015 ACM SIGMOD International Conference on Management of Data, pp. 1855-1870. ACM, 2015.

This GUI uses the tslearn.clustering.KShape implementation.

**See also:**

*API reference*

**Note:** This plot can be saved in an interactive form, see *Saving plots*

**Layout**

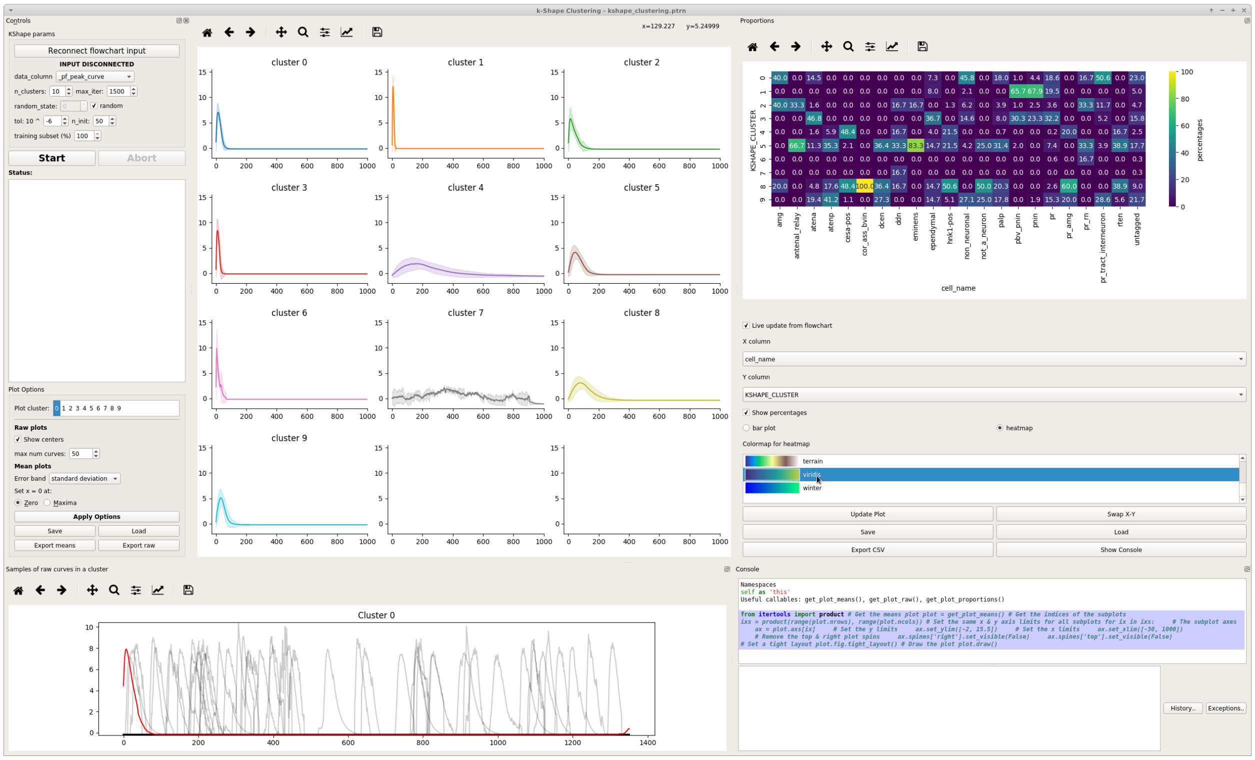


**Left:** KShape parameters and Plot parameters

**Bottom left:** Plot of a random sample of input data from a cluster.

**Center:** Plot of cluster mean and either confidence interval, standard deviation, or neither. Uses seaborn.lineplot

**Right:** Proportions plot. Exactly the same as *Proportions*.

**Bottom Right:** Console

#### 1.31.1 KShape Parameters

The parameters and input data are simply fed to tslearn.clustering.KShape

Parameters outlined here are simply as they appear in the tslearn docs.

**data_column:** Input data for clustering.

**n_clusters:** Number of clusters to form.

**max_iter:** Maximum number of iterations of the k-Shape algorithm.

**tol:** Inertia variation threshold. If at some point, inertia varies less than this threshold between two consecutive iterations, the model is considered to have converged and the algorithm stops.

**n_init:** Number of times the k-Shape algorithm will be run with different centroid seeds. The final results will be the best output of n_init consecutive runs in terms of inertia.

**random_state:** Generator used to initialize the centers. If an integer is given, it fixes the seed. Defaults to the global numpy random number generator.

**training subset:** The subset of the input data that are used for used for training. After training, the predictions are fit on all the input data.

##### 1.31.2 Plot Options

**Plot cluster:** The cluster from which to plot random samples of input data in the bottom left plot

**Show centers:** Show the centroids returned by the KShape model

> **Warning:** There's currently an issue where cluster centroids don't appear to be index correctly. See https://github.com/rtavenar/tslearn/issues/114

**max num curves:** Maximum number of input data samples to plot

**Error band:** The type of data to show for the the error band in the means plots.

**set x = 0 at:** The zero position of a means plots with respect to the cluster members in the plot.

**Save:** *Save the plot data and state in an interactive form*

##### 1.31.3 Console

The console can be useful for formatting plots, inspecting the underlying data etc.

**See also:**

*API reference*

###### Namespace

| reference | Description |
|---|---|
| this | The higher-level *KShape* widget instance, i.e. the entire widget |
| this.transmission | Current input *Transmission* |
| get_plot_means() | Returns the means plot |
| get_plot_raw() | Returns the raw plot |
| get_plot_proportions() | Returns the proportions plot, which is an instance of *Proportions Widget* |

###### Examples

**See also:**

matplotlib Axes

###### Set axis ranges

Set equal x & y axis ranges for the means plots. Also removes the top & right spines.

```
from itertools import product

# Get the means plot
plot = get_plot_means()

# Get the indices of the subplots
ixs = product(range(plot.nrows), range(plot.ncols))
```

(continues on next page)

(continued from previous page)

```

# Set the same x & y axis limits for all subplots
for ix in ixs:

    # The subplot axes
    ax = plot.axs[ix]

    # Set the y limits
    ax.set_ylim([-2, 15.5])

    # Set the x limits
    ax.set_xlim([-30, 1000])

    # Remove the top & right plot spins
    ax.spines['right'].set_visible(False)
    ax.spines['top'].set_visible(False)

# Set a tight layout
plot.fig.tight_layout()

# Draw the plot
plot.draw()
```

**Note:** You may need to resize the dock widget that the plot is present in to display the newly drawn plot, this is a Qt-matplotlib issue.

##### x tick labels

Set the x tick labels in time units instead of frames

**See also:**

matplotlib.axes.Axes.set_xticklabels | matplotlib.axes.Axes.set_xticks.

```
import numpy as np
from itertools import product
from mesmerize.analysis import get_sampling_rate

# Get the sampling rate of the data
sampling_rate = get_sampling_rate(this.transmission)

# Get the padded number of frames that are shown in the plots
num_frames = this.cluster_centers.shape[1]

# Set an appropriate interval
interval = 5 # This is in seconds, not frames

# Convert the padded frame number to time units
total_time = int(num_frames / sampling_rate)

```

(continues on next page)

(continued from previous page)

```
ixs = product(range(4), range(3))

# Set these time units for all the means plots
# For the raw plots just remove the loop
for ix in ixs:
    # Get the axes
    ax = get_plot_means().axs[ix]

    # Set the new ticks
    ax.set_xticks(np.arange(0, num_frames, interval * sampling_rate))

    # Set the tick labels
    # You can change the fontsize here
    ax.set_xticklabels(np.arange(0, total_time, interval), fontdict={'fontsize': 4}, rotation=90)

    # Set a title for the x axis. You can change the fontsize here
    ax.set_xlabel('Time (seconds)', fontdict={'fontsize': 6})

    # Set ylabel as well
    ax.set_ylabel('z-score', fontdict={'fontsize': 6})

# Set a tight layout
get_plot_means().fig.tight_layout()

# Draw the plot with these changes
get_plot_means().draw()
```

**Note:** You may need to resize the dock widget that the plot is present in to display the newly drawn plot, this is a Qt-matplotlib issue.

##### Hide legend

Hide/show legend in the proportions plot

```
get_plot_proportions().ax.legend().set_visible(True)
get_plot_proportions().draw()
```

##### Export

You can export any of the plots with a specific size & DPI.

Replace the `get_<plot>().fig` on *line 5* with the desired plot.

**See also:**

matplotlib API for: Figure.savefig, Figure.set_size_inches, Figure.get_size_inches

```
# Desired size (width, height)
size = (7.0, 10.0)

# Get the figure
fig = get_<plot>().fig

# original size to reset the figure after we save it
orig_size = fig.get_size_inches()

#Set the desired size
fig.set_size_inches(size)

# Save the figure as an png file with 600 dpi
fig.savefig('/share/data/temp/kushal/amazing_shapes.png', dpi=600, bbox_inches='tight', pad_inches=0)

# Reset the figure size and draw
fig.set_size_inches(orig_size)
get_<plot>().draw()
```

**Note:** The entire plot area might go gray after the figure is reset to the original size. I think this is a Qt-matplotlib issue. Just resize the window a bit and the plot will be visible again!

#### 1.32 Peak Editor

Visualize and edit detected peaks & bases. This GUI is accessible through the *PeakDetect node*.

##### 1.32.1 Video Tutorial

##### 1.32.2 Usage

- Optimize your peaks/bases detection through the datastreams that feed into the *Derivative* and *Normalize* terminals of the parent *PeakDetect node*. For example, play with filtering parameters for the *ButterWorth node* or *SavitzkyGolay node*.
- Optimize amplitude thresholds of the parent *PeakDetect node*.
- Disconnect from the flowchart (see below).
- Edit your peaks/bases
- Click "Send all curves to flowchart" (see below) to set the edited data as the output of the parent *PeakDetect node*.

#### Layout

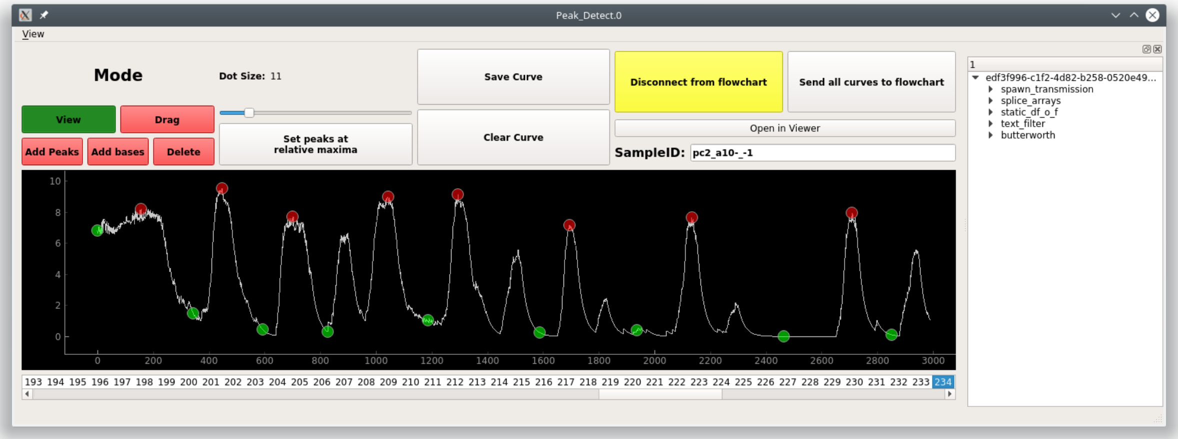


#### Bottom

List of curves from the Transmission inputted to the *Curve* or *PB_Input* terminal. See *PeakDetect node*

#### Top

**Mode buttons:** Set the current interactive mode for mouse events.

*View*: Just view, pan, and zoom the plot.

*Drag*: Click and drag peaks/bases along the curve.

*Add Peak/Base*: Click to add a peak/base onto the curve.

*Delete*: Delete a peak or base.

**Dot Size:** Move the slider to change the size of the dots representing peaks/bases.

**Set Peaks at relative maxima:** Not implemented yet.

**Save Curve:** Save the current curve. A curve auto-saved when you switch to another one.

**Clear Curve:** Not implemented.

**Disconnect from flowchart:** Disconnect the GUI from changes in the flowchart. Edits to the peaks/bases will be lost if this GUI is not disconnected while changes occur in the flowchart.

**Send all curves to flowchart:** Set the edited data as the output of the parent *PeakDetect node*

**Open in viewer:** Open the parent Sample of this curve in a *Viewer*.

##### Right

History Tree Widget

#### 1.33 Proportions

*API Reference*

**Note:** *This plot can be saved in an interactive form*

Compare proportions of categorical variables between different groups using bar charts.

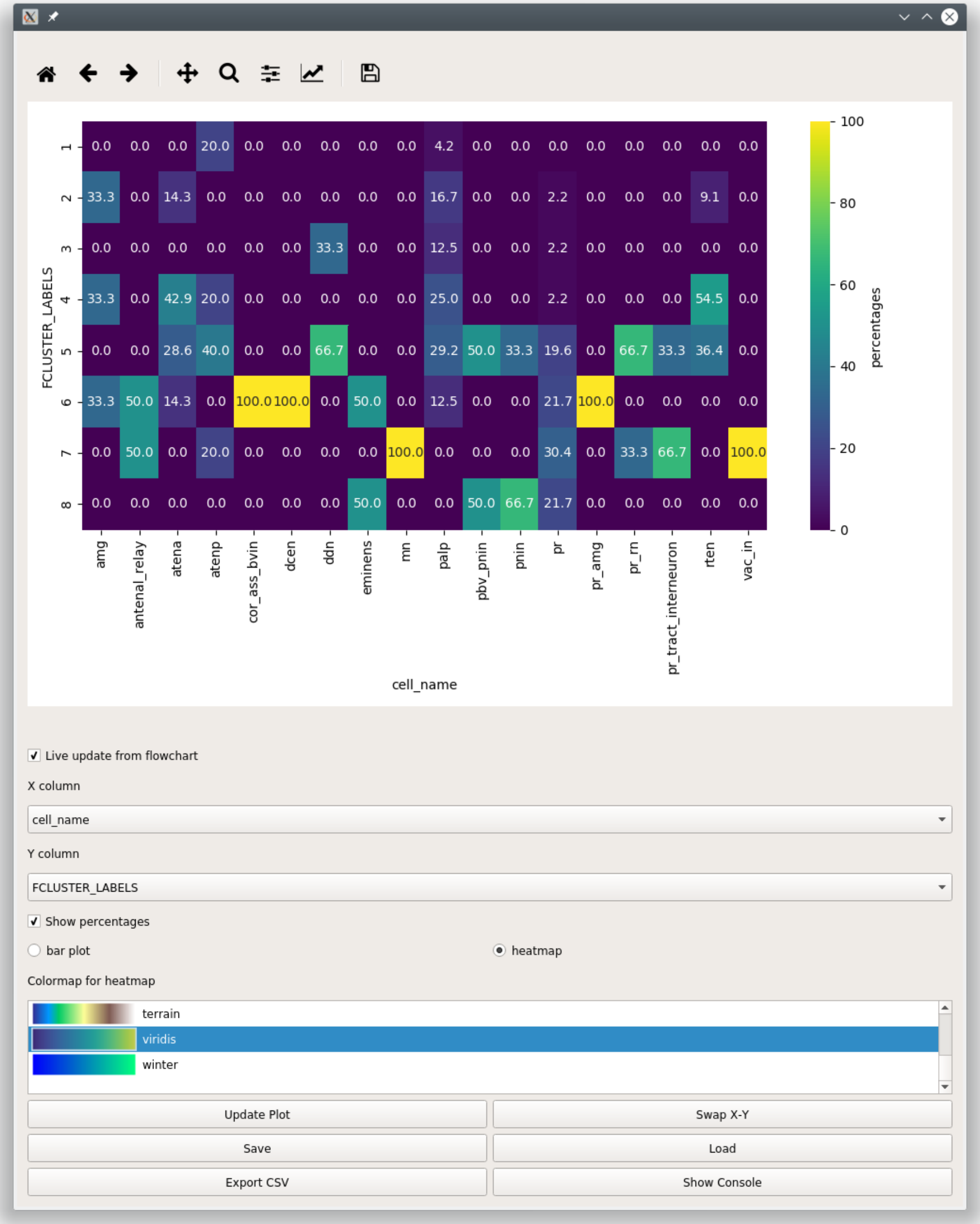
100
80
60
40
20
0
percentages
FCLUSTER_LABELS
1 0.0 0.0 0.0 20.0 0.0 0.0 0.0 0.0 0.0 4.2 0.0 0.0 0.0 0.0 0.0 0.0 0.0 0.0
2 33.3 0.0 14.3 0.0 0.0 0.0 0.0 0.0 0.0 16.7 0.0 0.0 2.2 0.0 0.0 0.0 9.1 0.0
3 0.0 0.0 0.0 0.0 0.0 0.0 33.3 0.0 0.0 12.5 0.0 0.0 2.2 0.0 0.0 0.0 0.0 0.0
4 33.3 0.0 42.9 20.0 0.0 0.0 0.0 0.0 0.0 25.0 0.0 0.0 2.2 0.0 0.0 0.0 54.5 0.0
5 0.0 0.0 28.6 40.0 0.0 0.0 66.7 0.0 0.0 29.2 50.0 33.3 19.6 0.0 66.7 33.3 36.4 0.0
6 33.3 50.0 14.3 0.0 100.0 100.0 0.0 50.0 0.0 12.5 0.0 0.0 21.7 100.0 0.0 0.0 0.0 0.0
7 0.0 50.0 0.0 20.0 0.0 0.0 0.0 0.0 100.0 0.0 0.0 0.0 30.4 0.0 33.3 66.7 0.0 100.0
8 0.0 0.0 0.0 0.0 0.0 0.0 0.0 50.0 0.0 0.0 50.0 66.7 21.7 0.0 0.0 0.0 0.0 0.0
amg
antenal_relay
atena
atenp
cor_ass_bvin
dcen
ddn
eminens
mn
palp
pbv_pnin
pnin
pr
pr_amg
pr_rn
pr_tract_interneuron
rten
vac_in
cell_name
Live update from flowchart
X column
cell_name
Y column
FCLUSTER_LABELS
Show percentages
bar plot
heatmap
Colormap for heatmap
terrain
viridis
winter
Update Plot
Swap X-Y
Save
Load
Export CSV
Show Console

| Parameter | Description |
|---|---|
| X column | DataFrame column containing the categorical labels used for grouping the data<br>Data in each X column sums to 100% if *Show percentages* is checked |
| Y column | DataFrame column containing the categorical labels that are counted for each group |
| Show percentages | When unchecked shows raw counts |
| bar plot | Visualize as bar plots |
| heatmap | Visualize as a heatmap |
| Update Plot | Update plot |
| Swap X-Y | Swap X & Y columns |
| Save | *Save this plot as a ptrn file* |
| Load | *Load from a ptrn file* |
| Export CSV | Export the data for the current plot as to a csv file. |
| Show Console | Show/hide the console |

#### 1.34 Scatter

*API Reference*

**Interactive scatter plot**

**Note:** *This plot can be saved in an interactive form*

##### 1.34.1 Video Tutorial

From 13:04 onward this tutorial shows how you can perform PCA and visualize the transformed data using the Scatter Plot.

##### 1.34.2 Layout

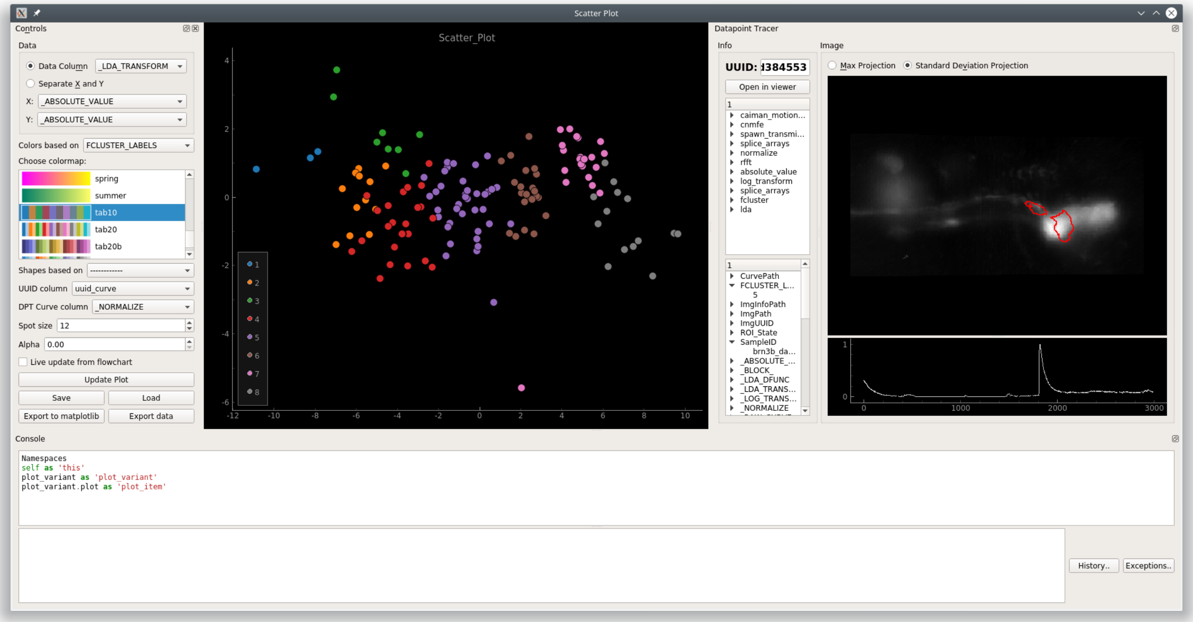


**Left:** Controls

| Control | |
|---|---|
| Data Column | Data column containing numerical arrays of size 2, X & Y values [x, y] |
| X | Data column containing only X values |
| Y | Data column containing only Y values |
| log x | Use $log_{10}$ of the X data |
| log y | Use $log_{10}$ of the Y data |
| Colors based on | Set spot colors based on categorical labels in this column |
| Choose colormap | Colromap for the the spot colors |
| Shapes based on | Set spot shapes based on categorical labels in this column |
| UUID Column | Column containing UUIDs that correspond to the plot data |
| DPT Curve column | Data column containing numerical arrays to show in the *Datapoint Tracer* |
| Spot size | Size of the spots |
| Alpha | Not implemented yet |
| Live update… | Update the plot with live inputs from the flowchart |
| Update Plot | Update the plot according to the input data from the flowchart and the parameters |
| Save | *Save the plot as a ptrn file* |
| Load | *Load a saved ptrn file* |
| Export to ma… | Not implemented yet |
| Export data | Not implemented yet |

**Below the plot:** Status label that displays plotting issues. Click the label to see more information.

**Right:** *Datapoint Tracer*. Click datapoints in the plot to set the Datapoint Tracer.

**Bottom:** *Console*

#### 1.35 Simple Plot

Just a very basic time-series plot. It will plot all the data in the selected data column.

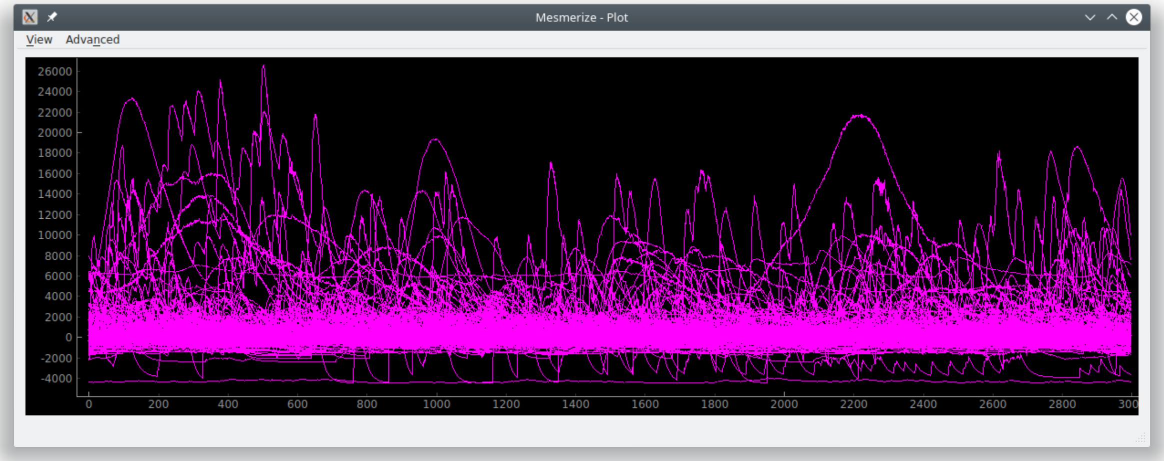


#### 1.36 SpaceMap

*API Reference*

**Note:** This plot can be saved in an interactive form, see *Saving plots*

Spatially map a categorical variable onto a projection of a Sample's image sequence

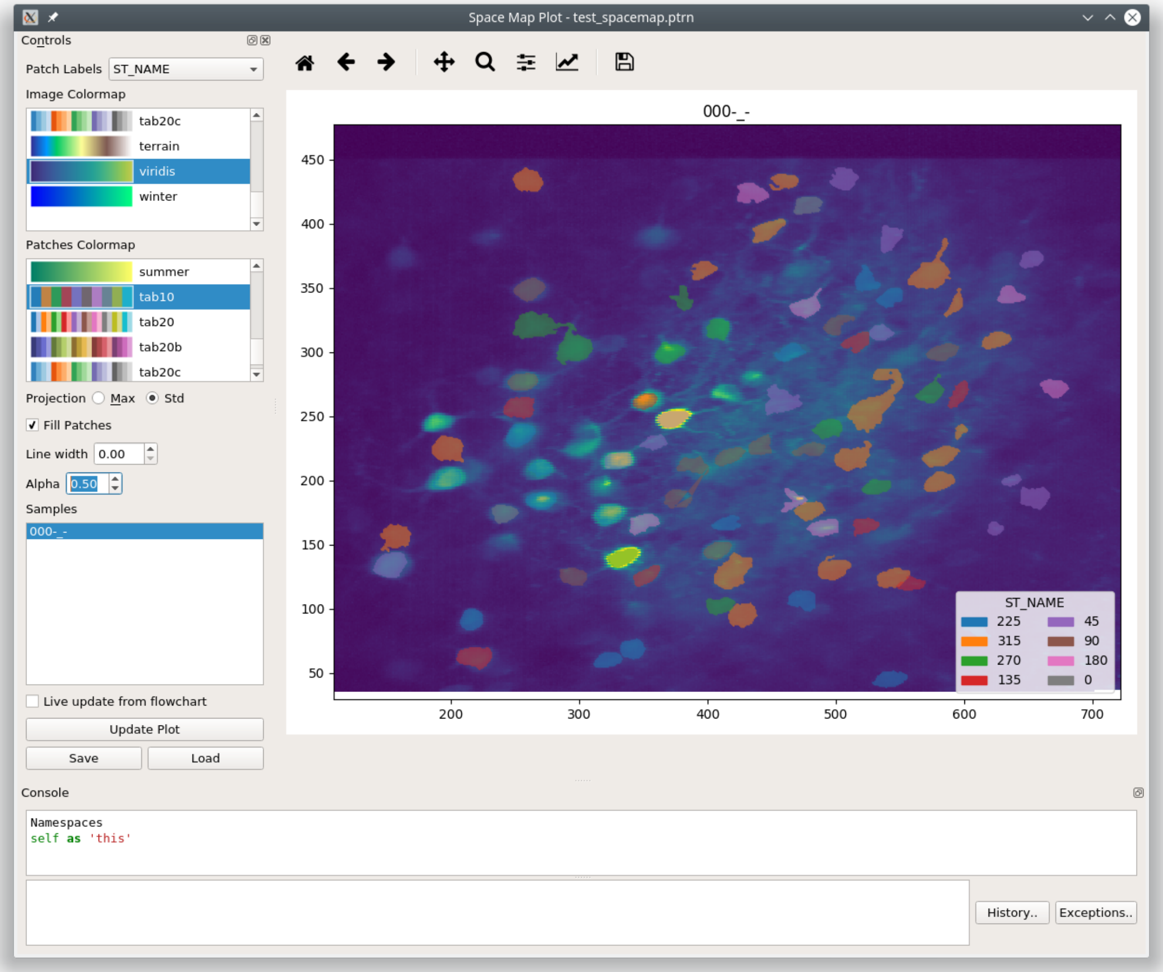


**Note:** Image produced from the following dataset: Garner, Aleena (2014): In vivo calcium imaging of layer 4 cells in the mouse using sinusoidal grating stimuli. CRCNS.org. http://dx.doi.org/10.6080/K0C8276G

#### 1.36.1 Video Tutorial

This shows how you can view a space map that shows the tuning of cells. The Space map plot itself is shows after 3:38.

##### 1.36.2 Controls

| Parameter | Description |
|---|---|
| Patch labels | Categorical column to use for the patch labels |
| Image Colormap | Colormap for the image |
| Patches Colormap | Colormap for the patches |
| Projection | Show the image as a "Max" or "Standard Deviation" projection |
| Fill Patches | Fill the patches |
| Line width | Line width of the patches |
| Alpha | Alpha level of the patches |
| Samples | Click on the sample to plot |
| Save | *Save the plot data and state in an interactive form* |
| Load | Load a plot that has been saved as a ".ptrn" file. |

##### 1.36.3 Console

**See also:**

*API Reference*

###### Namespace

| reference | Description |
|---|---|
| this | The *SpaceMapWidget* instance, i.e. the entire widget |
| this.transmission | Current input *Transmission* |
| get_plot() | Returns the plot area |
| get_plot().fig | Returns the matplotlib Figure instance |
| get_plot().ax | Returns the Axes for the current plot matplotlib Axes |

###### Examples

###### Export

**See also:**

matplotlib API for: Figure.savefig, Figure.set_size_inches, Figure.get_size_inches

```
# Desired size (width, height)
size = (6,5)

# Get the figure
fig = get_plot().fig

# original size to reset the figure after we save it
orig_size = fig.get_size_inches()

#Set the desired size
fig.set_size_inches(size)

```

(continues on next page)

(continued from previous page)

```
# Save the figure as a png file with 600 dpi
fig.savefig('/share/data/temp/kushal/spacemap.png', dpi=600, bbox_inches='tight', pad_
↪inches=0)

# Reset to original size and draw
fig.set_size_inches(orig_size)
get_plot().draw()
```

**Note:** The entire plot area might go gray after the figure is reset to the original size. I think this is a Qt-matplotlib issue. Just resize the window a bit and the plot will be visible again!

###### Legend Title

**See also:**

matplotlib API for matplotlib.axes.Axes.get_legend

```
get_plot().ax.get_legend().set_title('New Title')
get_plot().draw()
```

###### Hide Axis Borders

**See also:**

matplotlib API for matplotlib.axes.Axes.axis

```
get_plot().ax.axis('off')
get_plot().draw()
```

#### 1.37 Stimulus Tuning

Get the stimulus tuning and tuning curves of neurons.

The output from this plot node can also be used for various things downstream, such as hierarchical clustering to sort your cells based on their tuning, visualizing the tuning of each neuron using a *SpaceMap plot*, and much more. See the video tutorial for examples.

##### 1.37.1 Video Tutorial

This tutorial uses part of the CRCNS pvc-7 dataset from the Allen Institute to get stimulus tuning curves, perform hierarchical clustering and dimensionality reduction.

##### 1.37.2 Layout

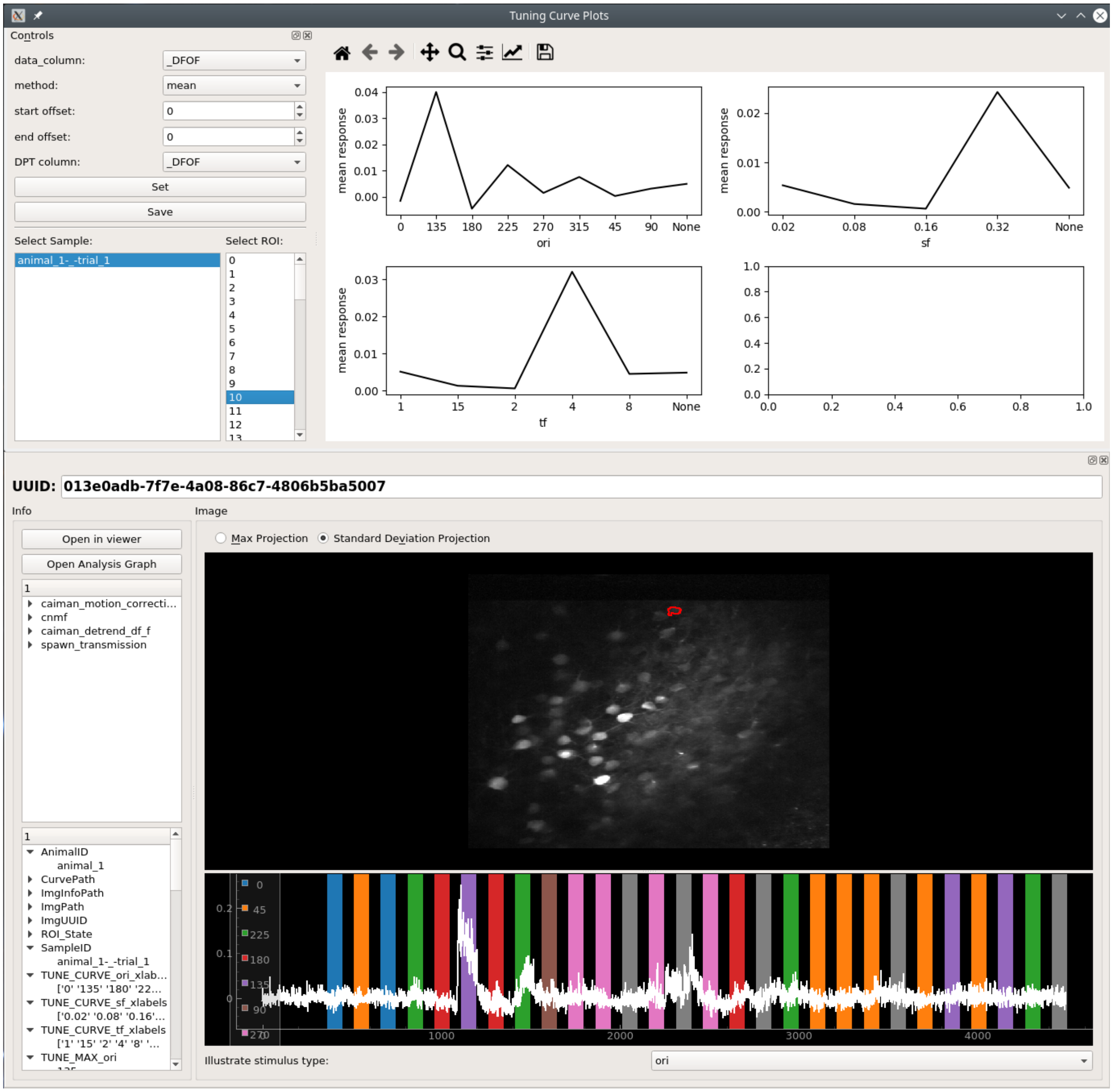


##### 1.37.3 Controls

| Parameter | Description |
|---|---|
| data_column | Data column used to determine the stimulus tuning of the cells |
| method | Use one of `mean`, `median`, `max` or `min` response within a stimulus period to determine the tuning |
| start offset | Use a start offset for the stimulus periods (can be either positive or negative) |
| end offset | Use a end offset for the stimulus periods (can be either positive or negative) |
| DPT column | Data column that is shown in the *Datapoint Tracer*. |
| Set | Set the stimulus extraction parameters defined above. |
| Save | *Save the plot data and state in an interactive form* |

##### 1.37.4 Usage

1. Set the desired parameters for **data_column**, **method**, **start offset**, **end offset** and **DPT column**.
2. Click **Set**.
3. Choose a Sample from the list.
4. Click on an ROI number to view the tuning curve, and corresponding spatial localization and curve in the *Datapoint Tracer*.
5. You can use the output of this plot node for further analysis, as show in the tutorial video.

#### 1.38 Welcome Window

The Welcome Window is the first window that you are presented with when you launch Mesmerize.

- Use the large buttons for opening new *Viewer* or *Flowchart* windows.
- Open a project using the button, or double-click a recent project from the list.
- Create a new project using the button.
- You basically have access to all objects in the Mesmerize instance through this console.

**See also:**

*User guide on creating new projects* and *Consoles*

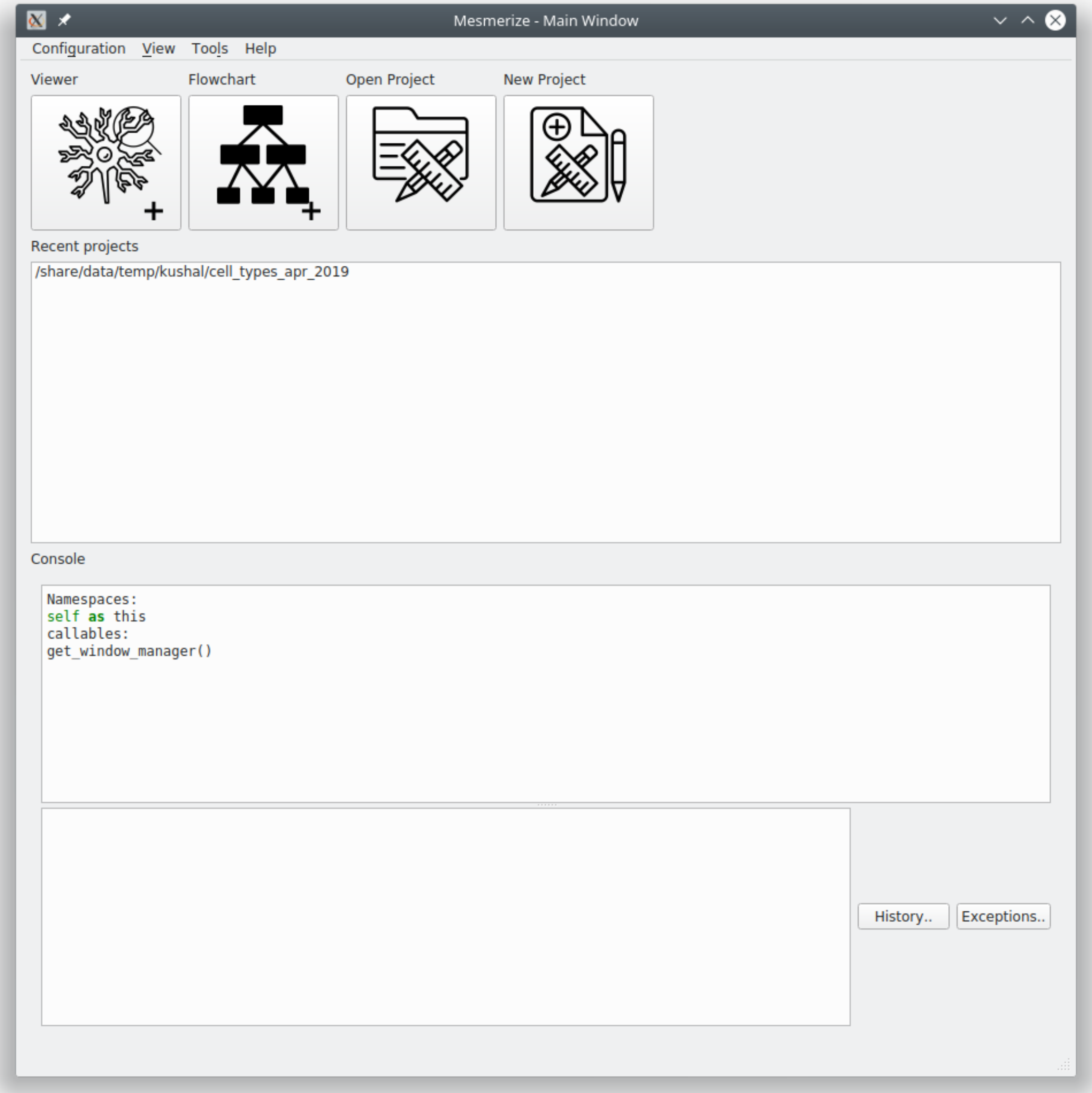


#### 1.39 Project Structure

A Mesmerize project is encapsulated within a single directory. It contains the following:

- config file - contains configuration data, such as roi type columns, stimulus type columns, and custom columns with their datatypes.

> **Warning:** Do not manually modify the config file

**Directories**

| Dir | Purpose |
|---|---|
| dataframes | Contains an file storing the project dataframe, root.dfr, and backups.<br>A new backup is created every time a a new *Sample* is *added to the project*.<br>Restore a backup by renaming it to "root.dfr". |
| images | Contains the image sequences and work environment data for all samples in the project |
| batches | Used for storing batches used by the *Batch Manager* if you wish. |
| flowcharts | Used for storing *.fc* flowchart files that save the layout of nodes in a flowchart. |
| plots | Used for storing *.ptrn* interactive plot files. |

**See also:**

*Flowchart Overview* and *Saving Plots*

**Warning:** Do not manually modify the data under the **images** or **curves** directories

#### 1.40 Consoles

A Python console is embedded in many parts of Mesmerize. You can use it to perform very specific operations, further automate tasks, save an analysis object, format plots, etc.

The console is accessible in many windows through View -> Console. Within the console namespace `this` refers to the window. For example `this` refers to the *Project Browser* Window instance in the Project Browser's console. A list of useful object references and helper functions are listed when you open most consoles.

You can run entire scripts within the console. You can also use import statements to import libraries that you have in your Python environment.

Keyboard controls:

**Execute:** Shift + Enter

**New line:** Enter

**Scroll up through history**: Page Up

**Scroll down through history:** Page Down

The history is stored in `~/.mesmerize`

#### 1.41 Saving plots

Some plots allow you to save them in an interactive form, along with the plot data and the plot state as a ".ptrn" file. If you save the file in the "plots" directory of your project it will be listed in the *Welcome Window* when you open your project.

This is currently possible with the following plots: *Heatmap*, *KShape*, *Proportions*, *Scatter*, and *SpaceMap*

#### 1.42 Plot Navbar

Many plots have a navigation toolbar which you can use to zoom, pan, configure plots, and export plots as images.

Official matplotlib docs about the navigation toolbar: https://matplotlib.org/2.1.2/users/navigation_toolbar.html

**Home:** Reset the plot (not applicable for all plots)

**Pan:** Pan the plot

**Zoom:** Zoom in/out a selection using the left/right mouse button respectively

**Subplot-configuration:** Options to adjust spacing, borders, set tight layout.

**Edit axis, curve…:** For some plots. Options for formating x & y axis limits, labels, select line style, color, etc.

**Save:** Export the figure as an image. **This is not the same as saving an interactive plot, see "Saving Plots" above**.

#### 1.43 System Configuration

**Set system configuration options**

This window is accessible through the *Welcome Window* menubar at Configuration -> System Configuration.

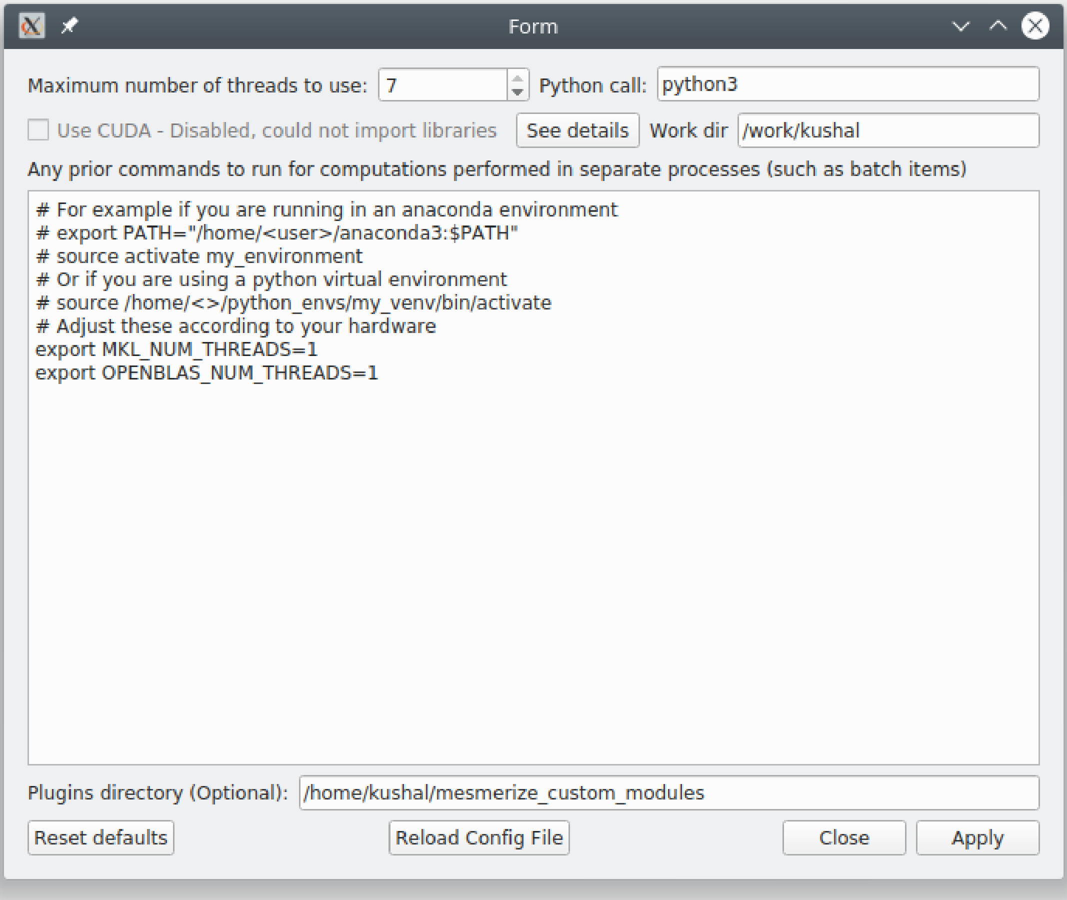

**Maximum number of threads to use:** The maximum number of threads that Mesmerize is allowed to use, this includes proccesses started by the *Batch Manager*, various analysis proccesses in the flowchart, and the viewer as well.

**Python call:** Many parts of Mesmerize, such as the *Batch Manager* use external processes to run a python script. This setting sets which python call should be used. `python3` should work for Linux & Mac OSX. We've found that this needs to be set to `python` to work within Anaconda environments on Windows.

**Use CUDA:** Use CUDA accerlation if you have a GPU with CUDA cores. You must have pycuda and scikit-cuda (as well as the nvidia CUDA toolkit) installed. CUDA accerlation isn't used much currently.

**Work dir:** Many parts of Mesmerize use a working directory for temporary files. If you have a fast filesystem you can use that for this purpose.

**Pre-run commands (large text entry):** Mesmerize runs some computationally intensive tasks in subprocesses. These commands are run prior to the python script that performs the task.

- If you are using Mesmerize in a virtual environment or conda environment you will need activate the environment so you must include the line `source /path_to_venv/bin/activate` or `conda activate <env_name>` to the pre-run commands.
- If you are using an Intel CPU you should get optimal performance by installing Math Kernel Library (MKL) and including `export MKL_NUM_THREADS=1` to the pre-run commands.
- If you are using an AMD CPU make sure you have OpenBLAS installed for optimal performance and include `export OPENBLAS_NUM_THREADS=1` to the pre-run commands. You may better performance by installing the AMD specific libraries.

**Plugins directory:** If you have a plugins dir include enter its path here.

#### 1.44 Nodes

The easiest way to create a new node is create a class that inherits from *`CtrlNode`*. You can place this class in one of the existing modules in mesmerize/pyqtgraphCore/flowchart/library

Become familiar with the *Transmission object* before creating a node. Almost all nodes work with a Transmission object for storing data. Make sure to conform to the conventions for naming of data columns and categorical columns.

##### 1.44.1 Simple node

The simplest type of node performs an operation on a user-specified data column and doesn't take any parameters.

**Basic structure**

```
class MyNode(CtrlNode):
"""Doc String that is shown when node is clicked on"""
nodeName = 'MyNode'
uiTemplate = <list of tuples, see below>

def processData(self, transmission: Transmission):
    self.t = transmission.copy()  #: input to this node

    # .. do stuff to the Transmission DataFrame

    params = <dict of analysis params>

    # log the analysis that was done
```

(continues on next page)

(continued from previous page)

```
    self.t.history_trace.add_operation('all', 'mynode', params)

    # Send the output
    return self.t
```

Required class attributes:

- **nodeName: (str)** The name prefix used when instances of this node are created in the flowchart
- **uiTemplate: (list)** *List of UI element tuples (see section)*

If the node only has one input and one output terminal it is sufficient to create a processData method that performs the node's analysis operation(s).

##### Example

```
class Derivative(CtrlNode):
"""Return the Derivative of a curve."""
nodeName = 'Derivative'
uiTemplate = [('data_column', 'combo', {}),
              ('Apply', 'check', {'checked': False, 'applyBox': True})
              ]

# If there is only one input and one output temrinal, processData will
# always have a single argument which is just the input transmission,
# i.e. the output from the previous node.
def processData(self, transmission: Transmission):
    # the input transmission
    self.t = transmission

    # If a comboBox widget named 'data_column' is specified in the
    # uiTemplate, you can update its contents using the following method.
    # This will populate the comboBox with all the data columns from the
    # input transmission and select the input data column as the
    # output data column from the previous node.
    self.set_data_column_combo_box()

    # Check if the Apply checkbox is checked
    if self.ctrls['Apply'].isChecked() is False:
        return

    # Make a copy of the input transmission so we can modify it to create an output
    self.t = transmission.copy()

    # By convention output columns are named after the node's name and in all caps
    # Columns containing numerical data have a leading underscore
    output_column = '_DERIVATIVE'

    # Perform this nodees operation
    self.t.df[output_column] = self.t.df[self.data_column].apply(np.gradient)

    # Set tranmission's `last_output` attribute as the name of the output column
```

(continues on next page)

(continued from previous page)

```
    # This is used by the next node to know what thet last output data was
    self.t.last_output = output_column

    # Create a dict of parameters that this node used
    # Usually a dict that captures the state of the uiTemplate
    # the transmission `last_unit` attribute is the data units of the data
    # in the output column (i.e. `t.last_output`). Change it only if the data units change
    params = {'data_column': self.data_column,
              'units': self.t.last_unit
              }

    # Add a log of this node's operation to the transmission's `HistoryTrace` instance
    # Nodes usually perform an operation on all datablocks pass 'all' to the data_block_id argument
    # By convention the operation name is the name of the node in lowercase letters
    self.t.history_trace.add_operation(data_block_id='all', operation='derivative', parameters=params)

    # return the modified transmission instance, which is then the output of this node
    return self.t
```

##### 1.44.2 Complex node

For a more complex node with multiple inputs and/or outputs you will need to explicitly specify the terminals when instantiating the parent *CtrlNode* and create a simple override of the process() method.

Format of the dict specifying the node's terminals:

```
{
    <terminal name (str)>:            {'io': <'in' or 'out'>},
    <another terminal name (str)>:    {'io', <'in' or 'out'>},
    <another terminal name (str)>:    {'io', <'in' or 'out'>}
    ...
}
```

Override the process() method simply pass all kwargs to a processData() method and return the output The processData() method must return a dict. This dict must have keys that correspond to the specified output terminals. The values of these keys are the outputs from the respective terminals.

Here is a trimmed down example from the *LDA node*:

```
class LDA(CtrlNode):
"""Linear Discriminant Analysis, uses sklearn"""
nodeName = "LDA"
uiTemplate = [('train_data', 'list_widget', {'selection_mode': QtWidgets.QAbstractItemView.ExtendedSelection,
                                             'toolTip': 'Column containing the training data'}),
              ('train_labels', 'combo', {'toolTip': 'Column containing training labels'}),
              ('solver', 'combo', {'items': ['svd', 'lsqr', 'eigen']}),
              ('shrinkage', 'combo', {'items': ['None', 'auto', 'value']}),
```

(continues on next page)

(continued from previous page)

```
              ('shrinkage_val', 'doubleSpin', {'min': 0.0, 'max': 1.0, 'step': 0.1, 'value': 0.5}),
              ('n_components', 'intSpin', {'min': 2, 'max': 1000, 'step': 1, 'value': 2}),
              ('tol', 'intSpin', {'min': -50, 'max': 0, 'step': 1, 'value': -4}),
              ('score', 'lineEdit', {}),
              ('predict_on', 'list_widget', {'selection_mode': QtWidgets.QAbstractItemView.ExtendedSelection,
                                             'toolTip': 'Data column of the input "predict" Transmission\n'
                                                        'that is used for predicting from the model'}),
              ('Apply', 'check', {'applyBox': True, 'checked': False})
              ]

def __init__(self, name, **kwargs):
    # Specify the terminals with a dict
    CtrlNode.__init__(self, name, terminals={'train': {'io': 'in'},
                                             'predict': {'io': 'in'},

                                             'T': {'io': 'out'},
                                             'coef': {'io': 'out'},
                                             'means': {'io': 'out'},
                                             'predicted': {'io': 'out'}
                                             },
                      **kwargs)
    self.ctrls['score'].setReadOnly(True)

# Very simple override
def process(self, **kwargs):
    return self.processData(**kwargs)

def processData(self, train: Transmission, predict: Transmission):
    self.t = train.copy()  #: Transmisison instance containing the training data with the labels
    self.to_predict = predict.copy()  #: Transmission instance containing the data to predict after fitting on the the training data

    # function from mesmerize.analysis.utils
    dcols, ccols, ucols = organize_dataframe_columns(self.t.df.columns)

    # Set available options for training data & labels
    self.ctrls['train_data'].setItems(dcols)
    self.ctrls['train_labels'].setItems(ccols)

    dcols = organize_dataframe_columns(self.to_predct.df.columns)
    # Set available data column options for predicting on
    self.ctrls['predict_on'].setItems(dcols)

    # Process further only if Apply is checked
    if not self.ctrls['Apply'].isChecked():
        return
```

(continues on next page)

(continued from previous page)

```

    # Get the user-set parameters
    train_column = self.ctrls['train_data'].currentText()

    # ... get other params
    n_components = self.ctrls['n_components'].value()

    # ... do stuff

    # This node ouputs separate transmissions that are all logged
    self.t.history_trace.add_operation('all', 'lda', params)
    self.t_coef.history_trace.add_operation('all', 'lda', params)
    self.t_means.history_trace.add_operation('all', 'lda', params)

    # the `to_predict` transmission is logged differently
    self.to_predict.history_trace.add_operations('all', 'lda-predict', params_predict)

    # dict for organizing this node's outputs
    # The keys MUST be the same those specified for this node's output terminals
    out = {'T': self.t,
           'coef': self.t_coef,
           'means': self.t_means,
           'predicated': self.to_predct
           }

    return out
```

##### 1.44.3 uiTemplate

Specify the uiTemplate attribute as a list of tuples.

One tuple per UI element with the following structure:

(<name (str)>, <type (str)>, <dict of attributes to set>)

Examples:

```
('dist_metric', 'combo', {'items': ['euclidean', 'wasserstein', 'bah'], 'toolTip':
↪'distance metric to use'})
('n_components', 'intSpin', {'min': 2, 'max': 10, 'value': 2, 'step': 1, 'toolTip':
↪'number of components'})
('data_columns', 'list_widget', {'selection_mode': QtWidgets.QAbstractItemView.
↪ExtendedSelection})
```

The name can be anything. Accepted types and accepted attributes are outlined below

| widget type | attributes that can be set |
|---|---|
| intSpin | *min (int):* minimum value allowed in the spinbox<br>*max (int):* maximum value allowed<br>*step (int):* step size<br>*value (int):* default value |
| doubleSpin | *min (float):* minimum value allowed in the spinbox<br>*max (float):* maximum value allowed<br>*step (float):* step size<br>*value (float):* default value |
| check | *checked (bool):* default state of the checkBox<br>*applyBox (bool):* Whether this is an "Apply checkbox" |
| radioBtn | *checked (bool):* default state of this radioButton |
| combo | *items (list):* default list of items that will be set in the comboBox |
| list_widget | *items (list):* default list of items that will be set in the list_widget<br>*selection_mode:* One of the accepted QAbstractItemView selection modes |
| lineEdit | *text (str):* default text in the line edit<br>*placeHolder (str):* placeholder text<br>*readOnly (bool):* set as read only |
| plainTextEdit | *text (str):* default text in the text edit<br>*placeHolder (str):* placeholder text |
| label | *text (str):* default text |
| button | *text (str):* default text on the button<br>*checkable (bool):* whether this button is checkable |
| color | *Does not take any attributes* |

**All UI widget types outlined above take 'toolTip' as an attribute which can be used to display tooltips**

#### 1.45 Plots

The easiest way to create a plot module is by subclassing the *BasePlotWidget*. You could also subclass the abstract base if you need to define all the common functionality differently.

##### 1.45.1 General Design

This shows how you can design a plot using the *SpaceMapPlot* as a simple example. It will generally consist of a class for the main plot area, plot control, and the plot window which contains the controls and plot area.

##### 1.45.2 Plot Area

A class which holds the actual plot, could be a matplotlib widget or pyqtgraph plot widget for example. In the *SpaceMapPlot* this is simply a subclass of the pyqtgraph matplotlib widget with a few more attributes and a helper method. The *error_label* attribute is simply a QLabel used for displaying a plot error summary and is handled by the *exceptions_label* decorator from *qdialogs*.

##### 1.45.3 Plot Controls

A class which manages the plot controls. Generally useful to use a QDockWidget for this and design the actual GUI layout using QtDesigner. The *WidgetRegistry* provides a simple way to package the plot control values (plot parameters) into a dict.

Register a widget to the registry using the *WidgetRegistry* instance's *register()* method. The **getter** method corresponds to the widget's method which will return the value of the widget (such as text or a number) that is set in the parameters dict which is created when *widget_registry.get_state()* is called. Correspondingly, **setter** method is the widget's method that is used to set a value to the widget and is used when saved plots are restored. In essense, **setter** and **getter** must be interoperable.

The Space Map plot uses a *sig_changed* class attribute that simply emits when any of the widgets are changed. This is later used in the main plot window to update the plot.

A *fill_widget()* method is useful for populating the controls in the dock widget when the input data to the plot window changes.

In the Space Map widget, *get_state()* and *set_state()* simply wrap the corresponding methods from the *WidgetRegistry* instance.

##### 1.45.4 Plot Window

Subclass from QMainWindow and *BasePlotWidget*. **Mandatory** to specify a *drop_opts* class attribute of type *list*. This list contains the name of any widgets in the dict return from the *WidgetRegistry* that should be exluded when saving the plot. This should be used if you are using data types that are not JSON serializable, however it is rarely necessary. Support for *drop_opts* may be removed in the future.

In general specifying the methods described below should be sufficient to create a saveable plot. If you need finer control of the data struture for saving/opening plots you can subclass from the *abstract base class*.

##### __init__

Setting things up, connection signals, etc. Useful to have a console dock widget.

##### set_update_live()

A method that interacts with a "live update" checkbox in the plot controls.

##### set_input()

Set the input transmission for this plot if it is in "live update" mode or if the plot instance is new (has not had input data previously).

Useful to have a *BasePlotWidget.signal_blocker* decorator so that the plot doesn't constantly update while the new data comes in, since it could cause plot options to change etc.

##### fill_control_widget()

Organize the plot options that are available to the user and set the control widgets.

Useful to have a *BasePlotWidget.signal_blocker* decorator here as well for same reasons as described above.

##### update_plot()

This is the core of plot. Use the input transmission and the user-selected plot parameters to draw the plot in the plot area. Generally interacts with the Plot Area instance. You can use the *get_state()* method of the control widget's *WidgetRegistry* to conveniently get a dict of all the user-selected plot parameters.

Useful to have an *exceptions_label* or *present_exceptions* decorator from the *qdialogs module*. The *exceptions_label* provides a less annoying way to present exceptions that occured when updating the plot.

##### get_plot_opts()

Usually just returns the dict from the widget registry containing all user-set plot parameters.

##### set_plot_opts()

Usually just calls the widget registry's set_state() method to set the plot parameters from a dict.

Useful to have a *BasePlotWidget.signal_blocker* decorator. In general you would use the *BasePlotWidget*.open_plot() method to open a saved plot and it takes care of updating the plot after the input transmission and plot parameters are set.

###### show_exception_info()

Called when the *exceptions_label* is clicked. Opens a QMessageBox to show the entire stack trace.

#### 1.46 Viewer Modules

Viewer modules appear as either QDockWidgets or QWidgets to the user. They must consist of a main `ModuleGUI` class which inherits from either QDockWidget or QWidget. They can utilize any additional python modules, classes, etc.

##### 1.46.1 Instructions

1. Create a directory plugins directory if you don't have one. If you are using a snap installation this has to be within your home folder. Set this plugins directory in the *System Configuration*. This directory can contain as many custom modules as you want. All python modules within the plugins directory are automatically imported.
2. Download the __init__.py and place it within the plugins directory.
3. Create the main module file for your custom module. This file can be named as you wish and must use the struture outlined below. In addition to this main module file you can create a directory to house any other modules, files etc. You can create Qt templates using Qt Creator and convert them to .py template files using pyuic5 and use them for your custom module.

   **Basic Structure**

```python
from PyQt5 import QtWidgets

module_name = 'Example Module'

# You must define module_name.
# This is the name that will be displayed in the "Plugins" menu of the Viewer Window.
# You can use this to reference the ModuleGUI instance through the Viewer Console via ``get_module(<module_name>)``


# The main GUI class MUST be named ModuleGUI.
# You can have other classes and more GUIs however ModuleGUI is the one that the Viewer Window directly calls.

class ModuleGUI(QtWidgets.QDockWidget):
    # The Viewer MainWindow will pass its Viewer instance that can be used to interact with the viewer and work environment.
    def __init__(self, parent, viewer_instance):
        QtWidgets.QDockWidget.__init__(self, parent)
        self.setFloating(True) # Must be floating
```

4. The module will be accessible through the Viewer Window's "Plugins" menu. The names in the plugins menu will correspond to the aforementioned `module_name` variable.

#### 1.47 Common

mesmerize.common.**get_proj_config**(*proj_path: Optional[str] = None*) → configparser.RawConfigParser

**Parameters** **proj_path** – Full project path

mesmerize.common.**get_project_manager**()
Get the project manager for this Mesmerize instance

mesmerize.common.**get_sys_config**() → dict
Get the user-set system configuration

mesmerize.common.**get_window_manager**()
Get the Window Manager for this Mesmerize instance

##### 1.47.1 Utils

**Some frequently used utility functions**

mesmerize.common.utils.**make_workdir**(*prefix: str = ''*) → str
Make a workdir within the mesmerize_tmp directory of the workdir specified in the configuration The name of the created workdir is the date & time of its creation. You can add a prefix to this name.

**Parameters** **prefix** – Prefix for the workdir namec

**Returns** full workdir path

**Return type** str

mesmerize.common.utils.**make_runfile**(*module_path: str, savedir: str, args_str: Optional[str] = None, filename: Optional[str] = None, pre_run: Optional[str] = None, post_run: Optional[str] = None*) → str
Make an executable bash script. Used for running python scripts in external processes.

**Parameters**

- **module_path** (*str*) – absolute module path
- **args_str** (*str*) – str of args that is directly passed with the python command in the bash script
- **savedir** (*Optional[str]*) – working directory
- **filename** (*Optional[str]*) – optional, specific filename for the script
- **pre_run** (*Optional[str]*) – optional, str to run before module is ran
- **post_run** (*Optional[str]*) – optional, str to run after module has run

**Returns** path to the shell script that can be run

**Return type** str

**class** mesmerize.common.utils.**HdfTools**
Functions for saving and loading HDF5 data

**static save_dataframe**(*path: str, dataframe: pandas.core.frame.DataFrame, metadata: Optional[dict] = None, metadata_method: str = 'json', raise_meta_fail: bool = True*)
Save DataFrame to hdf5 file along with a meta data dict.

Meta data dict can either be serialized with json and stored as a str in the hdf5 file, or recursively saved into hdf5 groups if the dict contains types that hdf5 can deal with. Experiment with both methods and see what works best

Currently the hdf5 method can work with these types: [str, bytes, int, float, np.int, np.int8, np.int16, np.int32, np.int64, np.float, np.float16, np.float32, np.float64, np.float128, np.complex].

If it encounters an object that is not of these types it will store whatever that object's __str__() method returns if on_meta_fail is False, else it will raise an exception.

**Parameters**

- **path** (*str*) – path to save the file to
- **dataframe** (*pd.DataFrame*) – DataFrame to save in the hdf5 file
- **metadata** (*Optional[dict]*) – Any associated meta data to store along with the DataFrame in the hdf5 file
- **metadata_method** (*str*) – method for storing the metadata dict, either 'json' or 'recursive'
- **raise_meta_fail** (*bool*) – raise an exception if recursive metadata saving encounters an unsupported object If false, it will save the unsupported object's __str__() return value

`static load_dataframe(filepath: str) → Tuple[pandas.core.frame.DataFrame, Optional[dict]]`

Load a DataFrame along with meta data that were saved using `HdfTools.save_dataframe`

**Parameters** **filepath** (*str*) – file path to the hdf5 file

**Returns** tuple, (DataFrame, meta data dict if present else None)

**Return type** Tuple[pd.DataFrame, Union[dict, None]]

`static save_dict(d: dict, filename: str, group: str, raise_type_fail=True)`

Recursively save a dict to an hdf5 group.

**Parameters**

- **d** (*dict*) – dict to save
- **filename** (*str*) – filename
- **group** (*str*) – group name to save the dict to
- **raise_type_fail** (*bool*) – whether to raise if saving a piece of data fails

`static load_dict(filename: str, group: str) → dict`

Recursively load a dict from an hdf5 group.

**Parameters**

- **filename** (*str*) – filename
- **group** (*str*) – group name of the dict

**Returns** dict recursively loaded from the hdf5 group

**Return type** dict

`mesmerize.common.utils.draw_graph(l: List[dict], filename: Optional[str] = None, view: bool = False) → str`

Draw a graph from a list of dicts.

**Parameters**

- **l** (*List[dict]*) – list of dicts
- **filename** (*Optional[str]*) – full path for storing the draw graph pdf file

- **view** (*Optional[bool]*) – view the graph in the system's default pdf reader after it is rendered

**Returns** full path to the graph pdf file

**Return type** str

#### 1.47.2 QDialogs

**Decorators for Qt Dialog GUIs used throughout Mesmerize**

`mesmerize.common.qdialogs.`**`present_exceptions`**(*title: str = 'error', msg: str = 'The following error occurred.'*)

Use to catch exceptions and present them to the user in a QMessageBox warning dialog. The traceback from the exception is also shown.

This decorator can be stacked on top of other decorators.

Example:

**Parameters**

- **title** – Title of the dialog box
- **msg** – Message to display above the traceback in the dialog box
- **help_func** – A helper function which is called if the user clicked the "Help" button

`mesmerize.common.qdialogs.`**`exceptions_label`**(*label: str, exception_holder: Optional[str] = None, title: str = 'error', msg: str = 'The following error occured'*)

Use a label to display an exception instead of a QMessageBox

**Parameters**

- **label** – name of a QLabel instance
- **exception_holder** – name of an exception_holder attribute where the exception message is stored. This can be used to view the whole exception when the label is clicked on for example.
- **title** – title supplied for the QMessageBox (if used later)
- **msg** – message supplied for the QMessageBox (if used later)

`mesmerize.common.qdialogs.`**`use_open_file_dialog`**(*title: str = 'Choose file', start_dir: Optional[str] = None, exts: Optional[List[str]] = None*)

Use to pass a file path, for opening, into the decorated function using QFileDialog.getOpenFileName

**Parameters**

- **title** – Title of the dialog box
- **start_dir** – Directory that is first shown in the dialog box.
- **exts** – List of file extensions to set the filter in the dialog box

`mesmerize.common.qdialogs.`**`use_save_file_dialog`**(*title: str = 'Save file', start_dir: Optional[str] = None, ext: Optional[str] = None*)

Use to pass a file path, for saving, into the decorated function using QFileDialog.getSaveFileName

**Parameters**

- **title** – Title of the dialog box
- **start_dir** – Directory that is first shown in the dialog box.

- **exts** – List of file extensions to set the filter in the dialog box

mesmerize.common.qdialogs.**use_open_dir_dialog**(*title: str = 'Open directory', start_dir: Optional[str] = None*)

Use to pass a dir path, to open, into the decorated function using QFileDialog.getExistingDirectory

**Parameters**

- **title** – Title of the dialog box
- **start_dir** – Directory that is first shown in the dialog box.

Example:

```
@use_open_dir_dialog('Select Project Directory', '')
def load_data(self, path, *args, **kwargs):
    my_func_to_do_stuff_and_load_data(path)
```

#### 1.48 Viewer Core

##### 1.48.1 Video Tutorial

##### 1.48.2 ViewerWorkEnv

This objects stores the data that the *Viewer* interacts with.

**class** mesmerize.viewer.core.**ViewerWorkEnv**(*imgdata: Optional[mesmerize.viewer.core.data_types.ImgData] = None, sample_id='', UUID=None, meta=None, stim_maps=None, roi_manager=None, roi_states=None, comments='', origin_file='', custom_cols=None, history_trace: Optional[list] = None, additional_data: Optional[dict] = None, misc: Optional[dict] = None, **kwargs*)

**_UUID**

UUID, if opened from a project Sample refers to the ImgUUID

**__init__**(*imgdata: Optional[mesmerize.viewer.core.data_types.ImgData] = None, sample_id='', UUID=None, meta=None, stim_maps=None, roi_manager=None, roi_states=None, comments='', origin_file='', custom_cols=None, history_trace: Optional[list] = None, additional_data: Optional[dict] = None, misc: Optional[dict] = None, **kwargs*)

A class that encapsulates the main work environment objects (img sequence, ROIs, and ROI associated curves) of the viewer. Allows for a work environment to be easily spawned from different types of sources and allows for a work environment to be easily saved in different ways regardless of the type of original data source.

**Parameters**

- **roi_states** (*dict*) – roi states from ROI Manager module
- **stim_maps** (*dict*) – {'units': str, 'dataframe': pd.DataFrame}
- **history_trace** (*list*) – list of dicts containing a traceable history of what what done with the work environment, such as params used from modules to process the data

**`__weakref__`**
list of weak references to the object (if defined)

**`static _organize_meta`**(*meta: dict*, *origin: str*) → dict
Organize input meta data dict into a uniform structure :param meta: meta data dict, origin from a json file for example :param origin: name of the origin source of the meta data, such a program or microscope etc. :return: dict organized with keys that are used throughout Mesmerize.

**`clear`**()
Cleanup of the work environment

**`classmethod from_mesfile`**(*mesfile_object: mesmerize.viewer.core.mesfile.MES*, *img_reference: str*)
Return instance of work environment with MesmerizeCore.ImgData class object using seq returned from MES.load_img from MesmerizeCore.FileInput module and any stimulus map that the user may have specified.

**Parameters**

- **`mesfile_object`** – MES object, created from .mes file
- **`img_reference`** – image reference to load, see *`mesmerize.viewer.core.mesfile.MES.get_image_references()`*

**`classmethod from_pickle`**(*pickle_file_path: str*, *tiff_path: Optional[str] = None*)
Get pickled image data from a pickle file & image sequence from a npz or tiff. Used after motion correction & to view a sample from a project DataFrame. Create ImgData class object (See MesmerizeCore.DataTypes) and return instance of the work environment.

**Param** pickle_file_path: full path to the pickle containing image metadata, stim maps, and roi_states

**Param** tiff_path: str of the full path to a tiff file containing the image sequence

**`classmethod from_tiff`**(*path: str*, *method: str*, *meta_path: Optional[str] = None*, *axes_order: Optional[str] = None*, *meta_format: Optional[str] = None*)
Return instance of work environment with ImgData.seq set from the tiff file.

**Parameters**

- **`path`** – path to the tiff file
- **`method`** – one of 'imread', 'asarray', or 'asarray-multi'. Refers to usage of either tifffile.imread or tifffile.asarray. 'asarray-multi' will load multi-page tiff files.
- **`meta_path`** – path to a file containing meta data
- **`meta_format`** – meta data format, must correspond to the name of a function in viewer.core.organize_meta
- **`axes_order`** – Axes order as a 3 or 4 letter string for 2D or 3D data respectively. Axes order is assumed to be "txy" or "tzxy" if not specified.

**`history_trace`**
history log

**`imgdata:`** *`mesmerize.viewer.core.data_types.ImgData`*
ImgData instance

**`isEmpty`**
Return True if the work environment is empty

**static load_mesfile**(*path: str*) → *mesmerize.viewer.core.mesfile.MES*

Just passes the path of a .mes file to the constructor of class MES in MesmerizeCore.FileInput. Loads .mes file & constructs MES obj from which individual images & their respective metadata can be loaded to construct viewer work environments using the classmethod viewerWorkEnv.from_mesfile.

**Parameters** `path` – full path to a single .mes file.

**roi_manager**

reference to the back-end ROI Manager that is currently in use

**sample_id**

SampleID, if opened from a project Sample

**stim_maps**

Stimulus maps

**to_pandas**(*proj_path: str*, *modify_options: Optional[dict] = None*) → list

Used for saving the work environment as a project Sample.

**Parameters**

- `proj_path` – Root path of the current project
- `modify_options` –

**Returns** list of dicts that each correspond to a single curve that can be appended as rows to the project dataframe

**to_pickle**(*dir_path: str*, *filename: Optional[str] = None*, *save_img_seq=True*, *UUID=None*) → str

Package the current work Env ImgData class object (See MesmerizeCore.DataTypes) and any paramteres such as for motion correction and package them into a pickle & image seq array. Used for batch motion correction and for saving current sample to the project. Image sequence is saved as a tiff and other information about the image is saved in a pickle.

##### 1.48.3 ImgData

**class** mesmerize.viewer.core.data_types.**ImgData**(*seq: Optional[numpy.ndarray] = None*, *meta: Optional[dict] = None*)

Object that stores the image sequence and meta data from the imaging source

**__init__**(*seq: Optional[numpy.ndarray] = None*, *meta: Optional[dict] = None*)

**Parameters**

- `seq` – Image sequence as a numpy array, shape is [x, y, t] or [x, y, t, z]
- `meta` – Meta data dict from the imaging source.

##### 1.48.4 ViewerUtils

The *Viewer* is usually not interacted with directly from modules outside of the viewer (such as viewer modules. They instead use the ViewerUtils class which includes helper functions and a reference to the viewer.

**class** mesmerize.viewer.core.**ViewerUtils**(*viewer_reference: <module 'mesmerize.pyqtgraphCore.imageview.ImageView' from '/home/docs/checkouts/readthedocs.org/user_builds/mesmerize/envs/master/lib/pyt[illegible] packages/mesmerize-0.6.1-py3.7.egg/mesmerize/pyqtgraphCore/imageview/ImageView.py'>*)

Some utility functions for interfacing viewer.core.ViewerWorkEnv with the pyqtgraphCore.ImageView widget

**__init__**(*viewer_reference: <module 'mesmerize.pyqtgraphCore.imageview.ImageView' from '/home/docs/checkouts/readthedocs.org/user_builds/mesmerize/envs/master/lib/python3.7/site-packages/mesmerize-0.6.1-py3.7.egg/mesmerize/pyqtgraphCore/imageview/ImageView.py'>*)

**_clear_workEnv**(*clear_sample_id=False*)
Cleanup of the ViewerWorkEnv and ImageView widget

**discard_workEnv**(*clear_sample_id=False*)
Ask the user if they want to discard their work environment. If Yes, calls _clear_workEnv()

**set_statusbar**(*msg*)
Set the status bar message in the viewer window.

**Parameters** **msg** – text to display in the status bar

**update_workEnv**()
Update the ImageView widget with the ViewerWorkEnv

**viewer**
reference to the pyqtgraph ImageView widget instance (viewer)

**work_env**
ViewerWorkEnv instance

##### 1.48.5 Mesfile

**class** mesmerize.viewer.core.mesfile.**MES**(*filename: str*)
Handles of opening .mes files and organizing the images and meta data. The load_img() method returns a 3D array (dims are [time, cols, rows]) of the image sequence and its associated meta data.

Usage: Create a MES instance by passing the path of your mes file, example:

mesfile = MES('/path/to/mesfile/experiment_Feb_31.mes')

Call the get_image_references() method to get a list of references for images that can be loaded.

To load an image that is available in the instance, just pass one of the references from get_image_references() to the load_img method:

img_array, meta_dict = mesfile.load_img('IF0001_0001')

**__init__**(*filename: str*)

**Parameters** **filename** – full path of a single .mes file

**__weakref__**
list of weak references to the object (if defined)

**get_image_references**() → list
Get a list of all image references available in the instance

**load_img**(*img_reference: str) -> (<class 'numpy.ndarray'>, <class 'dict'>*)

**Parameters** **img_reference** – The image reference, usually something like IFxxxx_xxxx or Ifxxxx_xxxx

**Returns** (image sequence array, meta data dict)

##### 1.48.6 Examples

These examples can be run in the *Viewer Console*.

###### Working with meta data

```
# view meta data
>>> get_meta()

{'origin': 'AwesomeImager', 'version': '4107ff58a0c3d4d5d3c15c3d6a69f8798a20e3de', 'fps': 10.0, 'date': '20190426_152034', 'vmin': 323, 'vmax': 1529, 'orig_meta': {'source': 'AwesomeImager', 'version': '4107ff58a0c3d4d5d3c15c3d6a69f8798a20e3de', 'level_min': 323, 'stims': {}, 'time': '152034', 'date': '20190426', 'framerate': 10.0, 'level_max': 1529}}

# manually set meta data entries
>>> get_meta()['fps'] = 30.0
```

###### Open image

Use the *Viewer Core API* to open any arbitrary image

This example loads an image stored using numpy.save(), but this is applicable to images stored in any format that can eventually be represented as a numpy array in python. For example, you could also load image files stored in HDF5 format and load the numpy array that represents your image sequence.

```
import numpy as np

# clear the viewer work environment
clear_workEnv()

a = np.load('/path_to_image.npy')

# check what the axes order is
a.shape

# (1000, 512, 512) # for example
# looks like this is in [t, x, y]
# this can be transposed so we get [x, y, t]
# ImgData takes either [x, y, t] or [x, y, t, z] axes order

# Define a meta data dict
meta = \
    {
        "origin":      "Tutorial example",
        "fps":         10.0,
        "date":        "20200629_171823",
        "scanner_pos": [0, 1, 2, 3, 4, 5, 6]
    }

# Create ImgData instance
imgdata = ImgData(a.T, meta)  # use a.T to get [x, y, t]
```

(continues on next page)

(continued from previous page)

```

# Create a work environment instance
work_env = ViewerWorkEnv(imgdata)

# Set the current Viewer Work Environment from this new instance
vi.viewer.workEnv = work_env

# Update the viewer with the new work environment
# this MUST be run whenever you replace the viewer work environment (the previous line)
update_workEnv()
```

##### Image data

Image sequences are simply numpy arrays. For example extract the image sequence between frame 1000 and 2000.

**See also:**

Numpy array indexing

```
# Get the current image sequence
seq = get_image()

# Trim the image sequence
trim = seq[:, :, 1000:2000]

# Set the viewer work environment image sequence to the trim one
vi.viewer.workEnv.imgdata.seq = trim

# Update the GUI with the new work environment
update_workEnv()
```

##### View analysis log

View the analysis log, such as batch manager processing history.

```
>>> get_workEnv().history_trace

[{'caiman_motion_correction': {'max_shifts_x': 32, 'max_shifts_y': 32, 'iters_rigid': 1,
↪'name_rigid': 'Does not matter', 'max_dev': 20, 'strides': 196, 'overlaps': 98,
↪'upsample': 4, 'name_elas': 'a1_t2', 'output_bit_depth': 'Do not convert', 'bord_px':␣
↪5}}, {'cnmfe': {'Input': 'Current Work Environment', 'frate': 10.0, 'gSig': 10, 'bord_
↪px': 5, 'min_corr': 0.9600000000000001, 'min_pnr': 10, 'min_SNR': 1, 'r_values_min': 0.
↪7, 'decay_time': 2, 'rf': 80, 'stride': 40, 'gnb': 8, 'nb_patch': 8, 'k': 8, 'name_
↪corr_pnr': 'a8_t1', 'name_cnmfe': 'a1_t2', 'do_corr_pnr': False, 'do_cnmfe': True}}, {
↪'cnmfe': {'Input': 'Current Work Environment', 'frate': 10.0, 'gSig': 10, 'bord_px': 5,
↪ 'min_corr': 0.9600000000000001, 'min_pnr': 14, 'min_SNR': 1, 'r_values_min': 0.7,
↪'decay_time': 4, 'rf': 80, 'stride': 40, 'gnb': 8, 'nb_patch': 8, 'k': 8, 'name_corr_
↪pnr': '', 'name_cnmfe': 'a1_t2', 'do_corr_pnr': False, 'do_cnmfe': True}}]
```

##### Running scripts

You can use the *Script Editor* to run scripts in the Viewer console for automating tasks such as batch creation. It basically allows you to use the *viewer console* more conveniently with a text editor. The execution environment of the viewer console and script editor are identical.

Some example are provided for caiman modules and *stimulus mapping*.

### 1.49 Viewer Modules

#### 1.49.1 Batch Manager

**class** mesmerize.viewer.modules.batch_manager.**ModuleGUI**(*parent*, *run_batch: Optional[list] = None*, *testing: bool = False*)

GUI for the Batch Manager

**__init__**(*parent*, *run_batch: Optional[list] = None*, *testing: bool = False*)

Initialize self. See help(type(self)) for accurate signature.

**add_item**(*module: str*, *input_workEnv:* mesmerize.viewer.core.viewer_work_environment.ViewerWorkEnv, *input_params: dict*, *name: str = ''*, *info: dict = ''*) → uuid.UUID

Add an item to the currently open batch

**Parameters**

- **module** – The module to run from /batch_run_modules.
- **input_workEnv** – Input workEnv that the module will use
- **input_params** – Input params that the module will use. Depends on your subclass of BatchRunInterface.process() method
- **name** – A name for the batch item
- **info** – A dictionary with any metadata information to display in the scroll area label.

**Returns** UUID of the added item

**del_item**()

Delete the currently selected item from the batch and any corresponding dependents of the item's output

**df**

pandas.DataFrame that stores a "database" of information on the batch

**get_item_index**(*u: Union[uuid.UUID, str]*) → int

Get DataFrame index from UUID

**Parameters** **u** – UUID or str representing UUID

**Returns** numerical index of the DataFrame corresponding to the UUID

**get_selected_items**() → Tuple[List[int], List[uuid.UUID]]

Returns a list of numeric indices and uuids for the currently selected items

**load_item_input**(*viewers: Union[mesmerize.viewer.main_window.MainWindow, collections.UserList]*, *r: pandas.core.series.Series = None*, *UUID: uuid.UUID = None*, **args*)

Pass either the batch DataFrame row or UUID of the item of which to load the input into a viewer

**Parameters**

- **viewers** – ViewerWindow or list of ImageView

- **r** – Row of batch DataFrame corresponding to the selected item
- **UUID** – UUID of the item to load input from

**load_item_output**(*module: str, viewers: Union[mesmerize.viewer.main_window.MainWindow, mesmerize.pyqtgraphCore.imageview.ImageView.ImageView, collections.UserList], UUID: uuid.UUID*)

Calls subclass of BatchRunInterface.show_output()

**Parameters**

- **module** – The module name under /batch_run_modules that the batch item is from
- **viewers** – ViewerWindow, ImageView, or list of ViewerWindows
- **UUID** – UUID of the item to load output from

**process_batch**(*start_ix: Union[int, uuid.UUID] = 0, clear_viewers=False*)

Process everything in the batch by calling subclass of BatchRunInterface.process() for all items in batch

**Parameters**

- **start_ix** – Either DataFrame index (int) or UUID of the item to start from.
- **clear_viewers** – Clear work environments in all viewers that are open

**show_item_info**(*s: PyQt5.QtWidgets.QListWidgetItem*)

Shows any info (such as the batch module's params) in the meta-info label

##### 1.49.2 Tiff Module

**Uses the tifffile package created by Christoph Gohlke:** https://pypi.org/project/tifffile/

Can be used with scripts within Mesmerize for loading tiff files without using the API of *Viewer Core*

**class** mesmerize.viewer.modules.tiff_io.**ModuleGUI**(*parent, viewer_reference*)

**check_meta_path**() → bool

check if a file exists with the same name and the meta data extension specified by the selected meta data format

**load**(*tiff_path: str, method: str, axes_order: Optional[str] = None, meta_path: Optional[str] = None, meta_format: Optional[str] = None*) → *mesmerize.viewer.core.viewer_work_environment.ViewerWorkEnv*

Load a tiff file along with associated meta data

**Parameters**

- **tiff_path** – path to the tiff file
- **meta_path** – path to the json meta data file
- **method** – one of "asarray", "asarray-multi", or "imread" "asarray" and "asarray-multi" uses `tifffile.asarray()` "asarray-multi" is for multi-page tiffs "imread" uses `tifffile.imread()`
- **axes_order** – axes order, examples: txy, xyt, tzxy, xytz etc.
- **meta_format** – name of function from viewer.core.organize_meta that should be used to organize the meta data.

##### 1.49.3 Caiman Motion Correction

Front-end for Caiman NoRMCorre parameters entry

Can be used with scripts for adding batch items.

**See also:**

*User guide*

**class** mesmerize.viewer.modules.caiman_motion_correction.**ModuleGUI**(*parent*, *viewer_reference*)

**get_params**(*group_params: bool = False*) → dict
Get a dict of the set parameters :return: parameters dict :rtype: dict

**add_to_batch_elas_corr**()
Add a batch item with the currently set parameters and the current work environment.

##### 1.49.4 CNMF

Front-end for Caiman CNMF parameter entry

Can be used with scripts for adding batch items.

**See also:**

*User guide*

**class** mesmerize.viewer.modules.cnmf.**ModuleGUI**(*parent*, *viewer_reference*)

**get_params**(**args*, *group_params: bool = False*) → dict
Get a dict of the set parameters. If the work environment was loaded from a motion correction batch item it put the bord_px in the dict. Doesn't use any arguments

**Returns** parameters dict

**Return type** dict

**add_to_batch**(*params: dict = None*) → uuid.UUID
Add a CNMF batch item with the currently set parameters and the current work environment.

##### 1.49.5 CNMFE

Front-end for Caiman CNMFE parameter entry

Can be used with scripts for adding batch items.

**See also:**

*User guide*

**class** mesmerize.viewer.modules.cnmfe.**ModuleGUI**(*parent*, *viewer_reference*)

**get_params**(*item_type: str*, *group_params: bool = False*) → dict
Get a dict of the set parameters. If the work environment was loaded from a motion correction batch item it put the bord_px in the dict. Doesn't use any arguments

**Parameters** **item_type** – one of *corr_pnr* or *cnmfe*

**add_to_batch_corr_pnr**(*params: Optional[dict] = None*) → uuid.UUID
Add a Corr PNR batch item with the currently set parameters and the current work environment.

**add_to_batch_cnmfe**(*params: Optional[dict] = None*) → uuid.UUID
Add a CNMFE batch item with the currently set parameters and the current work environment.

##### 1.49.6 MESc Importer

MESc importer for exploring & importing image sequences from .mesc HDF5 files.

###### ModuleGUI

class mesmerize.viewer.modules.femtonics_mesc.**ModuleGUI**(*parent*, *viewer_ref*)

**mesc_navigator**
instance of `MEScNavigator`

**plot_widgets**
list of plot widgets

**set_file**(*path: str*, **args*)
Create an h5py file handle from the *.mesc* file at the given `path`.

`*args` are not used, its just there for compatibility with the decorator.

**Parameters**

- **path** (*str*) – path to the *.mes* file
- **args** – not used

**close_file**(**args*)
Close the file handle

**Parameters** **args** – `*args` not used, just there for compatibility with the decorator

**import_recording**(**args*)
Imports the chosen recording into the Viewer Work Environment based on the user selected hpath from the list widgets

`*args` not used, just there for compatibility with the decorator

###### MEScNavigator

Takes care of navigating through the HDF5 data structure of the .mesc file.

class mesmerize.viewer.modules.femtonics_mesc.**MEScNavigator**(*parent*, *list_widgets: List[mesmerize.pyqtgraphCore.widgets.ListWidget.ListW*

**sig_hpath_changed: PyQt5.QtCore.pyqtSignal**
emitted every time the hdf path changes

**sig_channel_doubleclicked: PyQt5.QtCore.pyqtSignal**
emitted when a `Channel` or `Curve` item is double clicked

**path: str**
system path to the hdf5 file

**file: h5py._hl.files.File**
h5py file handle

**session: str**
currently selected MSession

**sessions: List[str]**
list of `MSession` options available in current file

**listw_sessions: mesmerize.pyqtgraphCore.widgets.ListWidget.ListWidget**
ui list of `MSession` options

**unit: str**
currently selected `MUnit`

**units: List[str]**
list of `MUnit` options available in current `MSession`

**listw_units: mesmerize.pyqtgraphCore.widgets.ListWidget.ListWidget**
ui list of `MUnit` options

**channel: str**
currently selected `Channel`

**channels: List[str]**
list of `Channel` options available in current `MUnit`

**listw_channels: mesmerize.pyqtgraphCore.widgets.ListWidget.ListWidget**
ui list of `Channel` options

**set_file_path**(*path: str*)
set the path to the .mesc file

**Parameters** **path** – path to the .mesc hdf5 file

**close_file**()
Close the h5py file handle that is currently open.

**set_session**(*key: Union[str, PyQt5.QtWidgets.QListWidgetItem]*)
Set the MSession option

**Parameters** **key** – a valid `MSession`

**Returns**

**set_unit**(*key: Union[str, PyQt5.QtWidgets.QListWidgetItem]*)
Set the MUnit option

**Parameters** **key** – a valid `MUnit`

**Returns**

**set_channel**(*key: Union[str, PyQt5.QtWidgets.QListWidgetItem]*)
Set a Channel or Curve option

**Parameters** **key** – a valid `Channel` or `Curve`

**Returns**

**get_hpath**(*astype: type*) → Union[str, list, dict]
get the current hdf path

**Parameters** **astype** – one of *str*, *list*, or *dict*

**Returns** the hdf path as the chosen data type

#### 1.50 ROI Manager

##### 1.50.1 Video Tutorial

##### 1.50.2 ModuleGUI

The GUI QDockWidget that communicates with the *back-end managers*

**class** mesmerize.viewer.modules.roi_manager.**ModuleGUI**(*parent*, *viewer_reference*)
The GUI front-end for the ROI Manager module

**__init__**(*parent*, *viewer_reference*)
Instantiate attributes

**manager**
The back-end manager instance.

**eventFilter**(*QObject*, *QEvent*)
Set some keyboard shortcuts

**slot_delete_roi_menu**()
Delete the currently selected ROI

**start_backend**(*type_str: str*)
Choose backend, one of the Manager classes in the managers module.

**start_manual_mode**()
Start in manual mode. Creates a new back-end manager instance (Uses ManagerManual)

**add_manual_roi**(*shape: str*)
Add a manual ROI. Just calls ManagerManual.add_roi

**package_for_project**() → dict
Gets all the ROI states so that they can be packaged along with the rest of the work environment to be saved as a project Sample

**set_all_from_states**(*states: dict*)
Set all the ROIs from a states dict. Instantiates the appropriate back-end Manager

**import_from_imagej**()
Import ROIs from ImageJ zip file

##### 1.50.3 Managers

The back-end managers that are used by the *ROI Manager ModuleGUI*

The managers hold instances of *ROIs* in an instance of *ROIList*

##### AbstractBaseManager

Subclass this if you want to make your own Manager Back-end.

**class** mesmerize.viewer.modules.roi_manager_modules.managers.**AbstractBaseManager**(*parent*, *ui*, *viewer_interface: mesmerize.viewer.core.common.View*

Base ROI Manager

**__init__**(*parent*, *ui*, *viewer_interface: mesmerize.viewer.core.common.ViewerUtils*)
Set the common attributes

**Parameters**

- **parent** – The ModuleGUI QDockWidget instance
- **ui** – The ui of the ModuleGUI QDockWidget instance,
- **viewer_interface** – A ViewerUtils instance for accessing the Viewer the parent QDockWidget belongs to

**roi_list**
The ROIList instance that stores the list of ROIs

**abstract add_roi**(**args*, ***kwargs*)
Method for adding an ROI, must be implemented in subclass

**is_empty**() → bool
Return true if the ROI list is empty, else return False

**get_all_states**() → dict
Get the ROI states for all ROIs in self.roi_list so that they can be restored. The appropriate manager is instantiated based on the 'roi_type' key of the returned dict

**get_plot_item**() → mesmerize.pyqtgraphCore.graphicsItems.PlotDataItem.PlotDataItem
Get the viewer plot item that is associated to these ROIs

**clear**()
Cleanup of all ROIs in the list

**__del__**()
Cleanup of all ROIs in the list and deletes the manager instance. Used when switching modes.

**__weakref__**
list of weak references to the object (if defined)

##### ManagerManual

**class** mesmerize.viewer.modules.roi_manager_modules.managers.**ManagerManual**(*parent*, *ui*, *viewer_interface*)

Bases: *mesmerize.viewer.modules.roi_manager_modules.managers.AbstractBaseManager*

The Manager for the Manual mode

**__init__**(*parent*, *ui*, *viewer_interface*)
Set the common attributes

**Parameters**

- **parent** – The ModuleGUI QDockWidget instance
- **ui** – The ui of the ModuleGUI QDockWidget instance,

- **viewer_interface** – A ViewerUtils instance for accessing the Viewer the parent QDockWidget belongs to

**create_roi_list**()
Create a new empty ROI list instance for storing Manual ROIs

**add_roi**(*shape: str*) → *mesmerize.viewer.modules.roi_manager_modules.roi_types.ManualROI*
Add an ROI to the list

**Parameters** **shape** – either "PolyLineROI" or "EllipseROI"

**restore_from_states**(*states: dict*)
Restore ROIs from states

**get_all_states**() → dict
Get the ROI states so that they can be restored later

**import_from_imagej**(*path: str*)
Uses read-roi package created by Hadrien Mary. https://pypi.org/project/read-roi/

**Parameters** **path** – Full path to the ImageJ ROIs zip file

#### ManagerScatterROI

class mesmerize.viewer.modules.roi_manager_modules.managers.**ManagerScatterROI**(*parent*, *ui*, *viewer_interface:* mesmerize.viewer.core.common.ViewerU

Bases: *mesmerize.viewer.modules.roi_manager_modules.managers.AbstractBaseManager*

Manager for unmoveable ROIs drawn using scatterplots

**__init__**(*parent*, *ui*, *viewer_interface:* mesmerize.viewer.core.common.ViewerUtils)
Set the common attributes

**Parameters**

- **parent** – The ModuleGUI QDockWidget instance
- **ui** – The ui of the ModuleGUI QDockWidget instance,
- **viewer_interface** – A ViewerUtils instance for accessing the Viewer the parent QDockWidget belongs to

**add_roi**(*curve: numpy.ndarray*, *xs: numpy.ndarray*, *ys: numpy.ndarray*, *metadata: Optional[dict] = None*, *dfof_data: Optional[numpy.ndarray] = None*, *spike_data: Optional[numpy.ndarray] = None*) → *mesmerize.viewer.modules.roi_manager_modules.roi_types.ScatterROI*
Add a single ROI

*xs* and *ys* arguments are 1D numpy arrays.

**Parameters**

- **curve** – curve data, 1-D array, y values/intensity values
- **xs** – x-values for the scatter plot to spatially illustrate the ROI
- **ys** – corresponding y-values for the scatter plot to spatially illustrate the ROI
- **metadata** – Any metadata for this ROI

**Returns** ScatterROI object

**restore_from_states**(*states: dict*)
Restore from states, such as when these ROIs are saved with a Project Sample

**create_roi_list**()
Create empty ROI List

**set_spot_size**(*size: int*)
Set the spot size for the scatter plot which illustrates the ROI

#### ManagerVolROI

class mesmerize.viewer.modules.roi_manager_modules.managers.**ManagerVolROI**(*parent*, *ui*, *viewer_interface: mesmerize.viewer.core.common.ViewerUtils*)

Bases: *mesmerize.viewer.modules.roi_manager_modules.managers.ManagerScatterROI*

Manager for 3D ROIs

**__init__**(*parent*, *ui*, *viewer_interface: mesmerize.viewer.core.common.ViewerUtils*)
Set the common attributes

**Parameters**

- **parent** – The ModuleGUI QDockWidget instance
- **ui** – The ui of the ModuleGUI QDockWidget instance,
- **viewer_interface** – A ViewerUtils instance for accessing the Viewer the parent QDockWidget belongs to

**set_zlevel**(*z: int*)
Set the current z-level to be visible in the viewer

**create_roi_list**()
Create new empty ROI list

#### ManagerVolCNMF

class mesmerize.viewer.modules.roi_manager_modules.managers.**ManagerVolCNMF**(*parent*, *ui*, *viewer_interface*)

Bases: *mesmerize.viewer.modules.roi_manager_modules.managers.ManagerVolROI*

Manager for 3D CNMF based ROIs

**__init__**(*parent*, *ui*, *viewer_interface*)
Set the common attributes

**Parameters**

- **parent** – The ModuleGUI QDockWidget instance
- **ui** – The ui of the ModuleGUI QDockWidget instance,
- **viewer_interface** – A ViewerUtils instance for accessing the Viewer the parent QDockWidget belongs to

**create_roi_list**()
Create new empty ROI list

**add_all_components**(*cnmf_data_dict: dict*, *input_params_dict: dict*)
Add all components from a CNMF(E) output. Arguments correspond to CNMF(E) outputs

**Parameters**

- **cnmf_data_dict** – CNMF results data directly from the HDF5 file
- **input_params_dict** – dict of input params, from the batch manager
- **calc_raw_min_max** – Calculate raw min & max for each ROI

**Returns**

**add_roi**()
Not implemented, uses add_all_components to import all ROIs instead

**restore_from_states**(*states: dict*)
Restore from states, such as when these ROIs are saved with a Project Sample

**get_all_states**() → dict
Get all states so that they can be restored

**update_idx_components**(*ix: int*)
Update idx_components if the user manually delete an ROI

**set_spot_size**(*size: int*)
Set the spot size for the scatter plot which illustrates the ROI

#### ManagerCNMFROI

class mesmerize.viewer.modules.roi_manager_modules.managers.**ManagerCNMFROI**(*parent*, *ui*, *viewer_interface*)

Bases: *mesmerize.viewer.modules.roi_manager_modules.managers.AbstractBaseManager*

Manager for ROIs imported from CNMF or CNMFE outputs

**__init__**(*parent*, *ui*, *viewer_interface*)
Instantiate necessary attributes

**create_roi_list**()
Create empty CNMFROI list

**add_all_components**(*cnmf_data_dict*, *input_params_dict*, *calc_raw_min_max=False*)
Add all components from a CNMF(E) output. Arguments correspond to CNMF(E) outputs

**Parameters**

- **cnmf_data_dict** – CNMF results data directly from the HDF5 file
- **input_params_dict** – dict of input params, from the batch manager
- **calc_raw_min_max** – Calculate raw min & max for each ROI

**Returns**

**add_roi**()
Not implemented, uses add_all_components to import all ROIs instead

**restore_from_states**(*states: dict*)
Restore from states, such as when these ROIs are saved with a Project Sample

**get_all_states**() → dict
Get all states so that they can be restored

**update_idx_components**(*ix: int*)
Update idx_components if the user manually delete an ROI

##### 1.50.4 ROI List

Used for holding instance of *ROIs*

class mesmerize.viewer.modules.roi_manager_modules.roi_list.**ROIList**(*ui*, *roi_types: type*, *viewer_interface:* mesmerize.viewer.core.common.ViewerUtils)

A list for holding ROIs of one type

**__init__**(*ui*, *roi_types: type*, *viewer_interface:* mesmerize.viewer.core.common.ViewerUtils)
Instantiate

**Parameters**

- **ui** – The ui from the parent ModuleGUI, used to interact with the ROI list widget etc.
- **roi_types** – The type of ROI that this list will hold
- **viewer_interface** – ViewerUtils instance for interacting with the parent Viewer

**list_widget**
ROI list widget

**list_widget_tags**
Tags list widget

**vi**
ViewrUtils instance

**current_index**
Current index (int)

**previous_index**
Previous index (int)

**append**(*roi: Union[*mesmerize.viewer.modules.roi_manager_modules.roi_types.ScatterROI, mesmerize.viewer.modules.roi_manager_modules.roi_types.ManualROI, *mesmerize.viewer.modules.roi_manager_modules.roi_types.VolMultiCNMFROI]*, *add_to_list_widget: bool = True*)
Add an ROI instance to the list

**clear_**()
Cleanup of the list

**__delitem__**(*key*)
Delete an ROI from the list and cleanup from the viewer, reindex the colors etc.

**disconnect_all**()
Disconnect signals from the parent GUI

**_reindex_list_widget**()
Reindex ROI list

**reindex_colormap**(*random_shuffle=False*)
Reindex the colors so they sequentially follow the HSV colormap

**__getitem__**(*item*) → Union[*mesmerize.viewer.modules.roi_manager_modules.roi_types.ManualROI*, *mesmerize.viewer.modules.roi_manager_modules.roi_types.ScatterROI*]
Get an item (ROI) from the list

**set_current_index**(*ix: int*)
Set the current index

**highlight_roi**(*roi: Union[mesmerize.viewer.modules.roi_manager_modules.roi_types.ManualROI, mesmerize.viewer.modules.roi_manager_modules.roi_types.ScatterROI]*)
Highlight an ROI in white, both the spatial visualization and the curve

**highlight_curve**(*ix: int*)
Highlight the curve corresponding to the ROI at the passed index

**set_previous_index**()
Set the previous_index attribute

**slot_show_all_checkbox_clicked**(*b: bool*)
Show all ROIs in the viewer overlay visualization and curves

**_show_graphics_object**(*ix: int*)
Show the ROI at the passed index in the viewer overlay visualization

**_hide_graphics_object**(*ix: int*)
Hide the ROI at the passed index in the viewer overlay visualization

**_show_all_graphics_objects**()
Show all ROIs in the viewer overlay visualization

**_hide_all_graphics_objects**()
Hide all ROIs in the viewer overlay visualization

**plot_manual_roi_regions**()
Plot the ROI curves from the regions of all ManualROI instances in the list

**set_pg_roi_plot**(*ix: int*)
Plot the ROI curve from the region of the ManualROI instance at the passed index

**set_list_widget_tags**()
Set the tags list for the ROI at the current index

**update_roi_defs_from_configuration**()
Update ROI_DEFs in the Tags list from the project configuration

**__weakref__**
list of weak references to the object (if defined)

#### 1.50.5 ROI Types

A list of these are held by an instance of *ROIList*

##### AbstractBaseROI

**class** mesmerize.viewer.modules.roi_manager_modules.roi_types.**_AbstractBaseROI**(*curve_plot_item: mesmerize.pyqtgraphCore.graphicsItems view_box: mesmerize.pyqtgraphCore.graphicsItems state: Optional[dict]*)
Abstract base class defining an ROI that works with the ROIList and ROI Managers. Inherit from this or BaseROI to make a new ROI class

**abstract __init__**(*curve_plot_item: mesmerize.pyqtgraphCore.graphicsItems.PlotDataItem.PlotDataItem*, *view_box: mesmerize.pyqtgraphCore.graphicsItems.ViewBox.ViewBox.ViewBox*, *state: Optional[dict]*)

Minimum required attributes

**Parameters**

- **curve_plot_item** – The plot item that is used for display the curves in the viewer
- **view_box** – ViewBox containing the image sequence, used for overlaying the ROIs on top of the image
- **state** – ROI state, used for restoring the ROIs. Pass None is not restoring an ROI from a state dict

**abstract get_roi_graphics_object**() → PyQt5.QtWidgets.QGraphicsObject

Get the QGraphicsObject used for visualization of the spatial localization of the ROI

**abstract set_roi_graphics_object**(**args*, ***kwargs*)

Set the QGraphicsObject used for visualization of the spatial localization of the ROI

**abstract reset_color**()

Reset the color of this ROI back to the original color

**abstract set_original_color**(*color*)

Set the original color for this ROI

**Parameters** **color** – 1D numpy array of 4 floating point numbers (range 0 - 255) in RBGA format, [R, G, B, A]

**abstract get_color**() → numpy.ndarray

Get the current color of this ROI

**Returns** 1D numpy array of 4 floating point numbers (range 0 - 255) in RBGA format, [R, G, B, A]

**Return type** np.ndarray

**abstract set_color**(*color*, **args*, ***kwargs*)

Set the current color of this ROI

**Parameters** **color** – 1D numpy array of 4 floating point numbers (range 0 - 255) in RBGA format, [R, G, B, A]

**abstract set_text**(*text: str*)

Not implemented

**abstract set_tag**(*roi_def: str*, *tag: str*)

Set a tag for the passed roi_def

**Parameters**

- **roi_def** – The ROI_DEF that should be tagged
- **tag** – The tag to label for the passed ROI_DEF/ROI Type

**abstract get_tag**(*roi_def*) → str

Get the tag that is set to the passed 'roi_def'

**Return type** str

**abstract get_all_tags**() → dict

Get all the tags for all the ROI_DEFs

**Return type** dict

`abstract add_to_viewer()`
Add this ROI to the viewer.

`abstract remove_from_viewer()`
Remove this ROI from the viewer

`abstract to_state()`
Get the current state for this ROI so that it can be restored later

`abstract classmethod from_state`(*curve_plot_item: mesmerize.pyqtgraphCore.graphicsItems.PlotDataItem.PlotDataItem*, *view_box: mesmerize.pyqtgraphCore.graphicsItems.ViewBox.ViewBox.ViewBox*, *state: dict*)
Restore this ROI from a state

**Parameters**

- **curve_plot_item** – The plot item that is used for display the curves in the viewer
- **view_box** – ViewBox containing the image sequence, used for overlaying the ROIs on top of the image
- **state** – ROI state, used for restoring the ROIs. Pass None is not restoring an ROI from a state dict

`__weakref__`
list of weak references to the object (if defined)

#### BaseROI

Subclass from this if you want to make your own ROI Type.

`class mesmerize.viewer.modules.roi_manager_modules.roi_types.BaseROI`(*curve_plot_item: mesmerize.pyqtgraphCore.graphicsItems.PlotDataIte[illegible]*, *view_box: mesmerize.pyqtgraphCore.graphicsItems.ViewBox.Vi[illegible]*, *state: Optional[dict] = None*, *metadata: Optional[dict] = None*)

Bases: `mesmerize.viewer.modules.roi_manager_modules.roi_types._AbstractBaseROI`

A base class that is used by ManualROI and CNMFEROI Inherit from this to make a new ROI class

`__init__`(*curve_plot_item: mesmerize.pyqtgraphCore.graphicsItems.PlotDataItem.PlotDataItem*, *view_box: mesmerize.pyqtgraphCore.graphicsItems.ViewBox.ViewBox.ViewBox*, *state: Optional[dict] = None*, *metadata: Optional[dict] = None*)
Instantiate common attributes

**Parameters**

- **curve_plot_item** – The plot item that is used for display the curves in the viewer
- **view_box** – ViewBox containing the image sequence, used for overlaying the ROIs on top of the image
- **state** – ROI state, used for restoring the ROIs. Pass None is not restoring an ROI from a state dict

**get_roi_graphics_object**() → PyQt5.QtWidgets.QGraphicsObject
Get the QGraphicsObject used for visualization of the spatial localization of the ROI

**set_roi_graphics_object**(**args*, ***kwargs*)
Set the QGraphicsObject used for visualization of the spatial localization of the ROI

**reset_color**()
Reset the color of this ROI back to the original color

**set_original_color**(*color*)
Set the original color for this ROI

**Parameters** **color** – 1D numpy array of 4 floating point numbers (range 0 - 255) in RBGA format, [R, G, B, A]

**get_color**()
Get the current color of this ROI

**Returns** 1D numpy array of 4 floating point numbers (range 0 - 255) in RBGA format, [R, G, B, A]

**Return type** np.ndarray

**set_color**(*color: Union[numpy.ndarray, str]*, **args*, ***kwargs*)
Set the current color of this ROI

**Parameters** **color** – 1D numpy array of 4 floating point numbers (range 0 - 255) in RBGA format, [R, G, B, A]

**set_text**(*text: str*)
Not implemented

**set_tag**(*roi_def: str*, *tag: str*)
Set a tag for the passed roi_def

**Parameters**

- **roi_def** – The ROI_DEF that should be tagged
- **tag** – The tag to label for the passed ROI_DEF/ROI Type

**get_tag**(*roi_def*) → str
Get the tag that is set to the passed 'roi_def'

**Return type** str

**get_all_tags**() → dict
Get all the tags for all the ROI_DEFs

**Return type** dict

**add_to_viewer**()
Add this ROI to the viewer.

**remove_from_viewer**()
Remove this ROI from the viewer

**to_state**()
Must be implemented in subclass

**classmethod from_state**(*curve_plot_item: mesmerize.pyqtgraphCore.graphicsItems.PlotDataItem.PlotDataItem*, *view_box: mesmerize.pyqtgraphCore.graphicsItems.ViewBox.ViewBox.ViewBox*, *state: dict*)
Restore this ROI from a state

**Parameters**

- **curve_plot_item** – The plot item that is used for display the curves in the viewer
- **view_box** – ViewBox containing the image sequence, used for overlaying the ROIs on top of the image
- **state** – ROI state, used for restoring the ROIs. Pass None is not restoring an ROI from a state dict

#### ManualROI

class mesmerize.viewer.modules.roi_manager_modules.roi_types.**ManualROI**(*curve_plot_item: mesmerize.pyqtgraphCore.graphicsItems.PlotDat[illegible], roi_graphics_object: mesmerize.pyqtgraphCore.graphicsItems.ROI.RO[illegible], view_box: mesmerize.pyqtgraphCore.graphicsItems.ViewBo[illegible], state: Optional[dict] = None*, *spike_data: Optional[numpy.ndarray] = None*, *dfof_data: Optional[numpy.ndarray] = None*)

Bases: `mesmerize.viewer.modules.roi_manager_modules.roi_types.BaseROI`

A class manually drawn ROIs

**__init__**(*curve_plot_item: mesmerize.pyqtgraphCore.graphicsItems.PlotDataItem.PlotDataItem*, *roi_graphics_object: mesmerize.pyqtgraphCore.graphicsItems.ROI.ROI*, *view_box: mesmerize.pyqtgraphCore.graphicsItems.ViewBox.ViewBox.ViewBox*, *state: Optional[dict] = None*, *spike_data: Optional[numpy.ndarray] = None*, *dfof_data: Optional[numpy.ndarray] = None*)

**property curve_data: tuple**

tuple of (xs, ys)

**Type** return

**get_roi_graphics_object**() → mesmerize.pyqtgraphCore.graphicsItems.ROI.ROI

Get the QGraphicsObject used for visualization of the spatial localization of the ROI

**set_roi_graphics_object**(*graphics_object: mesmerize.pyqtgraphCore.graphicsItems.ROI.ROI*)

Set the QGraphicsObject used for visualization of the spatial localization of the ROI

**to_state**()

Must be implemented in subclass

**classmethod from_state**(*curve_plot_item: mesmerize.pyqtgraphCore.graphicsItems.PlotDataItem.PlotDataItem*, *view_box: mesmerize.pyqtgraphCore.graphicsItems.ViewBox.ViewBox.ViewBox*, *state: dict*)

Restore this ROI from a state

**Parameters**

- **curve_plot_item** – The plot item that is used for display the curves in the viewer
- **view_box** – ViewBox containing the image sequence, used for overlaying the ROIs on top of the image
- **state** – ROI state, used for restoring the ROIs. Pass None is not restoring an ROI from a state dict

**reset_color**()
Reset the color of this ROI back to the original color

**set_original_color**(*color*)
Set the original color for this ROI

**Parameters** **color** – 1D numpy array of 4 floating point numbers (range 0 - 255) in RBGA format, [R, G, B, A]

**get_color**()
Get the current color of this ROI

**Returns** 1D numpy array of 4 floating point numbers (range 0 - 255) in RBGA format, [R, G, B, A]

**Return type** np.ndarray

**set_color**(*color: Union[numpy.ndarray, str]*, **args*, ***kwargs*)
Set the current color of this ROI

**Parameters** **color** – 1D numpy array of 4 floating point numbers (range 0 - 255) in RBGA format, [R, G, B, A]

**set_text**(*text: str*)
Not implemented

**set_tag**(*roi_def: str*, *tag: str*)
Set a tag for the passed roi_def

**Parameters**

- **roi_def** – The ROI_DEF that should be tagged
- **tag** – The tag to label for the passed ROI_DEF/ROI Type

**get_tag**(*roi_def*) → str
Get the tag that is set to the passed 'roi_def'

**Return type** str

**get_all_tags**() → dict
Get all the tags for all the ROI_DEFs

**Return type** dict

**add_to_viewer**()
Add this ROI to the viewer.

**remove_from_viewer**()
Remove this ROI from the viewer

##### ScatterROI

**class** mesmerize.viewer.modules.roi_manager_modules.roi_types.**ScatterROI**(*curve_plot_item: mesmerize.pyqtgraphCore.graphicsItems.PlotDa* *view_box: mesmerize.pyqtgraphCore.graphicsItems.ViewBo* *state: Optional[dict] = None, curve_data: Optional[numpy.ndarray] = None, xs: Optional[numpy.ndarray] = None, ys: Optional[numpy.ndarray] = None, metadata: Optional[dict] = None, spike_data: Optional[numpy.ndarray] = None, dfof_data: Optional[numpy.ndarray] = None, **kwargs*)

Bases: `mesmerize.viewer.modules.roi_manager_modules.roi_types.BaseROI`

A class for unmoveable ROIs drawn using scatter points

**__init__**(*curve_plot_item: mesmerize.pyqtgraphCore.graphicsItems.PlotDataItem.PlotDataItem, view_box: mesmerize.pyqtgraphCore.graphicsItems.ViewBox.ViewBox.ViewBox, state: Optional[dict] = None, curve_data: Optional[numpy.ndarray] = None, xs: Optional[numpy.ndarray] = None, ys: Optional[numpy.ndarray] = None, metadata: Optional[dict] = None, spike_data: Optional[numpy.ndarray] = None, dfof_data: Optional[numpy.ndarray] = None, **kwargs*)

**Parameters**

- **curve_plot_item** –
- **view_box** –
- **state** –
- **curve_data** – 1D numpy array of y values
- **kwargs** –

**set_curve_data**(*y_vals: numpy.ndarray, set_plot: bool = True*)
Set the curve data

**to_state**() → dict
Must be implemented in subclass

**set_roi_graphics_object**(*xs: numpy.ndarray, ys: numpy.ndarray*)
Set the QGraphicsObject used for visualization of the spatial localization of the ROI

**get_roi_graphics_object**() → mesmerize.pyqtgraphCore.graphicsItems.ScatterPlotItem.ScatterPlotItem
Get the QGraphicsObject used for visualization of the spatial localization of the ROI

**classmethod from_state**(*curve_plot_item: mesmerize.pyqtgraphCore.graphicsItems.PlotDataItem.PlotDataItem*, *view_box: mesmerize.pyqtgraphCore.graphicsItems.ViewBox.ViewBox.ViewBox*, *state: dict*, ***kwargs*)

Restore this ROI from a state

**Parameters**

- **curve_plot_item** – The plot item that is used for display the curves in the viewer
- **view_box** – ViewBox containing the image sequence, used for overlaying the ROIs on top of the image
- **state** – ROI state, used for restoring the ROIs. Pass None is not restoring an ROI from a state dict

**reset_color**()

Reset the color of this ROI back to the original color

**set_original_color**(*color*)

Set the original color for this ROI

**Parameters color** – 1D numpy array of 4 floating point numbers (range 0 - 255) in RBGA format, [R, G, B, A]

**get_color**()

Get the current color of this ROI

**Returns** 1D numpy array of 4 floating point numbers (range 0 - 255) in RBGA format, [R, G, B, A]

**Return type** np.ndarray

**set_color**(*color: Union[numpy.ndarray, str]*, **args*, ***kwargs*)

Set the current color of this ROI

**Parameters color** – 1D numpy array of 4 floating point numbers (range 0 - 255) in RBGA format, [R, G, B, A]

**set_text**(*text: str*)

Not implemented

**set_tag**(*roi_def: str*, *tag: str*)

Set a tag for the passed roi_def

**Parameters**

- **roi_def** – The ROI_DEF that should be tagged
- **tag** – The tag to label for the passed ROI_DEF/ROI Type

**get_tag**(*roi_def*) → str

Get the tag that is set to the passed 'roi_def'

**Return type** str

**get_all_tags**() → dict

Get all the tags for all the ROI_DEFs

**Return type** dict

**add_to_viewer**()

Add this ROI to the viewer.

**remove_from_viewer**()

Remove this ROI from the viewer

#### VolCNMF

class mesmerize.viewer.modules.roi_manager_modules.roi_types.**VolCNMF**(*curve_plot_item: mesmerize.pyqtgraphCore.graphicsItems.PlotDataIte*[illegible] *view_box: mesmerize.pyqtgraphCore.graphicsItems.ViewBox.Vi*[illegible] *cnmf_idx: Optional[int] = None*, *curve_data: Optional[numpy.ndarray] = None*, *contour: Optional[dict] = None*, *state: Optional[dict] = None*, *spike_data: Optional[numpy.ndarray] = None*, *dfof_data: Optional[numpy.ndarray] = None*, *metadata: Optional[dict] = None*, *zlevel: int = 0*)

Bases: `mesmerize.viewer.modules.roi_manager_modules.roi_types.ScatterROI`

3D ROI for CNMF data

**__init__**(*curve_plot_item: mesmerize.pyqtgraphCore.graphicsItems.PlotDataItem.PlotDataItem*, *view_box: mesmerize.pyqtgraphCore.graphicsItems.ViewBox.ViewBox.ViewBox*, *cnmf_idx: Optional[int] = None*, *curve_data: Optional[numpy.ndarray] = None*, *contour: Optional[dict] = None*, *state: Optional[dict] = None*, *spike_data: Optional[numpy.ndarray] = None*, *dfof_data: Optional[numpy.ndarray] = None*, *metadata: Optional[dict] = None*, *zlevel: int = 0*)

**Parameters**

- **curve_plot_item** –
- **view_box** –
- **state** –
- **curve_data** – 1D numpy array of y values
- **kwargs** –

**to_state**() → dict

Must be implemented in subclass

**set_roi_graphics_object**()

Set the QGraphicsObject used for visualization of the spatial localization of the ROI

**set_zlevel**(*z: int*)

Set the z-level of the ROI to correspond with the z-level of the image.

Different from *setZValue*!!

**get_roi_graphics_object**() → mesmerize.pyqtgraphCore.graphicsItems.ScatterPlotItem.ScatterPlotItem

Get the QGraphicsObject used for visualization of the spatial localization of the ROI

**reset_color()**
Reset the color of this ROI back to the original color

**set_original_color**(*color*)
Set the original color for this ROI

**Parameters** **color** – 1D numpy array of 4 floating point numbers (range 0 - 255) in RBGA format, [R, G, B, A]

**get_color()**
Get the current color of this ROI

**Returns** 1D numpy array of 4 floating point numbers (range 0 - 255) in RBGA format, [R, G, B, A]

**Return type** np.ndarray

**set_color**(*color: Union[numpy.ndarray, str]*, **args*, ***kwargs*)
Set the current color of this ROI

**Parameters** **color** – 1D numpy array of 4 floating point numbers (range 0 - 255) in RBGA format, [R, G, B, A]

**set_text**(*text: str*)
Not implemented

**set_tag**(*roi_def: str*, *tag: str*)
Set a tag for the passed roi_def

**Parameters**

- **roi_def** – The ROI_DEF that should be tagged
- **tag** – The tag to label for the passed ROI_DEF/ROI Type

**get_tag**(*roi_def*) → str
Get the tag that is set to the passed 'roi_def'

**Return type** str

**get_all_tags()** → dict
Get all the tags for all the ROI_DEFs

**Return type** dict

**add_to_viewer()**
Add this ROI to the viewer.

**remove_from_viewer()**
Remove this ROI from the viewer

**classmethod from_state**(*curve_plot_item: mesmerize.pyqtgraphCore.graphicsItems.PlotDataItem.PlotDataItem*, *view_box: mesmerize.pyqtgraphCore.graphicsItems.ViewBox.ViewBox.ViewBox*, *state: dict*, ***kwargs*)
Restore this ROI from a state

**Parameters**

- **curve_plot_item** – The plot item that is used for display the curves in the viewer
- **view_box** – ViewBox containing the image sequence, used for overlaying the ROIs on top of the image

- **state** – ROI state, used for restoring the ROIs. Pass None is not restoring an ROI from a state dict

**set_curve_data**(*y_vals: numpy.ndarray*, *set_plot: bool = True*)
Set the curve data

#### CNMFROI

class mesmerize.viewer.modules.roi_manager_modules.roi_types.**CNMFROI**(*curve_plot_item: mesmerize.pyqtgraphCore.graphicsItems.PlotDataIte[illegible]*, *view_box: mesmerize.pyqtgraphCore.graphicsItems.ViewBox.Vi[illegible]*, *cnmf_idx: Optional[int] = None*, *curve_data: Optional[numpy.ndarray] = None*, *contour: Optional[dict] = None*, *state: Optional[dict] = None*, *spike_data: Optional[numpy.ndarray] = None*, *dfof_data: Optional[numpy.ndarray] = None*, *metadata: Optional[dict] = None*, ***kwargs*)

Bases: *mesmerize.viewer.modules.roi_manager_modules.roi_types.ScatterROI*

A class for ROIs imported from CNMF(E) output data

**get_roi_graphics_object**() → mesmerize.pyqtgraphCore.graphicsItems.ScatterPlotItem.ScatterPlotItem
Get the QGraphicsObject used for visualization of the spatial localization of the ROI

**set_roi_graphics_object**(*xs: numpy.ndarray*, *ys: numpy.ndarray*)
Set the QGraphicsObject used for visualization of the spatial localization of the ROI

**reset_color**()
Reset the color of this ROI back to the original color

**set_original_color**(*color*)
Set the original color for this ROI

**Parameters** **color** – 1D numpy array of 4 floating point numbers (range 0 - 255) in RBGA format, [R, G, B, A]

**get_color**()
Get the current color of this ROI

**Returns** 1D numpy array of 4 floating point numbers (range 0 - 255) in RBGA format, [R, G, B, A]

**Return type** np.ndarray

**set_color**(*color: Union[numpy.ndarray, str]*, **args*, ***kwargs*)
Set the current color of this ROI

**Parameters** **color** – 1D numpy array of 4 floating point numbers (range 0 - 255) in RBGA format, [R, G, B, A]

**set_text**(*text: str*)
Not implemented

**set_tag**(*roi_def: str*, *tag: str*)
Set a tag for the passed roi_def

**Parameters**

- **roi_def** – The ROI_DEF that should be tagged
- **tag** – The tag to label for the passed ROI_DEF/ROI Type

**get_tag**(*roi_def*) → str
Get the tag that is set to the passed 'roi_def'

**Return type** str

**get_all_tags**() → dict
Get all the tags for all the ROI_DEFs

**Return type** dict

**add_to_viewer**()
Add this ROI to the viewer.

**remove_from_viewer**()
Remove this ROI from the viewer

**classmethod from_state**(*curve_plot_item: mesmerize.pyqtgraphCore.graphicsItems.PlotDataItem.PlotDataItem*, *view_box: mesmerize.pyqtgraphCore.graphicsItems.ViewBox.ViewBox.ViewBox*, *state: dict*, ***kwargs*)
Restore this ROI from a state

**Parameters**

- **curve_plot_item** – The plot item that is used for display the curves in the viewer
- **view_box** – ViewBox containing the image sequence, used for overlaying the ROIs on top of the image
- **state** – ROI state, used for restoring the ROIs. Pass None is not restoring an ROI from a state dict

**set_curve_data**(*y_vals: numpy.ndarray*, *set_plot: bool = True*)
Set the curve data

**__init__**(*curve_plot_item: mesmerize.pyqtgraphCore.graphicsItems.PlotDataItem.PlotDataItem*, *view_box: mesmerize.pyqtgraphCore.graphicsItems.ViewBox.ViewBox.ViewBox*, *cnmf_idx: Optional[int] = None*, *curve_data: Optional[numpy.ndarray] = None*, *contour: Optional[dict] = None*, *state: Optional[dict] = None*, *spike_data: Optional[numpy.ndarray] = None*, *dfof_data: Optional[numpy.ndarray] = None*, *metadata: Optional[dict] = None*, ***kwargs*)
Instantiate attributes.

**Type** curve_data: np.ndarray

**Parameters**

- **curve_data** – 1D numpy array of y values
- **cnmf_idx** – original index of the ROI from cnmf idx_components

**to_state**() → dict
Must be implemented in subclass

##### 1.50.6 Basic Examples

These examples can be run through the viewer console or *Script editor* to interact with the ROIs.

**See also:**

*Back-end ROI Manager APIs*, *ROIList API*, *ROI Type APIs*

Get the back-end ROI Manager, see *ROI Manager APIs*

```
>>> get_workEnv().roi_manager

<mesmerize.viewer.modules.roi_manager_modules.managers.ManagerCNMFROI object at␣
↪0x7f01b8780668>``
```

Get the ROI List, see *ROIList API*

```
>>> get_workEnv().roi_manager.roi_list

[<mesmerize.viewer.modules.roi_manager_modules.roi_types.CNMFROI object at␣
↪0x7f01bc78b278>, <mesmerize.viewer.modules.roi_manager_modules.roi_types.CNMFROI␣
↪object at 0x7f01bc817630>, <mesmerize.viewer.modules.roi_manager_modules.roi_types.
↪CNMFROI object at 0x7f01bc817668>, <mesmerize.viewer.modules.roi_manager_modules.roi_
↪types.CNMFROI object at 0x7f01bc7c5438>, <mesmerize.viewer.modules.roi_manager_modules.
↪roi_types.CNMFROI object at 0x7f01bc7c5208>]
```

Work with an ROI object, see *ROI Type APIs*

```
# Get the curve data of an ROI
>>> get_workEnv().roi_manager.roi_list[3].curve_data

(array([   0,     1,     2, ..., 2995, 2996, 2997]), array([ -207.00168389,  -161.78229208,
↪  -157.62522988, ..., -1017.73174502,
   -1030.27047731, -1042.26989668]))

# Get the tags of an ROI
>>> get_workEnv().roi_manager.roi_list[2].get_all_tags()

{'anatomical_location': 'tail', 'cell_name': 'dcen', 'morphology': 'untagged'}

# Get a single tag
>>> get_workEnv().roi_manager.roi_list[2].get_tag('cell_name')

'dcen'
```

#### 1.51 Stimulus Mapping

##### 1.51.1 ModuleGUI

**class** mesmerize.viewer.modules.stimulus_mapping.**ModuleGUI**(*parent*, *viewer*)

**property maps: dict**

Returns a dictionary of the stimulus maps

##### 1.51.2 Page

**Each Page instance contains the mapping data for one stimulus type**

`class mesmerize.viewer.modules.stimmap_modules.page.Page`(*parent*, *stim_type: str*)

`set_data`(*dataframe: pandas.core.frame.DataFrame*)
Set the stimulus map

**Parameters** `dataframe` – DataFrame with the appropriate rows (see add_row())

`get_dataframe()` → pandas.core.frame.DataFrame
Get the stimulus map as a DataFrame

`set_units`(*units: str*)
Set the time units

**Parameters** `units` – One of 'frames' or 'seconds'

`get_units()` → str
Get the time units

`add_row`(*pd_series: Optional[pandas.core.series.Series] = None*)
Add a row to the stimulus map

**Parameters** `pd_series` – pandas series containing the following: stimulus name, start, end, and color

**Returns**

`delete_row`(*row: Union[mesmerize.viewer.modules.stimmap_modules.row.Row, int]*)
Delete a row from the stimulus map

**Parameters** `row` – The Row object to remove or the numerical index of the row

`clear()`
Clear the stimulus map

##### 1.51.3 DataFrame Format

**Page.set_data() expects a DataFrame in the following format**

**Columns**

| Column | Description |
|---|---|
| name | Stimulus name |
| start | Start time of stimulus period |
| end | End time of stimulus period |
| color | Color to display in the viewer curve plot |

Data types:

| Column | Data type |
|---|---|
| name | str |
| start | numpy.float64 |
| end | numpy.float64 |
| color | Tuple in RGBA format<br><br>(int, int, int, int)<br><br>Each int must be within the 0 - 255 range |

**Example**

| name | start | end | color |
|---|---|---|---|
| control | 0.0 | 328.0 | (0, 75, 0, 255) |
| stim_A | 328.0 | 1156.0 | (0, 0, 125, 255) |
| stim_C | 1156.0 | 2987.0 | (125, 0, 0, 255) |

#### 1.51.4 Example

This example creates a pandas DataFrame from a csv file to set the stimulus mappings. It uses the csv file from the pvc-7 dataset availble on CRCNS: http://dx.doi.org/10.6080/K0C8276G

You can also download the csv here: `stimulus_pvc7.csv`

This example is meant to be run through the *Viewer Script Editor*

```
import pandas as pd
from mesmerize.plotting.utils import get_colormap

# Load dataframe from CSV
df = pd.read_csv('path_to_csv_file')

# Sort according to time
df.sort_values(by='start').reset_index(drop=True, inplace=True)

# Trim off the stimulus periods that are not in the current image sequence
trim = get_image().shape[2]
df = df[df['start'] <= trim]

# get one dataframe for each of the stimulus types
ori_df = df.drop(columns=['sf', 'tf', 'contrast'])  # contains ori stims
sf_df = df.drop(columns=['ori', 'tf', 'contrast'])  # contains sf stims
tf_df = df.drop(columns=['sf', 'ori', 'contrast'])  # contains tf stims

# Rename the stimulus column of interest to "name"
ori_df.rename(columns={'ori': 'name'}, inplace=True)
sf_df.rename(columns={'sf': 'name'}, inplace=True)
tf_df.rename(columns={'tf': 'name'}, inplace=True)


# Get the stimulus mapping module
```

(continues on next page)

(continued from previous page)

```
smm = get_module('stimulus_mapping')

# set the stimulus map in Mesmerize for each of the 3 stimulus types
for stim_type, _df in zip(['ori', 'sf', 'tf'], [ori_df, sf_df, tf_df]):
    # data in the name column must be `str` type for stimulus mapping module
    _df['name'] = _df['name'].apply(str)

    # Get the names of the stimulus periods
    stimuli = _df['name'].unique()
    stimuli.sort()

    # Create colormap with the stimulus names
    stimuli_cmap = get_colormap(stimuli, 'tab10', output='pyqt', alpha=0.6)

    # Create a column with colors that correspond to the stimulus names
    # This is for illustrating the stimulus periods in the viewer plot
    _df['color'] = _df['name'].map(stimuli_cmap)

    # Set the data in the Stimulus Mapping module
    smm.maps[stim_type].set_data(_df)
```

#### 1.52 Data types used for analysis

##### 1.52.1 Transmission

**Inherits from BaseTransmission**

**class** mesmerize.**Transmission**(*df: pandas.core.frame.DataFrame*, *history_trace: mesmerize.analysis.data_types.HistoryTrace*, *proj_path: Optional[str] = None*, *last_output: Optional[str] = None*, *last_unit: Optional[str] = None*, *ROI_DEFS: Optional[list] = None*, *STIM_DEFS: Optional[list] = None*, *CUSTOM_COLUMNS: Optional[list] = None*, *plot_state: Optional[dict] = None*)

The transmission object used throughout the flowchart

**__init__**(*df: pandas.core.frame.DataFrame*, *history_trace: mesmerize.analysis.data_types.HistoryTrace*, *proj_path: Optional[str] = None*, *last_output: Optional[str] = None*, *last_unit: Optional[str] = None*, *ROI_DEFS: Optional[list] = None*, *STIM_DEFS: Optional[list] = None*, *CUSTOM_COLUMNS: Optional[list] = None*, *plot_state: Optional[dict] = None*)

Base class for common Transmission functions

**Parameters**

- **df** (*pd.DataFrame*) – Transmission dataframe
- **history_trace** (HistoryTrace) – HistoryTrace object, keeps track of the nodes & node parameters the transmission has been processed through
- **proj_path** (*str*) – Project path, necessary for the datapoint tracer
- **last_output** (*str*) – Last data column that was appended via a node's operation
- **last_unit** (*str*) – Current units of the data. Refers to the units of column in last_output

- **plot_state** (*dict*) – State of a plot, such as data and label columns. Used when saving interactive plots.

**Variables**

- **df** – DataFrame instance
- **history_trace** – *HistoryTrace instance*
- **last_output** – Name of the DataFrame column that contains data from the most recent node
- **last_unit** – The data units for the data in the column of 'last_output'
- **plot_state** – State of a plot, containing user entered plot parameters. Used for storing interactive plot states.

**static empty_df**(*transmission*, *addCols: Optional[list] = None*) → pandas.core.frame.DataFrame

Just a helper method to return an empty DataFrame with the same columns

**Parameters**

- **transmission** – Transmission object that forms the basis
- **addCols** – list of columns to add

**Returns** The input transmission with an empty dataframe containing the same columns and any additional columns that were passed

**classmethod from_pickle**(*path*)

Load Transmission from a pickle.

**Parameters path** – file path, usually ends in .trn

**to_pickle**(*path: str*)

Save Transmission as a pickle. Not recommended for sharing data, use *to_hdf5()*

**Parameters path** – file path, usually ends in .trn

**classmethod from_hdf5**(*path: str*)

Create Transmission from an hdf5 file. See *HdfTools* for information on the file structure.

**Parameters path** – file path, usually ends in .trn (.ptrn for plots)

**to_hdf5**(*path: str*)

Save as an hdf5 file. Uses pytables to save the DataFrame, serielizes the HistoryTrace using JSON. See *HdfTools*

**Parameters path** – file path, usually ends in .trn

**get_proj_path**() → str

Get the project root dir associated to this Transmission.

**Returns** Root directory of the project

**set_proj_path**(*path: str*)

Set the project root dir for this transmission.

Used for finding associated project files, for example the Datapoint Tracer uses it to find max and std projections of image sequences.

**Parameters path** – Root directory of the project

**to_dict**() → dict

Package Transmission as a dict, useful for saving to hdf5 or pickle

**classmethod from_proj**(*proj_path: str, dataframe: pandas.core.frame.DataFrame, sub_dataframe_name: str = 'root', dataframe_filter_history: Optional[dict] = None*)

**Parameters**

- **proj_path** – root directory of the project
- **dataframe** – Chosen Child DataFrame from the Mesmerize Project
- **sub_dataframe_name** – Name of the sub DataFrame to load
- **dataframe_filter_history** – Filter history of the child dataframe

**static _load_files**(*proj_path: str, row: pandas.core.series.Series*) → pandas.core.series.Series

Loads npz of curve data and pickle files containing metadata using the paths specified in each row of the chosen sub-dataframe of the project

**classmethod merge**(*transmissions: list*)

Merges a list of Transmissions into one transmission. A single DataFrame is created by simple concatenation. HistoryTrace objects are also merged using HistoryTrace.merge.

**Parameters** **transmissions** – A list containing Transmission objects to merge

**Returns** Merged transmission

##### 1.52.2 HistoryTrace

**class mesmerize.analysis.data_types.HistoryTrace**(*history: Optional[Dict[Union[uuid.UUID, str], List[Dict]]] = None, data_blocks: Optional[List[Union[uuid.UUID, str]]] = None*)

Structure of a history trace:

A dict with keys that are the block_ids. Each dict value is a list of operation_dicts. Each operation_dict has a single key which is the name of the operation and the value of that key is the operation parameters.

```
{block_id_1: [
        {operation_1:
            {
              param_1: a,
              param_2: b,
              param_n, z
              }
          },
        {operation_2:
              {
                param_1: a,
                param_n, z
                }
            },
            ...
```

{operation_n:

{

param_n: x

}

}

]

block_id_2: <list of operation dicts>,

…

block_id_n: <list of operation dicts>

}

**The main dict illustrated above should never be worked with directly.**

**You must use the helper methods of this class to query or add information**

**`__init__`**(*history: Optional[Dict[Union[uuid.UUID, str], List[Dict]]] = None*, *data_blocks: Optional[List[Union[uuid.UUID, str]]] = None*)

**Parameters**

- **`history`** – Dict containing a data block UUIDs as keys. The values are a list of dicts containing operation parameters.
- **`data_blocks`** – List of data block UUIDs

**Variables**

- **`_history`** – The dict of the actual data, as illustrated above. Should not be accessed directly. Use the `history` property or call *get_all_data_blocks_history()*.
- **`_data_blocks`** – List of all data blocks. Should not be accessed directly, use the `data_blocks` property instead.

**`property data_blocks: list`**

List of UUIDs that allow you to pin down the history of specific rows of the dataframe to their history as stored in the history trace data structure (self.history)

**`property history: dict`**

The analysis log that is stored in the structure outlined in the doc string

**`create_data_block`**(*dataframe: pandas.core.frame.DataFrame*) → Tuple[pandas.core.frame.DataFrame, uuid.UUID]

Creates a new UUID, assigns it to the input dataframe by setting the UUID in the _BLOCK_ column

**Parameters** **`dataframe`** – Assigns a block ID to this entire DataFrame.

**`_add_data_block`**(*data_block_id: uuid.UUID*)

Adds new datablock UUID to the list of datablocks in this instance. Throws exception if UUID already exists.

**`add_operation`**(*data_block_id: Union[uuid.UUID, str]*, *operation: str*, *parameters: dict*)

Add a single operation, that is usually performed by a node, to the history trace. Added to all or specific datablock(s), depending on which datablock(s) the node performed the operation on

**Parameters**

- **data_block_id** – data_block_id to log the operation on to. either a UUID or 'all' to append the operation to all data blocks
- **operation** – name of the operation, usually the same as the name of the node in all lowercase
- **parameters** – operation parameters.

**get_data_block_history**(*data_block_id: Union[str, uuid.UUID], copy: bool = False*) → List[dict]
Get the full history trace of a single data block.

Use copy=False if you want to modify the history trace of the data block.

**Parameters**

- **data_block_id** (*Union[str, UUID]*) – data block ID
- **copy** (*bool*) – If true, returns a deepcopy

**Returns** data block history

**Return type** List[dict]

**get_all_data_blocks_history**() → dict
Returns history trace of all datablocks

**get_operations_list**(*data_block_id: Union[uuid.UUID, str]*) → list
Returns just a simple list of operations in the order that they were performed on the given datablock. To get the operations along with their parameters call get_data_block_history()

**get_operation_params**(*data_block_id: Union[uuid.UUID, str], operation: str*) → dict
Get the parameters dict for a specific operation that was performed on a specific data block

**check_operation_exists**(*data_block_id: uuid.UUID, operation: str*) → bool
Check if a specific operation was performed on a specific datablock

**static _to_uuid**(*u: Union[str, uuid.UUID]*) → uuid.UUID
If argument 'u' is type <str> that can be formatted as a UUID, return it as UUID type. If argument 'u' is a UUID, just return it.

**to_dict**() → dict
Package the HistoryTrace instance as a dict. Converts all UUIDs to <str> representation for JSON compatibility.

**static from_dict**(*d: dict*) → dict
Format a dict stored using HistoryTrace.to_dict so that it can be used to create a HistoryTrace instance. Converts all the <str> representations of UUID back to <`uuid.UUID`> types.

**Parameters d** – dict containing appropriate 'history' and 'datablocks' keys. Must be packaged by HistoryTrace.to_dict()

**Returns** dict formatted so that it can be used to instantiate a HistoryTrace instance recapitulating the HistoryTrace it was packaged from.

**to_json**(*path: str*)
Save HistoryTrace to a JSON file.

**Parameters path** – file path, usually ends with .json

**classmethod from_json**(*path: str*)
Instantiate HistoryTrace from JSON file (that was saved using HistoryTrace.to_json)

**Parameters path** – file path, usually ends with .json

**to_pickle**(*path: str*)
Dump this instance to a pickle

**Parameters** **path** – file path

**classmethod from_pickle**(*path: str*)
Load HistoryTrace that was pickled

**Parameters** **path** – file path

**classmethod merge**(*history_traces: list*)
Merge a list of HistoryTrace instances into one HistoryTrace instance. Useful when merging Transmission objects.

**Parameters** **history_traces** – list of HistoryTrace instances

**draw_graph**(*data_block_id: Union[str, uuid.UUID]*, ***kwargs*) → str
Draw graph of a data block. kwargs are passed to *mesmerize.common.utils.draw_graph*

**Parameters** **data_block_id** (*Union[str, UUID]*) – data block ID from which to get history to draw in a graph

**Returns** file path to the graph pdf file

**Return type** str

**static clean_history_trace**(*db_history: list*) → list
Cleans up excessive data such as frequencies linspaces and linkage matrices so the graph is viewable.

**Parameters** **db_history** – data block history

**Returns** data block history with excessive params removed

##### 1.52.3 Examples

You can save a Transmission files using the *Save node* and work with the data directly in scripts, jupyter notebooks etc. You can also save them through the flowchart console (and plot consoles) through `Transmission.to_hdf5`.

###### Working with Transmission files

Load a saved Transmission instance using `Transmission.from_hdf5`

```
>>> from mesmerize import Transmission
>>> from uuid import UUID

# load transmission file
>>> t = Transmission.from_hdf5('/share/data/temp/kushal/data.trn')
<mesmerize.analysis.data_types.Transmission at 0x7f4d42f386a0>

# The DataFrame is always the 'df' attribute
>>> t.df.head()

                                           CurvePath  ... FCLUSTER_LABELS
0  curves/a2-_-1-_-843c2d43-75f3-421a-9fef-483d1e...  ...               8
1  curves/brn3b_a6-_-2-_-21557a64-6868-4ff4-8db1-...  ...               4
2  curves/brn3b_a6-_-2-_-21557a64-6868-4ff4-8db1-...  ...               5
3  curves/brn3b_day1_3-_-2-_-ff3e95df-0e15-495c-9...  ...               8
4  curves/brn3b_day1_3-_-2-_-ff3e95df-0e15-495c-9...  ...               6
```

(continues on next page)

(continued from previous page)

```

[5 rows x 27 columns]

# the `df` is just a pandas dataframe
# View a list of samples in the current file
>>> t.df.SampleID.unique()

array(['a2-_-1', 'a5-_-1', 'brn3b_a6-_-2', 'brn3b_day1_3-_-2',
    'brn3b_day1_a1-_-2', 'brn3b_day1_a2-_-2', 'brn3b_day1_a4-_-2',
    'brn3b_day2_a1-_-2', 'brn3b_day2_a1-_-t', 'brn3b_day2_a10-_-2',
    'brn3b_day2_a2-_-1', 'brn3b_day2_a2-_-3', 'brn3b_day2_a8-_-1',
    'cesa_a1-_-1', 'cesa_a1-_-2', 'cesa_a1_jan_2019-_-1',
    'cesa_a1_jan_2019-_-2', 'cesa_a2-_-2', 'cesa_a6-_-1',
    'cesa_a7-_-1', 'cesa_a7-_-2', 'cesa_a8-_-1', 'cesa_a9-_-1',
    'cng_ch4_day1_a2-_-t1', 'cng_ch4_day1_a2-_-t2',
    'cng_ch4_day2_a4-_-t1', 'dmrt1_day1_a2-_-2', 'dmrt1_day1_a4-_-t2',
    'dmrt1_day1_a5-_-', 'dmrt1_day1_a6-_-t', 'dmrt1_day1_a6-_-t2',
    'dmrt1_day2_a1-_-t1', 'dmrt1_day2_a1-_-t2', 'dmrt1_day2_a2-_-t1',
    'dmrt1_day2_a3-_-t1', 'dmrt1_day2_a3-_-t2', 'dmrt1_day2_a4-_-t1',
    'dmrt1_day2_a4-_-t2', 'hnk1_a5-_-2', 'hnk1_a6-_-1', 'hnk1_a7-_-1',
    'hnk1_a7-_-2', 'hnk1_a8-_-1', 'pc2_a10-_-1', 'pc2_a11-_-1',
    'pc2_a13-_-1', 'pc2_a14-_-1', 'pc2_a15-_-1', 'pc2_a16-_-1',
    'pc2_a9-_-1', 'pde9_day1_a2-_-2', 'pde9_day1_a3-_-1',
    'pde9_day1_a4-_-1', 'pde9_day1_a4-_-2', 'pde9_day2_a2-_-t2',
    'pde9_day2_a2-_-t4', 'pde9_day2_a4-_-t1', 'pde9_day2_a4-_-t2',
    'pde9_day2_a4-_-t3', 'pde9_day2_a5-_-t1', 'pde9_day2_a5-_-t2',
    'pde9_day2_a6-_-t1', 'pde9_day2_a7-_-t1', 'pde9_day2_a7-_-t2'],
    dtype=object)

# Show data associated with a single sample
>>> t.df[t.df['SampleID'] == 'brn3b_day1_a1-_-2']

                                            CurvePath  ... FCLUSTER_LABELS
6   curves/brn3b_day1_a1-_-2-_-d3c5f225-7039-4abd-...  ...               6
7   curves/brn3b_day1_a1-_-2-_-d3c5f225-7039-4abd-...  ...               6
8   curves/brn3b_day1_a1-_-2-_-d3c5f225-7039-4abd-...  ...               5
9   curves/brn3b_day1_a1-_-2-_-d3c5f225-7039-4abd-...  ...               7
10  curves/brn3b_day1_a1-_-2-_-d3c5f225-7039-4abd-...  ...               5

# View the data associated with one ROI
# the `uuid_curve` is a unique identifier for each curve/ROI
>> t.df[t.df['SampleID'] == 'brn3b_day1_a1-_-2'].iloc[0]

CurvePath               curves/brn3b_day1_a1-_-2-_-d3c5f225-7039-4abd-...
ImgInfoPath             images/brn3b_day1_a1-_-2-_-d3c5f225-7039-4abd-...
ImgPath                 images/brn3b_day1_a1-_-2-_-d3c5f225-7039-4abd-...
ImgUUID                              d3c5f225-7039-4abd-a7a1-5e9ef2150013
ROI_State               {'roi_xs': [554, 553, 553, 552, 552, 551, 551,...
SampleID                                                brn3b_day1_a1-_-2
anatomical_location                                                  palp
cell_name                                                            palp
comments                                                         untagged
```

(continues on next page)

(continued from previous page)

```
date                                                       20190425_110103
dorso_ventral_axis                                                untagged
misc                                                                    {}
morphology                                                        untagged
promoter                                                             brn3b
rostro_caudal_axis                                                untagged
stimulus_name                                                   [untagged]
uuid_curve                            f44fbd3d-6eaa-4e19-a677-496908565fde
_RAW_CURVE              [81.41972198848178, 75.61356993008134, 70.0493...
meta                    {'origin': 'AwesomeImager', 'version': '4107ff...
stim_maps                                                         [[None]]
_BLOCK_                               3e069e2d-d012-47ee-830c-93d85197e2f4
_SPLICE_ARRAYS          [2.646593459501195, 1.8252819116136887, 1.7422...
_NORMALIZE              [0.0681729940259753, 0.06533186950232853, 0.06...
_RFFT                   [443.19357880089615, -66.8777897472859, 55.244...
_ABSOLUTE_VALUE         [443.19357880089615, 66.8777897472859, 55.2443...
_LOG_TRANSFORM          [2.646593459501195, 1.8252819116136887, 1.7422...
FCLUSTER_LABELS                                                          6
Name: 6, dtype: object

# Show the ROI object data
>>> t.df[t.df['SampleID'] == 'brn3b_day1_a1-_-2'].iloc[0]['ROI_State']

{'roi_xs': array([554, 553, 553, 552, 552, 551, 551, 551, 551, 550, 550, 550, 549,
    548, 547, 547, 546, 546, 545, 545, 544, 543, 543, 542, 541, 541,
    540, 540, 539, 539, 538, 537, 536, 535, 534, 533, 532, 531, 531,
    530, 529, 528, 527, 527, 526, 526, 525, 525, 525, 524, 524, 523,
    522, 522, 521, 521, 520, 521, 521, 521, 521, 521, 522, 522, 522,
    522, 522, 522, 522, 522, 521, 521, 521, 521, 521, 521, 522, 523,
    524, 524, 525, 525, 525, 526, 526, 527, 528, 528, 529, 529, 529,
    530, 530, 531, 532, 532, 533, 534, 535, 535, 536, 536, 537, 538,
    539, 540, 540, 541, 541, 542, 542, 543, 544, 545, 546, 546, 547,
    548, 548, 549, 549, 549, 549, 550, 550, 550, 550, 551, 551, 551,
    552, 552, 552, 553, 553, 553, 554, 554, 554, 553, 554, 554, 554,
    554, 554]),
'roi_ys': array([155, 156, 156, 157, 157, 158, 159, 160, 160, 161, 162, 162, 162,
        162, 163, 163, 164, 164, 165, 165, 165, 166, 166, 166, 167, 167,
        167, 166, 167, 167, 167, 167, 167, 167, 167, 167, 167, 168, 168,
        168, 168, 168, 168, 167, 167, 166, 166, 165, 164, 164, 163, 163,
        163, 162, 162, 161, 161, 160, 160, 159, 158, 157, 156, 156, 155,
        154, 153, 152, 151, 150, 150, 149, 148, 147, 146, 145, 144, 144,
        144, 144, 143, 143, 142, 141, 141, 140, 140, 140, 139, 139, 138,
        137, 137, 136, 136, 136, 135, 135, 135, 136, 136, 137, 137, 137,
        137, 137, 138, 138, 138, 137, 137, 136, 136, 136, 136, 137, 137,
        137, 138, 138, 139, 140, 141, 141, 142, 143, 144, 144, 145, 146,
        146, 147, 148, 148, 149, 150, 150, 151, 151, 152, 152, 153, 154,
        155, 155]),
'curve_data': (array([   0,    1,    2, ..., 2996, 2997, 2998]),
array([ 81.41972199,  75.61356993,  70.04934883, ..., 195.4416283 ,
        184.8844155 , 174.76708104])),
'tags': {'anatomical_location': 'palp',
'cell_name': 'palp',
```

(continues on next page)

(continued from previous page)

```
'morphology': 'untagged'},
'roi_type': 'CNMFROI',
'cnmf_idx': 2}
```

##### View History Log

Transmissions have a *history_trace* attribute which is an instance of *HistoryTrace*.

Use the *get_data_block_history* and *get_operations_list* methods to view the history log of a data block.

```
# To view the history log, first get the block UUID of the dataframe row of which you want the history log

# Block UUIDs are stored in the _BLOCK_ column
>>> bid = t.df.iloc[10]._BLOCK_
>>> bid

'248a6ece-e60e-4a09-845e-188a5199d262'

# Get the history log of this data block
# HistoryTrace.get_operations_list() returns a list of operations, without parameters
# HistoryTrace.get_data_block_history() returns the operations list with the parameters
>>> t.history_trace.get_operations_list(bid)

['spawn_transmission',
 'splice_arrays',
 'normalize',
 'rfft',
 'absolute_value',
 'log_transform',
 'splice_arrays',
 'fcluster']

# View the entire history log with all params
>>> t.history_trace.get_data_block_history(bid)

[{'spawn_transmission': {'sub_dataframe_name': 'neuronal',
'dataframe_filter_history': {'dataframe_filter_history': ['df[~df["promoter"].isin([\'cesa\', \'hnk1\'])]',
    'df[~df["promoter"].isin([\'cesa\', \'hnk1\'])]',
    'df[~df["cell_name"].isin([\'not_a_neuron\', \'non_neuronal\', \'untagged\', \'ependymal\'])]']}}},
{'splice_arrays': {'data_column': '_RAW_CURVE',
'start_ix': 0,
'end_ix': 2990,
'units': 'time'}},
{'normalize': {'data_column': '_SPLICE_ARRAYS', 'units': 'time'}},
{'rfft': {'data_column': '_NORMALIZE',
'frequencies': [0.0,
    0.0033444816053511705,
    0.0033444816053511705,
```

(continues on next page)

(continued from previous page)

```
    0.006688963210702341,
    ...

# Get the parameters for the 'fcluster' operation
>>> fp = t.history_trace.get_operation_params(bid, 'fcluster')

# remove the linkage matrix first so we can view the other params
>>> fp.pop('linkage_matrix');fp

{'threshold': 8.0,
 'criterion': 'maxclust',
 'depth': 1,
 'linkage_params': {'method': 'complete',
 'metric': 'wasserstein',
 'optimal_ordering': True}}

# Draw the analysis history as a graph
# This will open your defeault pdf viewer with the graph
>>> t.history_trace.draw_graph(bid, view=True)

# If you are using the API to perform analysis on
# transmission files, you can use the `HistoryTrace`
# to log the analysis history
# For example, add a number `3.14` to all datapoints in a curve
>>> t.df['_RAW_CURVE'] = t.df['_RAW_CURVE'].apply(lambda x: x + 3.14)

# Append the analysis log
>>> t.history_trace.add_operation(data_block_id='all', operation='addition', parameters={
↪'value': 3.14}
```

#### 1.53 Analysis

Analysis helper functions

##### 1.53.1 Utils

`mesmerize.analysis.utils.`**`get_array_size`**(*transmission: mesmerize.analysis.data_types.Transmission*, *data_column: str*) → int

Returns the size of the 1D arrays in the specified data column. Throws an exception if they do not match

**Parameters**

- **transmission** (`Transmission`) – Desired Transmission
- **data_column** (*str*) – Data column of the Transmission from which to retrieve the size

**Returns** Size of the 1D arrays of the specified data column

**Return type** int

`mesmerize.analysis.utils.`**`get_frequency_linspace`**(*transmission:* mesmerize.analysis.data_types.Transmission) → Tuple[numpy.ndarray, float]

Get the frequency linspace.

Throwns an exception if all datablocks do not have the same linspace & Nyquist frequencies

**Parameters** **`transmission`** – Transmission containing data from which to get frequency linspace

**Returns** tuple: (frequency linspace as a 1D numpy array, nyquist frequency)

**Return type** Tuple[np.ndarray, float]

`mesmerize.analysis.utils.`**`get_proportions`**(*xs: Union[pandas.core.series.Series, numpy.ndarray, list]*, *ys: Union[pandas.core.series.Series, numpy.ndarray]*, *xs_name: str = 'xs'*, *ys_name: str = 'ys'*, *swap: bool = False*, *percentages: bool = True*) → pandas.core.frame.DataFrame

Get the proportions of xs vs ys.

xs & ys are categorical data.

**Parameters**

- **`xs`** (*`Union[pd.Series, np.ndarray]`*) – data plotted on the x axis
- **`ys`** (*`Union[pd.Series, np.ndarray]`*) – proportions of unique elements in ys are calculated per xs
- **`xs_name`** (*`str`*) – name for the xs data, useful for labeling the axis in plots
- **`ys_name`** (*`str`*) – name for the ys data, useful for labeling the axis in plots
- **`swap`** (*`bool`*) – swap x and y

**Returns** DataFrame that can be plotted in a proportions bar graph

**Return type** pd.DataFrame

`mesmerize.analysis.utils.`**`get_sampling_rate`**(*transmission:* mesmerize.analysis.data_types.Transmission, *tolerance: Optional[float] = 0.1*) → float

Returns the mean sampling rate of all data in a Transmission if it is within the specified tolerance. Otherwise throws an exception.

**Parameters**

- **`transmission`** (`Transmission`) – Transmission object of the data from which sampling rate is obtained.
- **`tolerance`** (*`float`*) – Maximum tolerance (in Hertz) of sampling rate variation between different samples

**Returns** The mean sampling rate of all data in the Transmission

**Return type** float

`mesmerize.analysis.utils.`**`organize_dataframe_columns`**(*columns: Iterable[str]*) → Tuple[List[str], List[str], List[str]]

Organizes DataFrame columns into data column, categorical label columns, and uuid columns.

**Parameters** **`columns`** – All DataFrame columns

**Returns** (data_columns, categorical_columns, uuid_columns)

**Return type** Tuple[List[str], List[str], List[str]]

`mesmerize.analysis.utils.`**`pad_arrays`**(*a: numpy.ndarray*, *method: str = 'random'*, *output_size: Optional[int] = None*, *mode: str = 'minimum'*, *constant: Optional[Any] = None*) → numpy.ndarray

Pad all the input arrays so that are of the same length. The length is determined by the largest input array. The padding value for each input array is the minimum value in that array.

Padding for each input array is either done after the array's last index to fill up to the length of the largest input array (method 'fill-size') or the padding is randomly flanked to the input array (method 'random') for easier visualization.

**Parameters**

- **a** (*np.ndarray*) – 1D array where each element is a 1D array
- **method** (*str*) – one of 'fill-size' or 'random', see docstring for details
- **output_size** – not used
- **mode** (*str*) – one of either 'constant' or 'minimum'. If 'minimum' the min value of the array is used as the padding value. If 'constant' the values passed to the "constant" argument is used as the padding value.
- **constant** (*Any*) – padding value if 'mode' is set to 'constant'

**Returns** Arrays padded according to the chosen method. 2D array of shape [n_arrays, size of largest input array]

**Return type** np.ndarray

##### 1.53.2 Cross correlation

###### functions

**Helper functions. Uses tslearn.cycc**

`mesmerize.analysis.math.cross_correlation.`**`ncc_c`**(*x: numpy.ndarray*, *y: numpy.ndarray*) → numpy.ndarray

Must pass 1D array to both x and y

**Parameters**

- **x** – Input array [x1, x2, x3, … xn]
- **y** – Input array [y2, y2, x3, … yn]

**Returns** Returns the normalized cross correlation function (as an array) of the two input vector arguments "x" and "y"

**Return type** np.ndarray

`mesmerize.analysis.math.cross_correlation.`**`get_omega`**(*x: Optional[numpy.ndarray] = None*, *y: Optional[numpy.ndarray] = None*, *cc: Optional[numpy.ndarray] = None*) → int

Must pass a 1D array to either both "x" and "y" or a cross-correlation function (as an array) to "cc"

**Parameters**

- **x** – Input array [x1, x2, x3, … xn]
- **y** – Input array [y2, y2, x3, … yn]
- **cc** – cross-correlation function represented as an array [c1, c2, c3, … cn]

**Returns** index (x-axis position) of the global maxima of the cross-correlation function

**Return type** np.ndarray

`mesmerize.analysis.math.cross_correlation.`**`get_lag`**(*x: Optional[numpy.ndarray] = None*, *y: Optional[numpy.ndarray] = None*, *cc: Optional[numpy.ndarray] = None*) → float

Must pass a 1D array to either both "x" and "y" or a cross-correlation function (as an array) to "cc"

**Parameters**

- **x** – Input array [x1, x2, x3, … xn]
- **y** – Input array [y2, y2, x3, … yn]
- **cc** – cross-correlation function represented as a array [c1, c2, c3, … cn]

**Returns** Position of the maxima of the cross-correlation function with respect to middle point of the cross-correlation function

**Return type** np.ndarray

`mesmerize.analysis.math.cross_correlation.`**`get_epsilon`**(*x: Optional[numpy.ndarray] = None*, *y: Optional[numpy.ndarray] = None*, *cc: Optional[numpy.ndarray] = None*) → float

Must pass a 1D vector to either both "x" and "y" or a cross-correlation function to "cc"

**Parameters**

- **x** – Input array [x1, x2, x3, … xn]
- **y** – Input array [y2, y2, x3, … yn]
- **cc** – cross-correlation function represented as an array [c1, c2, c3, … cn]

**Returns** Magnitude of the global maxima of the cross-correlationn function

**Return type** np.ndarray

`mesmerize.analysis.math.cross_correlation.`**`get_lag_matrix`**(*curves: Optional[numpy.ndarray] = None*, *ccs: Optional[numpy.ndarray] = None*) → numpy.ndarray

Get a 2D matrix of lags. Can pass either a 2D array of 1D curves or cross-correlations

**Parameters**

- **curves** – 2D array of 1D curves
- **ccs** – 2D array of 1D cross-correlation functions represented by arrays

**Returns** 2D matrix of lag values, shape is [n_curves, n_curves]

**Return type** np.ndarray

`mesmerize.analysis.math.cross_correlation.`**`get_epsilon_matrix`**(*curves: Optional[numpy.ndarray] = None*, *ccs: Optional[numpy.ndarray] = None*) → numpy.ndarray

Get a 2D matrix of maximas. Can pass either a 2D array of 1D curves or cross-correlations

**Parameters**

- **curves** – 2D array of 1D curves
- **ccs** – 2D array of 1D cross-correlation functions represented by arrays

**Returns** 2D matrix of maxima values, shape is [n_curves, n_curves]

**Return type** np.ndarray

`mesmerize.analysis.math.cross_correlation.`**`compute_cc_data`**(*curves: numpy.ndarray*) → *mesmerize.analysis.math.cross_correlation.CC_Data*

Compute cross-correlation data (cc functions, lag and maxima matrices)

**Parameters** **`curves`** – input curves as a 2D array, shape is [n_samples, curve_size]

**Returns** cross correlation data for the input curves as a CC_Data instance

**Return type** *CC_Data*

`mesmerize.analysis.math.cross_correlation.`**`compute_ccs`**(*a: numpy.ndarray*) → numpy.ndarray

Compute cross-correlations between all 1D curves in a 2D input array

**Parameters** **`a`** – 2D input array of 1D curves, shape is [n_samples, curve_size]

**Return type** np.ndarray

#### CC_Data

##### Data container

> **Warning:** All arguments MUST be numpy.ndarray type for CC_Data for the save to be saveable as an hdf5 file. Set `numpy.unicode` as the **dtype** for the `curve_uuids` and `labels` arrays. If the **dtype** is `'O'` (object) the **to_hdf5()** method will fail.

**`class`** `mesmerize.analysis.cross_correlation.`**`CC_Data`**(*input_data: Optional[numpy.ndarray] = None, ccs: Optional[numpy.ndarray] = None, lag_matrix: Optional[numpy.ndarray] = None, epsilon_matrix: Optional[numpy.ndarray] = None, curve_uuids: Optional[numpy.ndarray] = None, labels: Optional[numpy.ndarray] = None*)

**`__init__`**(*input_data: Optional[numpy.ndarray] = None, ccs: Optional[numpy.ndarray] = None, lag_matrix: Optional[numpy.ndarray] = None, epsilon_matrix: Optional[numpy.ndarray] = None, curve_uuids: Optional[numpy.ndarray] = None, labels: Optional[numpy.ndarray] = None*)

Object for organizing cross-correlation data

types must be numpy.ndarray to be compatible with hdf5

**Parameters**

- **`ccs`** (*`np.ndarray`*) – array of cross-correlation functions, shape: [n_curves, n_curves, func_length]
- **`lag_matrix`** (*`np.ndarray`*) – the lag matrix, shape: [n_curves, n_curves]
- **`epsilon_matrix`** (*`np.ndarray`*) – the maxima matrix, shape: [n_curves, n_curves]
- **`curve_uuids`** (*`np.ndarray`*) – uuids (str representation) for each of the curves, length: n_curves
- **`labels`** (*`np.ndarray`*) – labels for each curve, length: n_curves

**`ccs`**

array of cross-correlation functions

**`lag_matrix`**

lag matrix

**epsilon_matrix**
maxima matrix

**curve_uuids**
uuids for each curve

**labels**
labels for each curve

**get_threshold_matrix**(*matrix_type: str*, *lag_thr: float*, *max_thr: float*, *lag_thr_abs: bool = True*) → numpy.ndarray
Get lag or maxima matrix with thresholds applied. Values outside the threshold are set to NaN

**Parameters**

- **matrix_type** – one of 'lag' or 'maxima'
- **lag_thr** – lag threshold
- **max_thr** – maxima threshold
- **lag_thr_abs** – threshold with the absolute value of lag

**Returns** the requested matrix with the thresholds applied to it.

**Return type** np.ndarray

**classmethod from_dict**(*d: dict*)
Load data from a dict

**to_hdf5**(*path: str*)
Save as an HDF5 file

**Parameters path** – path to save the hdf5 file to, file must not exist.

**classmethod from_hdf5**(*path: str*)
Load cross-correlation data from an hdf5 file

**Parameters path** – path to the hdf5 file

##### 1.53.3 Clustering metrics

mesmerize.analysis.clustering_metrics.**get_centerlike**(*cluster_members: numpy.ndarray*, *metric: Optional[Union[str, callable]] = None*, *dist_matrix: Optional[numpy.ndarray] = None*) → Tuple[numpy.ndarray, int]
Finds the 1D time-series within a cluster that is the most centerlike

**Parameters**

- **cluster_members** – 2D numpy array in the form [n_samples, 1D time_series]
- **metric** – Metric to use for pairwise distance calculation, simply passed to sklearn.metrics.pairwise_distances
- **dist_matrix** – Distance matrix of the cluster members

**Returns** The cluster member which is most centerlike, and its index in the cluster_members array mesmerize.analysis.clustering_metrics.**get_cluster_radius**(*cluster_members: numpy.ndarray*, *metric: Optional[Union[str, callable]] = None*, *dist_matrix: Optional[numpy.ndarray] = None*, *centerlike_index: Optional[int] = None*) → float

Returns the cluster radius according to chosen distance metric

**Parameters**

- **cluster_members** – 2D numpy array in the form [n_samples, 1D time_series]
- **metric** – Metric to use for pairwise distance calculation, simply passed to sklearn.metrics.pairwise_distances
- **dist_matrix** – Distance matrix of the cluster members
- **centerlike_index** – Index of the centerlike cluster member within the cluster_members array

**Returns** The cluster radius, average between the most centerlike member and all other members

mesmerize.analysis.clustering_metrics.**davies_bouldin_score**(*data: numpy.ndarray*, *cluster_labels: numpy.ndarray*, *metric: Union[str, callable]*) → Tuple[float, numpy.ndarray]

Adopted from sklearn.metrics.davies_bouldin_score to use any distance metric

**Parameters**

- **data** – Data that was used for clustering, [n_samples, 1D time_series]
- **metric** – Metric to use for pairwise distance calculation, simply passed to sklearn.metrics.pairwise_distances
- **cluster_labels** – Cluster labels

**Returns** Davies Bouldin Score using EMD

#### 1.54 Nodes

##### 1.54.1 Data

**class** mesmerize.pyqtgraphCore.flowchart.library.Data.**DropNa**(**args*, ***kwargs*)

Drop NaNs from the DataFrame

**class** mesmerize.pyqtgraphCore.flowchart.library.Data.**LoadFile**(*name*)

Load Transmission data object from pickled file

**class** mesmerize.pyqtgraphCore.flowchart.library.Data.**LoadProjDF**(*name*)

Load raw project DataFrames as Transmission

**class** mesmerize.pyqtgraphCore.flowchart.library.Data.**Merge**(*name*)

Merge transmissions

**class** mesmerize.pyqtgraphCore.flowchart.library.Data.**NormRaw**(*name*, *ui=None*, *terminals=None*, ***kwargs*)

Normalize between raw min and max values.

**class** mesmerize.pyqtgraphCore.flowchart.library.Data.**PadArrays**(*name*, *ui=None*, *terminals=None*, ***kwargs*)

Pad 1-D numpy arrays in a particular column

**class** mesmerize.pyqtgraphCore.flowchart.library.Data.**Save**(*name*)
Save Transmission data object

**class** mesmerize.pyqtgraphCore.flowchart.library.Data.**SelectColumns**(*name*, *ui=None*, *terminals=None*, ***kwargs*)

**class** mesmerize.pyqtgraphCore.flowchart.library.Data.**SelectRows**(*name*, *ui=None*, *terminals=None*, ***kwargs*)

**class** mesmerize.pyqtgraphCore.flowchart.library.Data.**SpliceArrays**(*name*, *ui=None*, *terminals=None*, ***kwargs*)
Splice 1-D numpy arrays in a particular column.

**class** mesmerize.pyqtgraphCore.flowchart.library.Data.**TextFilter**(*name*, *ui=None*, *terminals=None*, ***kwargs*)
Simple string filtering in a specified column

**class** mesmerize.pyqtgraphCore.flowchart.library.Data.**ViewHistory**(**args*, ***kwargs*)
View History Trace of the input Transmission

**class** mesmerize.pyqtgraphCore.flowchart.library.Data.**ViewTransmission**(*name*)
View transmission using the spyder object editor

**class** mesmerize.pyqtgraphCore.flowchart.library.Data.**iloc**(*name*, *ui=None*, *terminals=None*, ***kwargs*)
Pass only one or multiple DataFrame Indices

##### 1.54.2 Display

**class** mesmerize.pyqtgraphCore.flowchart.library.Display.**AnalysisGraph**(*name*)
Graph of the analysis log

**class** mesmerize.pyqtgraphCore.flowchart.library.Display.**BeeswarmPlots**(*name*)
Beeswarm and Violin plots

**class** mesmerize.pyqtgraphCore.flowchart.library.Display.**CrossCorr**(*name*)
Cross Correlation

**class** mesmerize.pyqtgraphCore.flowchart.library.Display.**FrequencyDomainMagnitude**(*name*, *ui=None*, *terminals=None*, ***kwargs*)
Plot Frequency vs. Frequency Domain Magnitude

**class** mesmerize.pyqtgraphCore.flowchart.library.Display.**Heatmap**(*name*)
Stack 1-D arrays and plot visually like a heatmap

**class** mesmerize.pyqtgraphCore.flowchart.library.Display.**Plot**(*name*)
Plot curves and/or scatter points

**class** mesmerize.pyqtgraphCore.flowchart.library.Display.**Proportions**(*name*)
Plot proportions of one categorical column vs another

**class** mesmerize.pyqtgraphCore.flowchart.library.Display.**ScatterPlot**(*name*)
Scatter Plot, useful for visualizing transformed data and clusters

**class** mesmerize.pyqtgraphCore.flowchart.library.Display.**SpaceMap**(*name*)
Visualize spatial maps of a categorical variable

**class** mesmerize.pyqtgraphCore.flowchart.library.Display.**Timeseries**(*name*)

Seaborn Timeseries plot

##### 1.54.3 Signal

**class** mesmerize.pyqtgraphCore.flowchart.library.Signal.**ButterWorth**(*name*, *ui=None*, *terminals=None*, ***kwargs*)

Butterworth Filter

**class** mesmerize.pyqtgraphCore.flowchart.library.Signal.**Normalize**(*name*, *ui=None*, *terminals=None*, ***kwargs*)

Normalize a column containing 1-D arrays such that values in each array are normalized between 0 and 1

Output Column -> Input Column

**class** mesmerize.pyqtgraphCore.flowchart.library.Signal.**PeakDetect**(*name*, ***kwargs*)

Detect peaks & bases by finding local maxima & minima. Use this after the Derivative Filter

**class** mesmerize.pyqtgraphCore.flowchart.library.Signal.**PeakFeatures**(**args*, ***kwargs*)

Extract peak features after peak detection

**class** mesmerize.pyqtgraphCore.flowchart.library.Signal.**PowerSpectralDensity**(*name*, *ui=None*, *terminals=None*, ***kwargs*)

Return the Power Spectral Density of a curve.

**class** mesmerize.pyqtgraphCore.flowchart.library.Signal.**RFFT**(*name*, *ui=None*, *terminals=None*, ***kwargs*)

Uses fftpack.rfft, 'Discrete Fourier transform of a real sequence.

https://docs.scipy.org/doc/scipy/reference/generated/scipy.fftpack.rfft.html#scipy.fftpack.rfft

**class** mesmerize.pyqtgraphCore.flowchart.library.Signal.**Resample**(*name*, *ui=None*, *terminals=None*, ***kwargs*)

Resample 1D data, uses scipy.signal.resample. "Rs" is the new sampling rate in "Tu" units of time. If "Tu" = 1, then Rs is the new sampling rate in Hertz.

**class** mesmerize.pyqtgraphCore.flowchart.library.Signal.**SavitzkyGolay**(*name*, *ui=None*, *terminals=None*, ***kwargs*)

Savitzky-Golay filter.

**class** mesmerize.pyqtgraphCore.flowchart.library.Signal.**ScalerMeanVariance**(*name*, *ui=None*, *terminals=None*, ***kwargs*)

Scaler for time series. Scales time series so that their mean (resp. standard deviation) in each dimension is mu (resp. std).

See https://tslearn.readthedocs.io/en/latest/gen_modules/preprocessing/tslearn.preprocessing.TimeSeriesScalerMeanVariance.html#tslearn.preprocessing.TimeSeriesScalerMeanVariance

**class** mesmerize.pyqtgraphCore.flowchart.library.Signal.**SigmaMAD**(*name*, *ui=None*, *terminals=None*, ***kwargs*)

**class** mesmerize.pyqtgraphCore.flowchart.library.Signal.**iRFFT**(*name*, *ui=None*, *terminals=None*, ***kwargs*)

Uses fftpack.irfft, 'Return inverse discrete Fourier transform of real sequence.'

https://docs.scipy.org/doc/scipy/reference/generated/scipy.fftpack.irfft.html#scipy.fftpack.irfft

Input must have an _RFFT column from the RFFT node.

##### 1.54.4 Math

class mesmerize.pyqtgraphCore.flowchart.library.Math.**AbsoluteValue**(*name*, *ui=None*, *terminals=None*, ***kwargs*)

Performs numpy.abs(<input>). Returns root-mean-square value if <input> is complex

class mesmerize.pyqtgraphCore.flowchart.library.Math.**ArgGroupStat**(*name*, *ui=None*, *terminals=None*, ***kwargs*)

Group by a certain column and return value of another column based on a data column statistic

class mesmerize.pyqtgraphCore.flowchart.library.Math.**ArrayStats**(*name*, *ui=None*, *terminals=None*, ***kwargs*)

Perform various statistical functions

class mesmerize.pyqtgraphCore.flowchart.library.Math.**Derivative**(*name*, *ui=None*, *terminals=None*, ***kwargs*)

Return the Derivative of a curve.

class mesmerize.pyqtgraphCore.flowchart.library.Math.**Integrate**(*name*, *ui=None*, *terminals=None*, ***kwargs*)

class mesmerize.pyqtgraphCore.flowchart.library.Math.**LinRegress**(*name*, *ui=None*, *terminals=None*, ***kwargs*)

Linear Regression

class mesmerize.pyqtgraphCore.flowchart.library.Math.**LogTransform**(*name*, *ui=None*, *terminals=None*, ***kwargs*)

Can perform various log transforms

class mesmerize.pyqtgraphCore.flowchart.library.Math.**TVDiff**(*name*, *ui=None*, *terminals=None*, ***kwargs*)

Total Variation Regularized Numerical Differentiation, Chartrand 2011 method

class mesmerize.pyqtgraphCore.flowchart.library.Math.**XpowerY**(*name*, *ui=None*, *terminals=None*, ***kwargs*)

Raise each element of arrays in data column to the exponent Y

class mesmerize.pyqtgraphCore.flowchart.library.Math.**ZScore**(*name*, *ui=None*, *terminals=None*, ***kwargs*)

Z-Score the input data. Uses scipy.stats.zscore. Computes over sub-DataFrames that are created according to the "group_by" column parameter

##### 1.54.5 Biology

class mesmerize.pyqtgraphCore.flowchart.library.Biology.**ExtractStim**(*name*, *ui=None*, *terminals=None*, ***kwargs*)

Extract portions of curves according to stimulus maps

class mesmerize.pyqtgraphCore.flowchart.library.Biology.**ManualDFoF**(*name*, *ui=None*, *terminals=None*, ***kwargs*)

Set Fo for dF/Fo using a particular time period. Useful for looking at stimulus responses

class mesmerize.pyqtgraphCore.flowchart.library.Biology.**StaticDFoFo**(*name*, *ui=None*, *terminals=None*, ***kwargs*)

Perform (F - Fo / Fo) without a rolling window to find Fo

**class** mesmerize.pyqtgraphCore.flowchart.library.Biology.**StimTuning**(*name*)
Get stimulus tuning curves

##### 1.54.6 Clustering

**class** mesmerize.pyqtgraphCore.flowchart.library.Clustering.**KMeans**(*name*, *ui=None*, *terminals=None*, ***kwargs*)
KMeans clustering https://scikit-learn.org/stable/modules/generated/sklearn.cluster.KMeans.html
Output column -> KMEANS_CLUSTER_<data_column>

**class** mesmerize.pyqtgraphCore.flowchart.library.Clustering.**KShape**(*name*)
k-Shape clustering

##### 1.54.7 Hierarchical

**class** mesmerize.pyqtgraphCore.flowchart.library.Hierarchical.**FCluster**(*name*, ***kwargs*)
Basically scipy.cluster.hierarchy.fcluster. Form flat clusters from the hierarchical clustering defined by the given linkage matrix.

**class** mesmerize.pyqtgraphCore.flowchart.library.Hierarchical.**Inconsistent**(*name*)
Calculate inconsistency statistics on a linkage matrix. Returns inconsistency matrix

**class** mesmerize.pyqtgraphCore.flowchart.library.Hierarchical.**Linkage**(*name*, *ui=None*, *terminals=None*, ***kwargs*)
Basically scipy.cluster.hierarchy.linkage Compute a linkage matrix for Hierarchical clustering

**class** mesmerize.pyqtgraphCore.flowchart.library.Hierarchical.**MaxIncStat**(*name*, ***kwargs*)
Return the maximum statistic for each non-singleton cluster and its children.

**class** mesmerize.pyqtgraphCore.flowchart.library.Hierarchical.**MaxInconsistent**(*name*, ***kwargs*)
Return the maximum inconsistency coefficient for each non-singleton cluster and its children.

##### 1.54.8 Transform

**class** mesmerize.pyqtgraphCore.flowchart.library.Transform.**LDA**(*name*, ***kwargs*)
Linear Discriminant Analysis, uses sklearn

**class** mesmerize.pyqtgraphCore.flowchart.library.Transform.**Manifold**(*name*, *ui=None*, *terminals=None*, ***kwargs*)
Manifold learning

**class** mesmerize.pyqtgraphCore.flowchart.library.Transform.**NeuralDynamics**(*name*)
Dimensionality reduction of neuronal dynamics

**class** mesmerize.pyqtgraphCore.flowchart.library.Transform.**PCA**(*name*, *ui=None*, *terminals=None*, ***kwargs*)
Principal component analysis. Uses sklearn.decomposition.PCA

##### 1.54.9 CtrlNode

Base for all nodes

**class** `mesmerize.pyqtgraphCore.flowchart.library.common.CtrlNode`(*name*, *ui=None*, *terminals=None*, ***kwargs*)

Abstract class for nodes with auto-generated control UI

**ctrlWidget**()

Return this Node's control widget.

By default, Nodes have no control widget. Subclasses may reimplement this method to provide a custom widget. This method is called by Flowcharts when they are constructing their Node list.

**process**(***kwargs*)

Process data through this node. This method is called any time the flowchart wants the node to process data. It will be called with one keyword argument corresponding to each input terminal, and must return a dict mapping the name of each output terminal to its new value.

This method is also called with a 'display' keyword argument, which indicates whether the node should update its display (if it implements any) while processing this data. This is primarily used to disable expensive display operations during batch processing.

**saveState**()

Return a dictionary representing the current state of this node (excluding input / output values). This is used for saving/reloading flowcharts. The default implementation returns this Node's position, bypass state, and information about each of its terminals.

Subclasses may want to extend this method, adding extra keys to the returned dict.

**restoreState**(*state*)

Restore the state of this node from a structure previously generated by saveState().

#### 1.55 Plotting utils

A few useful helper functions

`mesmerize.plotting.utils.`**get_colormap**(*labels: iter*, *cmap: str*, ***kwargs*) → collections.OrderedDict

Get a dict for mapping labels onto colors

Any kwargs are passed to auto_colormap()

**Parameters**

- **labels** – labels for creating a colormap. Order is maintained if it is a list of unique elements.
- **cmap** – name of colormap

**Returns** dict of labels as keys and colors as values

`mesmerize.plotting.utils.`**map_labels_to_colors**(*labels: iter*, *cmap: str*, ***kwargs*) → list

Map labels onto colors according to chosen colormap

Any kwargs are passed to auto_colormap()

**Parameters**

- **labels** – labels for mapping onto a colormap
- **cmap** – name of colormap

**Returns** list of colors mapped onto the labels

mesmerize.plotting.utils.**auto_colormap**(*n_colors: int*, *cmap: str = 'hsv'*, *output: str = 'mpl'*, *spacing: str = 'uniform'*, *alpha: float = 1.0*) → List[Union[PyQt5.QtGui.QColor, numpy.ndarray, str]]

If non-qualitative map: returns list of colors evenly spread through the chosen colormap. If qualitative map: returns subsequent colors from the chosen colormap

**Parameters**

- **n_colors** – Numbers of colors to return
- **cmap** – name of colormap
- **output** – option: 'mpl' returns RGBA values between 0-1 which matplotlib likes, option: 'pyqt' returns QtGui.QColor instances that correspond to the RGBA values option: 'bokeh' returns hex strings that correspond to the RGBA values which bokeh likes
- **spacing** – option: 'uniform' returns evenly spaced colors across the entire cmap range option: 'subsequent' returns subsequent colors from the cmap
- **alpha** – alpha level, 0.0 - 1.0

**Returns** List of colors as either `QColor`, `numpy.ndarray`, or hex `str` with length `n_colors`

class mesmerize.plotting.utils.**WidgetRegistry**

Register widgets to conveniently store and restore their states

**register**(*widget: PyQt5.QtWidgets.QWidget*, *setter: callable*, *getter: callable*, *name: str*)

Register a widget. The *setter* and *getter* methods must be interoperable

**Parameters**

- **widget** (*QtWidgets.QWidget*) – widget to register
- **setter** (*callable*) – widget's method to use for setting its value
- **getter** (*callable*) – widget's method to use for getting its value. This value must be settable through the specified "setter" method
- **name** (*str*) – a name for this widget

**get_state**() → dict

Get a dict of states for all registered widgets

**set_state**(*state: dict*)

Set all registered widgets from a dict

### 1.56 Plot Bases

#### 1.56.1 AbstractBasePlotWidget

class mesmerize.plotting.widgets.base.**_AbstractBasePlotWidget**

**abstract property transmission: mesmerize.analysis.data_types.Transmission**

The input transmission

**Return type** *Transmission*

**abstract set_input**(*transmission: mesmerize.analysis.data_types.Transmission*)

Set the input Transmission with data to plot

**Parameters** **transmission** – Input transmission

**abstract update_plot**(**args*, ***kwargs*)
Method that must must be used for updating the plot

**abstract get_plot_opts**() → dict
Package all necessary plot parameters that in combination with the transmission property are sufficient to restore the plot

**abstract set_plot_opts**(*opts: dict*)
Set plot parameters from a dict in the format returned by get_plot_opts()

**abstract save_plot**(**args*)
Package plot data and plot parameters and save to a file. Must contain all the information that is necessary to restore the plot

**abstract open_plot**(*ptrn_path: str*, *proj_path: str*) → Optional[Tuple[str, str]]
Open a plot file and restore the plot

##### 1.56.2 BasePlotWidget

Inherit from this to create interactive plots that can be saved and restored.

**class** mesmerize.plotting.widgets.base.**BasePlotWidget**
Bases: `mesmerize.plotting.widgets.base._AbstractBasePlotWidget`

Base for plot widgets.

Subclasses must define the class attribute "drop_opts" which is used to store a list of JSON incompatible keys returned by the get_plot_opts() method

**__init__**()
Initialize self. See help(type(self)) for accurate signature.

**block_signals_list**
List of QObjects included in dynamic signal blocking. Used for storing plot parameter widgets so that changing all of them quickly (like when restoring a plot) doesn't cause the plot to constantly update.

**property transmission: mesmerize.analysis.data_types.Transmission**
The input transmission

**Return type** *Transmission*

**set_input**(*transmission:* mesmerize.analysis.data_types.Transmission)
Set the input Transmission with data to plot

**Parameters** **transmission** – Input transmission

**fill_control_widget**(*data_columns: list*, *categorical_columns: list*, *uuid_columns: list*)
Method for filling the control widget(s) when inputs are set. Must be implemented in subclass

**update_plot**(**args*, ***kwargs*)
Must be implemented in subclass

**get_plot_opts**(*drop: bool*) → dict
Must be implemented in subclass

**set_plot_opts**(*opts: dict*)
Must be implemented in subclass

**classmethod signal_blocker**(*func*)
Use as a decorator. Block Qt signals from all QObjects instances in the block_signals_list

**save_plot_dialog**(*path*, **args*)
Plot save dialog

**save_plot**(*path*)
Save the plot as a Transmission in an HDF5 file. Plot parameters are stored as a JSON string within the HDF5 file. See Transmission.to_hdf5

**Parameters** **path** – Path to save the file to. For easy identification use ".ptrn" extension.

**open_plot_dialog**(*filepath*, *dirpath*, **args*, ***kwargs*)
Open plot dialog

**open_plot**(*ptrn_path: str*, *proj_path: str*) → Optional[Tuple[str, str]]
Open a plot saved by the save_plot() method

**Parameters**

- **ptrn_path** – Path to the HDF5 Transmission file. By convention file extension is ".ptrn"
- **proj_path** – Project path of the associated plot data.

#### 1.57 Datapoint Tracer

*User guide*

**class** mesmerize.plotting.**DatapointTracerWidget**

**set_widget**(*datapoint_uuid: uuid.UUID*, *data_column_curve: str*, *row: pandas.core.series.Series*, *proj_path: str*, *history_trace: Optional[list] = None*, *peak_ix: Optional[int] = None*, *tstart: Optional[int] = None*, *tend: Optional[int] = None*, *roi_color: Optional[Union[str, float, int, tuple]] = 'ff0000'*, *clear_linear_regions: bool = True*)
Set the widget from the datapoint.

**Parameters**

- **datapoint_uuid** – appropriate UUID for the datapoint (such as uuid_curve or _pfeature_uuid)
- **data_column_curve** – data column containing an array to plot
- **row** – DataFrame row that corresponds to the datapoint
- **proj_path** – root dir of the project the datapoint comes from, used for finding max & std projections
- **history_trace** – history trace of the datablock the datapoint comes from
- **peak_ix** – Deprecated
- **tstart** – lower bounds for drawing *LinearRegionItem*
- **tend** – upper bounds for drawing *LinearRegionItem*
- **roi_color** – color for drawing the spatial bounds of the ROI

**set_image**(*projection: str*)
Set either the max or std projection image

**Parameters** **projection** – one of either 'max' or 'std'

**open_in_viewer**()
Open the parent Sample of the current datapoint.

#### 1.58 Heatmap

##### 1.58.1 Widgets

Higher level widgets that are directly used by the end-user. Both Heatmap widgets use the same plot variant.

###### HeatmapSplitterWidget

Heatmap with a vertical splitter that can be used to house another widget. The plot is compatible with both 'row' and 'item' selection modes.

**class** mesmerize.plotting.**HeatmapSplitterWidget**(*highlight_mode='row'*)

Widget for interactive heatmaps

**__init__**(*highlight_mode='row'*)

**Parameters** **highlight_mode** – Interactive mode, one of 'row' or 'item'

**set_data**(*dataframes: Union[pandas.core.frame.DataFrame, list], data_column: str, labels_column: str, cmap: str = 'jet', transmission: Optional[mesmerize.analysis.data_types.Transmission] = None, sort: bool = True, reset_sorting: bool = True, **kwargs*)

Set the data and then set the plot

**Parameters**

- **dataframes** – list of dataframes or a single DataFrame
- **data_column** – data column of the dataframe that is plotted in the heatmap
- **labels_column** – dataframe column (usually categorical labels) used to generate the y-labels and legend.
- **cmap** – colormap choice
- **transmission** – transmission object that dataframe originates, used to calculate data units if passed
- **sort** – if False, the sort comboBox is ignored
- **reset_sorting** – reset the order of the rows in the heatmap
- **kwargs** – Passed to Heatmap.set

**_set_sort_order**(*column: str*)

Set the sort order of the heatmap rows according to a dataframe column. The column must contain categorical values. The rows are grouped together according to the categorical values.

**Parameters** **column** – DataFrame column containing categorical values used for sorting the heatmap rows

**set_transmission**(*transmission: mesmerize.analysis.data_types.Transmission*)

Set the input transmission

**get_transmission**() → *mesmerize.analysis.data_types.Transmission*

Get the input transmission

**highlight_row**(*ix: int*)

Highlight a row on the heatmap

##### HeatmapTracerWidget

Heatmap with an embedded *Datapoint Tracer* that can be saved and restored.

**class** mesmerize.plotting.**HeatmapTracerWidget**

Bases: *mesmerize.plotting.widgets.base.BasePlotWidget*, *mesmerize.plotting.widgets.heatmap.widget.HeatmapSplitterWidget*

Heatmap with an embedded datapoint tracer

**drop_opts = ['dataframes', 'transmission']**

keys of the plot_opts dict that are not JSON compatible and not required for restoring this plot

**live_datapoint_tracer**

The embedded *Datapoint Tracer <API_DatapointTracer>*

**set_current_datapoint**(*ix: tuple*)

Set the currently selected datapoint in the *Datapoint Tracer*.

**Parameters** **ix** – index, (x, y). x is always 0 for this widget since it only uses 'row' selection mode and not 'item'

**get_plot_opts**(*drop: bool = False*) → dict

Get the plot options

**Parameters** **drop** – Drop the non-json compatible objects that are not necessary to restore this plot

**update_plot**()

Calls set_data and passes dict from get_plot_opts() as keyword arguments

**get_cluster_kwargs**() → dict

Organize kwargs for visualization of hierarchical clustering

**set_data**(**args*, *datapoint_tracer_curve_column: str = None*, ***kwargs*)

Set the plot data, parameters and draw the plot. If the input Transmission comes directly from the FCluster it will pass a dict from get_cluster_kwargs() to the cluster_kwargs argument. Else it will pass None to cluster_kwargs.

**Parameters**

- **args** – arguments to pass to superclass set_data() method
- **datapoint_tracer_curve_column** – Data column containing curves to use in the datapoint tracer
- **kwargs** – keyword arguments, passed to superclass set_data() method

**property transmission: mesmerize.analysis.data_types.Transmission**

The input transmission

**Return type** *Transmission*

**save_plot_dialog**(*path*, **args*)

Plot save dialog

**save_plot**(*path*)

Save the plot as a Transmission in an HDF5 file. Plot parameters are stored as a JSON string within the HDF5 file. See Transmission.to_hdf5

**Parameters** **path** – Path to save the file to. For easy identification use ".ptrn" extension.

**open_plot_dialog**(*filepath*, *dirpath*, **args*, ***kwargs*)

Open plot dialog

**open_plot**(*ptrn_path: str*, *proj_path: str*) → Optional[Tuple[str, str]]

Open a plot saved by the save_plot() method

**Parameters**

- **ptrn_path** – Path to the HDF5 Transmission file. By convention file extension is ".ptrn"
- **proj_path** – Project path of the associated plot data.

##### 1.58.2 Variant

Lower level widget that handles the actual plotting and user interaction

class mesmerize.plotting.variants.**Heatmap**(*highlight_mode='row'*)

Bases: `mesmerize.pyqtgraphCore.widgets.MatplotlibWidget.MatplotlibWidget`

Heatmap plot variant

**sig_selection_changed**

Emits indices of data coordinates (x, y) from mouse-click events on the heatmap

**__init__**(*highlight_mode='row'*)

**Parameters** **highlight_mode** – The selection mode for the heatmap. One of either 'row' or 'item'

**data**

2D numpy array of the heatmap data

**selector**

Selection instance that organizes mouse click events on the heatmap

**plot**

ClusterGrid object instance containing the plot Axes

**set**(*data: numpy.ndarray*, **args*, *ylabels: Optional[Union[pandas.core.series.Series, numpy.ndarray, list]] = None*, *ylabels_cmap: str = 'tab20'*, *cluster_kwargs: Optional[dict] = None*, ***kwargs*)

**Parameters**

- **data** – 2D numpy array
- **args** – Additional args that are passed to sns.heatmap()
- **ylabels** – Labels used to create the ylabels bar
- **ylabels_cmap** – colormap for the ylabels bar
- **cluster_kwargs** – keyword arguments for visualizing hierarchical clustering
- **kwargs** – Additional kwargs that are passed to sns.heatmap()

**add_stimulus_indicator**(*start: int*, *end: int*, *color: str*)

Add lines to indicate the start and end of a stimulus or behavioral period

**Parameters**

- **start** – start index
- **end** – end index
- **color** – line color

#### 1.59 KShape

**class** mesmerize.plotting.**KShapeWidget**(*parent=None*)

Bases: PyQt5.QtWidgets.QMainWindow, *mesmerize.plotting.widgets.base.BasePlotWidget*

User-end KShape widget

**sig_output_changed**
Emits output Transmission containing cluster labels

**drop_opts = None**
Unused by this plot widget

**property input_arrays: numpy.ndarray**
The input arrays for clustering

**Returns** 2D array, shape is [num_samples, padded_peak_curve_length]

**Return type** np.ndarray

**property ks: tslearn.clustering.KShape**
tslearn KShape object

**property n_clusters: int**
Number of clusters

**Return type** int

**property train_data: numpy.ndarray**
The training data for clustering

**Returns** Training data as a 2D array, shape is [n_samples, padded_curve_length]

**Return type** np.ndarray

**property y_pred: numpy.ndarray**
Predicted cluster labels after the model converges

**Returns** 1D array of cluster labels that correspond to the input_data

**Return type** np.ndarray

**property cluster_centers: numpy.ndarray**
Cluster centroids

**Returns** 2D array, shape is [n_clusters, centroid_array]

**Return type** np.ndarray

**property cluster_means: numpy.ndarray**
The cluster means

**Returns** 2D array, shape is [cluster_label, mean_array]

**Return type** np.ndarray

**property params: dict**
Parameters dict.

**Return type** dict

**set_input**(*transmission:* mesmerize.analysis.data_types.Transmission)
Set the input Transmission for the widget

**Parameters** **transmission** – Input Transmission

**pad_input_data**(*a: numpy.ndarray*, *method: str = 'random'*) → numpy.ndarray
Pad all the input arrays so that are of the same length. The length is determined by the largest input array. The padding value for each input array is the minimum value in that array.

Padding for each input array is either done after the array's last index to fill up to the length of the largest input array (method 'fill-size') or the padding is randomly flanked to the input array (method 'random) for easier visualization.

**Parameters**

- **a** – 1D array of input arrays where each element is a sample array
- **method** – 'fill-size' or 'random'

**Returns** 2D array of the padded arrays in the rows

**start_process**(**args*, ***kwargs*)
Start the the KShape clustering in a QProcess

**property transmission: mesmerize.analysis.data_types.Transmission**
The input transmission

**Return type** *Transmission*

**class** mesmerize.plotting.widgets.kshape.widget.**KShapeMeansPlot**(*parent*)
Bases: mesmerize.pyqtgraphCore.widgets.MatplotlibWidget.MatplotlibWidget

Means plots grouped by cluster membership

**axs**
array of axis objects used for drawing the means plots, shape is [nrows, ncols]

**set_plots**(*input_arrays: numpy.ndarray*, *n_clusters: int*, *y_pred: numpy.ndarray*, *xzero_pos: str*, *error_band*)
Set the subplots

**Parameters**

- **input_arrays** – padded input arrays (2D), shape is [num_samples, padded_peak_curve_length]
- **n_clusters** – number of clusters
- **y_pred** – cluster predictions (labels)
- **xzero_pos** – set the zero position as the 'zero' position of the input array or the 'maxima' of the input array
- **error_band** – Type of error band to show, one of either 'ci' or 'std'

**class** mesmerize.plotting.widgets.kshape.widget.**KShapePlot**(*parent*)
Bases: PyQt5.QtWidgets.QDockWidget

Curves plots, showing a sample of individual curves from a single cluster

**ax**
The Axes object for this plot

**plot**
MatplotlibWidget() instance

#### 1.60 Proportions

**class** mesmerize.plotting.**ProportionsWidget**

Bases: *mesmerize.plotting.widgets.base.BasePlotWidget*, mesmerize.pyqtgraphCore.widgets.MatplotlibWidget.MatplotlibWidget

**drop_opts = ['xs', 'ys']**

Drop the 'xs' and 'ys' since they are pd.Series objects and not required for restoring the plot

**property ax: matplotlib.axes._axes.Axes**

The Axes object for this plot

**Returns** The Axes object for this plot

**Return type** AXes

**get_plot_opts**(*drop: bool = False*)

Get the plot options

**Parameters drop** – Drop the 'xs' and 'ys' objects when saving the returned dict for saving to an hdf5 file

**update_plot**()

Update the plot data and draw

**export**(**args, **kwargs*)

Export as a csv file

**property transmission: mesmerize.analysis.data_types.Transmission**

The input transmission

**Return type** *Transmission*

**save_plot_dialog**(*path, *args*)

Plot save dialog

**save_plot**(*path*)

Save the plot as a Transmission in an HDF5 file. Plot parameters are stored as a JSON string within the HDF5 file. See Transmission.to_hdf5

**Parameters path** – Path to save the file to. For easy identification use ".ptrn" extension.

**open_plot_dialog**(*filepath, dirpath, *args, **kwargs*)

Open plot dialog

**open_plot**(*ptrn_path: str, proj_path: str*) → Optional[Tuple[str, str]]

Open a plot saved by the save_plot() method

**Parameters**

- **ptrn_path** – Path to the HDF5 Transmission file. By convention file extension is ".ptrn"
- **proj_path** – Project path of the associated plot data.

#### 1.61 Scatter Plot

##### 1.61.1 ScatterPlotWidget

Higher level widget that is directly used by the end-user. Scatter plot with docked Control Widget, *Datapoint Tracer*, and Console.

**class** mesmerize.plotting.**ScatterPlotWidget**

Bases: PyQt5.QtWidgets.QMainWindow, *mesmerize.plotting.widgets.base.BasePlotWidget*

**live_datapoint_tracer**

Instance of *DatapointTracer*

**set_update_live**(*b: bool*)

Must be implemented in subclass

**set_input**(**args*, ***kwds*)

Set the input Transmission with data to plot

**Parameters** **transmission** – Input transmission

**get_plot_opts**(*drop: bool = False*) → dict

Get the plot options

**Parameters** **drop** – no drop opts are specified for this plot

**set_plot_opts**(**args*, ***kwds*)

Must be implemented in subclass

**update_plot**()

Update the plot data and draw

**set_current_datapoint**(*identifier: uuid.UUID*)

Set the UUID of the current datapoint and update the Datapoint Tracer

**property transmission: mesmerize.analysis.data_types.Transmission**

The input transmission

**Return type** *Transmission*

**save_plot_dialog**(*path*, **args*)

Plot save dialog

**save_plot**(*path*)

Save the plot as a Transmission in an HDF5 file. Plot parameters are stored as a JSON string within the HDF5 file. See Transmission.to_hdf5

**Parameters** **path** – Path to save the file to. For easy identification use ".ptrn" extension.

**open_plot_dialog**(*filepath*, *dirpath*, **args*, ***kwargs*)

Open plot dialog

**open_plot**(*ptrn_path: str*, *proj_path: str*) → Optional[Tuple[str, str]]

Open a plot saved by the save_plot() method

**Parameters**

- **ptrn_path** – Path to the HDF5 Transmission file. By convention file extension is ".ptrn"
- **proj_path** – Project path of the associated plot data.

##### 1.61.2 Variant

Lower level widget that interfaces with pqytgraph.ScatterPlotItem and has some helper methods.

**class** `mesmerize.plotting.variants.`**`PgScatterPlot`**(*graphics_view: mesmerize.pyqtgraphCore.widgets.GraphicsLayoutWidget.GraphicsLayoutWidg* [illegible] *parent=None*)

Bases: `PyQt5.QtCore.QObject`

**`signal_spot_clicked`**
Emits the UUID of a spot when it is clicked

**`__init__`**(*graphics_view: mesmerize.pyqtgraphCore.widgets.GraphicsLayoutWidget.GraphicsLayoutWidget*, *parent=None*)

**Parameters** **`graphics_view`** – This plot will instantiate within this GraphicsLayoutWidget

**`add_data`**(*xs: numpy.ndarray*, *ys: numpy.ndarray*, *uuid_series: pandas.core.series.Series*, *color: Union[str, PyQt5.QtGui.QColor, PyQt5.QtGui.QBrush, List[Union[PyQt5.QtGui.QBrush, PyQt5.QtGui.QColor, str]]]*, *size: int = 10*, ***kwargs*)
Add data to the plot

**Parameters**

- **`xs`** (*`np.ndarray`*) – array of x values, indices must correspond to the "ys" array
- **`ys`** (*`np.ndarray`*) – array of y values, indices must correspond to the "xs" array
- **`uuid_series`** (*`pd.Series`*) – series of UUID values. Each SpotItem on the plot is tagged with these UUIDs, therefore the indices must correspond to the "xs" and "ys" arrays.
- **`color`** (*`Union[str, QtGui.QColor, QtGui.QBrush, List[Union[QtGui.QBrush, QtGui.QColor, str]]]`*) – Either a single color or list of colors that pqytgraph.fn.mkBrush() can accept
- **`size`** (*`int`*) – spot size
- **`kwargs`** – any additional kwargs that are passed to ScatterPlotItem.addPoints()

**`_clicked`**(*plot*, *points*)
Called when the plot is clicked

**`set_legend`**(*colors: dict*, *shapes: Optional[dict] = None*)
Set the legend.

**Parameters**

- **`colors`** (*`Dict[str, Union[QtGui.QColor, QtGui.QBrush, str]]`*) – dict mapping of labels onto their corresponding colors {'label': QtGui.QColor}
- **`shapes`** (*`Dict[str, str]`*) – dict mapping of labels onto their corresponding shapes {'label': <shape as a str>}

**`clear_legend`**()
Clear the legend

**`clear`**()
Clear the plot

#### 1.62 SpaceMap

**class** mesmerize.plotting.**SpaceMapWidget**

Bases: PyQt5.QtWidgets.QMainWindow, *mesmerize.plotting.widgets.base.BasePlotWidget*

**sample_df**

sub-dataframe of the current sample

**update_plot**(**args*, ***kwargs*)

Must be implemented in subclass

**property transmission: mesmerize.analysis.data_types.Transmission**

The input transmission

**Return type** *Transmission*

**get_plot_opts**(*drop: bool = False*) → dict

Must be implemented in subclass

**save_plot_dialog**(*path*, **args*)

Plot save dialog

**save_plot**(*path*)

Save the plot as a Transmission in an HDF5 file. Plot parameters are stored as a JSON string within the HDF5 file. See Transmission.to_hdf5

**Parameters path** – Path to save the file to. For easy identification use ".ptrn" extension.

**open_plot_dialog**(*filepath*, *dirpath*, **args*, ***kwargs*)

Open plot dialog

**open_plot**(*ptrn_path: str*, *proj_path: str*) → Optional[Tuple[str, str]]

Open a plot saved by the save_plot() method

**Parameters**

- **ptrn_path** – Path to the HDF5 Transmission file. By convention file extension is ".ptrn"
- **proj_path** – Project path of the associated plot data.

**set_plot_opts**(**args*, ***kwds*)

Must be implemented in subclass

CHAPTER

TWO

### INDICES AND TABLES

- genindex
- modindex
- search

### PYTHON MODULE INDEX

## m


mesmerize.analysis.clustering_metrics, 215
mesmerize.analysis.math.cross_correlation, 212
mesmerize.analysis.utils, 210
mesmerize.common, 166
mesmerize.common.qdialogs, 168
mesmerize.common.utils, 166
mesmerize.pyqtgraphCore.flowchart.library.Biology, 219
mesmerize.pyqtgraphCore.flowchart.library.Clustering, 220
mesmerize.pyqtgraphCore.flowchart.library.Data, 216
mesmerize.pyqtgraphCore.flowchart.library.Display, 217
mesmerize.pyqtgraphCore.flowchart.library.Hierarchical, 220
mesmerize.pyqtgraphCore.flowchart.library.Math, 219
mesmerize.pyqtgraphCore.flowchart.library.Signal, 218
mesmerize.pyqtgraphCore.flowchart.library.Transform, 220

### INDEX

#### Symbols


`_AbstractBasePlotWidget` (*class in mesmerize.plotting.widgets.base*), 222
`_AbstractBaseROI` (*class in mesmerize.viewer.modules.roi_manager_modules.roi_types*), 186
`_UUID` (*mesmerize.viewer.core.ViewerWorkEnv attribute*), 169
`__del__()` (*mesmerize.viewer.modules.roi_manager_modules.managers.AbstractBaseManager method*), 181
`__delitem__()` (*mesmerize.viewer.modules.roi_manager_modules.roi_list.ROIList method*), 185
`__getitem__()` (*mesmerize.viewer.modules.roi_manager_modules.roi_list.ROIList method*), 185
`__init__()` (*mesmerize.Transmission method*), 201
`__init__()` (*mesmerize.analysis.cross_correlation.CC_Data method*), 214
`__init__()` (*mesmerize.analysis.data_types.HistoryTrace method*), 204
`__init__()` (*mesmerize.plotting.HeatmapSplitterWidget method*), 225
`__init__()` (*mesmerize.plotting.variants.Heatmap method*), 227
`__init__()` (*mesmerize.plotting.variants.PgScatterPlot method*), 232
`__init__()` (*mesmerize.plotting.widgets.base.BasePlotWidget method*), 223
`__init__()` (*mesmerize.viewer.core.ViewerUtils method*), 171
`__init__()` (*mesmerize.viewer.core.ViewerWorkEnv method*), 169
`__init__()` (*mesmerize.viewer.core.data_types.ImgData method*), 171
`__init__()` (*mesmerize.viewer.core.mesfile.MES method*), 172
`__init__()` (*mesmerize.viewer.modules.batch_manager.ModuleGUI method*), 175
`__init__()` (*mesmerize.viewer.modules.roi_manager.ModuleGUI method*), 180
`__init__()` (*mesmerize.viewer.modules.roi_manager_modules.managers.AbstractBaseManager method*), 181
`__init__()` (*mesmerize.viewer.modules.roi_manager_modules.managers.* method), 184
`__init__()` (*mesmerize.viewer.modules.roi_manager_modules.managers.* method), 181
`__init__()` (*mesmerize.viewer.modules.roi_manager_modules.managers.* method), 182
`__init__()` (*mesmerize.viewer.modules.roi_manager_modules.managers.* method), 183
`__init__()` (*mesmerize.viewer.modules.roi_manager_modules.managers.* method), 183
`__init__()` (*mesmerize.viewer.modules.roi_manager_modules.roi_list.RO* method), 185
`__init__()` (*mesmerize.viewer.modules.roi_manager_modules.roi_types.E* method), 188
`__init__()` (*mesmerize.viewer.modules.roi_manager_modules.roi_types.C* method), 197
`__init__()` (*mesmerize.viewer.modules.roi_manager_modules.roi_types.M* method), 190
`__init__()` (*mesmerize.viewer.modules.roi_manager_modules.roi_types.S* method), 192
`__init__()` (*mesmerize.viewer.modules.roi_manager_modules.roi_types.V* method), 194
`__init__()` (*mesmerize.viewer.modules.roi_manager_modules.roi_types._* method), 186
`__weakref__` (*mesmerize.viewer.core.ViewerWorkEnv attribute*), 169
`__weakref__` (*mesmerize.viewer.core.mesfile.MES attribute*), 172
`__weakref__` (*mesmerize.viewer.modules.roi_manager_modules.managers.AbstractBase attribute*), 181
`__weakref__` (*mesmerize.viewer.modules.roi_manager_modules.roi_list.ROIList attribute*), 186
`__weakref__` (*mesmerize.viewer.modules.roi_manager_modules.roi_types._AbstractBas attribute*), 188
`_add_data_block()` (*mesmerize.analysis.data_types.HistoryTrace method*), 204
`_clear_workEnv()` (*mesmerize.viewer.core.ViewerUtils*

*method*), 172
_clicked() (*mesmerize.plotting.variants.PgScatterPlot method*), 232
_hide_all_graphics_objects() (*mesmerize.viewer.modules.roi_manager_modules.roi_list.ROIList method*), 186
_hide_graphics_object() (*mesmerize.viewer.modules.roi_manager_modules.roi_list.ROIList method*), 186
_load_files() (*mesmerize.Transmission static method*), 203
_organize_meta() (*mesmerize.viewer.core.ViewerWorkEnv static method*), 170
_reindex_list_widget() (*mesmerize.viewer.modules.roi_manager_modules.roi_list.ROIList method*), 185
_set_sort_order() (*mesmerize.plotting.HeatmapSplitterWidget method*), 225
_show_all_graphics_objects() (*mesmerize.viewer.modules.roi_manager_modules.roi_list.ROIList method*), 186
_show_graphics_object() (*mesmerize.viewer.modules.roi_manager_modules.roi_list.ROIList method*), 186
_to_uuid() (*mesmerize.analysis.data_types.HistoryTrace static method*), 205

# A

AbsoluteValue (*class in mesmerize.pyqtgraphCore.flowchart.library.Math*), 219
AbstractBaseManager (*class in mesmerize.viewer.modules.roi_manager_modules.managers*), 181
add_all_components() (*mesmerize.viewer.modules.roi_manager_modules.managers.ManagerCNMFROI method*), 184
add_all_components() (*mesmerize.viewer.modules.roi_manager_modules.managers.ManagerVolCNMF method*), 183
add_data() (*mesmerize.plotting.variants.PgScatterPlot method*), 232
add_item() (*mesmerize.viewer.modules.batch_manager.ModuleGUI method*), 175
add_manual_roi() (*mesmerize.viewer.modules.roi_manager.ModuleGUI method*), 180
add_operation() (*mesmerize.analysis.data_types.HistoryTrace method*), 204
add_roi() (*mesmerize.viewer.modules.roi_manager_modules.managers.AbstractBaseManager method*), 181
add_roi() (*mesmerize.viewer.modules.roi_manager_modules.managers.M method*), 184
add_roi() (*mesmerize.viewer.modules.roi_manager_modules.managers.M method*), 182
add_roi() (*mesmerize.viewer.modules.roi_manager_modules.managers.M method*), 182
add_roi() (*mesmerize.viewer.modules.roi_manager_modules.managers.M method*), 184
add_row() (*mesmerize.viewer.modules.stimmap_modules.page.Page method*), 199
add_stimulus_indicator() (*mesmerize.plotting.variants.Heatmap method*), 227
add_to_batch() (*mesmerize.viewer.modules.cnmf.ModuleGUI method*), 177
add_to_batch_cnmfe() (*mesmerize.viewer.modules.cnmfe.ModuleGUI method*), 178
add_to_batch_corr_pnr() (*mesmerize.viewer.modules.cnmfe.ModuleGUI method*), 177
add_to_batch_elas_corr() (*mesmerize.viewer.modules.caiman_motion_correction.ModuleGUI method*), 177
add_to_viewer() (*mesmerize.viewer.modules.roi_manager_modules.roi_types._AbstractBas method*), 188
add_to_viewer() (*mesmerize.viewer.modules.roi_manager_modules.roi_types.BaseROI method*), 189
add_to_viewer() (*mesmerize.viewer.modules.roi_manager_modules.roi_types.CNMFROI method*), 197
add_to_viewer() (*mesmerize.viewer.modules.roi_manager_modules.roi_types.ManualROI method*), 191
add_to_viewer() (*mesmerize.viewer.modules.roi_manager_modules.roi_types.ScatterROI method*), 193
add_to_viewer() (*mesmerize.viewer.modules.roi_manager_modules.roi_types.VolCNMF method*), 195
AnalysisGraph (*class in mesmerize.pyqtgraphCore.flowchart.library.Display*), 217
append() (*mesmerize.viewer.modules.roi_manager_modules.roi_list.ROIL method*), 185
ArgGroupStat (*class in mesmerize.pyqtgraphCore.flowchart.library.Math*), 219
ArrayStats (*class in mesmerize.pyqtgraphCore.flowchart.library.Math*), 219
auto_colormap() (*in module mesmerize.plotting.utils*),

221
ax (*mesmerize.plotting.ProportionsWidget property*), 230
ax (*mesmerize.plotting.widgets.kshape.widget.KShapePlot attribute*), 229
axs (*mesmerize.plotting.widgets.kshape.widget.KShapeMeansPlot attribute*), 229

## B

BasePlotWidget (*class in mesmerize.plotting.widgets.base*), 223
BaseROI (*class in mesmerize.viewer.modules.roi_manager_modules.roi_types*), 188
BeeswarmPlots (*class in mesmerize.pyqtgraphCore.flowchart.library.Display*), 217
block_signals_list (*mesmerize.plotting.widgets.base.BasePlotWidget attribute*), 223
ButterWorth (*class in mesmerize.pyqtgraphCore.flowchart.library.Signal*), 218

## C

CC_Data (*class in mesmerize.analysis.cross_correlation*), 214
ccs (*mesmerize.analysis.cross_correlation.CC_Data attribute*), 214
channel (*mesmerize.viewer.modules.femtonics_mesc.MEScNavigator attribute*), 179
channels (*mesmerize.viewer.modules.femtonics_mesc.MEScNavigator attribute*), 179
check_meta_path() (*mesmerize.viewer.modules.tiff_io.ModuleGUI method*), 176
check_operation_exists() (*mesmerize.analysis.data_types.HistoryTrace method*), 205
clean_history_trace() (*mesmerize.analysis.data_types.HistoryTrace static method*), 206
clear() (*mesmerize.plotting.variants.PgScatterPlot method*), 232
clear() (*mesmerize.viewer.core.ViewerWorkEnv method*), 170
clear() (*mesmerize.viewer.modules.roi_manager_modules.managers.AbstractBaseManager method*), 181
clear() (*mesmerize.viewer.modules.stimmap_modules.page.Page method*), 199
clear_() (*mesmerize.viewer.modules.roi_manager_modules.roi_list.ROIList method*), 185
clear_legend() (*mesmerize.plotting.variants.PgScatterPlot method*), 232
close_file() (*mesmerize.viewer.modules.femtonics_mesc.MEScNavigator method*), 179
close_file() (*mesmerize.viewer.modules.femtonics_mesc.ModuleGUI method*), 178
cluster_centers (*mesmerize.plotting.KShapeWidget property*), 228
cluster_means (*mesmerize.plotting.KShapeWidget property*), 228
CNMFROI (*class in mesmerize.viewer.modules.roi_manager_modules.roi_types*), 196
compute_cc_data() (*in module mesmerize.analysis.math.cross_correlation*), 213
compute_ccs() (*in module mesmerize.analysis.math.cross_correlation*), 214
create_data_block() (*mesmerize.analysis.data_types.HistoryTrace method*), 204
create_roi_list() (*mesmerize.viewer.modules.roi_manager_modules.managers.ManagerCN method*), 184
create_roi_list() (*mesmerize.viewer.modules.roi_manager_modules.managers.ManagerMa method*), 182
create_roi_list() (*mesmerize.viewer.modules.roi_manager_modules.managers.ManagerSca method*), 183
create_roi_list() (*mesmerize.viewer.modules.roi_manager_modules.managers.ManagerVol method*), 183
create_roi_list() (*mesmerize.viewer.modules.roi_manager_modules.managers.ManagerVol method*), 183
CrossCorr (*class in mesmerize.pyqtgraphCore.flowchart.library.Display*), 217
CtrlNode (*class in mesmerize.pyqtgraphCore.flowchart.library.common*), 221
ctrlWidget() (*mesmerize.pyqtgraphCore.flowchart.library.common.CtrlNode method*), 221
current_index (*mesmerize.viewer.modules.roi_manager_modules.roi_list.ROIList attribute*), 185
curve_data (*mesmerize.viewer.modules.roi_manager_modules.roi_types.M property*), 190
curve_uuids (*mesmerize.analysis.cross_correlation.CC_Data attribute*), 215

# D


data (*mesmerize.plotting.variants.Heatmap attribute*), 227
data_blocks (*mesmerize.analysis.data_types.HistoryTrace property*), 204
DatapointTracerWidget (*class in mesmerize.plotting*), 224
davies_bouldin_score() (*in module mesmerize.analysis.clustering_metrics*), 216
del_item() (*mesmerize.viewer.modules.batch_manager.ModuleGUI method*), 175
delete_row() (*mesmerize.viewer.modules.stimmap_modules.page.Page method*), 199
Derivative (*class in mesmerize.pyqtgraphCore.flowchart.library.Math*), 219
df (*mesmerize.viewer.modules.batch_manager.ModuleGUI attribute*), 175
discard_workEnv() (*mesmerize.viewer.core.ViewerUtils method*), 172
disconnect_all() (*mesmerize.viewer.modules.roi_manager_modules.roi_list.ROIList method*), 185
draw_graph() (*in module mesmerize.common.utils*), 167
draw_graph() (*mesmerize.analysis.data_types.HistoryTrace method*), 206
drop_opts (*mesmerize.plotting.HeatmapTracerWidget attribute*), 226
drop_opts (*mesmerize.plotting.KShapeWidget attribute*), 228
drop_opts (*mesmerize.plotting.ProportionsWidget attribute*), 230
DropNa (*class in mesmerize.pyqtgraphCore.flowchart.library.Data*), 216


# E


empty_df() (*mesmerize.Transmission static method*), 202
epsilon_matrix (*mesmerize.analysis.cross_correlation.CC_Data attribute*), 214
eventFilter() (*mesmerize.viewer.modules.roi_manager.ModuleGUI method*), 180
exceptions_label() (*in module mesmerize.common.qdialogs*), 168
export() (*mesmerize.plotting.ProportionsWidget method*), 230
ExtractStim (*class in mesmerize.pyqtgraphCore.flowchart.library.Biology*), 219


# F


FCluster (*class in mesmerize.pyqtgraphCore.flowchart.library.Hierarchical*), 220
file (*mesmerize.viewer.modules.femtonics_mesc.MEScNavigator attribute*), 178
fill_control_widget() (*mesmerize.plotting.widgets.base.BasePlotWidget method*), 223
FrequencyDomainMagnitude (*class in mesmerize.pyqtgraphCore.flowchart.library.Display*), 217
from_dict() (*mesmerize.analysis.cross_correlation.CC_Data class method*), 215
from_dict() (*mesmerize.analysis.data_types.HistoryTrace static method*), 205
from_hdf5() (*mesmerize.analysis.cross_correlation.CC_Data class method*), 215
from_hdf5() (*mesmerize.Transmission class method*), 202
from_json() (*mesmerize.analysis.data_types.HistoryTrace class method*), 205
from_mesfile() (*mesmerize.viewer.core.ViewerWorkEnv class method*), 170
from_pickle() (*mesmerize.analysis.data_types.HistoryTrace class method*), 206
from_pickle() (*mesmerize.Transmission class method*), 202
from_pickle() (*mesmerize.viewer.core.ViewerWorkEnv class method*), 170
from_proj() (*mesmerize.Transmission class method*), 202
from_state() (*mesmerize.viewer.modules.roi_manager_modules.roi_types._AbstractBas class method*), 188
from_state() (*mesmerize.viewer.modules.roi_manager_modules.roi_types.BaseROI class method*), 189
from_state() (*mesmerize.viewer.modules.roi_manager_modules.roi_types.CNMFROI class method*), 197
from_state() (*mesmerize.viewer.modules.roi_manager_modules.roi_types.ManualROI class method*), 190
from_state() (*mesmer-

*ize.viewer.modules.roi_manager_modules.roi_types.ScatterROI class method*), 192

from_state() (*mesmerize.viewer.modules.roi_manager_modules.roi_types.VolCNMF class method*), 195

from_tiff() (*mesmerize.viewer.core.ViewerWorkEnv class method*), 170

# G

get_all_data_blocks_history() (*mesmerize.analysis.data_types.HistoryTrace method*), 205

get_all_states() (*mesmerize.viewer.modules.roi_manager_modules.managers.AbstractBaseManager method*), 181

get_all_states() (*mesmerize.viewer.modules.roi_manager_modules.managers.ManagerCNMFROI method*), 184

get_all_states() (*mesmerize.viewer.modules.roi_manager_modules.managers.ManagerManual method*), 182

get_all_states() (*mesmerize.viewer.modules.roi_manager_modules.managers.ManagerVolCNMF method*), 184

get_all_tags() (*mesmerize.viewer.modules.roi_manager_modules.roi_types._AbstractBaseROI method*), 187

get_all_tags() (*mesmerize.viewer.modules.roi_manager_modules.roi_types.BaseROI method*), 189

get_all_tags() (*mesmerize.viewer.modules.roi_manager_modules.roi_types.CNMFROI method*), 197

get_all_tags() (*mesmerize.viewer.modules.roi_manager_modules.roi_types.ManualROI method*), 191

get_all_tags() (*mesmerize.viewer.modules.roi_manager_modules.roi_types.ScatterROI method*), 193

get_all_tags() (*mesmerize.viewer.modules.roi_manager_modules.roi_types.VolCNMF method*), 195

get_array_size() (*in module mesmerize.analysis.utils*), 210

get_centerlike() (*in module mesmerize.analysis.clustering_metrics*), 215

get_cluster_kwargs() (*mesmerize.plotting.HeatmapTracerWidget method*), 226

get_cluster_radius() (*in module mesmerize.analysis.clustering_metrics*), 215

get_color() (*mesmerize.viewer.modules.roi_manager_modules.roi_types._AbstractBaseROI method*), 187

get_color() (*mesmerize.viewer.modules.roi_manager_modules.roi_types.BaseROI method*), 189

get_color() (*mesmerize.viewer.modules.roi_manager_modules.roi_types.CNMFROI method*), 196

get_color() (*mesmerize.viewer.modules.roi_manager_modules.roi_types.ManualROI method*), 191

get_color() (*mesmerize.viewer.modules.roi_manager_modules.roi_types.ScatterROI method*), 193

get_color() (*mesmerize.viewer.modules.roi_manager_modules.roi_types.VolCNMF method*), 195

get_colormap() (*in module mesmerize.plotting.utils*), 221

get_data_block_history() (*mesmerize.analysis.data_types.HistoryTrace method*), 205

get_dataframe() (*mesmerize.viewer.modules.stimmap_modules.page.Page method*), 199

get_epsilon() (*in module mesmerize.analysis.math.cross_correlation*), 213

get_epsilon_matrix() (*in module mesmerize.analysis.math.cross_correlation*), 213

get_frequency_linspace() (*in module mesmerize.analysis.utils*), 210

get_hpath() (*mesmerize.viewer.modules.femtonics_mesc.MEScNavigator method*), 179

get_image_references() (*mesmerize.viewer.core.mesfile.MES method*), 172

get_item_index() (*mesmerize.viewer.modules.batch_manager.ModuleGUI method*), 175

get_lag() (*in module mesmerize.analysis.math.cross_correlation*), 213

get_lag_matrix() (*in module mesmerize.analysis.math.cross_correlation*), 213

get_omega() (*in module mesmerize.analysis.math.cross_correlation*), 212

get_operation_params() (*mesmerize.analysis.data_types.HistoryTrace method*), 205

get_operations_list() (*mesmerize.analysis.data_types.HistoryTrace method*), 205

get_params() (*mesmerize.viewer.modules.caiman_motion_correction.ModuleGUI method*), 177

get_params() (*mesmerize.viewer.modules.cnmf.ModuleGUI method*),

177
get_params() (mesmerize.viewer.modules.cnmfe.ModuleGUI method), 177
get_plot_item() (mesmerize.viewer.modules.roi_manager_modules.managers.AbstractBaseManager method), 181
get_plot_opts() (mesmerize.plotting.HeatmapTracerWidget method), 226
get_plot_opts() (mesmerize.plotting.ProportionsWidget method), 230
get_plot_opts() (mesmerize.plotting.ScatterPlotWidget method), 231
get_plot_opts() (mesmerize.plotting.SpaceMapWidget method), 233
get_plot_opts() (mesmerize.plotting.widgets.base._AbstractBasePlotWidget method), 223
get_plot_opts() (mesmerize.plotting.widgets.base.BasePlotWidget method), 223
get_proj_config() (in module mesmerize.common), 166
get_proj_path() (mesmerize.Transmission method), 202
get_project_manager() (in module mesmerize.common), 166
get_proportions() (in module mesmerize.analysis.utils), 211
get_roi_graphics_object() (mesmerize.viewer.modules.roi_manager_modules.roi_types._AbstractBaseROI method), 187
get_roi_graphics_object() (mesmerize.viewer.modules.roi_manager_modules.roi_types.BaseROI method), 188
get_roi_graphics_object() (mesmerize.viewer.modules.roi_manager_modules.roi_types.CNMFROI method), 196
get_roi_graphics_object() (mesmerize.viewer.modules.roi_manager_modules.roi_types.ManualROI method), 190
get_roi_graphics_object() (mesmerize.viewer.modules.roi_manager_modules.roi_types.ScatterROI method), 192
get_roi_graphics_object() (mesmerize.viewer.modules.roi_manager_modules.roi_types.VolCNMF method), 194
get_sampling_rate() (in module mesmerize.analysis.utils), 211
get_selected_items() (mesmerize.viewer.modules.batch_manager.ModuleGUI method), 175
get_state() (mesmerize.plotting.utils.WidgetRegistry method), 222
get_sys_config() (in module mesmerize.common), 166
get_tag() (mesmerize.viewer.modules.roi_manager_modules.roi_types._A[illegible] method), 187
get_tag() (mesmerize.viewer.modules.roi_manager_modules.roi_types.Ba[illegible] method), 189
get_tag() (mesmerize.viewer.modules.roi_manager_modules.roi_types.Cl[illegible] method), 197
get_tag() (mesmerize.viewer.modules.roi_manager_modules.roi_types.M[illegible] method), 191
get_tag() (mesmerize.viewer.modules.roi_manager_modules.roi_types.Sc[illegible] method), 193
get_tag() (mesmerize.viewer.modules.roi_manager_modules.roi_types.Vo[illegible] method), 195
get_threshold_matrix() (mesmerize.analysis.cross_correlation.CC_Data method), 215
get_transmission() (mesmerize.plotting.HeatmapSplitterWidget method), 225
get_units() (mesmerize.viewer.modules.stimmap_modules.page.Page method), 199
get_window_manager() (in module mesmerize.common), 166

## H

HdfTools (class in mesmerize.common.utils), 166
Heatmap (class in mesmerize.plotting.variants), 227
Heatmap (class in mesmerize.pyqtgraphCore.flowchart.library.Display), 217
HeatmapSplitterWidget (class in mesmerize.plotting), 225
HeatmapTracerWidget (class in mesmerize.plotting), 226
highlight_curve() (mesmerize.viewer.modules.roi_manager_modules.roi_list.ROIList method), 186
highlight_roi() (mesmerize.viewer.modules.roi_manager_modules.roi_list.ROIList method), 185
highlight_row() (mesmerize.plotting.HeatmapSplitterWidget method), 225
history (mesmerize.analysis.data_types.HistoryTrace property), 204
history_trace (mesmerize.viewer.core.ViewerWorkEnv attribute), 170
HistoryTrace (class in mesmerize.analysis.data_types), 203

## I


iloc (*class in mesmerize.pyqtgraphCore.flowchart.library.Data*), 217
ImgData (*class in mesmerize.viewer.core.data_types*), 171
imgdata (*mesmerize.viewer.core.ViewerWorkEnv attribute*), 170
import_from_imagej() (*mesmerize.viewer.modules.roi_manager.ModuleGUI method*), 180
import_from_imagej() (*mesmerize.viewer.modules.roi_manager_modules.managers.ManagerManual method*), 182
import_recording() (*mesmerize.viewer.modules.femtonics_mesc.ModuleGUI method*), 178
Inconsistent (*class in mesmerize.pyqtgraphCore.flowchart.library.Hierarchical*), 220
input_arrays (*mesmerize.plotting.KShapeWidget property*), 228
Integrate (*class in mesmerize.pyqtgraphCore.flowchart.library.Math*), 219
iRFFT (*class in mesmerize.pyqtgraphCore.flowchart.library.Signal*), 218
is_empty() (*mesmerize.viewer.modules.roi_manager_modules.managers.AbstractBaseManager method*), 181
isEmpty (*mesmerize.viewer.core.ViewerWorkEnv attribute*), 170


## K


KMeans (*class in mesmerize.pyqtgraphCore.flowchart.library.Clustering*), 220
ks (*mesmerize.plotting.KShapeWidget property*), 228
KShape (*class in mesmerize.pyqtgraphCore.flowchart.library.Clustering*), 220
KShapeMeansPlot (*class in mesmerize.plotting.widgets.kshape.widget*), 229
KShapePlot (*class in mesmerize.plotting.widgets.kshape.widget*), 229
KShapeWidget (*class in mesmerize.plotting*), 228


## L


labels (*mesmerize.analysis.cross_correlation.CC_Data attribute*), 215
lag_matrix (*mesmerize.analysis.cross_correlation.CC_Data attribute*), 214
LDA (*class in mesmerize.pyqtgraphCore.flowchart.library.Transform*), 220
Linkage (*class in mesmerize.pyqtgraphCore.flowchart.library.Hierarchical*), 220
LinRegress (*class in mesmerize.pyqtgraphCore.flowchart.library.Math*), 219
list_widget (*mesmerize.viewer.modules.roi_manager_modules.roi_list.ROIList attribute*), 185
list_widget_tags (*mesmerize.viewer.modules.roi_manager_modules.roi_list.ROIList attribute*), 185
listw_channels (*mesmerize.viewer.modules.femtonics_mesc.MEScNavigator attribute*), 179
listw_sessions (*mesmerize.viewer.modules.femtonics_mesc.MEScNavigator attribute*), 179
listw_units (*mesmerize.viewer.modules.femtonics_mesc.MEScNavigator attribute*), 179
live_datapoint_tracer (*mesmerize.plotting.HeatmapTracerWidget attribute*), 226
live_datapoint_tracer (*mesmerize.plotting.ScatterPlotWidget attribute*), 231
load() (*mesmerize.viewer.modules.tiff_io.ModuleGUI method*), 176
load_dataframe() (*mesmerize.common.utils.HdfTools static method*), 167
load_dict() (*mesmerize.common.utils.HdfTools static method*), 167
load_img() (*mesmerize.viewer.core.mesfile.MES method*), 172
load_item_input() (*mesmerize.viewer.modules.batch_manager.ModuleGUI method*), 175
load_item_output() (*mesmerize.viewer.modules.batch_manager.ModuleGUI method*), 176
load_mesfile() (*mesmerize.viewer.core.ViewerWorkEnv static method*), 170
LoadFile (*class in mesmerize.pyqtgraphCore.flowchart.library.Data*), 216
LoadProjDF (*class in mesmerize.pyqtgraphCore.flowchart.library.Data*), 216
LogTransform (*class in mesmer-*

ize.pyqtgraphCore.flowchart.library.Math), 219

## M

make_runfile() (*in module mesmerize.common.utils*), 166
make_workdir() (*in module mesmerize.common.utils*), 166
manager (*mesmerize.viewer.modules.roi_manager.ModuleGUI attribute*), 180
ManagerCNMFROI (*class in mesmerize.viewer.modules.roi_manager_modules.managers*), 184
ManagerManual (*class in mesmerize.viewer.modules.roi_manager_modules.managers*), 181
ManagerScatterROI (*class in mesmerize.viewer.modules.roi_manager_modules.managers*), 182
ManagerVolCNMF (*class in mesmerize.viewer.modules.roi_manager_modules.managers*), 183
ManagerVolROI (*class in mesmerize.viewer.modules.roi_manager_modules.managers*), 183
Manifold (*class in mesmerize.pyqtgraphCore.flowchart.library.Transform*), 220
ManualDFoF (*class in mesmerize.pyqtgraphCore.flowchart.library.Biology*), 219
ManualROI (*class in mesmerize.viewer.modules.roi_manager_modules.roi_types*), 190
map_labels_to_colors() (*in module mesmerize.plotting.utils*), 221
maps (*mesmerize.viewer.modules.stimulus_mapping.ModuleGUI property*), 198
MaxInconsistent (*class in mesmerize.pyqtgraphCore.flowchart.library.Hierarchical*), 220
MaxIncStat (*class in mesmerize.pyqtgraphCore.flowchart.library.Hierarchical*), 220
Merge (*class in mesmerize.pyqtgraphCore.flowchart.library.Data*), 216
merge() (*mesmerize.analysis.data_types.HistoryTrace class method*), 206
merge() (*mesmerize.Transmission class method*), 203
MES (*class in mesmerize.viewer.core.mesfile*), 172
mesc_navigator (*mesmerize.viewer.modules.femtonics_mesc.ModuleGUI attribute*), 178
MEScNavigator (*class in mesmerize.viewer.modules.femtonics_mesc*), 178
mesmerize.analysis.clustering_metrics
  module, 215
mesmerize.analysis.math.cross_correlation
  module, 212
mesmerize.analysis.utils
  module, 210
mesmerize.common
  module, 166
mesmerize.common.qdialogs
  module, 168
mesmerize.common.utils
  module, 166
mesmerize.pyqtgraphCore.flowchart.library.Biology
  module, 219
mesmerize.pyqtgraphCore.flowchart.library.Clustering
  module, 220
mesmerize.pyqtgraphCore.flowchart.library.Data
  module, 216
mesmerize.pyqtgraphCore.flowchart.library.Display
  module, 217
mesmerize.pyqtgraphCore.flowchart.library.Hierarchical
  module, 220
mesmerize.pyqtgraphCore.flowchart.library.Math
  module, 219
mesmerize.pyqtgraphCore.flowchart.library.Signal
  module, 218
mesmerize.pyqtgraphCore.flowchart.library.Transform
  module, 220
module
  mesmerize.analysis.clustering_metrics, 215
  mesmerize.analysis.math.cross_correlation, 212
  mesmerize.analysis.utils, 210
  mesmerize.common, 166
  mesmerize.common.qdialogs, 168
  mesmerize.common.utils, 166
  mesmerize.pyqtgraphCore.flowchart.library.Biology, 219
  mesmerize.pyqtgraphCore.flowchart.library.Clustering, 220
  mesmerize.pyqtgraphCore.flowchart.library.Data, 216
  mesmerize.pyqtgraphCore.flowchart.library.Display, 217
  mesmerize.pyqtgraphCore.flowchart.library.Hierarchical, 220
  mesmerize.pyqtgraphCore.flowchart.library.Math, 219
  mesmerize.pyqtgraphCore.flowchart.library.Signal, 218

mesmerize.pyqtgraphCore.flowchart.library.Transform), 220
ModuleGUI (class in mesmerize.viewer.modules.batch_manager), 175
ModuleGUI (class in mesmerize.viewer.modules.caiman_motion_correction), 177
ModuleGUI (class in mesmerize.viewer.modules.cnmf), 177
ModuleGUI (class in mesmerize.viewer.modules.cnmfe), 177
ModuleGUI (class in mesmerize.viewer.modules.femtonics_mesc), 178
ModuleGUI (class in mesmerize.viewer.modules.roi_manager), 180
ModuleGUI (class in mesmerize.viewer.modules.stimulus_mapping), 198
ModuleGUI (class in mesmerize.viewer.modules.tiff_io), 176

## N

n_clusters (mesmerize.plotting.KShapeWidget property), 228
ncc_c() (in module mesmerize.analysis.math.cross_correlation), 212
NeuralDynamics (class in mesmerize.pyqtgraphCore.flowchart.library.Transform), 220
Normalize (class in mesmerize.pyqtgraphCore.flowchart.library.Signal), 218
NormRaw (class in mesmerize.pyqtgraphCore.flowchart.library.Data), 216

## O

open_in_viewer() (mesmerize.plotting.DatapointTracerWidget method), 224
open_plot() (mesmerize.plotting.HeatmapTracerWidget method), 226
open_plot() (mesmerize.plotting.ProportionsWidget method), 230
open_plot() (mesmerize.plotting.ScatterPlotWidget method), 231
open_plot() (mesmerize.plotting.SpaceMapWidget method), 233
open_plot() (mesmerize.plotting.widgets.base._AbstractBasePlotWidget method), 223
open_plot() (mesmerize.plotting.widgets.base.BasePlotWidget method), 224
open_plot_dialog() (mesmerize.plotting.HeatmapTracerWidget method), 226
open_plot_dialog() (mesmerize.plotting.ProportionsWidget method), 230
open_plot_dialog() (mesmerize.plotting.ScatterPlotWidget method), 231
open_plot_dialog() (mesmerize.plotting.SpaceMapWidget method), 233
open_plot_dialog() (mesmerize.plotting.widgets.base.BasePlotWidget method), 224
organize_dataframe_columns() (in module mesmerize.analysis.utils), 211

## P

package_for_project() (mesmerize.viewer.modules.roi_manager.ModuleGUI method), 180
pad_arrays() (in module mesmerize.analysis.utils), 211
pad_input_data() (mesmerize.plotting.KShapeWidget method), 228
PadArrays (class in mesmerize.pyqtgraphCore.flowchart.library.Data), 216
Page (class in mesmerize.viewer.modules.stimmap_modules.page), 199
params (mesmerize.plotting.KShapeWidget property), 228
path (mesmerize.viewer.modules.femtonics_mesc.MEScNavigator attribute), 178
PCA (class in mesmerize.pyqtgraphCore.flowchart.library.Transform), 220
PeakDetect (class in mesmerize.pyqtgraphCore.flowchart.library.Signal), 218
PeakFeatures (class in mesmerize.pyqtgraphCore.flowchart.library.Signal), 218
PgScatterPlot (class in mesmerize.plotting.variants), 232
Plot (class in mesmerize.pyqtgraphCore.flowchart.library.Display), 217
plot (mesmerize.plotting.variants.Heatmap attribute), 227
plot (mesmerize.plotting.widgets.kshape.widget.KShapePlot attribute), 229
plot_manual_roi_regions() (mesmerize.viewer.modules.roi_manager_modules.roi_list.ROIList method), 186

plot_widgets (*mesmerize.viewer.modules.femtonics_mesc.ModuleGUI attribute*), 178
PowerSpectralDensity (*class in mesmerize.pyqtgraphCore.flowchart.library.Signal*), 218
present_exceptions() (*in module mesmerize.common.qdialogs*), 168
previous_index (*mesmerize.viewer.modules.roi_manager_modules.roi_list.ROIList attribute*), 185
process() (*mesmerize.pyqtgraphCore.flowchart.library.common.CtrlNode method*), 221
process_batch() (*mesmerize.viewer.modules.batch_manager.ModuleGUI method*), 176
Proportions (*class in mesmerize.pyqtgraphCore.flowchart.library.Display*), 217
ProportionsWidget (*class in mesmerize.plotting*), 230

## R

register() (*mesmerize.plotting.utils.WidgetRegistry method*), 222
reindex_colormap() (*mesmerize.viewer.modules.roi_manager_modules.roi_list.ROIList method*), 185
remove_from_viewer() (*mesmerize.viewer.modules.roi_manager_modules.roi_types._AbstractBaseROI method*), 188
remove_from_viewer() (*mesmerize.viewer.modules.roi_manager_modules.roi_types.BaseROI method*), 189
remove_from_viewer() (*mesmerize.viewer.modules.roi_manager_modules.roi_types.CNMFROI method*), 197
remove_from_viewer() (*mesmerize.viewer.modules.roi_manager_modules.roi_types.ManualROI method*), 191
remove_from_viewer() (*mesmerize.viewer.modules.roi_manager_modules.roi_types.ScatterROI method*), 193
remove_from_viewer() (*mesmerize.viewer.modules.roi_manager_modules.roi_types.VolCNMF method*), 195
Resample (*class in mesmerize.pyqtgraphCore.flowchart.library.Signal*), 218
reset_color() (*mesmerize.viewer.modules.roi_manager_modules.roi_types._AbstractBaseROI method*), 187
reset_color() (*mesmerize.viewer.modules.roi_manager_modules.roi_types.BaseROI method*), 189
reset_color() (*mesmerize.viewer.modules.roi_manager_modules.roi_types.CNMFROI method*), 196
reset_color() (*mesmerize.viewer.modules.roi_manager_modules.roi_types.ManualROI method*), 191
reset_color() (*mesmerize.viewer.modules.roi_manager_modules.roi_types.ScatterROI method*), 193
reset_color() (*mesmerize.viewer.modules.roi_manager_modules.roi_types.VolCNMF method*), 194
restore_from_states() (*mesmerize.viewer.modules.roi_manager_modules.managers.ManagerCN[illegible] method*), 184
restore_from_states() (*mesmerize.viewer.modules.roi_manager_modules.managers.ManagerMa[illegible] method*), 182
restore_from_states() (*mesmerize.viewer.modules.roi_manager_modules.managers.ManagerSca[illegible] method*), 182
restore_from_states() (*mesmerize.viewer.modules.roi_manager_modules.managers.ManagerVol[illegible] method*), 184
restoreState() (*mesmerize.pyqtgraphCore.flowchart.library.common.CtrlNode method*), 221
RFFT (*class in mesmerize.pyqtgraphCore.flowchart.library.Signal*), 218
roi_list (*mesmerize.viewer.modules.roi_manager_modules.managers.Abs[illegible] attribute*), 181
roi_manager (*mesmerize.viewer.core.ViewerWorkEnv attribute*), 171
ROIList (*class in mesmerize.viewer.modules.roi_manager_modules.roi_list*), 185

## S

sample_df (*mesmerize.plotting.SpaceMapWidget attribute*), 233
sample_id (*mesmerize.viewer.core.ViewerWorkEnv attribute*), 171
Save (*class in mesmerize.pyqtgraphCore.flowchart.library.Data*), 216
save_dataframe() (*mesmerize.common.utils.HdfTools static method*), 166
save_dict() (*mesmerize.common.utils.HdfTools static method*), 167
save_plot() (*mesmerize.plotting.HeatmapTracerWidget method*), 226
save_plot() (*mesmerize.plotting.ProportionsWidget*

*method*), 230
save_plot() (*mesmerize.plotting.ScatterPlotWidget method*), 231
save_plot() (*mesmerize.plotting.SpaceMapWidget method*), 233
save_plot() (*mesmerize.plotting.widgets.base._AbstractBasePlotWidget method*), 223
save_plot() (*mesmerize.plotting.widgets.base.BasePlotWidget method*), 224
save_plot_dialog() (*mesmerize.plotting.HeatmapTracerWidget method*), 226
save_plot_dialog() (*mesmerize.plotting.ProportionsWidget method*), 230
save_plot_dialog() (*mesmerize.plotting.ScatterPlotWidget method*), 231
save_plot_dialog() (*mesmerize.plotting.SpaceMapWidget method*), 233
save_plot_dialog() (*mesmerize.plotting.widgets.base.BasePlotWidget method*), 223
saveState() (*mesmerize.pyqtgraphCore.flowchart.library.common.CtrlNode method*), 221
SavitzkyGolay (*class in mesmerize.pyqtgraphCore.flowchart.library.Signal*), 218
ScalerMeanVariance (*class in mesmerize.pyqtgraphCore.flowchart.library.Signal*), 218
ScatterPlot (*class in mesmerize.pyqtgraphCore.flowchart.library.Display*), 217
ScatterPlotWidget (*class in mesmerize.plotting*), 231
ScatterROI (*class in mesmerize.viewer.modules.roi_manager_modules.roi_types*), 192
SelectColumns (*class in mesmerize.pyqtgraphCore.flowchart.library.Data*), 217
selector (*mesmerize.plotting.variants.Heatmap attribute*), 227
SelectRows (*class in mesmerize.pyqtgraphCore.flowchart.library.Data*), 217
session (*mesmerize.viewer.modules.femtonics_mesc.MEScNavigator attribute*), 179
sessions (*mesmerize.viewer.modules.femtonics_mesc.MEScNavigator attribute*), 179
set() (*mesmerize.plotting.variants.Heatmap method*), 227
set_all_from_states() (*mesmerize.viewer.modules.roi_manager.ModuleGUI method*), 180
set_channel() (*mesmerize.viewer.modules.femtonics_mesc.MEScNavigator method*), 179
set_color() (*mesmerize.viewer.modules.roi_manager_modules.roi_types._AbstractBas method*), 187
set_color() (*mesmerize.viewer.modules.roi_manager_modules.roi_types.BaseROI method*), 189
set_color() (*mesmerize.viewer.modules.roi_manager_modules.roi_types.CNMFROI method*), 196
set_color() (*mesmerize.viewer.modules.roi_manager_modules.roi_types.ManualROI method*), 191
set_color() (*mesmerize.viewer.modules.roi_manager_modules.roi_types.ScatterROI method*), 193
set_color() (*mesmerize.viewer.modules.roi_manager_modules.roi_types.VolCNMF method*), 195
set_current_datapoint() (*mesmerize.plotting.HeatmapTracerWidget method*), 226
set_current_datapoint() (*mesmerize.plotting.ScatterPlotWidget method*), 231
set_current_index() (*mesmerize.viewer.modules.roi_manager_modules.roi_list.ROIList method*), 185
set_curve_data() (*mesmerize.viewer.modules.roi_manager_modules.roi_types.CNMFROI method*), 197
set_curve_data() (*mesmerize.viewer.modules.roi_manager_modules.roi_types.ScatterROI method*), 192
set_curve_data() (*mesmerize.viewer.modules.roi_manager_modules.roi_types.VolCNMF method*), 196
set_data() (*mesmerize.plotting.HeatmapSplitterWidget method*), 225
set_data() (*mesmerize.plotting.HeatmapTracerWidget method*), 226
set_data() (*mesmerize.viewer.modules.stimmap_modules.page.Page method*), 199
set_file() (*mesmerize.viewer.modules.femtonics_mesc.ModuleGUI method*), 178
set_file_path() (*mesmerize.viewer.modules.femtonics_mesc.MEScNavigator method*), 179
set_image() (*mesmerize.plotting.DatapointTracerWidget method*),

224
set_input() (mesmerize.plotting.KShapeWidget method), 228
set_input() (mesmerize.plotting.ScatterPlotWidget method), 231
set_input() (mesmerize.plotting.widgets.base._AbstractBasePlotWidget method), 222
set_input() (mesmerize.plotting.widgets.base.BasePlotWidget method), 223
set_legend() (mesmerize.plotting.variants.PgScatterPlot method), 232
set_list_widget_tags() (mesmerize.viewer.modules.roi_manager_modules.roi_list.ROIList method), 186
set_original_color() (mesmerize.viewer.modules.roi_manager_modules.roi_types._AbstractBaseROI method), 187
set_original_color() (mesmerize.viewer.modules.roi_manager_modules.roi_types.BaseROI method), 189
set_original_color() (mesmerize.viewer.modules.roi_manager_modules.roi_types.CNMFROI method), 196
set_original_color() (mesmerize.viewer.modules.roi_manager_modules.roi_types.ManualROI method), 191
set_original_color() (mesmerize.viewer.modules.roi_manager_modules.roi_types.ScatterROI method), 193
set_original_color() (mesmerize.viewer.modules.roi_manager_modules.roi_types.VolCNMF method), 195
set_pg_roi_plot() (mesmerize.viewer.modules.roi_manager_modules.roi_list.ROIList method), 186
set_plot_opts() (mesmerize.plotting.ScatterPlotWidget method), 231
set_plot_opts() (mesmerize.plotting.SpaceMapWidget method), 233
set_plot_opts() (mesmerize.plotting.widgets.base._AbstractBasePlotWidget method), 223
set_plot_opts() (mesmerize.plotting.widgets.base.BasePlotWidget method), 223
set_plots() (mesmerize.plotting.widgets.kshape.widget.KShapeMeansPlot method), 229
set_previous_index() (mesmerize.viewer.modules.roi_manager_modules.roi_list.ROIList method), 186
set_proj_path() (mesmerize.Transmission method), 202
set_roi_graphics_object() (mesmerize.viewer.modules.roi_manager_modules.roi_types._AbstractBas[illegible] method), 187
set_roi_graphics_object() (mesmerize.viewer.modules.roi_manager_modules.roi_types.BaseROI method), 189
set_roi_graphics_object() (mesmerize.viewer.modules.roi_manager_modules.roi_types.CNMFROI method), 196
set_roi_graphics_object() (mesmerize.viewer.modules.roi_manager_modules.roi_types.ManualROI method), 190
set_roi_graphics_object() (mesmerize.viewer.modules.roi_manager_modules.roi_types.ScatterROI method), 192
set_roi_graphics_object() (mesmerize.viewer.modules.roi_manager_modules.roi_types.VolCNMF method), 194
set_session() (mesmerize.viewer.modules.femtonics_mesc.MEScNavigator method), 179
set_spot_size() (mesmerize.viewer.modules.roi_manager_modules.managers.ManagerSca[illegible] method), 183
set_spot_size() (mesmerize.viewer.modules.roi_manager_modules.managers.ManagerVol[illegible] method), 184
set_state() (mesmerize.plotting.utils.WidgetRegistry method), 222
set_statusbar() (mesmerize.viewer.core.ViewerUtils method), 172
set_tag() (mesmerize.viewer.modules.roi_manager_modules.roi_types._A[illegible] method), 187
set_tag() (mesmerize.viewer.modules.roi_manager_modules.roi_types.Ba[illegible] method), 189
set_tag() (mesmerize.viewer.modules.roi_manager_modules.roi_types.C[illegible] method), 197
set_tag() (mesmerize.viewer.modules.roi_manager_modules.roi_types.M[illegible] method), 191
set_tag() (mesmerize.viewer.modules.roi_manager_modules.roi_types.Sc[illegible] method), 193
set_tag() (mesmerize.viewer.modules.roi_manager_modules.roi_types.Vo[illegible] method), 195
set_text() (mesmerize.viewer.modules.roi_manager_modules.roi_types._[illegible] method), 187
set_text() (mesmerize.viewer.modules.roi_manager_modules.roi_types.[illegible] method), 189
set_text() (mesmerize.viewer.modules.roi_manager_modules.roi_types.[illegible] method), 196
set_text() (mesmerize.viewer.modules.roi_manager_modules.roi_types.[illegible] method), 191
set_text() (mesmerize.viewer.modules.roi_manager_modules.roi_types.[illegible]

method), 193
set_text() (mesmerize.viewer.modules.roi_manager_modules.roi_types.VolCNMF method), 195
set_transmission() (mesmerize.plotting.HeatmapSplitterWidget method), 225
set_unit() (mesmerize.viewer.modules.femtonics_mesc.MEScNavigator method), 179
set_units() (mesmerize.viewer.modules.stimmap_modules.page.Page method), 199
set_update_live() (mesmerize.plotting.ScatterPlotWidget method), 231
set_widget() (mesmerize.plotting.DatapointTracerWidget method), 224
set_zlevel() (mesmerize.viewer.modules.roi_manager_modules.manager.ManagerVolROI method), 183
set_zlevel() (mesmerize.viewer.modules.roi_manager_modules.roi_types.VolCNMF method), 194
show_item_info() (mesmerize.viewer.modules.batch_manager.ModuleGUI method), 176
sig_channel_doubleclicked (mesmerize.viewer.modules.femtonics_mesc.MEScNavigator attribute), 178
sig_hpath_changed (mesmerize.viewer.modules.femtonics_mesc.MEScNavigator attribute), 178
sig_output_changed (mesmerize.plotting.KShapeWidget attribute), 228
sig_selection_changed (mesmerize.plotting.variants.Heatmap attribute), 227
SigmaMAD (class in mesmerize.pyqtgraphCore.flowchart.library.Signal), 218
signal_blocker() (mesmerize.plotting.widgets.base.BasePlotWidget class method), 223
signal_spot_clicked (mesmerize.plotting.variants.PgScatterPlot attribute), 232
slot_delete_roi_menu() (mesmerize.viewer.modules.roi_manager.ModuleGUI method), 180
slot_show_all_checkbox_clicked() (mesmerize.viewer.modules.roi_manager_modules.roi_list.ROIList method), 186
SpaceMap (class in mesmerize.pyqtgraphCore.flowchart.library.Display), 217
SpaceMapWidget (class in mesmerize.plotting), 233
SpliceArrays (class in mesmerize.pyqtgraphCore.flowchart.library.Data), 217
start_backend() (mesmerize.viewer.modules.roi_manager.ModuleGUI method), 180
start_manual_mode() (mesmerize.viewer.modules.roi_manager.ModuleGUI method), 180
start_process() (mesmerize.plotting.KShapeWidget method), 229
StaticDFoFo (class in mesmerize.pyqtgraphCore.flowchart.library.Biology), 219
stim_maps (mesmerize.viewer.core.ViewerWorkEnv attribute), 171
StimTuning (class in mesmerize.pyqtgraphCore.flowchart.library.Biology), 219

T

TextFilter (class in mesmerize.pyqtgraphCore.flowchart.library.Data), 217
Timeseries (class in mesmerize.pyqtgraphCore.flowchart.library.Display), 217
to_dict() (mesmerize.analysis.data_types.HistoryTrace method), 205
to_dict() (mesmerize.Transmission method), 202
to_hdf5() (mesmerize.analysis.cross_correlation.CC_Data method), 215
to_hdf5() (mesmerize.Transmission method), 202
to_json() (mesmerize.analysis.data_types.HistoryTrace method), 205
to_pandas() (mesmerize.viewer.core.ViewerWorkEnv method), 171
to_pickle() (mesmerize.analysis.data_types.HistoryTrace method), 205
to_pickle() (mesmerize.Transmission method), 202
to_pickle() (mesmerize.viewer.core.ViewerWorkEnv method), 171
to_state() (mesmerize.viewer.modules.roi_manager_modules.roi_types._
method), 188
to_state() (mesmerize.viewer.modules.roi_manager_modules.roi_types.E
method), 189
to_state() (mesmerize.viewer.modules.roi_manager_modules.roi_types.C
method), 197
to_state() (mesmerize.viewer.modules.roi_manager_modules.roi_types.M
method), 190
to_state() (mesmerize.viewer.modules.roi_manager_modules.roi_types.S
method), 192

to_state() (mesmerize.viewer.modules.roi_manager_modules.roi_types.VolCNMF method), 194
train_data (mesmerize.plotting.KShapeWidget property), 228
Transmission (class in mesmerize), 201
transmission (mesmerize.plotting.HeatmapTracerWidget property), 226
transmission (mesmerize.plotting.KShapeWidget property), 229
transmission (mesmerize.plotting.ProportionsWidget property), 230
transmission (mesmerize.plotting.ScatterPlotWidget property), 231
transmission (mesmerize.plotting.SpaceMapWidget property), 233
transmission (mesmerize.plotting.widgets.base._AbstractBasePlotWidget property), 222
transmission (mesmerize.plotting.widgets.base.BasePlotWidget property), 223
TVDiff (class in mesmerize.pyqtgraphCore.flowchart.library.Math), 219

## U

unit (mesmerize.viewer.modules.femtonics_mesc.MEScNavigator attribute), 179
units (mesmerize.viewer.modules.femtonics_mesc.MEScNavigator attribute), 179
update_idx_components() (mesmerize.viewer.modules.roi_manager_modules.managers.ManagerCNMFROI method), 184
update_idx_components() (mesmerize.viewer.modules.roi_manager_modules.managers.ManagerVolCNMF method), 184
update_plot() (mesmerize.plotting.HeatmapTracerWidget method), 226
update_plot() (mesmerize.plotting.ProportionsWidget method), 230
update_plot() (mesmerize.plotting.ScatterPlotWidget method), 231
update_plot() (mesmerize.plotting.SpaceMapWidget method), 233
update_plot() (mesmerize.plotting.widgets.base._AbstractBasePlotWidget method), 222
update_plot() (mesmerize.plotting.widgets.base.BasePlotWidget method), 223
update_roi_defs_from_configuration() (mesmerize.viewer.modules.roi_manager_modules.roi_list.ROIList method), 186
update_workEnv() (mesmerize.viewer.core.ViewerUtils method), 172
use_open_dir_dialog() (in module mesmerize.common.qdialogs), 169
use_open_file_dialog() (in module mesmerize.common.qdialogs), 168
use_save_file_dialog() (in module mesmerize.common.qdialogs), 168

## V

vi (mesmerize.viewer.modules.roi_manager_modules.roi_list.ROIList attribute), 185
viewer (mesmerize.viewer.core.ViewerUtils attribute), 172
ViewerUtils (class in mesmerize.viewer.core), 171
ViewerWorkEnv (class in mesmerize.viewer.core), 169
ViewHistory (class in mesmerize.pyqtgraphCore.flowchart.library.Data), 217
ViewTransmission (class in mesmerize.pyqtgraphCore.flowchart.library.Data), 217
VolCNMF (class in mesmerize.viewer.modules.roi_manager_modules.roi_types), 194

## W

WidgetRegistry (class in mesmerize.plotting.utils), 222
work_env (mesmerize.viewer.core.ViewerUtils attribute), 172

## X

XpowerY (class in mesmerize.pyqtgraphCore.flowchart.library.Math), 219

## Y

y_pred (mesmerize.plotting.KShapeWidget property), 228

## Z

ZScore (class in mesmerize.pyqtgraphCore.flowchart.library.Math), 219
