## Supplementary Figures and Tables for "Mesmerize: a dynamically adaptable user-friendly analysis platform for 2D & 3D calcium imaging data"

Supplementary Figure 1: pvc-7 (mouse) dataset analysis graph

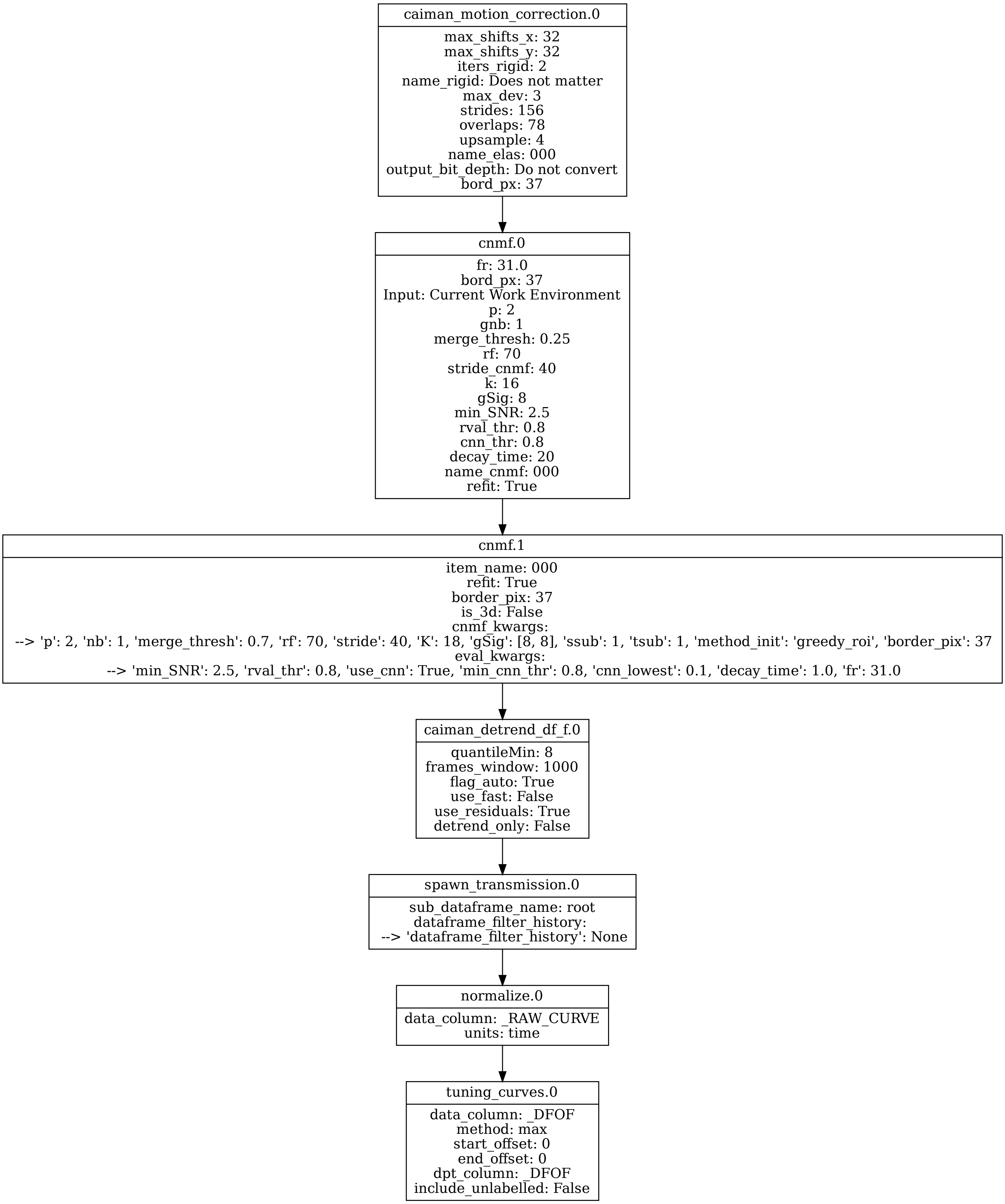

Supplementary Figure 2: k-means clustering of different cell types

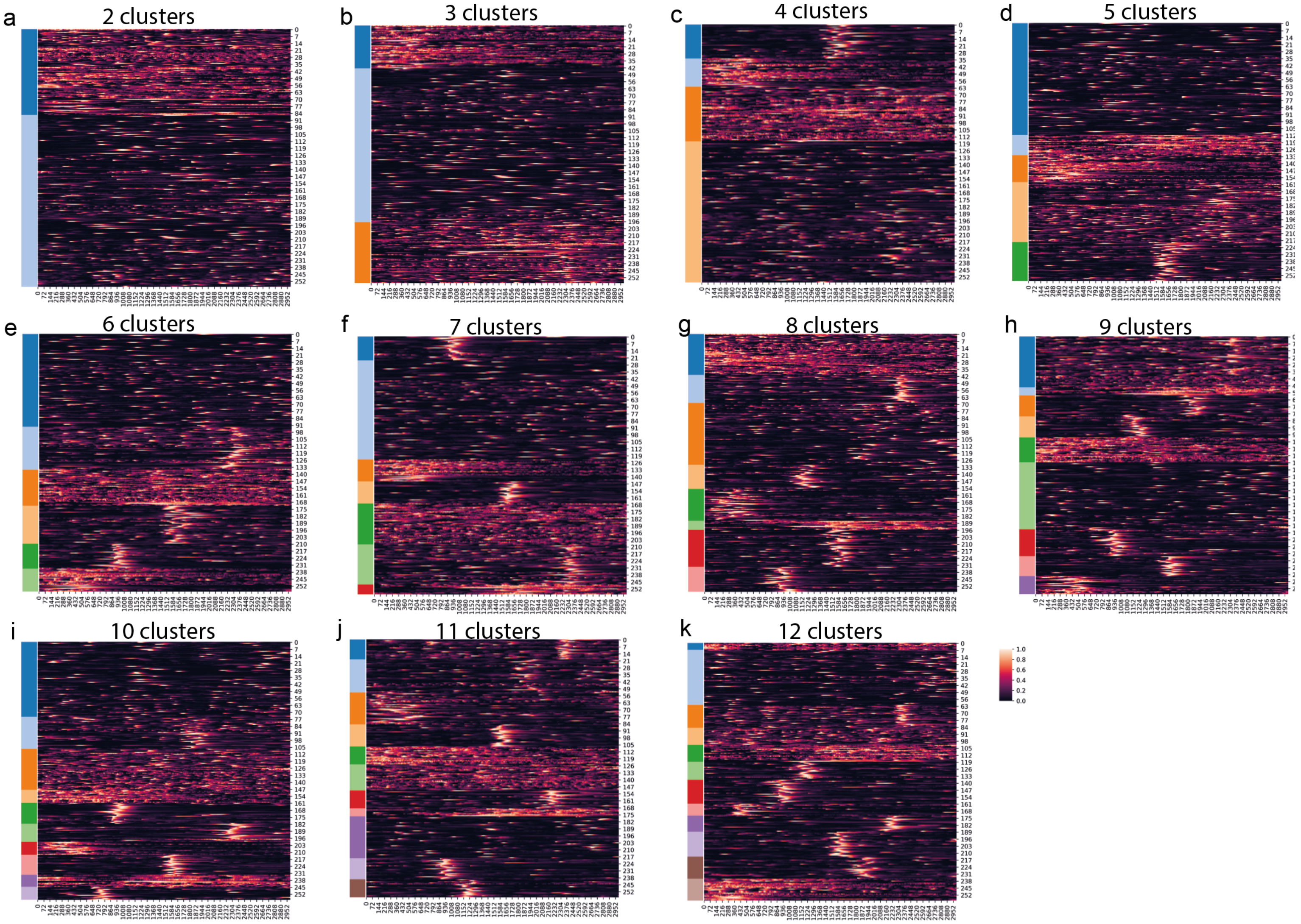

Supplementary Figure 3: Cell type identification in C. intestinalis

Example 1

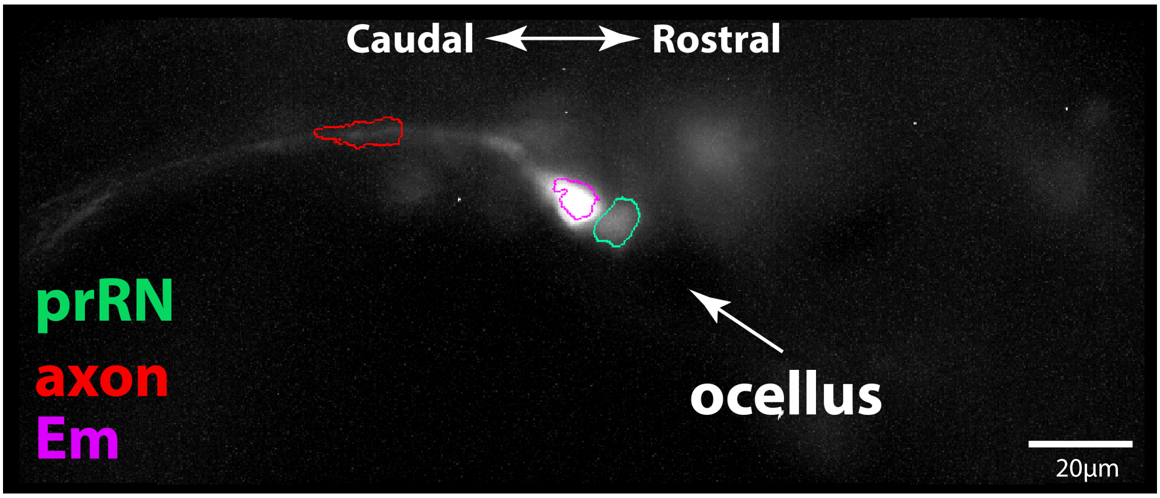


Example 2

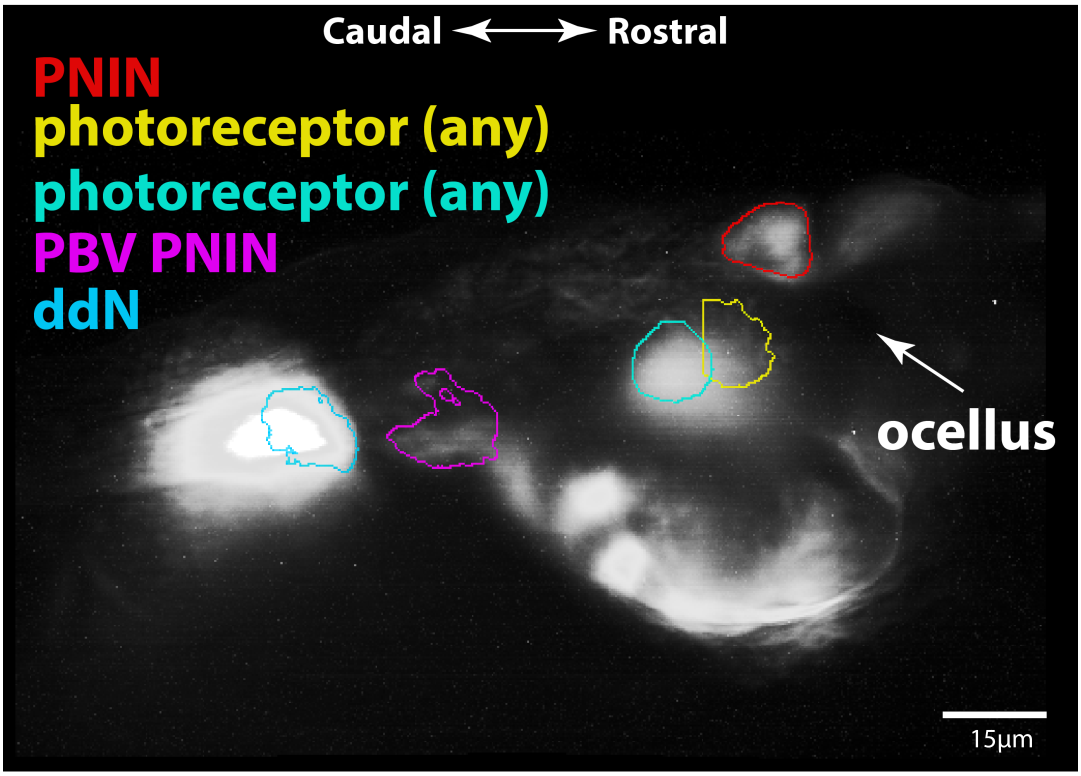


Example 3

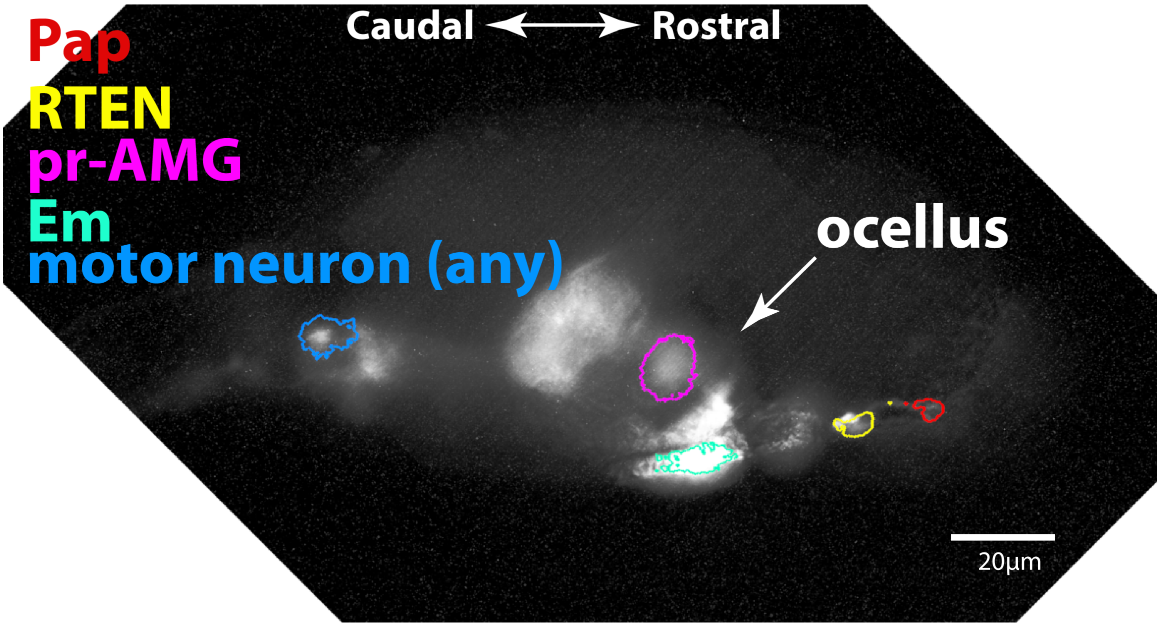

Supplementary  Figure 3: Cell type identification in *C. intestinalis*

Example 1:

**Em**

Eminens neurons are located near the dorsal surface, large and prominent with thick axons.

**prRN**

Photoreceptor relay neurons are located near the ocellus, and ventral to eminens neurons. Could also possibly be pr-AMG neuron or other photoceptor neuron

**axon**

axon of the Em neuron (excluded in analysis)

Example 2:

**PNIN**

Triangular cell body, near ocellus & near dorsal surface

**photoreceptor (any)**

neurons in general proximity to the ocellus

**PBV PNIN**

located near the neck, slightly ventral

**ddN**

located in the neck, rostral to other motor neurons

Example 3:

**Pap**

Palp neuron, this is an axon, the palp is out of plane

**RTEN**

Directly downstream of palp neurons

**pr-AMG**

near ocellus, below eminens, could possibly be another type of photoreceptor neuron

**Em**

Eminens neuron, large, close to dorsal surface, near ocellus

**motor neuron (any)**

Located in the neck region

Supplementary Figure 4: SOSD plot between the raw curves and interpolated Inverse Fourier Transforms (IFTs) of the DFTs with a step-wise increase in the frequency cut-off.

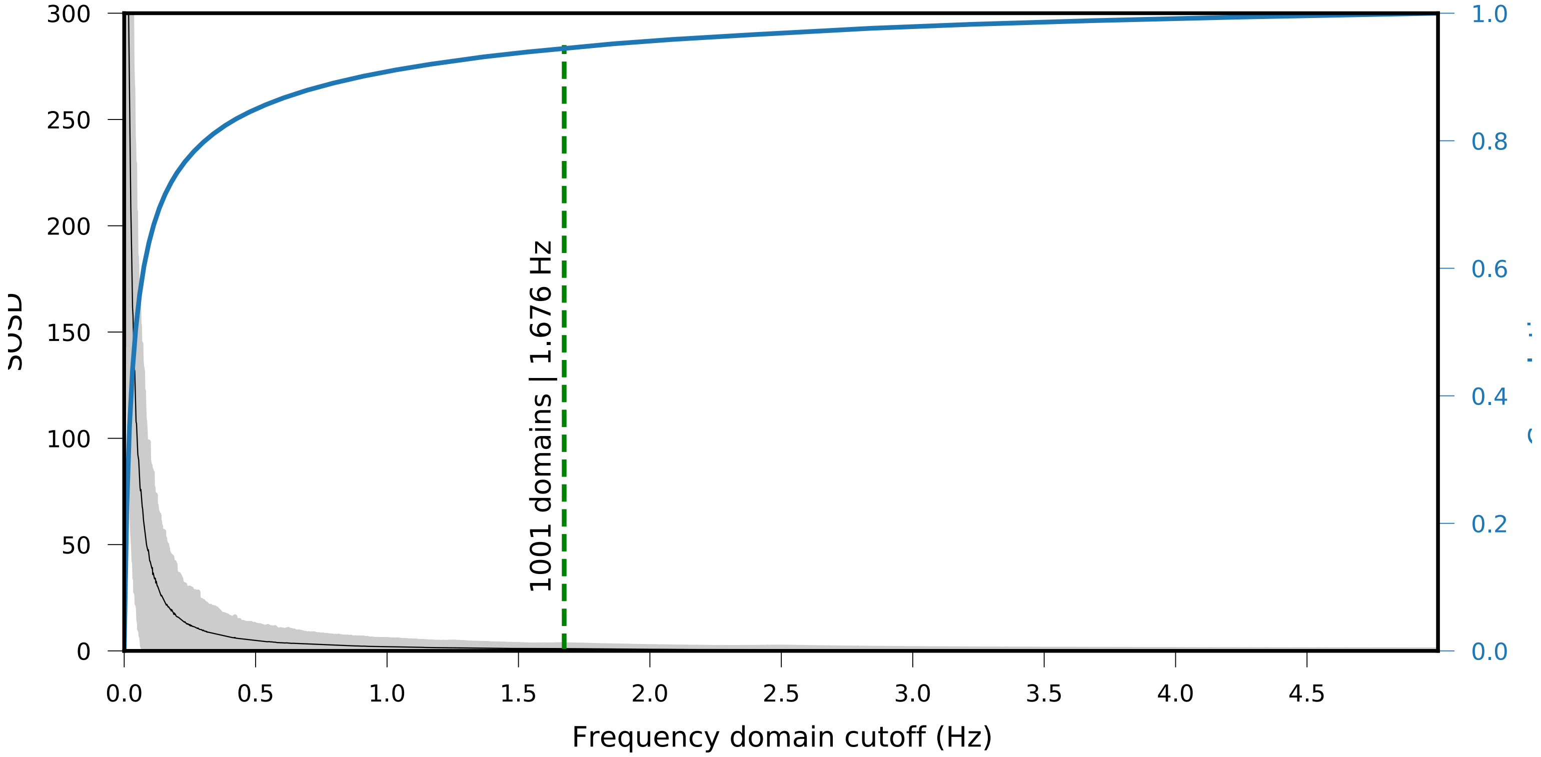

### Supplementary Figure 5: k-Shape clustering analysis graph

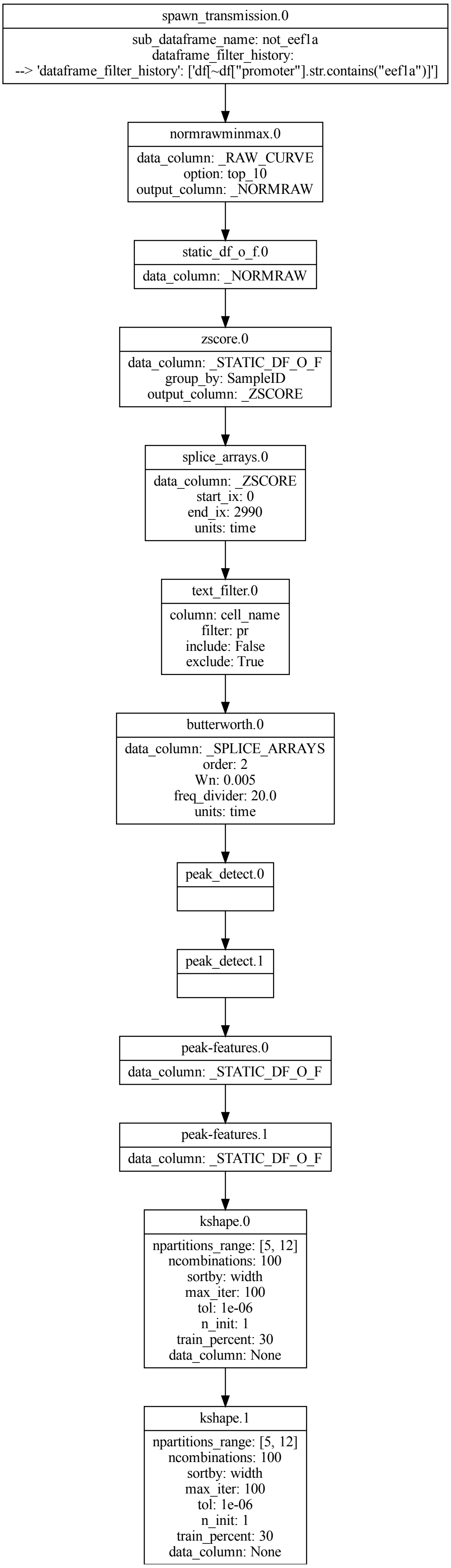

### Supplementary Figure 6: Zebrafish dataset analysis graph

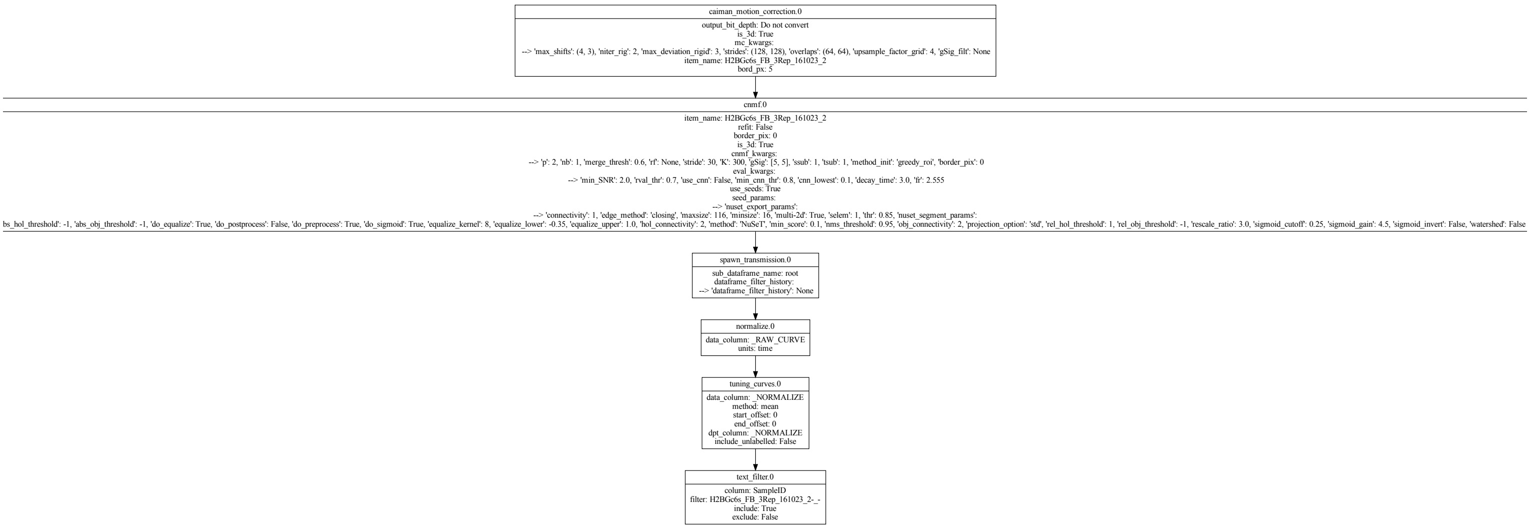

**Table 1: Number of animals and trials per promoter.**

| Promoter | Animals | Trials | Labelled cell types |
|---|---|---|---|
| pc2 | 7 | 7 | amg, atenp, cor_ass_bvin, dcen, eminens, mn, palp, pbv_pnin, pr, pr_amg, rten |
| brn3b | 10 | 12 | amg, antenal_relay, atena, atenp, ddn, eminens, palp, pr, pr_rn, rten |
| cesa | 8 | 11 | epidermis |
| cng_ch4 | 2 | 3 | pr |
| dmrt1 | 8 | 13 | atena, atenp, ddn, ependymal, palp, pbv_pnin, pnin, pr, pr_tract_interneuron, rten, vac_in |
| hnk1 | 5 | 5 | TLCs |
| pde9 | 9 | 15 | pr |
| eef1a | 10 | 15 | Ubiquitous expression |

**Table 2: Table describing cell types show in legend of Fig 5a**

| Abbreviation | Cell identity |
|---|---|
| AMG | Ascending motor ganglion peripheral interneurons |
| antRN | Antenna 1 relay neurons |
| aATEN | Anterior apical trunk epidermal neurons |
| pATEN | Posterior apical trunk epidermal neurons |
| CESA positive | CesA positive cell, epidermis cells. Labelled using the CesA promoter |
| cor-ass BVIN | Coronet associated ciliated brain vesicle interneurons |
| DCEN | Dorsal caudal epidermal neurons |
| ddN | Descending decussating neurons |
| Em | Eminens cell |
| ependymal | Ependymal cell |
| HNK-1 positive | HNK1 positive cell (TLCs). Labelled using the HNK1 promoter |
| motor neuron (any) | Motor Neuron. Exact pair not determined |
| Pap | Palp cell |
| PBV PNIN | Posterior brain vesicle peripheral interneurons |
| PNIN | Peripheral interneurons |
| photoreceptor (any) | Photoreceptor cell. Type not determined |
| pr-AMG | Photoreceptor-ascending motor ganglion neuron relay neurons |
| prRN | Photoreceptor relay neurons |
| trIN | Photoreceptor tract interneuron |
| RTEN | rostral trunk epidermal neurons |
| vacIN | Photoreceptor associated vacuolated neurons |
